# Key traits and genes associate with salinity tolerance independent from vigor in cultivated sunflower (*Helianthus annuus* L.)

**DOI:** 10.1101/2020.05.12.090837

**Authors:** Andries A. Temme, Kelly L. Kerr, Rishi R. Masalia, John M. Burke, Lisa A. Donovan

## Abstract

With rising food demands, crop production on salinized lands is increasingly necessary. Sunflower, a moderately salt tolerant crop, exhibits a trade-off where more vigorous, high-performing genotypes have a greater proportional decline in biomass under salinity stress. Prior research has found deviations from this relationship across genotypes; the magnitude and direction of these deviations provides a useful metric of tolerance. Here, we identified the traits and genomic regions underlying variation in this expectation-deviation tolerance. We grew a diversity panel under control and salt-stressed conditions and measured a suite of morphological (growth, allocation, plant and leaf morphology) and leaf ionomic traits. The genetic basis of variation in these traits and their plasticity was investigated via genome-wide association studies, which also enabled the identification of genomic regions (i.e., haplotypic blocks) influencing multiple traits. We found that the magnitude of plasticity in whole root mass fraction, fine root mass fraction, and chlorophyll content, as well as leaf Na and K content under saline conditions, were the traits most strongly correlated with expectation-deviation tolerance. Additionally, we identified multiple genomic regions underlying these traits as well as a single gene directly associated with this tolerance metric. Our results show that, by taking the vigor-salinity effect trade-off into account, we can identify unique traits and genes associated with salinity tolerance. Since these traits and genomic regions are distinct from those associated with high vigor (i.e., growth in benign conditions), they provide an avenue for increasing salinity tolerance in high-performing sunflower genotypes without compromising vigor.

**Single sentence summary:** Despite a trade-off between vigor and salinity-induced decline in biomass, distinct traits and genomic regions exist that could modulate this trade-off in cultivated sunflower.

## Introduction

The rapid rise of global population levels has increased strain on our food production systems (Ramankutty et al., 2018). With demands projected to nearly double by the middle of this century, expanded efforts to improve crop productivity are warranted. One factor limiting crop productivity is high soil salinity (generally NaCl) caused by poor irrigation practices, salt water encroachment, and/or drought. With over 20% of the world’s irrigated agricultural land being impacted by salinity and expansion needing to occur on less favorable lands, developing crop varieties more suitable for salinized soils is vital (Munns, 2005; Munns et al., 2020a)FAO 2005). However, mitigating the physiological problems imposed by high soil NaCl concentrations remains a challenging task (Munns et al., 2020a).

High soil salinity imposes two types of stress on plants. First, as a solute, dissolved NaCl imposes an osmotic stress which limits leaf expansion (Rawson and Munns, 1984) and photosynthesis through reduced transpiration, similar to drought (Munns, 2002; Munns and Tester, 2008). Second, either deliberately to combat the imposed osmotic stress or through unavoidable net leakage into the roots, plants take up NaCl from the soil (Munns et al., 2020a). Accumulated Na poses a risk of ion toxicity (Munns and Tester, 2008) and must either be excreted (a trait limited to salt-adapted species; (Cheeseman, 2015)) or sequestered in roots and stems (Cuin et al., 2011; Munns et al., 2012; Guan et al., 2014) or vacuoles (Mansour et al., 2003; Hasegawa, 2013; Bassil et al., 2019; Shabala et al., 2019). An improved understanding of the genetic basis of the traits underlying salt tolerance would accelerate the development of increasingly resilient cultivars (Zhu et al., 2016; Morton et al., 2018). Given the large set of traits and mechanisms related to salinity tolerance (Munns and Tester, 2008; Munns et al., 2020a; Munns et al., 2020b), however, the ability to maintain productivity under saline conditions is likely to be genetically complex (Flowers, 2004; Munns, 2005).

Numerous metrics for stress tolerance exist, leading to potentially conflicting conclusions depending on which metric is employed (Zhu et al., 2016; Morton et al., 2018). For example, a high performing genotype under saline conditions (suggesting ‘tolerance’) could also experience a large percentage performance decrease in salt stress (suggesting ‘sensitivity’). Furthermore, if there is a trade-off (or negative correlation; (Agrawal, 2020)) between performance and performance under stress (Mayrose et al., 2011; Koziol et al., 2012), then optimizing for yield under ideal conditions could favor genotypes that are more negatively impacted by stress. Conversely, optimizing for genotypes that are least impacted by stress could result in poor overall performance. An alternative approach is to decouple the metric of tolerance from performance (**Fig. 1**). Given an observed relationship between performance under benign conditions and the impact of stress across genotypes, the deviation of a given genotype from this overall relationship can be viewed as a measure of tolerance/sensitivity. In other words, if a genotype outperforms the expectation given its ideal performance, it exhibits evidence of stress tolerance relative to other genotypes. Similarly, if a genotype underperforms relative to the expectation, it exhibits evidence of stress sensitivity.

**Figure 1.**
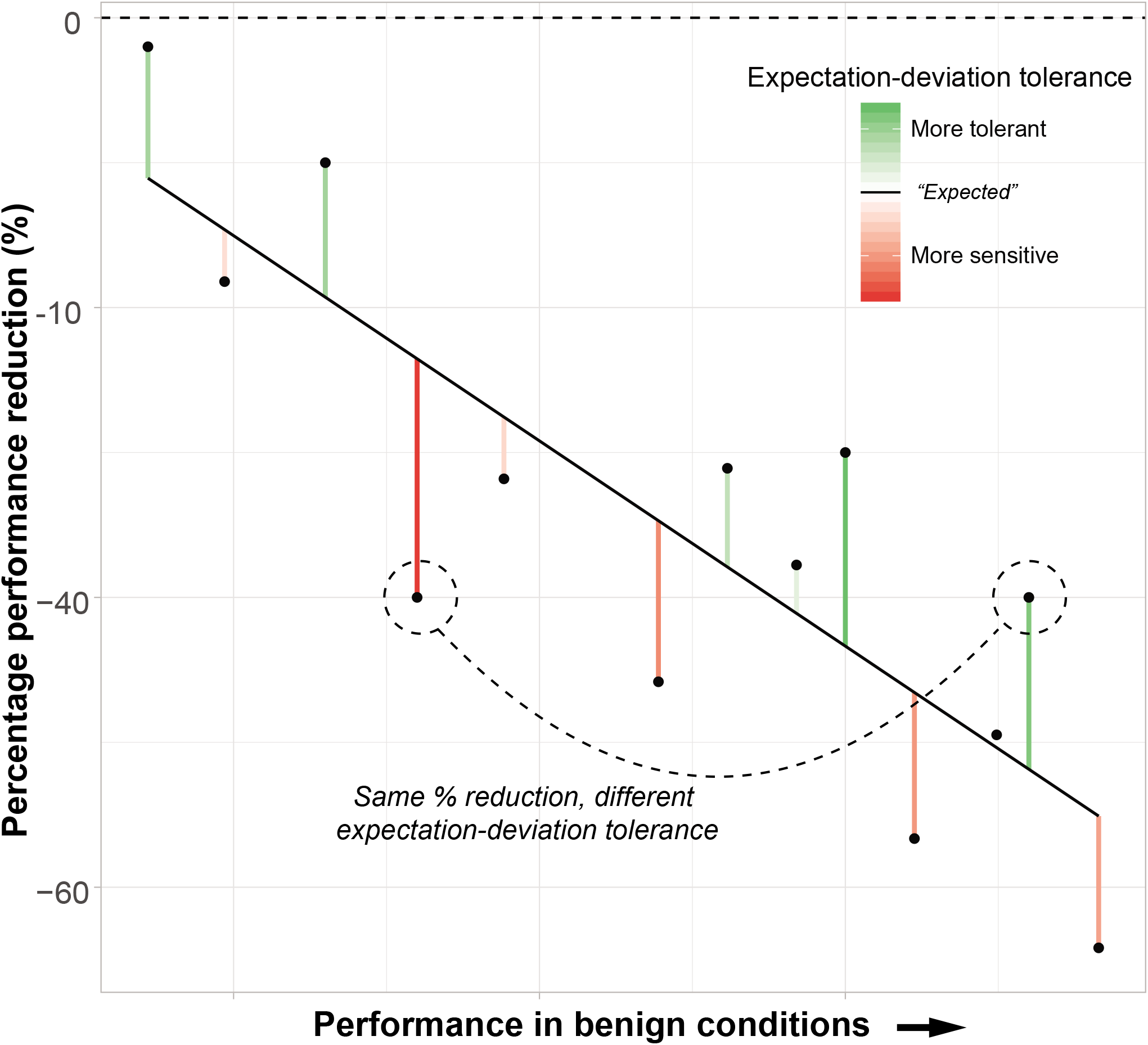
Conceptual diagram of expectation-deviation tolerance. Given a negative relationship between performance (estimated using any relevant measure) and the proportional effect of stress, along with variation in both, it can be difficult to define tolerance vs. sensitivity. The deviation of a genotype from the best-fit line provides an estimate of tolerance/sensitivity that is independent of differences in performance under benign conditions (see text for details).

In cultivated sunflower (*Helianthus annuus* L.), prior work has found a trade-off between vigor and the effect of salinity, with the more vigorous genotypes (higher growth under ideal conditions) exhibiting a greater decrease in biomass under stress (Temme et al., 2019a; Temme et al., 2019b). In these small sets of genotypes, it was possible to take this trade-off into account and quantify tolerance using a genotype’s deviation from the expected response (**Fig. 1**). This expectation-deviation tolerance was weakly correlated with Na sequestration in the stem and this differed from trait associations with a more common proportional-reduction tolerance metric (Temme et al., 2019a). By selecting on traits associated with better-than-expected performance, it could be possible to modulate this vigor/stress response tradeoff. A clearer picture of the traits underlying vigor as well as the extent to which multivariate trait expression is driven by genetic correlations (i.e., pleiotropy or close linkage; (Auge et al., 2019) will inform on the feasibility of this approach to modulate this negative relationship.

Given the central role of Na in salinity stress, extensive research has been done on its effect on plant processes (Mäser et al., 2002; Broadley et al., 2012; Hasegawa, 2013). Due to its ionic size, charge, and processes involved in sequestration, Na uptake affects the accumulation of other essential macro- and micronutrients. Most notably, K uptake is greatly affected when Na passes through K channels and disrupts the K:Na balance across membranes during Na sequestration (Shabala and Cuin, 2008; Shabala et al., 2019; Munns et al., 2020a). However, other elements are also likely to play a role in salinity tolerance. Decreases in the cost of inductively coupled plasma mass spectroscopy (ICP-MS) have enabled the measurement of a range of elements, collectively referred to as the ionome (Salt et al., 2008), and how they relate to salinity tolerance (Wu et al., 2018; Temme et al., 2019a; Temme et al., 2019b; Munns et al., 2020b).

In stressful environments, plants exhibit different trait values than under benign conditions. Explicit consideration of this trait plasticity has gained momentum as a useful tool in understanding trait variation across environments (Nicotra et al., 2010; Kusmec et al., 2017; Laitinen and Nikoloski, 2018; Arnold et al., 2019). Moreover, plasticity in some traits can be required for robustness in others (e.g., performance or yield; (Laitinen and Nikoloski, 2018). For example, in salt marshes, the success of invasive Japanese Knotweed (*Fallopia japonica*) is linked to plasticity in leaf succulence (Richards et al., 2008). Similarly, in cultivated sunflower, changes in leaf S content are associated with the maintenance of growth under salinity stress (Temme et al., 2019b). Examples such as these highlight the potential utility of trait plasticity for improving salt tolerance. The finding of distinct genomic regions underlying trait variation and plasticity in corn (Kusmec et al., 2017) and sunflower (Mangin et al., 2017) further suggests that trait expression can be decoupled from trait adjustment under stress.

Here we examine the suite of traits and genomic regions underlying salt tolerance, independent from vigor, in cultivated sunflower. Sunflower is a globally important oilseed crop that exhibits moderate, but genetically variable, salt tolerance (Katerji et al., 2000; Shi and Sheng, 2005; Temme et al., 2019a; Temme et al., 2019b). As demands for seed oil are expected to increase 70% by 2050 (Ramankutty et al., 2018), improvements need to be made in cultivars under production. Understanding the genetic basis of variation in the physiological mechanisms conferring tolerance to salinity stress will allow for rapid selection on these traits (Flexas and Gago, 2018; York, 2018). We use detailed phenotypic and ionomic characterization and genome-wide association studies (GWAS) to address the following questions: (1) What is the effect of salinity on morphological traits (vegetative growth, biomass allocation, plant and leaf morphology) and leaf ionomic traits in cultivated sunflower? (2) What is the link between trait expression, trait plasticity, and salinity expectation-deviation tolerance? (3) How heritable are the traits underlying expectation-deviation tolerance? (4) Which genomic regions influence vigor, expectation-deviation tolerance, and associated traits, and to what extent are these traits genetically correlated?

## Methods

### Plant materials and experimental design

This work used the sunflower association mapping (SAM) population (Mandel et al., 2011; Mandel et al., 2013). This population includes representatives from all four cultivated sunflower heterotic group/market-type combinations (HA-Oil, HA-NonOil, RHA-Oil, RHA-NonOil) as well as landraces, open-pollinated varieties, and cultivated genotypes carrying wild species introgressions. These lines were previously advanced via single-seed descent to minimize residual heterozygosity. All lines in this population have been subjected to whole-genome shotgun re-sequencing (Hübner et al., 2018), and these data were used to identify single-nucleotide polymorphisms (SNPs) and extract genotypic information for each line.

Due to greenhouse space constraints, we limited this study to the 239 genotypes from the SAM population with the least residual heterozygosity. With two treatments and four replicates/treatment (see below), this resulted in a total 1912 individual plants. The experiment was initiated by planting up to sixteen achenes from each genotype in seedling germination trays filled with a 3:1 mixture of coarse sand and turface (Turface Athletics, PROFILE Products, LLC, Buffalo Grove, IL). The trays were treated with a 0.45 g/L solution of a broad-spectrum fungicide to inhibit fungal growth (Banrot, Everris NA Inc., Dublin, OH) and watered twice daily for five days to allow for seedling emergence. At that point, eight representative seedlings per genotype were transplanted into 2.83 L pots filled with the same 3:1 sand/turface mixture plus 40 g Osmocote Plus (15-12-9 NPK; ScottsMiracle-Gro, Marysville, OH) and supplemental Ca^2+^ in the form of 5 ml each of gypsum (Performance Minerals Corporation, Birmingham, AL) and lime powder (Austinville Limestone, Austinville, VA). Pots were assigned to one of two treatments (i.e., control [0 mM NaCl] and salt-stressed [100 mM NaCl]) and arranged in a split-plot design with four replicate plots split between treatments.

Plots were arranged orthogonally to the direction of airflow within the greenhouse and alternated control and stress from east to west. Each subplot consisted of one pot per genotype randomly arranged. The differential treatments were implemented two days after transplanting with pots being placed in shallow ponds with the bottom 10 cm standing in either fresh water (control) or a 100 mM NaCl solution (stress). Three times per week, these ponds were drained and refilled with new solutions. At that time, all pots were top-watered with ~300 ml of the solution from their pond to stimulate nutrient release from the Osmocote pellets and to prevent buildup of NaCl at the soil surface in the stress treatment. After three weeks (while the plants were still in the vegetative growth stage), all plants were harvested over a four day period with one entire plot (478 plants, control and stress treatment) being harvested each day. This experiment was carried out during the summer of 2017 in the Botany Greenhouses at the University of Georgia (Athens, GA, USA).

### Phenotypic and leaf elemental traits

At harvest, all plants were measured for height (from the base of the stem to the apical meristem), stem diameter (at the base of the stem), and chlorophyll content (MC-100, Apogee Instruments, Inc., Logan, UT) of the most recently fully expanded leaf (MRFEL). Plant biomass was separated into the MRFEL pair, all other leaves, stem (plus immature flower bud, if present), and roots. The MRFELs (both leaves in the pair) were scanned at 300 DPI on a flatbed scanner (Canon CanoScan LiDE120) and leaf area was determined using Image J (NIH, USA, http://rsb.info.nih.gov/ij/). Prior to scanning, half of the lamina of one member of the MRFEL pair was separated and preserved for use in a separate study on leaf anatomy; the amount of missing biomass was therefore estimated using the mass/area relationship (i.e., leaf mass per area; LMA) from the remaining leaf lamina. All aboveground biomass samples were bagged and oven-dried at 60°C for 48 hrs. Pots containing roots were placed in cold storage to prevent degradation prior to processing. After harvest, over the course of two weeks, all roots were gently washed to remove soil, bagged, and oven dried at 60°C for 48 hrs. All dried samples were placed into storage outside the oven and then re-dried prior to any further analysis.

After drying, any immature flower buds that might have been present were separated from the stem tissue, and root tissue was separated into taproot (up to the point at which the taproot and lateral roots had similar widths) vs. lateral roots. After weighing all biomass fractions (lateral root, tap root, stem, leaves, MRFEL, buds), all intact MRFELs (excluding the petiole) for each genotype were pooled across replicates (separately by treatment) and ground to a coarse powder using a Wiley Mill (Thomas Scientific, Swedesboro, NJ). After mixing evenly, a 2 ml subsample of the coarse powder was ground to a fine powder using a Qiagen TissueLyser (Qiagen, Germantown, MD). Finely powdered leaf tissue was then sent to Midwest Laboratories (Omaha, NE) for ICP-MS to quantify the presence of the following elements: phosphorous (P), potassium (K), calcium (Ca), sodium (Na), sulfur (S), iron (Fe), zinc (Zn), copper (Cu), magnesium (Mg), manganese (Mn), and boron (B). Nitrogen (N) content was determined via the Dumas method.

### Data analysis

All statistical analyses were performed using R (v3.6.2 R Core Team 2015). Genotype mean values and the effects of salinity stress for all non-elemental traits were estimated by fitting a linear mixed model to the data with genotype and salinity treatment as fixed factors and pond within plot as a random factor using the R package *lme4* (Bates et al., 2015). Genotype means were calculated by taking the estimated marginal means of genotype x treatment without the random pond effect using *emmeans* (Lenth, 2018). Significance of the fixed effects and their interaction was determined via Wald’s Chi-square test using the package *car (Fox and Weisberg, 2011)*. The effect of the salinity treatment on leaf elemental content was tested using two-sample *t*-tests with genotype means as the unit of replication. All further analyses were carried out on the genotype means with genotype as the unit of replication. We quantified trait plasticity in response to salinity as the difference in natural log-transformed trait values between salt and control treatments. This plasticity metric has the benefit of being proportional to the trait value (in either the control or stressed treatments) and is symmetric to which treatment is set as the control, changing only in sign, not magnitude.

Correlational analyses were performed using *corrr* (https://github.com/tidymodels/corrr) with Spearman correlations and visualised using *ggplot2* (Wickham, 2009). Principal component analyses were performed on all size-independent traits, as well as all elemental traits with PCA biplots being visualised using a modified version of *ggbiplot* (https://github.com/vqv/ggbiplot).

Narrow sense heritability (*h*^2^) was calculated using the package *heritability* (Kruijer et al., 2015) and a kinship matrix constructed from the SNP data (see GWAS below). For all non-elemental traits, individual plants could be included in the heritability analysis. However, for the elemental traits and plasticity metric we only had genotype average values; this required us to use only genotypes mean values resulting in reduced precision in the heritability estimate.

### Genotypic data analysis and GWAS

Genome-wide association analyses were conducted using SNPs derived from whole-genome shotgun (WGS) resequencing of the SAM population (Hübner et al., 2018) using a SNP set developed by (Todesco et al., 2019). Briefly, this involved aligning the WGS reads to the XRQv1 reference genome (Badouin et al., 2017) using NextGenMap v0.5 (Sedlazeck et al., 2013), calling SNPs using GATK4 (Van der Auwera et al., 2013), and processing the resulting SNPs using GATK’s Variant Quality Score Recalibration (Poplin et al., 2017). This initial dataset was filtered to retain SNPs in the 90% tranche with < 10% missing data and a minor allele frequency ≥ 1%. Missing genotype calls were then imputed using BEAGLE v5.0 (Browning and Browning, 2016). This set of ca. 2.16M high quality SNPs was then re-ordered based on the more recent, improved assembly of the HA412-HO genome (Todesco et al., 2019) because it has better localized ordering of chromosomal segments. Finally, this dataset was filtered based on the 239 genotypes used in this study to remove SNPs with MAF < 5% and heterozygosity > 10%, resulting in ca 1.48M SNPs.

To better visualize the haplotypic structure present within the sunflower gene pool, and to estimate the effective number of independent markers in the genome, we used the ordered SNPs to construct a haplotype map based on observed patterns of linkage disequilibrium (LD) using PLINK v1.9 (Purcell et al., 2007). Haplotypic blocks were identified based on D’ following Gabriel et al. (2002) with a standard confidence interval of 0.7005 - 0.98, an informative fraction of 0.9 instead of 0.95 to be modestly more permissive of misaligned SNPs within blocks, and a maximum block span of 100Mb to accommodate the largest observed block in the genome (Gabriel et al., 2002) (Supplemental Fig. S1).

For all traits, association analyses were performed using GEMMA v0.98.1 (Zhou and Stephens, 2012). These analyses included corrections for both kinship (calculated using GEMMA) and population structure (using the first four PC axes from an analysis using a pruned set of ca. 24k independent SNPs [i.e., D’ < 0.8 using a sliding window approach] in the R package SNPRelate (Zheng et al., 2012). To correct our significance thresholds for multiple comparisons, we used a modified Bonferroni correction with the number of independent tests set to the number of multi-SNP haplotype blocks identified in our haplotypic analysis (see above). While this procedure helps to account for non-independence amongst linked markers, it still results in a highly conservative significance threshold since no genome assembly is perfect with respect to sequence ordering and orientation, thereby inflating the inferred number of independent regions.

Following the identification of significant marker-trait associations, associated genomic regions were identified on the basis of our haplotypic analysis. More specifically, significantly associated SNPs that occurred within a single haplotypic block were assumed to mark a single genomic region. To protect against the possibility of haplotype blocks being broken up by localized genotyping/ordering errors, thereby resulting in an overestimate of the number of independent regions influencing a given trait, we repeated the LD-based haplotypic analysis using only the markers on each chromosome that were found to be significant for one or more traits; if previously independent markers/haplotypes collapsed into a single haplotype in this re-analysis, they were assumed to mark the same region.

For each associated region, the magnitude of its effect on the trait of interest was set as the maximum β (i.e., effect) value across all SNPs in the region that were significantly associated with that trait. The relative effect size was calculated as (2\**β*)/(trait range) (Masalia et al., 2018)with the relative effect size of a haplotype block set as the maximum of the significant SNPs in that block. Relative effect size values were summed across all regions per trait to calculate the total phenotypic range that was explained by the significantly associated genomic regions.

In cases where SNPs within an associated region were significantly associated with multiple traits, we considered those traits to co-localize, either due to close linkage amongst the causal factors or through pleiotropy. Because of our highly stringent significance thresholds, false negatives are possible and could give the erroneous appearance of independence across traits. Thus, we also searched for cases of suggestive co-localization, defined as instances where SNPs within a haplotypic block were significantly associated with a given trait and in the top 0.1% of SNPs for one or more other traits, even if they did not reach statistical significance for those traits.

Following the identification of significantly associated regions for all traits of interest, a list of the genes contained within each region was extracted from the annotation of the HA412-HOv2 genome (Todesco et al., 2019) based on the positions of the first and last SNPs in each region. When singleton SNPs exhibited significant associations, we extracted either the gene in which it was contained or the two flanking genes if it was in an intergenic region. Code for the GWAS pipeline including the trait colocalization steps and gene extraction is available online (at https://github.com/aatemme/Sunflower-GWAS-v2).

## Results

### Phenotypes, salinity stress, and tolerance

Across all genotypes, salinity stress greatly affected growth, morphology, and the leaf ionome. While there was substantial variation among genotypes, the median response to salt stress across genotypes was a 56.3% reduction in biomass, and substantially greater biomass allocation to roots (50.4% increase in RMF) at the expense of stem allocation (35.2% decrease in SMF). Leaf mass per area increased by 46.4%, indicating thicker or denser leaves. Changes to the leaf ionome were substantial, with a median increase of 101.4% in Mn content and 4395.2% in Na content (**Table 1, Supplemental Fig. S2).** For all growth and morphological traits (i.e., all non-elemental traits) analyzed, genotypes (G) differed significantly in their trait values (*P* < 0.001) as well as in their response to salinity (genotype-by-treatment [GxT] interaction; *P* < 0.001). Due to the presence of this interaction, the main effect of treatment was non-significant for chlorophyll content, and several root traits. This indicates substantial spread in the response to salinity with some genotypes increasing in these trait values and others decreasing.

**Table 1.** Effect of salt stress on plant traits. Traits measured and their median value (and range) at control (0 mM NaCl), salt stress (100 mM NaCl), and the plasticity between treatments. Plasticity was calculated as the difference in natural log transformed values (control-salt) but converted here to Δ% (via e^Δln(trait)^ - 1) change from control for ease of interpretation. For all non-elemental traits stars indicate Wald’s Chi-square significance of the main effects of genotype (G), treatment (T), and their interaction (GxT). As tissue was bulked for elemental content stars indicate t-test significance between control and salt stressed groups. ns: non-significant, *: *P* < 0.05, **: *P* < 0.01, ***: *P* < 0.001.

| Trait | 0 mM NaCl | 100 mM NaCl | Plasticity ($\Delta\%$) | T | G | GxT |
| --- | --- | --- | --- | --- | --- | --- |
| *Plant mass (g)* | 4.31 (1.01 - 10.65) | 1.86 (0.43 - 4.82) | -56.3 (-85.29 - 47.7) | ** | *** | *** |
| *Leaf mass (g)* | 2.2 (0.62 - 5.54) | 0.98 (0.27 - 2.48) | -53.1 (-83.89 - 43.36) | * | *** | *** |
| *LMF ($g_{\text{leaves}} g_{\text{plant}}^{-1}$)* | 0.5 (0.4 - 0.73) | 0.54 (0.39 - 0.68) | 7 (-14.62 - 42.75) | * | *** | *** |
| *LMA ($g m^{-2}$)* | 22.62 (14.12 - 32.69) | 32.87 (26.91 - 41.65) | 46.4 (7.12 - 123.92) | *** | *** | *** |
| *Height (cm)* | 57.75 (9.5 - 80.63) | 23.88 (6.63 - 43.38) | -57.7 (-77.2 - -21.24) | *** | *** | *** |
| *Stem mass (g)* | 1.52 (0.05 - 3.9) | 0.42 (0.06 - 1.35) | -70.9 (-94.92 - 4.15) | *** | *** | *** |
| *SMF ($g_{\text{stem}} g_{\text{plant}}^{-1}$)* | 0.36 (0.11 - 0.45) | 0.23 (0.1 - 0.35) | -35.2 (-61.36 - -4.4) | *** | *** | *** |
| *Stem diameter (mm)* | 8.7 (5.2 - 14.61) | 5.12 (2.71 - 7.55) | -39.9 (-65.56 - -8.82) | *** | *** | *** |
| *Shoot mass (g)* | 3.78 (0.8 - 9.45) | 1.48 (0.33 - 3.83) | -59.5 (-85.65 - 27.85) | ** | *** | *** |
| *All root mass (g)* | 0.56 (0.12 - 1.57) | 0.36 (0.03 - 0.89) | -36.7 (-94.47 - 242.13) | . | *** | *** |
| *Tap root mass (g)* | 0.23 (0.05 - 0.51) | 0.14 (0.02 - 0.35) | -36.3 (-91.21 - 150.67) | ns | *** | *** |
| *Lateral root mass (g)* | 0.33 (0.07 - 1.13) | 0.21 (0.01 - 0.69) | -35.6 (-97.34 - 298.95) | ns | *** | *** |
| *RMF ($g_{\text{roots}} g_{\text{plant}}^{-1}$)* | 0.13 (0.08 - 0.22) | 0.2 (0.11 - 0.28) | 50.4 (-34.79 - 134.02) | ns | *** | *** |
| *Tap root MF ($g_{\text{tap root}} g_{\text{plant}}^{-1}$)* | 0.05 (0.02 - 0.14) | 0.08 (0.03 - 0.17) | 49.5 (-56.86 - 229.86) | * | *** | *** |
| *Tap root RF ($g_{\text{tap root}} g_{\text{roots}}^{-1}$)* | 0.41 (0.2 - 0.71) | 0.41 (0.17 - 0.63) | -1.6 (-56.97 - 156.03) | ns | *** | *** |
| *Fine root MF ($g_{\text{fine root}} g_{\text{plant}}^{-1}$)* | 0.08 (0.04 - 0.14) | 0.12 (0.06 - 0.22) | 52.4 (-28.88 - 194.57) | ns | *** | *** |
| *Fine root RF ($g_{\text{fine root}} g_{\text{roots}}^{-1}$)* | 0.59 (0.29 - 0.8) | 0.59 (0.37 - 0.83) | 1 (-38.43 - 46.88) | ns | *** | *** |
| *Chlorophyll content (index)* | 14.71 (8.86 - 26.15) | 17.79 (5.64 - 32.27) | 20.3 (-47.6 - 127.69) | ns | *** | *** |
| *Boron (ppm)* | 100 (55 - 162) | 109 (52 - 213) | 9.3 (-38.82 - 98.77) | *** |  |  |
| *Calcium (%)* | 1.23 (0.79 - 2.17) | 1.28 (0.7 - 2.42) | 1.4 (-47.18 - 130.48) | ns |  |  |
| *Copper (ppm)* | 27 (15 - 49) | 36 (21 - 65) | 37.5 (-23.4 - 125) | *** |  |  |
| *Iron (ppm)* | 139 (95 - 535) | 127.5 (74 - 934) | -9.1 (-64.11 - 661.67) | ** |  |  |
| *Magnesium (%)* | 0.41 (0.26 - 0.64) | 0.41 (0.23 - 0.76) | -2.5 (-41.51 - 94.87) | ns |  |  |
| *Manganese (ppm)* | 670 (350 - 1363) | 1358.5 (429 - 3663) | 101.4 (-2.51 - 439.47) | *** |  |  |
| *Nitrogen (%)* | 7.19 (5.94 - 8.72) | 5.97 (4.65 - 7.1) | -16.2 (-34.94 - 3.24) | *** |  |  |
| *Phosphorus (%)* | 0.44 (0.25 - 0.84) | 0.3 (0.17 - 0.46) | -30.6 (-65.48 - 20) | *** |  |  |
| *Potassium (%)* | 5.54 (3.93 - 7.6) | 5 (2.15 - 7) | -9.3 (-52.51 - 41.34) | *** |  |  |
| *Sodium (%)* | 0.02 (0.01 - 0.1) | 1.14 (0.13 - 4.66) | 4395.2 (288.33 - 48916.67) | *** |  |  |
| *Sulfur (%)* | 0.71 (0.52 - 1.54) | 0.52 (0.36 - 0.95) | -27.5 (-52.13 - 3.28) | *** |  |  |
| *Zinc (ppm)* | 84 (47 - 238) | 108 (55 - 471) | 27.5 (-60.14 - 324.32) | *** |  |  |
| *Potassium/sodium ratio* | 225.2 (60.93 - 973.33) | 4.41 (0.67 - 49.06) | -98 (-99.87 - -75.19) | *** |  |  |

Due to our bulking of leaf tissue across replicates for each genotype, we could not contrast genotypes for elemental content. However, by using genotypes as the unit of replication, we could still assess the impact of salinity on the diversity panel as a whole. Of the twelve elements analyzed, all but Ca and Mg were significantly affected by salinity. This lack of overall effect for these two elements is likely due to the large variance in response with content halving in some genotypes and doubling in others. Under salt stress, foliar concentrations significantly increased for B (median 9.3%), Cu (median 37.5%), Mn (median 101.4%), Na (median 4395.2%), and Zn (median 27.5%). In contrast, foliar concentrations decreased for Fe (median 9.1%), N (median 16.2%), P (median 30.6%), K (median 9.3%), and S (median 27.5%) (**Table 1, Supplemental Fig. S2).**

While all but one genotype decreased in biomass accumulation under salinity stress, there was a strong positive relationship between biomass in control conditions and biomass in saline conditions. More vigorous genotypes (i.e., higher biomass in control conditions) tended to accumulate the most biomass under salt stressed conditions as well (*P* < 0.001, *R*^2^=0.49) **(Fig. 2a).** However, these more vigorous genotypes also tended to have a greater proportional decrease in biomass under saline conditions (*P* < 0.001, *R*^2^=0.23) (**Fig. 2b)**. Given this trade-off between vigor and the effect of salinity, we quantified tolerance as the deviation of each genotype from this expected response. Thus, residuals from the fitted line (**Fig. 2b**) served as estimates of these expectation-deviation tolerance values (**Fig. 1**).

**Figure 2.**
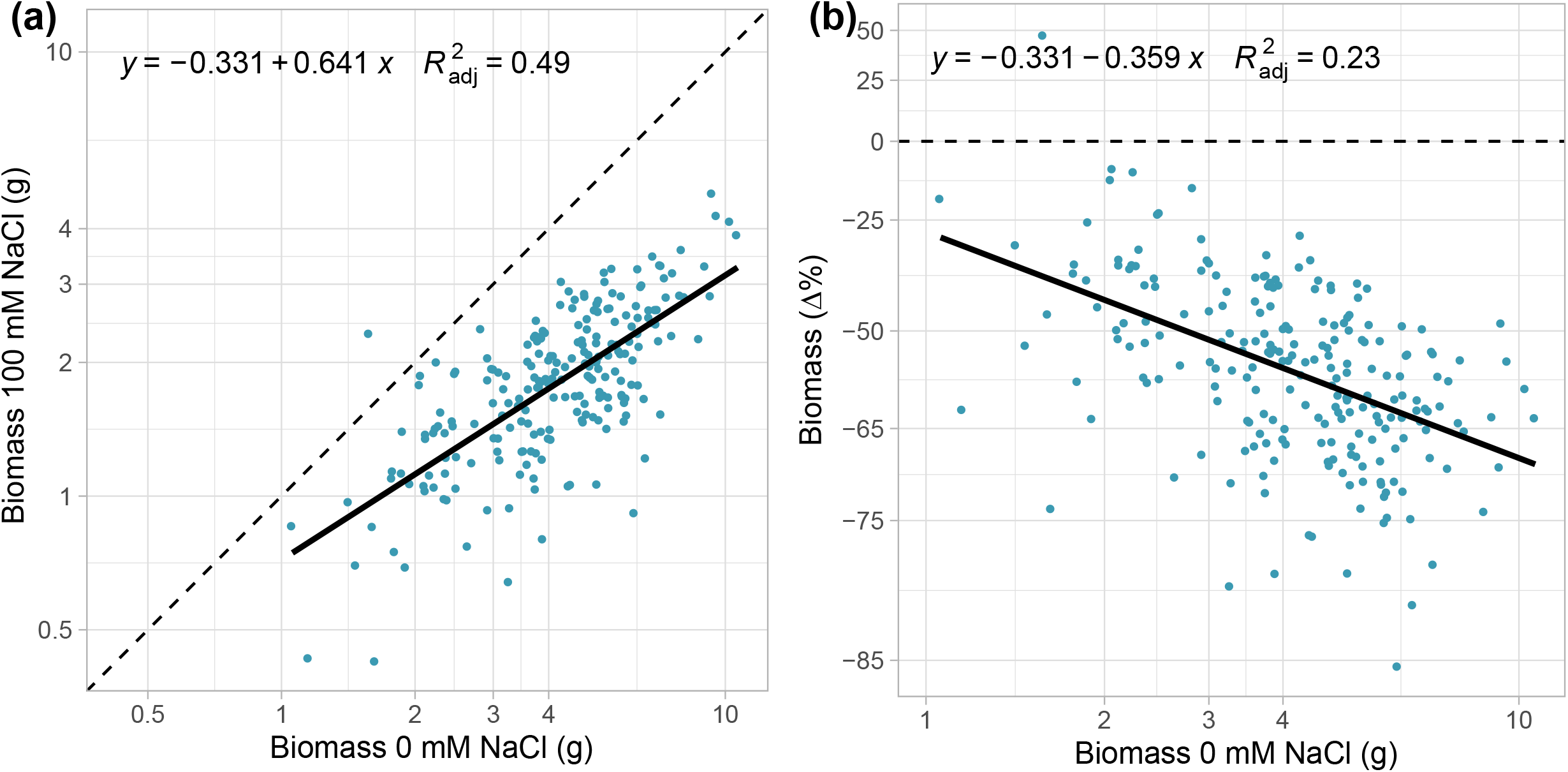
Effects of salt stress on whole-plant biomass. (a) Whole-plant biomass under control (0 mM NaCl) and salt stressed (100 mM NaCl) conditions. Dotted line indicates 1:1 and solid line fitted regression. (b) Plasticity in biomass under salt stress versus biomass at control conditions. Plasticity was calculated as the difference in natural log transformed values (control-salt) but converted here to Δ% (via e^Δln(trait)^-1) change from control for ease of interpretation. Dotted line indicates zero difference (no salt effect), solid line fitted regression. Note the natural log scaling of the axis ticks.

Numerous significant trait correlations were observed in both the control and salt-stressed environments (**Fig. 3a**). Some notable relationships in the control treatment were that vigor (biomass) was negatively correlated with overall root mass fraction (ρ = −0.27, *P* < 0.001) but positively correlated with the proportion of root mass made up by fine roots (ρ = 0.36, *P* < 0.001). While vigor was negatively correlated with leaf N (ρ = −0.26, *P* < 0.001) and P (ρ = −0.49, *P* < 0.001), it was positively correlated with Mg (ρ = 0.18, *P* < 0.05) and Mn (ρ = 0.38, *P* < 0.001). The trait correlations under saline conditions differed from those of the control treatment. For example, biomass was not correlated with Mg concentration, but was positively correlated with S (ρ = 0.146, *P* < 0.05), and negatively correlated with Na (ρ = −0.261, *P* < 0.001). Additionally, under saline conditions, Na was negatively correlated with K (ρ = −0.520, *P* < 0.001), S (ρ = −0.310, *P* < 0.001), and N (ρ = −0.199, *P* < 0.01), but positively correlated with P (ρ = 0.316, *P* < 0.001). All traits were positively correlated between treatments, though there was substantial variation in correlation strength across traits.

**Figure 3.**
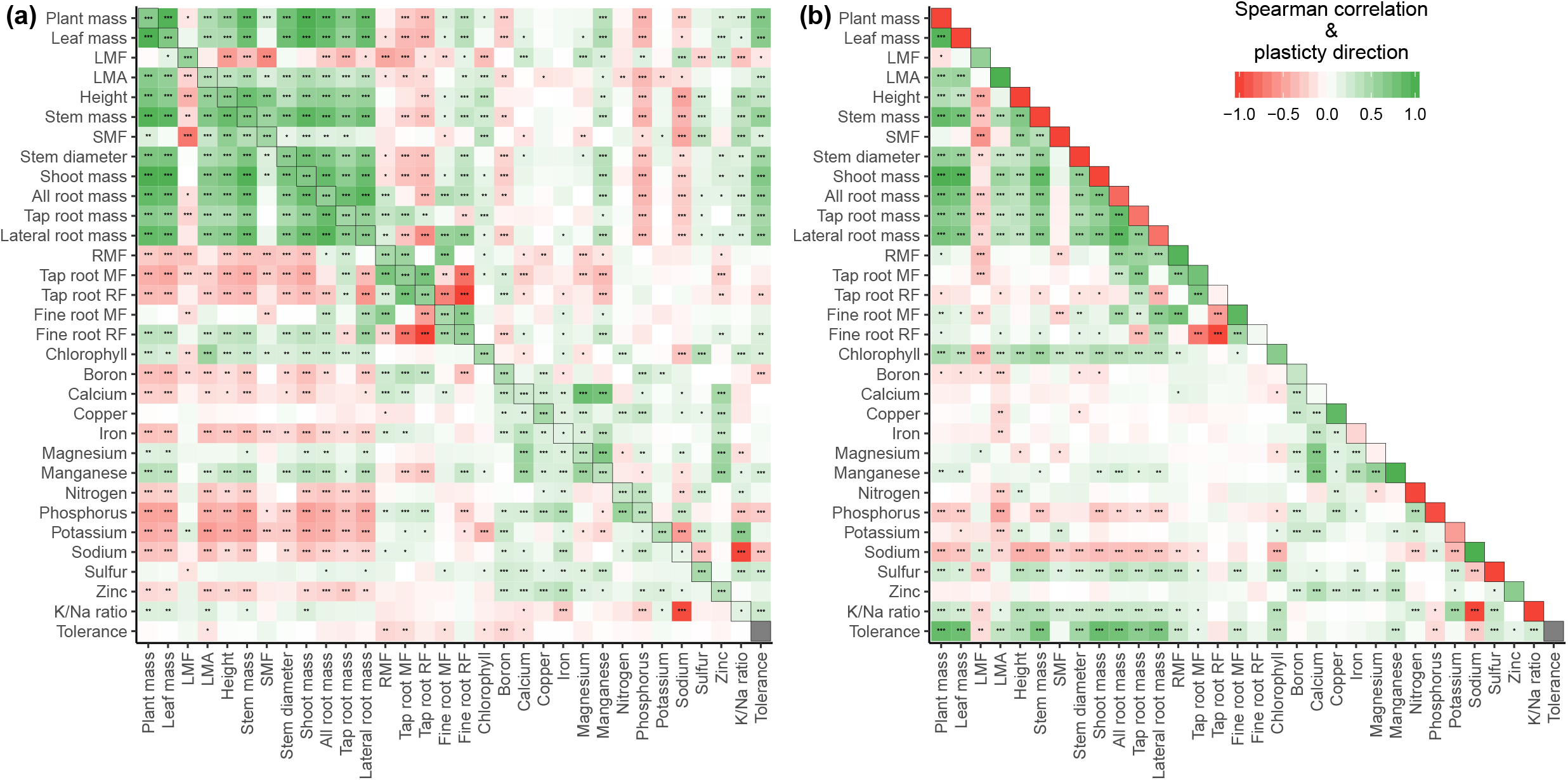
Trait correlations. (a) Spearman correlation matrix between all phenotypic and elemental traits under control (lower diagonal) and salt-stressed conditions (upper diagonal). Correlation within traits between the control and stressed conditions is on the diagonal. (b) Spearman correlation matrix of plasticity in trait value, Δln(trait), between all phenotypic and elemental traits. The average direction and uniformity of the shift in trait value due to salinity stress is on the diagonal (−1 = all genotypes decrease in trait value, +1= all genotypes increase in trait value). As tolerance is a compound trait based on both control and salt treatment, comparison across treatments does not apply. Correlation strengths are between −1 (red) and 1 (green). Stars in tiles indicate significance level of correlation. *: *P* < 0.05, **: *P* < 0.01, ***: *P* < 0.001.

Across genotypes, plasticity in response to salinity was also highly correlated among traits **(Fig. 3b**). In interpreting the sign of the correlation between trait shifts, it is important to note the sign of the change for each of the two correlated traits. For example, a positive correlation could result from two traits that both increase or both decrease with salinity but, perhaps counter-intuitively, also between one trait that increases and another that decreases with salinity when a smaller decrease in one is coupled with a larger increase in the other. Changes in biomass traits were all positively correlated, such that a greater decrease in any biomass component in response to salinity was coupled with a greater decrease in other components. However, in terms of biomass allocation changes, the leaf, stem, and root mass fractions were all negatively correlated indicating variability in how genotypes adjust their biomass allocation. For leaf elemental content, changes in leaf elemental concentration for Mg, Mn, and Ca were positively correlated. Sodium accumulation was correlated with multiple other elements, such that genotypes with a smaller increase in Na under salt stress had a smaller decrease in N (ρ = −0.21, *P* < 0.001), S (ρ = −0.29, *P* < 0.001), and K (ρ = −0.36, *P* < 0.001). For whole-plant biomass, genotypes with a greater proportional increase in leaf Na content had a greater decrease in biomass accumulation (ρ = −0.30, *P* < 0.001).

Given the substantial correlation among traits within treatments and among proportional changes across treatments, a multivariate approach incorporating trait correlations was used to provide an integrated view of the relationship of traits with both vigor and tolerance **(Table 3)**. Using principal component analysis (PCA), high vigor under benign conditions was associated most strongly with the first principal component of putatively size-independent traits (i.e., biomass ratios, LMA, and chlorophyll content; *P* < 0.001, *R^2^* = 0.27) and the second principal component of the leaf ionome (*P* < 0.001, *R*^2^ = 0.22). The top traits loading on these principal components were variation in root mass allocation for the putatively size-independent PCA and for the leaf element PCA a suite of concentrations including Mn, Mg, P, and N **(Table 4, Supplemental Fig. S3)**.

**Table 2.** Heritability and genome-wide association of traits. Narrow sense heritability (*h*^2^), number of independent regions with significant genome wide associations, and the sum of relative effect sizes (RES), (2\**β*)/(trait range), of those regions under benign (0 mM NaCl), salt stressed conditions (100 mM NaCl), and the shift in traits between treatments (plasticity). Narrow sense heritability was calculated using individual plants for non-elemental traits and genotype means for elemental traits and trait plasticity values. *h*^2^ confidence interval in parentheses. Relative effect sizes (%) as the sum of RES across all regions. RES > 100 likely indicates an overestimation of the number of independent regions, potentially due to errors in the underlying genome assembly.

| Trait | 0 mM NaCl |  |  | 100 mM NaCl |  |  | Plasticity |  |  |
| --- | --- | --- | --- | --- | --- | --- | --- | --- | --- |
| | $h^2$ | # Reg. | RES | $h^2$ | # Reg. | RES | $h^2$ | # Reg. | RES |
| *Plant mass (g)* | 0.47 (0.38-0.56) | 6 | 84.6 | 0.44 (0.35-0.54) | 1 | 15.3 | 0.01 (-0.03-0.05) | 1 | 10.7 |
| *Leaf mass (g)* | 0.44 (0.35-0.54) | 4 | 52.7 | 0.44 (0.35-0.53) | 3 | 56.8 | 0.01 (-0.04-0.06) |  |  |
| *LMF (g leaves g plant<sup>-1</sup>)* | 0.45 (0.37-0.53) | 15 | 310.1 | 0.32 (0.23-0.41) | 1 | 22 | 0.08 (0-0.15) |  |  |
| *LMA (g m<sup>-2</sup>)* | 0.28 (0.19-0.37) | 1 | 11.7 | 0.25 (0.16-0.34) | 2 | 49.7 | 0.03 (-0.02-0.07) | 3 | 55.8 |
| *Height (cm)* | 0.54 (0.47-0.62) | 5 | 100.8 | 0.64 (0.57-0.7) | 2 | 43 | 0.13 (0.03-0.24) |  |  |
| *Stem mass (g)* | 0.55 (0.47-0.63) | 3 | 41.2 | 0.5 (0.42-0.58) | 3 | 68.5 | 0.09 (-0.01-0.18) | 1 | 14.3 |
| *SMF (g stem g plant<sup>-1</sup>)* | 0.53 (0.45-0.6) | 6 | 125.8 | 0.49 (0.4-0.57) | 2 | 39.1 | 0.05 (-0.01-0.11) | 2 | 34.1 |
| *Stem diameter (mm)* | 0.45 (0.36-0.54) | 1 | 20.9 | 0.3 (0.21-0.38) | 2 | 29.8 | 0.11 (0.01-0.2) |  |  |
| *Shoot mass (g)* | 0.49 (0.4-0.58) | 3 | 41.3 | 0.47 (0.38-0.56) |  |  | 0.01 (-0.03-0.05) | 2 | 26.2 |
| *All root mass (g)* | 0.33 (0.23-0.42) | 4 | 69.7 | 0.33 (0.24-0.42) | 1 | 15.9 | 0.04 (-0.02-0.1) | 4 | 43.2 |
| *Tap root mass (g)* | 0.3 (0.21-0.39) |  |  | 0.3 (0.21-0.39) | 2 | 27.4 | 0.07 (-0.01-0.15) | 4 | 59.9 |
| *Lateral root mass (g)* | 0.35 (0.26-0.44) | 5 | 83.6 | 0.31 (0.22-0.39) |  |  | 0.03 (-0.02-0.08) | 4 | 38.5 |
| *RMF (g roots g plant<sup>-1</sup>)* | 0.19 (0.12-0.27) |  |  | 0.14 (0.07-0.21) | 1 | 17.7 | 0.06 (0-0.12) |  |  |
| *Tap root MF (g tap root g plant<sup>-1</sup>)* | 0.07 (0.04-0.11) | 1 | 16.3 | 0.09 (0.04-0.14) |  |  | 0.09 (0-0.17) |  |  |
| *Tap root RF (g tap root g roots<sup>-1</sup>)* | 0.37 (0.29-0.45) |  |  | 0.28 (0.19-0.37) |  |  | 0.04 (-0.02-0.09) | 1 | 10 |
| *Fine root MF (g fine root g plant<sup>-1</sup>)* | 0.07 (0.02-0.12) | 1 | 19 | 0.12 (0.05-0.19) | 4 | 72.3 | 0.03 (-0.01-0.07) |  |  |
| *Fine root RF (g fine root g roots<sup>-1</sup>)* | 0.37 (0.29-0.45) |  |  | 0.28 (0.19-0.37) |  |  | 0.03 (-0.02-0.08) | 1 | 14.5 |
| *Chlorophyll content (index)* | 0.59 (0.52-0.67) | 4 | 116.3 | 0.54 (0.46-0.62) | 2 | 36.6 | 0.12 (0.03-0.21) |  |  |
| *Boron (ppm)* | 0.29 (0.13-0.45) |  |  | 0.18 (0.05-0.31) | 4 | 84.1 | 0.09 (0.01-0.18) |  |  |
| *Calcium (%)* | 0.12 (0.03-0.22) | 1 | 10.3 | 0.18 (0.07-0.3) | 2 | 46.7 | 0.23 (0.1-0.35) | 2 | 50.8 |
| *Copper (ppm)* | 0.19 (0.07-0.31) | 3 | 85.7 | 0.23 (0.1-0.37) | 2 | 48.2 | 0.14 (0.03-0.25) |  |  |
| *Iron (ppm)* | 0.02 (-0.02-0.07) | 19 | 294.9 | 0.03 (-0.01-0.07) | 3 | 59.7 | 0.04 (0-0.09) |  |  |
| *Magnesium (%)* | 0.25 (0.1-0.4) |  |  | 0.28 (0.13-0.43) | 2 | 51 | 0.31 (0.15-0.46) | 1 | 18.6 |
| *Manganese (ppm)* | 0.07 (-0.01-0.15) |  |  | 0.15 (0.06-0.25) | 4 | 65.9 | 0.13 (0.03-0.22) | 6 | 104.4 |
| *Nitrogen (%)* | 0.16 (0.04-0.27) |  |  | 0.12 (0.02-0.23) | 1 | 24 | 0.07 (-0.01-0.16) | 3 | 48.1 |
| *Phosphorus (%)* | 0.11 (0.02-0.19) |  |  | 0.1 (0.02-0.18) | 1 | 15.6 | 0.01 (-0.02-0.04) |  |  |
| *Potassium (%)* | 0.11 (0.01-0.2) | 1 | 31.3 | 0.28 (0.14-0.43) | 2 | 49.6 | 0.25 (0.09-0.41) | 1 | 11.9 |
| *Sodium (%)* | 0.5 (0.4-0.6) | 1 | 21.6 | 0.22 (0.09-0.35) | 6 | 133.5 | 0.16 (0.05-0.28) | 2 | 42.2 |
| *Sulfur (%)* | 0.58 (0.39-0.77) | 3 | 55.9 | 0.38 (0.2-0.55) | 4 | 68.5 | 0.19 (0.06-0.33) | 2 | 29.8 |
| *Zinc (ppm)* | 0.01 (-0.03-0.05) | 1 | 14.5 | 0.06 (0.02-0.11) | 8 | 76.6 | 0.06 (0-0.11) | 4 | 81.8 |
| *Potassium/sodium ratio* | 0.01 (-0.03-0.05) | 2 | 48.5 | 0.17 (0.06-0.27) | 20 | 416.2 | 0.18 (0.06-0.3) | 3 | 56.8 |
| *Tolerance* | - | - | - | 0.05 (-0.01-0.11) | 1 | 19.9 | - | - | - |

**Table 3.** Relationship between principal components and plant vigor under benign conditions and salinity tolerance. *R*^2^ of the significant ordinary least-squares regression of PC1 and PC2 of the size-independent traits (chlorophyll content, fine root allocation (mass fraction and root fraction), LMA, LMF, RMF, SMF, tap root allocation (mass fraction and root fraction)) and elemental traits (B, Ca, Cu, Fe, Mg, Mn, N, P, K, Na, S, Zn, K/Na ratio) in control and salt stressed treatments and the plasticity in trait values between both environments with vigor (biomass at control conditions) and tolerance (deviation from the expected decline). Highlighted are the top two regressions with the highest explanatory power. *: *P* < 0.05, **: *P* < 0.01, ***: *P* < 0.001.

| Trait set | Treatment | PC axis | Vigor<br>$R^2_{adj.}$ | Tolerance<br>$R^2_{adj.}$ |
| --- | --- | --- | --- | --- |
| Size-independent traits | Control | PC1 | **0.27** *** | 0.02* |
|  |  | PC2 | 0.03** | - |
|  | Salt | PC1 | 0.08*** | 0.02* |
|  |  | PC2 | - | 0.04** |
|  | Plasticity | PC1 | 0.02* | - |
|  |  | PC2 | - | **0.17** *** |
| Elements | Control | PC1 | 0.1*** | - |
|  |  | PC2 | **0.22** *** | - |
|  | Salt | PC1 | 0.02* | - |
|  |  | PC2 | - | **0.14** *** |
|  | Plasticity | PC1 | 0.06*** | 0.04** |
|  |  | PC2 | - | 0.04** |

**Table 4.**
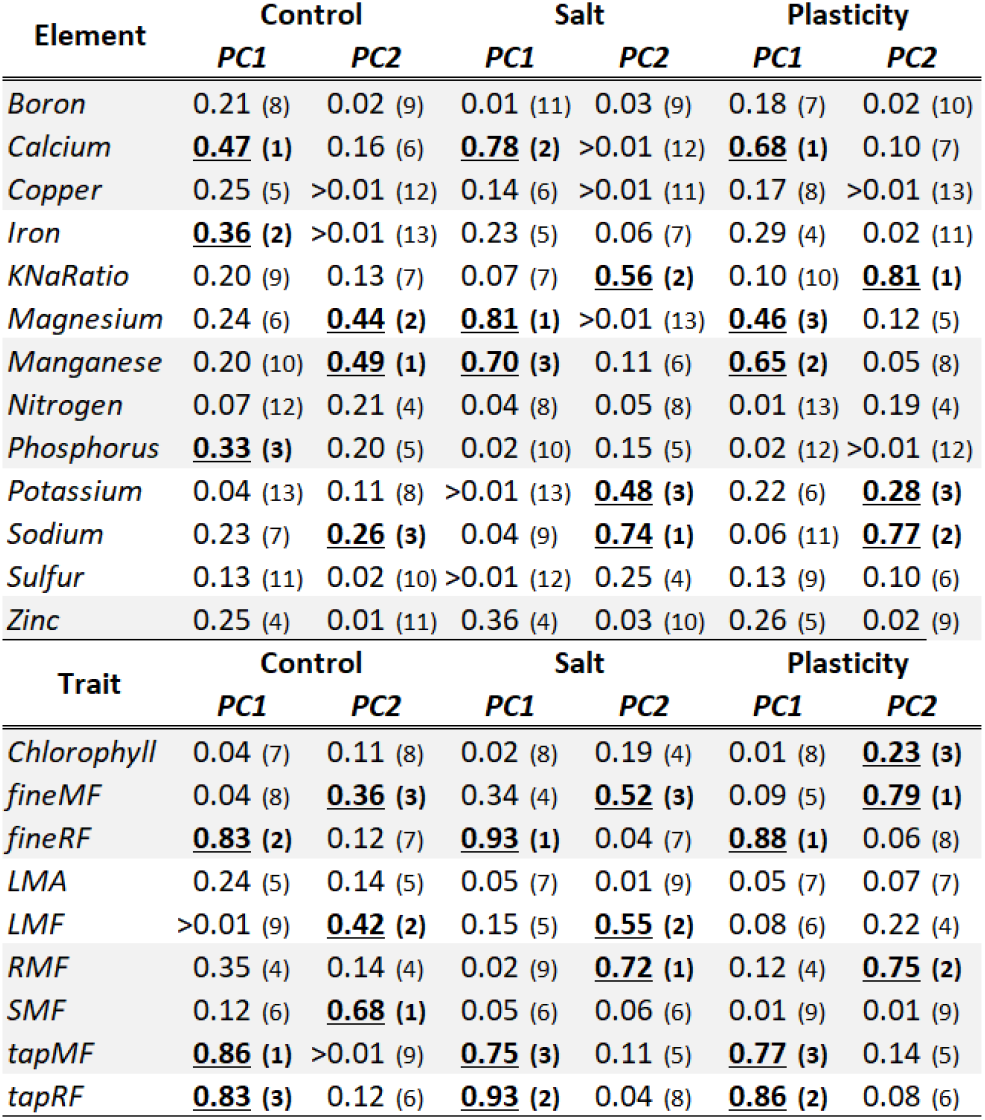
Principal component loadings of size-independent traits and elements. Loadings (fraction of variance in trait explained by principal component) on the PC1 and PC2 in control, salt stressed and the plasticity in trait values between both treatments. Highlighted are the top three size independent traits/elements per PC. Rank of loading in parentheses.

| Element | Control |  | Salt |  | Plasticity |  |
| --- | --- | --- | --- | --- | --- | --- |
|  | PC1 | PC2 | PC1 | PC2 | PC1 | PC2 |
| Boron | 0.21 (8) | 0.02 (9) | 0.01 (11) | 0.03 (9) | 0.18 (7) | 0.02 (10) |
| Calcium | **0.47** (1) | 0.16 (6) | **0.78** (2) | >0.01 (12) | **0.68** (1) | 0.10 (7) |
| Copper | 0.25 (5) | >0.01 (12) | 0.14 (6) | >0.01 (11) | 0.17 (8) | >0.01 (13) |
| Iron | **0.36** (2) | >0.01 (13) | 0.23 (5) | 0.06 (7) | 0.29 (4) | 0.02 (11) |
| KNaRatio | 0.20 (9) | 0.13 (7) | 0.07 (7) | **0.56** (2) | 0.10 (10) | **0.81** (1) |
| Magnesium | 0.24 (6) | **0.44** (2) | **0.81** (1) | >0.01 (13) | **0.46** (3) | 0.12 (5) |
| Manganese | 0.20 (10) | **0.49** (1) | **0.70** (3) | 0.11 (6) | **0.65** (2) | 0.05 (8) |
| Nitrogen | 0.07 (12) | 0.21 (4) | 0.04 (8) | 0.05 (8) | 0.01 (13) | 0.19 (4) |
| Phosphorus | **0.33** (3) | 0.20 (5) | 0.02 (10) | 0.15 (5) | 0.02 (12) | >0.01 (12) |
| Potassium | 0.04 (13) | 0.11 (8) | >0.01 (13) | **0.48** (3) | 0.22 (6) | **0.28** (3) |
| Sodium | 0.23 (7) | **0.26** (3) | 0.04 (9) | **0.74** (1) | 0.06 (11) | **0.77** (2) |
| Sulfur | 0.13 (11) | 0.02 (10) | >0.01 (12) | 0.25 (4) | 0.13 (9) | 0.10 (6) |
| Zinc | 0.25 (4) | 0.01 (11) | 0.36 (4) | 0.03 (10) | 0.26 (5) | 0.02 (9) |
| Trait | Control |  | Salt |  | Plasticity |  |
|  | PC1 | PC2 | PC1 | PC2 | PC1 | PC2 |
| Chlorophyll | 0.04 (7) | 0.11 (8) | 0.02 (8) | 0.19 (4) | 0.01 (8) | **0.23** (3) |
| fineMF | 0.04 (8) | **0.36** (3) | 0.34 (4) | **0.52** (3) | 0.09 (5) | **0.79** (1) |
| fineRF | **0.83** (2) | 0.12 (7) | **0.93** (1) | 0.04 (7) | **0.88** (1) | 0.06 (8) |
| LMA | 0.24 (5) | 0.14 (5) | 0.05 (7) | 0.01 (9) | 0.05 (7) | 0.07 (7) |
| LMF | >0.01 (9) | **0.42** (2) | 0.15 (5) | **0.55** (2) | 0.08 (6) | 0.22 (4) |
| RMF | 0.35 (4) | 0.14 (4) | 0.02 (9) | **0.72** (1) | 0.12 (4) | **0.75** (2) |
| SMF | 0.12 (6) | **0.68** (1) | 0.05 (6) | 0.06 (6) | 0.01 (9) | 0.01 (9) |
| tapMF | **0.86** (1) | >0.01 (9) | **0.75** (3) | 0.11 (5) | **0.77** (3) | 0.14 (5) |
| tapRF | **0.83** (3) | 0.12 (6) | **0.93** (2) | 0.04 (8) | **0.86** (2) | 0.08 (6) |

Tolerance to salinity, estimated as the residual from the fitted vigor vs. effect-of-salinity trade-off (**Fig. 2b)** – i.e expectation-deviation tolerance – was most strongly associated with principal components than differed from those associated with vigor. Expectation-deviation tolerance was most strongly associated with the second principal component of the PCA of plasticity in size-independent traits (*P* < 0.001, *R*^2^ = 0.17; **Fig. 4a,b)** and the second principal component of the leaf ionomic PCA under salinity stress (*P* < 0.001, *R*^2^ = 0.14, **Fig. 4c,d)**. The top traits loading onto these principal components were, respectively, plasticity in: root mass fraction, fine root mass fraction, and chlorophyll content; and Na content, K content, and K/Na ratio under saline conditions **(Table 4, Fig. 4a,c)**. Thus, genotypes that had a greater increase in root mass and fine root mass fraction as well as a lower Na content, and higher K/Na ratio under saline conditions had a greater estimated tolerance, resulting in a lower than expected (based on their vigor) decrease in performance.

**Figure 4.**
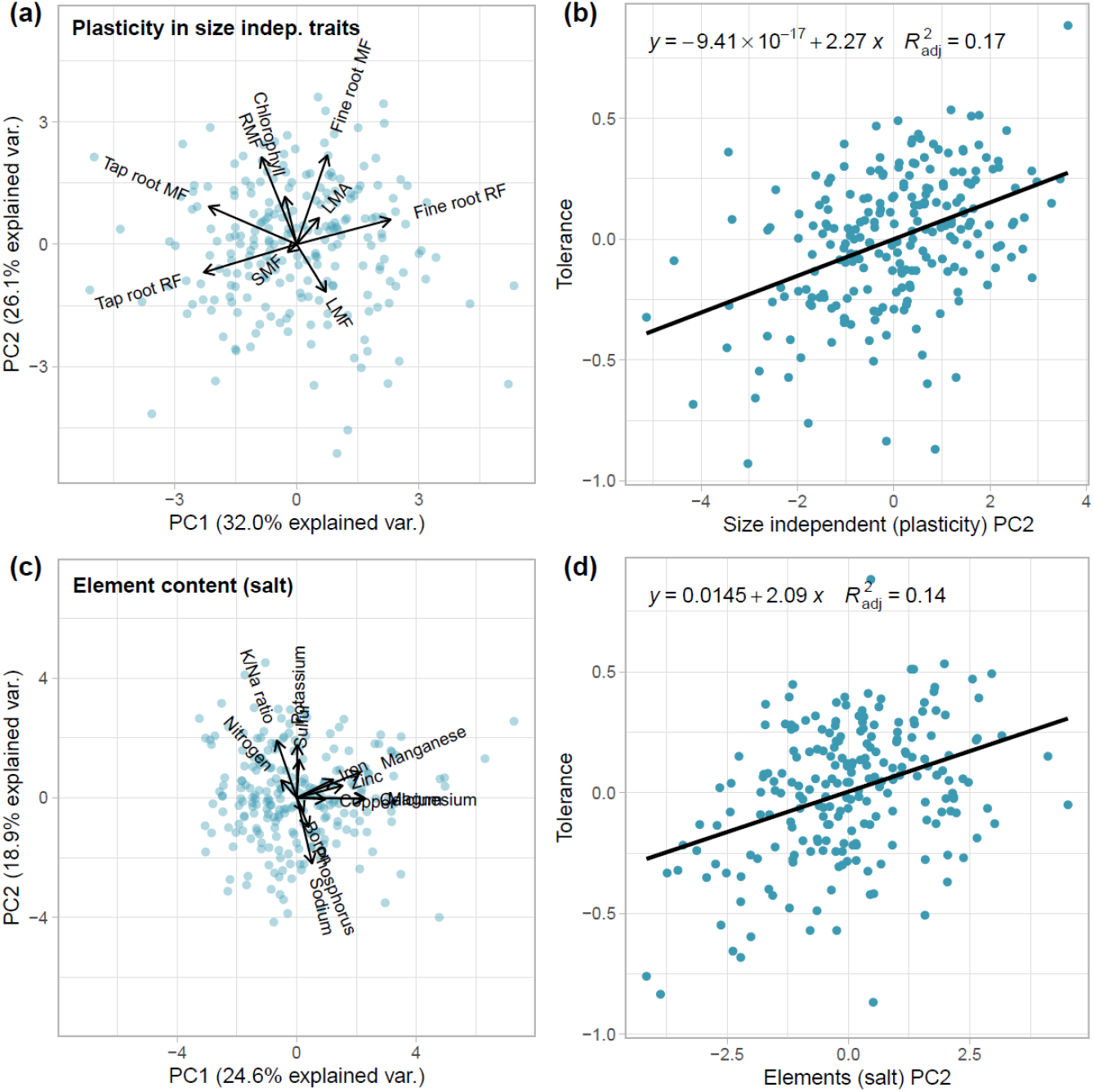
Salinity tolerance and associated trait complexes. Tolerance to salt stress (defined as the deviation of the expected decrease in biomass based on vigor) to the plastic adjustments in biomass independent traits (a & b) and to leaf elemental composition (c & d). Points indicate genotypes, whereas arrows represent eigenvectors of traits.

### Heritability and GWAS

Estimates of *h*^2^ revealed a broad range of additive genetic variance across traits in control and salt treatment with only modest differences between treatments (**Table 2**). A notable exception was that *h*^2^ of leaf Na concentration decreased from 0.5 to 0.22 under salt stress. It should be noted however that for the ionomic traits heritability values were based on genotypes’ bulked average with associated greater uncertainty in heritability estimate. For the growth and morphological traits, *h*^2^ was generally high with whole-plant biomass having an estimate of 0.47 in control conditions and 0.44 under saline conditions. Estimates for above ground biomass traits were greater than for belowground traits. Expectation-deviation tolerance had low *h*^2^ at just 0.05, although it should be noted again that this was necessarily estimated using genotype averages.

The haplotype block analysis resulted in the identification of 19,918 haplotypic blocks with multiple SNPs in strong LD along with 9,179 singletons (**Supplemental Fig. S1**). We identified 2,739 SNPs significantly associated with variation in one or more traits in the control and salt stress treatments, or with variation in the proportional shift between treatments. Based on the haploytpe map, these significant SNPs represented 242 haplotype blocks plus 17 singletons **(Supplemental Fig. S1, S4)**. Further clustering of these significant SNPs allowed for the combining of haplotype blocks (likely due to localized mis-ordering in the underlying genome assembly) resulting in 169 genomic regions (158 haplotypic blocks and 11 singletons) significantly associated with variation in trait values (**Supplemental Fig. S5**). Of these 169 regions, 29 were significantly associated with variation in multiple trait and/or treatment combinations (up to 11 for a single region spanning 93,336 kbp on chromosome 10) and 143 regions were significant for at least one trait and suggestive (top 0.1% of SNPs) for at least one other. Relative effect sizes of regions ranged from fairly high, 36% for chlorophyll content in control conditions at region 04-09, to fairly modest, 6% for zinc content under salt treatment for region 10-02 (**Supplemental Table S3, Supplemental Fig. S6**). Only 26 regions had a significant association with a single trait, but not even a suggestive association with others **(Supplemental Table S1, Supplemental Fig. S6).** Across all traits, only four instances were identified where a region was significantly associated with a trait in more than one of either control conditions, the salt treatment, or the plasticity between them (**Supplemental Table S1**). This suggests strong GxT interaction, but could also result from the stringent multiple comparison correction.

Surprisingly, many highly correlated traits did not share significant regions (**Supplemental Fig. S6**). For example, Na concentration, K concentration and leaf K/Na ratio shared no significant regions for control and salt treatment. However, suggestive regions did overlap, highlighting the issue of stringent multiple comparison correction. Within traits, across environments and plasticity, there were few significant shared regions as well (**Supplemental Fig. S7**) Although suggestive regions for control and salt treatment did overlap more frequently. Regions associated with trait plasticity were generally more distinct with even fewer significant and suggestive regions overlapping with control and salt treatment.

Based on the haplotype map, genome annotation, and GWAS results, we determined the genes contained in each significant region. When a region consisted of multiple haplotype blocks that could be combined after a second LD analysis (**Supplemental Fig. S4)**, we combined the genes contained in each individual haplotype block. Across the 169 genomic regions, we identified 4,167 genes potentially associated with variation in traits and trait plasticity (**Supplemental Table S2, Supplemental Fig. S8).** Of these genes, 1,586 were of unknown function, but the remaining 2,753 genes consisted of 720 unique proteins and 397 that were present two or more times (**Supplemental Table S2**). Contained in the gene list are several candidates that are of interest due to potential involvement with salt tolerance mechanism based on the literature.

For salinity expectation-deviation tolerance we identified a single, small, region on chromosome 16 with a relative effect size of 20%. This region (16-01) contained only a single putative alpha-mannosidase gene. While individuals carrying the major vs. minor allele at this locus did not differ in vigor (**Fig. 6b**), genotypes carrying the minor allele had a greater than expected (based on their vigor) decline in biomass under salt stress (**Fig. 6a**). It should be noted, however, that with over 38% genes of unknown function, and 135 regions containing multiple genes, identifying the causal genes underlying trait variation will be challenging.

## Discussion

Improving food security requires the development of crops that can withstand environmental stresses such as high salinity. Given the osmotic as well as ionic challenges presented by salt stress, the physiological and genetic basis of salinity tolerance is likely to be complex (Negrão et al., 2017; Morton et al., 2018). Here, we investigated the suites of traits underlying variation in salinity tolerance in sunflower, and examined their underlying genetic basis. Genotypes differed in terms of morphological, growth/biomass allocation, and leaf ionomic responses to salt stress. Morphological and ionomic traits were highly correlated amongst themselves under both control and saline conditions, as well as in their plasticity to salt stress. Given an observed trade-off between vigor (biomass accumulation in benign conditions) and the effect of salt stress, we decoupled our metric of tolerance from this expected response. Consistent with prior findings, the most vigorous genotypes in control conditions tended to remain the best performers (most biomass) under salt stress (**Fig. 2a;** (Temme et al., 2019a; Temme et al., 2019b). In addition, similar to previous work, these high vigor genotypes had the greatest proportional reduction in biomass in response to stress (Temme et al., 2019a; Temme et al., 2019b). The low vigor of the genotypes least affected by salinity stress might be attributable to the maintenance of costly mechanisms related to salinity tolerance (Munns and Tester, 2008; Shabala et al., 2019). However, by taking into account the relationship between vigor and the proportional decline in performance, we were able to focus on the traits associated with better/worse than expected performance, independent from vigor.

Using this metric of expectation-deviation tolerance, we found that genotypes that had a greater increase in root mass fraction, fine root mass fraction, and chlorophyll content had a lower than expected decline in biomass (i.e., they had a higher expectation-deviation tolerance; **Fig. 5**). Rather than variation in the traits themselves, it was the magnitude of trait plasticity that was most clearly linked to variation in expectation-deviation tolerance. Functionally, there are some known mechanisms that could play a role in these responses. For example, higher root mass fractions could allow for increased storage of Na in the roots, as has been found for soy (Guan et al., 2014). Alternatively, increased root branching (resulting in greater fine root mass fraction) has been suggested to reduce the energetic cost of Na exclusion (Zolla et al., 2010; Munns et al., 2020a)). Notably, this could come at the cost of reduced water uptake during the day (Arsova et al., 2019), suggesting a possible interaction between the osmotic and ionic components of salt stress through root branching. Greater chlorophyll content as measured could be the result of more chlorophyll per area or thicker leaves. Comparing this to the response in LMA, which also increased, our results could indicate greater leaf succulence under salinity stress, thereby diluting accumulated Na (Munns et al., 2016).

**Figure 5.**
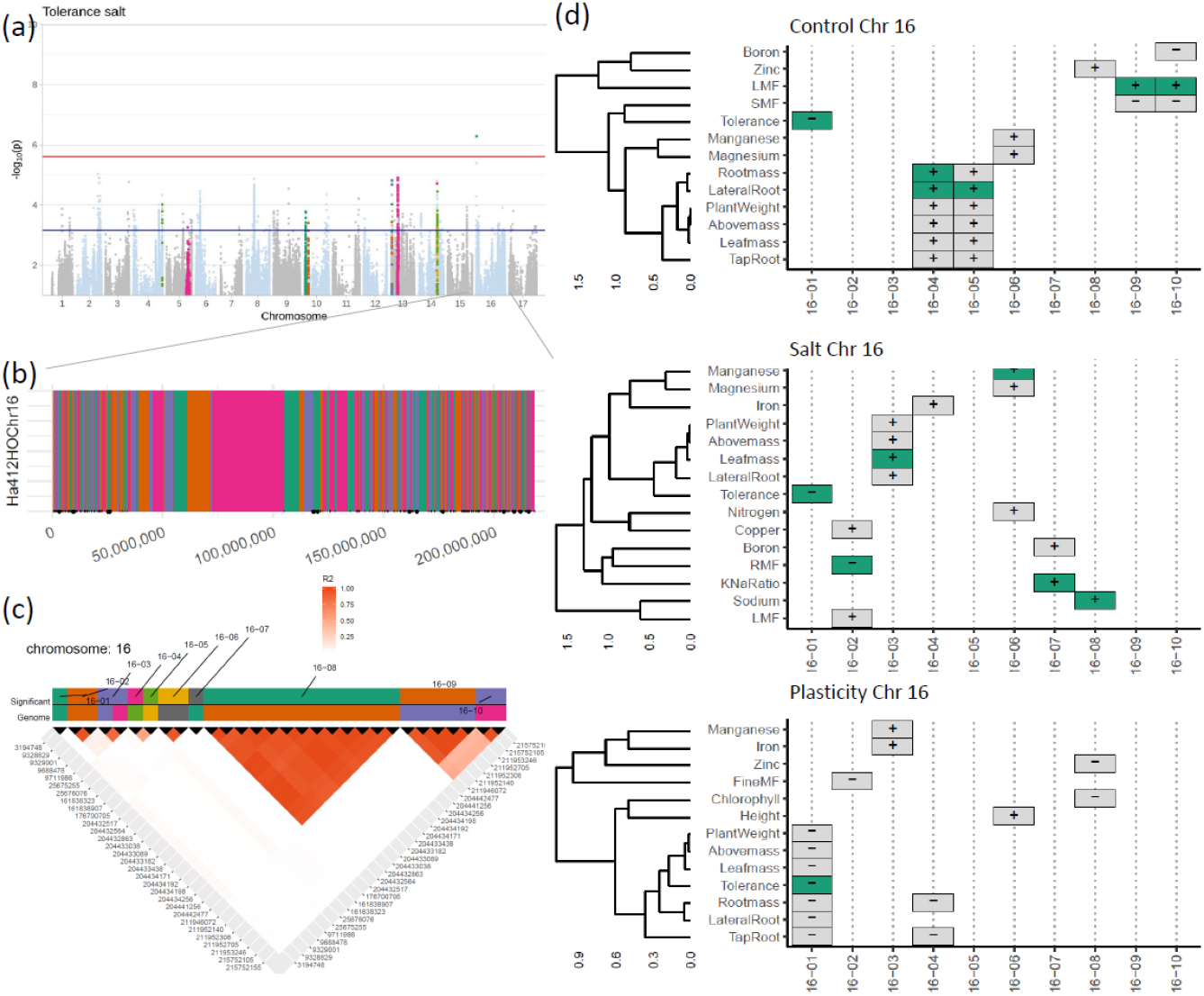
Overview of GWAS and trait co-localization analyses. Association testing was performed for trait values under control and salt stressed conditions, as well as the proportional change in trait value between treatments. **(a)** GWAS of salinity tolerance (defined as deviation from the expected response) resulted in a single independent region on chromosome 16 with a single significant association (above the red line). An additional 12 regions harbored suggestive associations (i.e., SNPs in the top 0.1% of likelihood values; above the blue line) for tolerance and significant associations for at least one other trait. **(b)** Zooming in on chromosome 16, the 103607 SNPs cluster together in 1732 haplotype blocks and 856 individual SNPs. Blocks that contain significant SNPs are marked by a black dot. **(c)** Evaluating all significant SNPs across all traits and “environments” on chromosome 16 reveals that these SNPs correspond to 11 unique haplotype blocks. However, after re-running the blocking algorithm (see methods for details), two blocks could be collapsed based on their linkage. **(d)** Visualising trait co-localization along chromosome 16 reveals relatively few instances of genomic overlap in traits with significant SNPs (green tiles), but substantial overlap in traits that are suggestive for a region (gray tiles). Plus or minus symbols in the tiles indicate the sign of beta (effect of minor allele on trait). Traits are ordered using hierarchical clustering based on their Spearman correlation strength.

Besides a substantial increase in Na content, the concentration of a suite of elements (N, P, K, S, Ca, Mg, Mn, Fe, Cu, Zn, B) was affected by salinity stress. We previously found a connection between leaf S content and salinity tolerance in sunflower (Temme et al., 2019b), possibly related to root sulpholipid content (Erdei et al., 1980). Additionally, Mn content has been linked to salt tolerance in Brassica (Wang et al., 2010; Chakraborty et al., 2016). However, here we found that lower leaf Na content, higher leaf K content, and higher K/Na ratio were associated most strongly with greater expectation-deviation tolerance. This is not surprising given that these traits are well known to be involved in salinity tolerance. Excluding Na from the leaf while maintaining K uptake and adequate levels in the cytosol are key to plants functioning under saline conditions (Munns and Tester, 2008; Munns et al., 2020a).

High vigor was correlated with a suite of traits including a relatively high fine root mass fraction (yet low overall root mass fraction) and high leaf Mn and Mg content **(Supplemental Fig. S3**). Taken together, these traits suggest nutrient uptake, and/or allometric scaling of root traits with vigor (Wang et al., 2019), as key components in vegetative growth. In a field setting, increased fertilization indeed produces increased growth in sunflower, though this did not ultimately translate into a yield increase; instead, it resulted in increased lodging and reduced yields (Schultz et al., 2018). This finding highlights the complexity of translating vegetative growth to yield.

In terms of GWAS results, we identified at least one genomic region with a significant association with the majority of traits either in the control treatment, salt treatment, or plasticity of the trait across treatments (**Table 2**). Heritability estimates for biomass, biomass fractions, and morphological traits were generally high, whereas estimates for ionomic traits were much lower, potentially owing to the lack of replicates for the ionomic traits (Kruijer et al., 2015). Relative effect sizes of alleles ranged from 6% for zinc under salt stress to 36% for chlorophyll content under control conditions **(Supplemental Table S3**). The identification of several potential regions of interest for the majority of traits combined with these sometimes sizable relative effect sizes of individual loci showcases the potential for adjusting trait expression through targeted breeding, or even genome editing once the underlying genes have been identified.

Considering trait expression across both treatments, the infrequent occurrence of genomic regions with a significant association for a trait in both treatments (**Supplemental Table S1**) was consistent with the occurrence of substantial GxT (**Table 1**) for the majority of traits (Des Marais et al., 2013). More commonly, regions were at least suggestively (or significant and suggestively) associated with traits in both environments (**Supplemental Fig. S7**). This highlights a potential consequence of stringent multiple comparison corrections: i.e., that false negatives may occur and give the appearance of environment specificity. Interestingly, there were far fewer significant and suggestive regions overlapping with plasticity in trait values. This suggests independent genomic control of the expression of a trait and the plasticity in that trait (Kusmec et al., 2017)).

Significant colocalizations across traits were also infrequent, with only 39 out of 189 regions being significantly associated with multiple traits. However, if we include suggestive co-localizations (i.e., regions that were significant for one trait and in the top 0.1% for at least one other trait), this number rises to 161 regions being significant for at least one trait and suggestive for at least one other. Thus, it may be that the apparent paucity of regions influencing multiple traits was due, at least in part, to the highly stringent significance thresholds. The colocalization of multiple traits, either significant or suggestive **(Supplemental Fig. S6, Supplemental Table S1**), in either control, salt stressed, or the plasticity of trait responses between treatments, demonstrates the multivariate nature of the genetic controls of trait expression (Wagner et al., 2007; Wagner and Zhang, 2011; Sella and Barton, 2019). This multivariate nature represents a constraint on the development of new trait combinations, and needs to be taken into account when attempting to improve salt tolerance. Indeed, adjustments in the expression of one trait will likely influence others (Pailles et al., 2019; Temme et al., 2019a), thereby restricting the response to selection for particular trait combinations. Nevertheless, since our metric of expectation-deviation tolerance is independent of vigor and correlated with plasticity in trait values (which are generally independent of the trait values themselves), it should be possible to improve tolerance while minimizing undesired trade-offs.

Based on the annotations of genes in the significantly associated regions, we were able to identify several candidates for genes underlying observed trait variation (**Supplemental Table S1, S2**). Here, we highlight a few with potential impacts on the traits most closely related to salinity tolerance. In region 8-4 (53 genes, 6,161 kbp), which was associated with Na content under salt stress, we found *hypersensitive-induced reaction 1 protein*, which has been linked to K efflux (impacting K/Na ratio) in plant cells (Jung and Hwang, 2007). Similarly, in region 12-14 (29 genes, 827 kbp), which was likewise associated with Na content under salt stress, we found two genes related to vacuolar function (*vacuolar (H+)-ATPase G subunit* and *vacuolar protein sorting-associate protein Vta1/callose synthase*) that could play a role in vacuolar sequestration of Na, which is known to be an important tolerance mechanism (Gaxiola et al., 2016; Schilling et al., 2017; Munns et al., 2020a). While these gene descriptions are consistent with putative mechanisms, they are often not the only genes in an independent block. For example, region 10-2 contains a *calcium permeable stress-gated cation channel 1 transmembrane domain-containing protein*, and has a significant association with K/Na ratio under salinity stress, yet this is also one of the larger regions (93,336 kb) and is associated with 11 traits (some in multiple environments **Supplemental Table S1**) and contains 1,397 genes. Future work on gene expression changes under salinity stress along with functional analyses of the most promising candidates will allow us to develop a more complete understanding of the molecular basis of observed trait variation.

We identified a single small region on chromosome 16, containing a single gene, associated with expectation-deviation tolerance. Interestingly, aside from biomass, none of the measured traits co-localized with that same region. This suggests the existence of a potentially unmeasured mechanism conferring salinity tolerance that is influenced by variation in this region. Genotypes carrying the minor allele at this locus were found to be less tolerant to salinity stress, but did not show any difference in performance under control conditions (**Fig. 6**) indicating strong GxT for this allele along with a lack of significant performance trade-offs. The gene of interest in this region is a putative alpha-mannosidase (**Supplemental Table S3**). In *Arabidopsis*, alpha-mannosidases are involved in a complex pathway involved in regulating glycoprotein abundance during salt stress (Liu et al., 2018) and they also play a role in lateral root formation (Veit et al., 2018). Given that we found that plasticity in root mass allocation was associated with expectation-deviation tolerance (though it did not co-localize with this region), and alpha-mannosidase is related to aspects of root formation in the literature, this gene appears to be a worthwhile candidate for future mechanistic work on sunflower root development and salinity tolerance.

**Figure 6.**
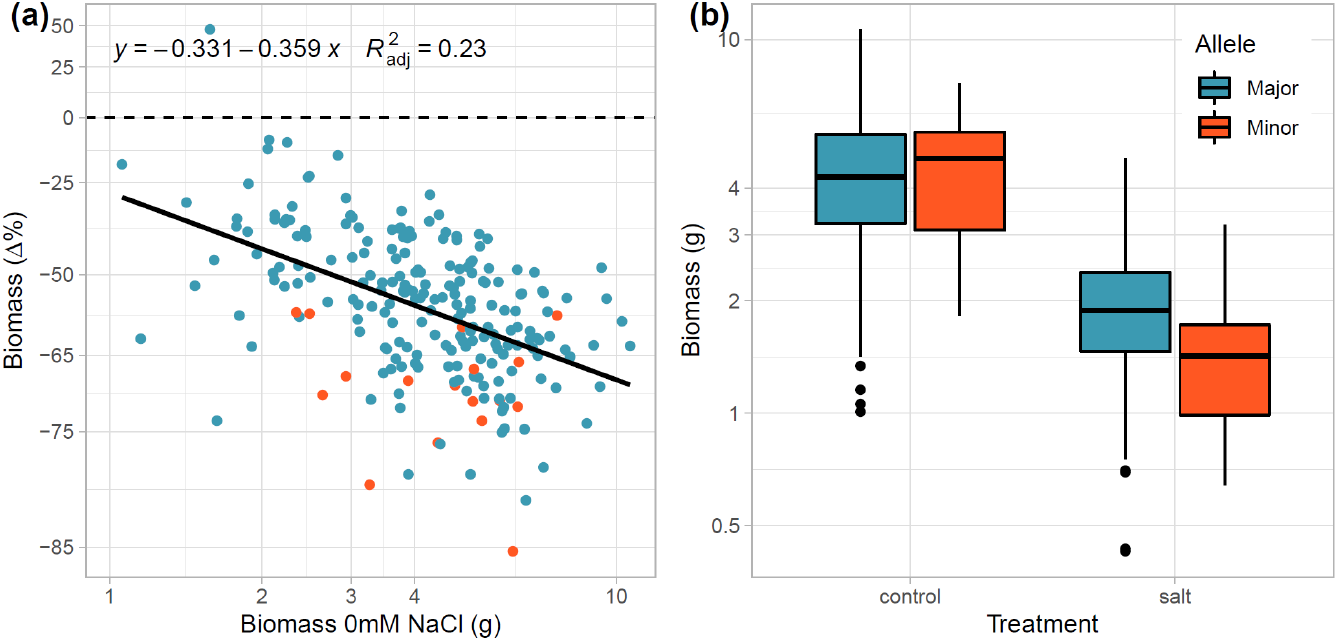
Allelic effects of region 16-1. (a) Plasticity vs. biomass under control conditions (i.e., vigor) for genotypes with the major (blue) vs. minor (orange) allele. Plasticity was estimated as the difference in natural log transformed values (control-salt) but converted here to Δ% (via e^Δln(trait)^-1) change from control for ease of interpretation. (b) The relationship between the treatment, allele (major vs. minor), and biomass accumulation. Note the natural log-scaling of the axes.

Taken together, our results indicate that salinity tolerance can be improved in sunflower while minimizing trade-offs with performance. Indeed, we found evidence that trait variation and plasticity influence salt tolerance independent of performance under benign conditions, and that these traits and vigor tend to map to distinct genomic regions. Since the traits of interest and their underlying genomic regions are distinct from those associated with high vigor (i.e., growth in benign conditions), this provides an avenue for increasing salt tolerance by fine-tuning trait variation/plasticity in high-performing genotypes without incurring costly trade-offs.

## Acknowledgements

We thank K. Bettinger, M. Boyd, K. Tarner, the rest of the greenhouse staff, the sunflower undergraduate army, and numerous members of the Donovan and Burke labs for help during the experiment. Special thanks to Greg Owens, Jean-Sébastien Légaré, and Loren Rieseberg for developing the sunflower SNPs used in this study, as well as their re-alignment to the HA412-HOv2 genome assembly. This work was supported by a grant from the NSF Plant Genome Research Program (IOS-1444522) to JMB and LAD.

## Supplementary materials

**Supplemental Table S1** List of significant genomic regions and associated significant/suggestive traits. Genomic regions of interest along with the number of significant and suggestive traits and number of genes in each region. Regions significant or suggestive for tolerance are bolded in the list of traits.

**Supplemental Table S2** List of genes per region. As detailed in the text, gene names were extracted from the annotation of the HA412-HOv2 assembly.

**Supplemental Table S3**. Detailed information on significant and suggestive regions per trait/treatment combination. Region notes the independent genomic region after rerunning the haplotype blocking on the significant SNPs. Haplotype block notes the unique block on the genome based on all SNPs. NSNP gives the number of significant SNPs in the haplotype block. Beta notes the effect of the minor allele (max of significant SNPs in that haplotype block). RES notes the max relative effect size of SNPs in that haplotype block.

**Supplemental Figure S1.** Haplotype block map of the sunflower genome. Haplotype blocks on the 17 individual sunflower genomes. Blocks are colored from the first SNP to the last SNP within each block. Adjacent blocks are colored in an alternating fashion repeating the same 4 colours. Black marks along the x-axis show singleton SNPs that did not fall within blocks. Note the large (93,336 kbp) region on chromosome 10, as well as the large region containing numerous singleton SNPs on chromosome 11.

**Supplemental Figure S2.** Trait reaction norms to salinity stress. Graphical representation of the response to salinity for the traits in table 1.

**Supplemental Figure S3.** Vigor and associated trait complexes. Vigor (i.e., biomass in control conditions) as compared to the biomass independent traits in control conditions (a & b) and to the leaf ionome in control conditions (c & d). Points indicate genotypes, whereas arrows represent eigenvectors of traits.

**Supplemental Figure S4.** Linkage disequilibrium heatmaps for significant SNPs on each chromosome. Linkage disequilibrium (estimated as *R^2^*) between all SNPs that are significant for at least one trait within and between both treatments. Colored bar notes the haplotype blocks SNPs belong to based on the haplotype map presented in **Fig S3**. Block membership is indicated for the calculation based on the genome-wide collection of SNPs (“genome”) and for the re-calculation based on just the significant SNPs (“significant”).

**Supplemental Figure S5.** Manhattan plots of all traits. Per-trait Manhattan plots are shown for trait values under control and salt-stressed conditions, as well as the plasticity between treatments (log difference). SNPs above the red line are significant after multiple comparison correction, SNPs above the blue line are in the top 0.1% of *P*-values. SNPs are colored by the “significant” haplotype regions per chromosome as displayed in **Fig S3**.

**Supplemental Figure S6.** Trait co-localization under control and salt-stressed conditions as well as their plasticity. Traits in (a) control, (b) salt stressed, and their (c) plasticity are clustered based on hierarchical clustering of their bivariate correlations. Traits grouped more closely together are more highly correlated. If a linkage block contained a significant SNP for a trait, all other traits were assessed to determine if any had a significant (green) or suggestive (i.e., top 0.1% of all SNPs; grey) in the same block. Plus/minus symbols in the tiles indicate the sign of *β* (effect of minor allele on trait). Linkage blocks are ordered by chromosome with black lines separating them. Sizes of the linkage blocks vary both in physical size as well as gene number.

**Supplemental Figure S7.** Co-localization of traits across environments. If a linkage block contained a significant SNP for a trait in one environment, or trait plasticity across environments, the alternate values were assessed to determine if there were significant (green) or suggestive (i.e., top 0.1% of all SNPs; grey) associations across environments. Plus/minus symbols in the tiles indicate the sign of *β* (effect of minor allele on trait). Linkage blocks are ordered by chromosome with black lines separating them. Sizes of the linkage blocks vary both in physical size as well as gene number.

**Supplemental Figure S8.** Number of genes per significant region. Points indicate the number of genes present in the 189 linkage blocks identified from GWAS of all traits/trait plasticities. Note the log scale on the y-axis to aid in visualization.

