## Supplemental Figure for "Key traits and genes associate with salinity tolerance independent from vigor in cultivated sunflower (*Helianthus annuus* L.)"

**Figure S1. Haplotype block map of the sunflower genome**.
Haplotype blocks on the 17 individual sunflower genomes. Blocks are colored from the first SNP to the last SNP within each block. Adjacent blocks are colored in an alternating fashion repeating the same 4 colours. Black marks along the x-axis show singleton SNPs that did not fall within blocks. Note the large (93,336 kbp) region on chromosome 10, as well as the large region containing numerous singleton SNPs on chromosome 11.

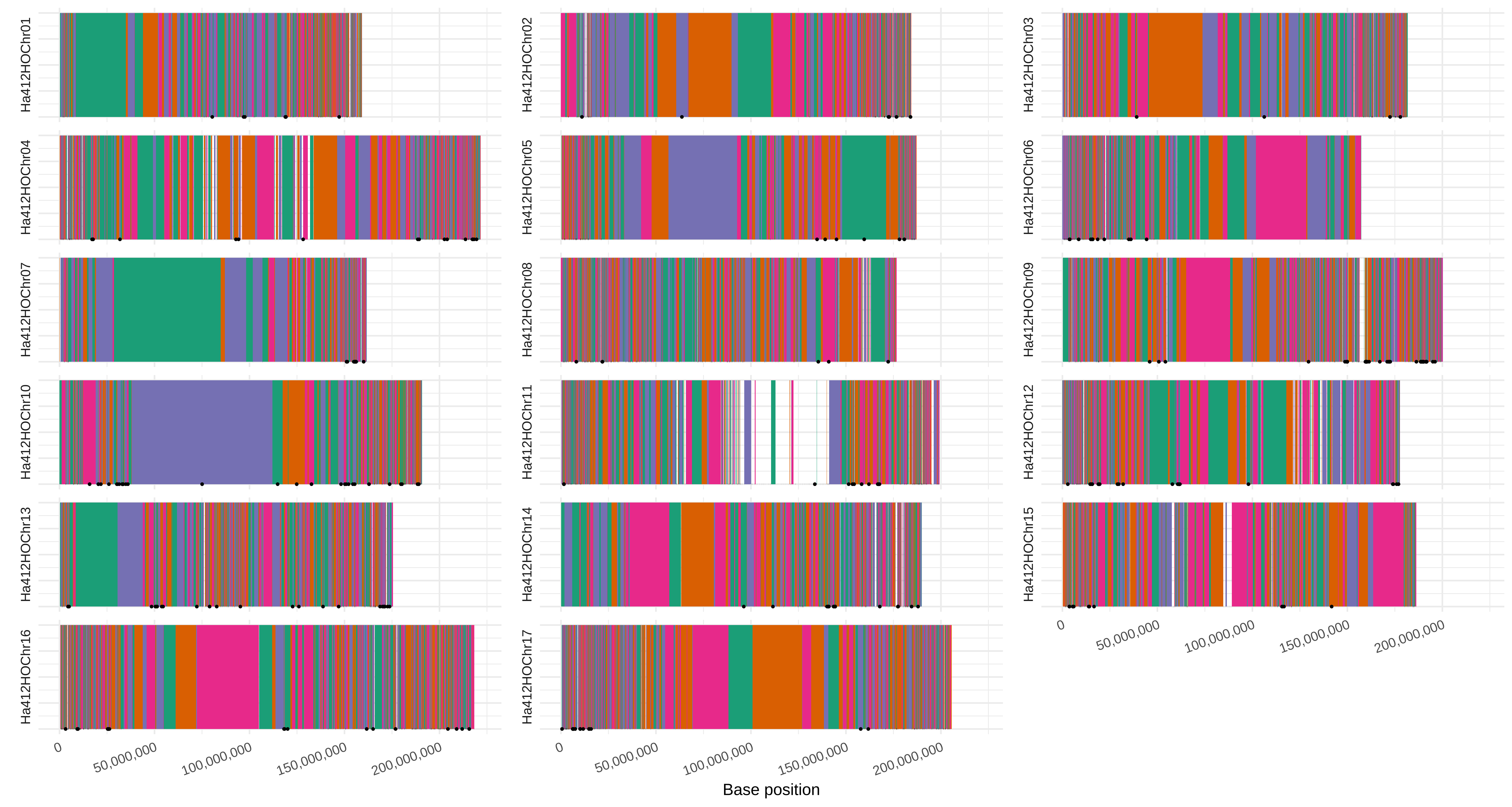

Ha412HOChr01
Ha412HOChr02
Ha412HOChr03
Ha412HOChr04
Ha412HOChr05
Ha412HOChr06
Ha412HOChr07
Ha412HOChr08
Ha412HOChr09
Ha412HOChr10
Ha412HOChr11
Ha412HOChr12
Ha412HOChr13
Ha412HOChr14
Ha412HOChr15
Ha412HOChr16
Ha412HOChr17
0
50,000,000
100,000,000
150,000,000
200,000,000
Base position

**Figure S2. Trait reaction norms to salinity stress.** Graphical representation of the response to salinity for the traits in table 1.

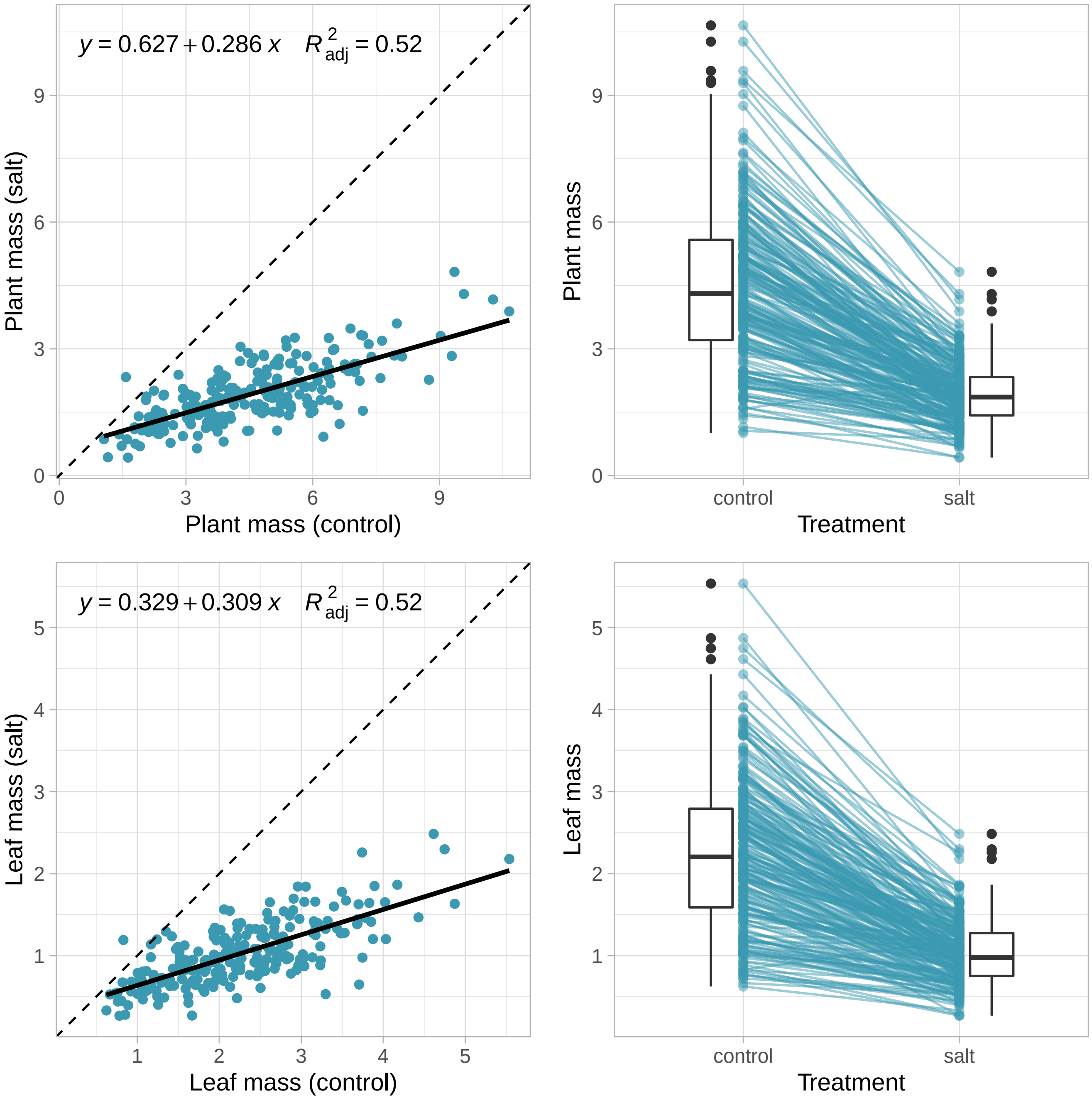

y = 0.627 + 0.286 x   R²adj = 0.52
Plant mass (salt)
Plant mass (control)
0
3
6
9
Plant mass
Treatment
control
salt
y = 0.329 + 0.309 x   R²adj = 0.52
Leaf mass (salt)
Leaf mass (control)
1
2
3
4
5
Leaf mass

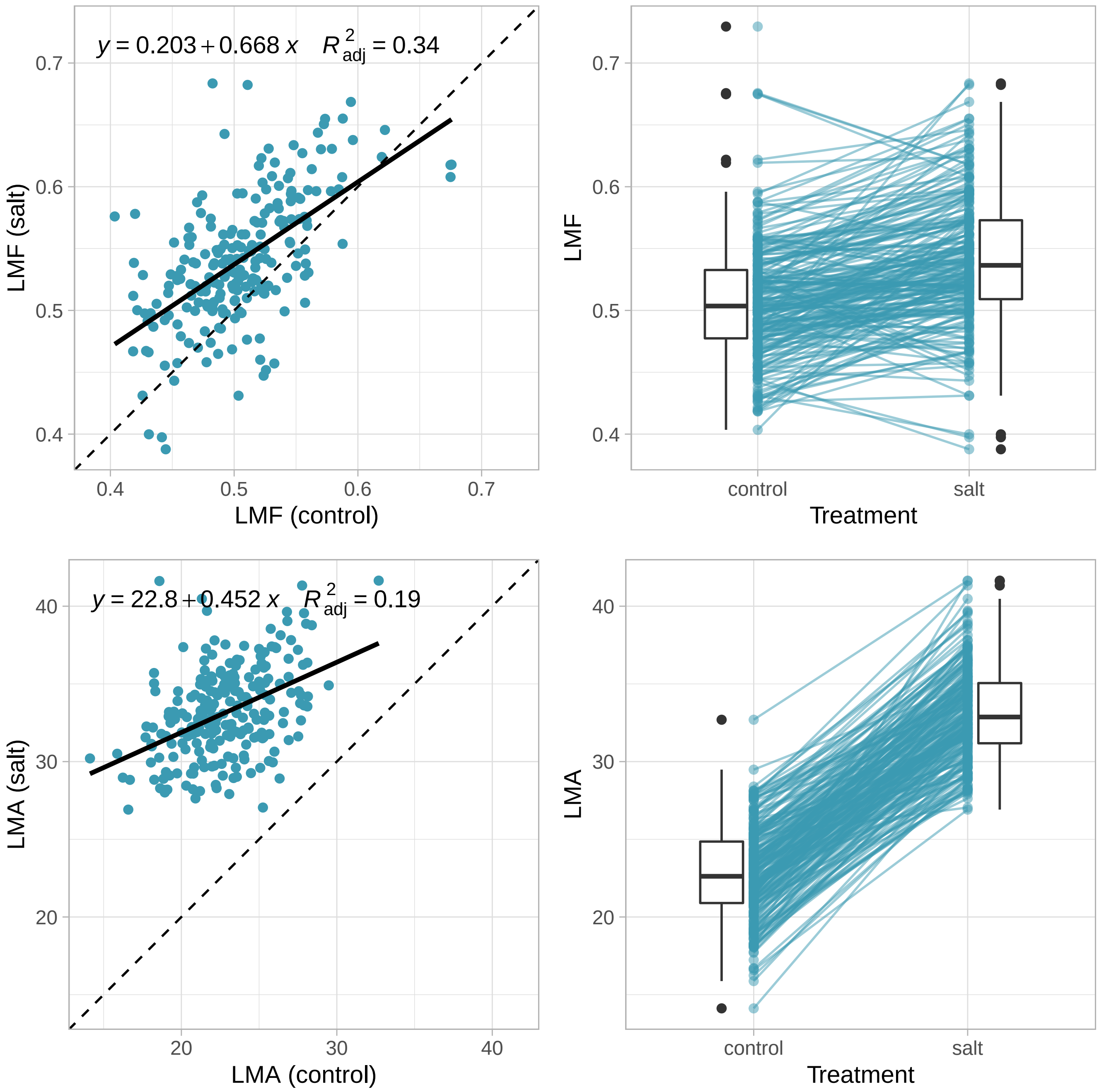

y = 0.203 + 0.668 x  R²adj = 0.34
0.7
0.6
0.5
0.4
LMF (salt)
0.4
0.5
0.6
0.7
LMF (control)
0.7
0.6
0.5
0.4
LMF
control
salt
Treatment
y = 22.8 + 0.452 x  R²adj = 0.19
40
30
20
LMA (salt)
20
30
40
LMA (control)
40
30
20
LMA
control
salt
Treatment

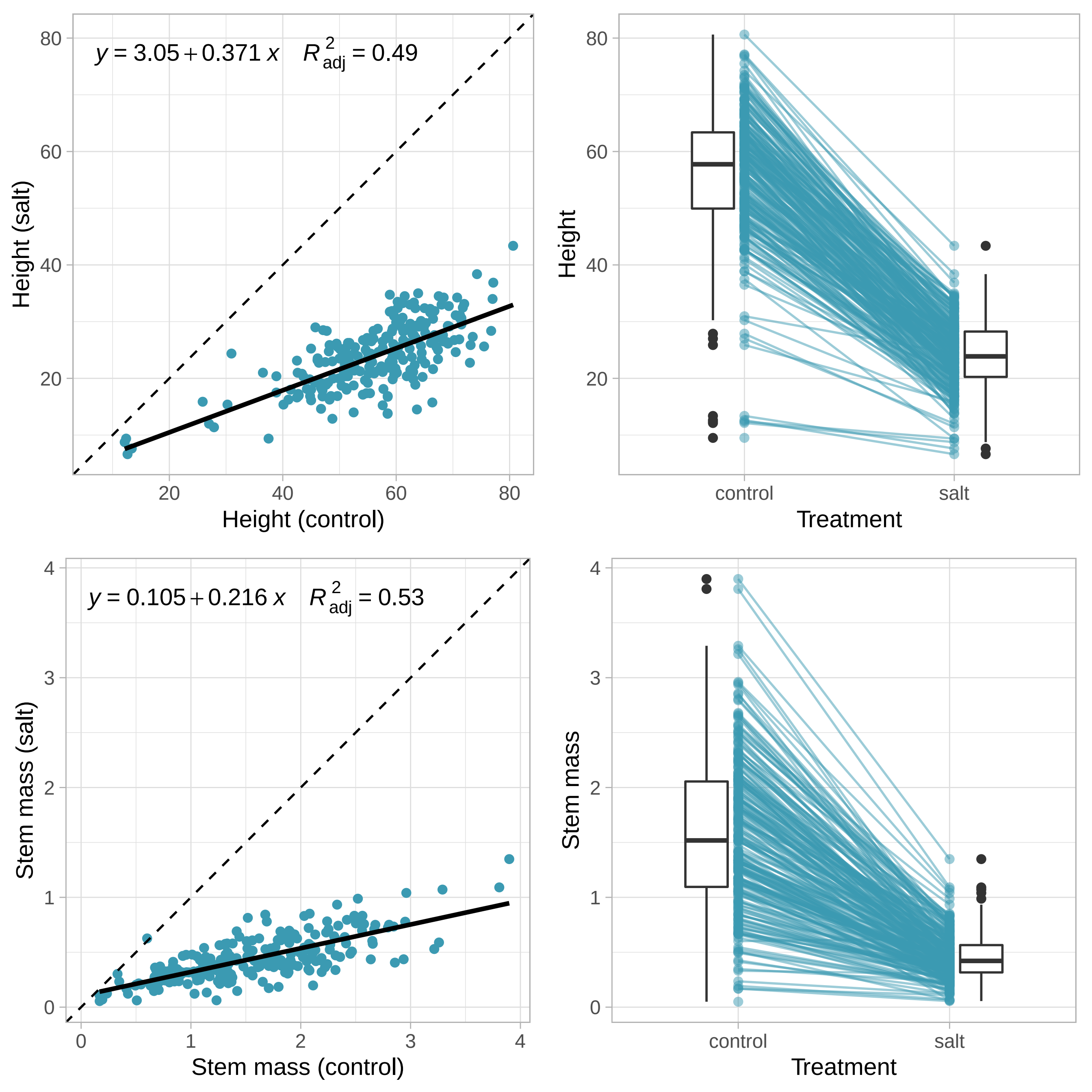

$y = 3.05 + 0.371\,x \quad R^2_{adj} = 0.49$
Height (salt)
Height (control)
Height
Treatment
control
salt
$y = 0.105 + 0.216\,x \quad R^2_{adj} = 0.53$
Stem mass (salt)
Stem mass (control)
Stem mass
Treatment
control
salt

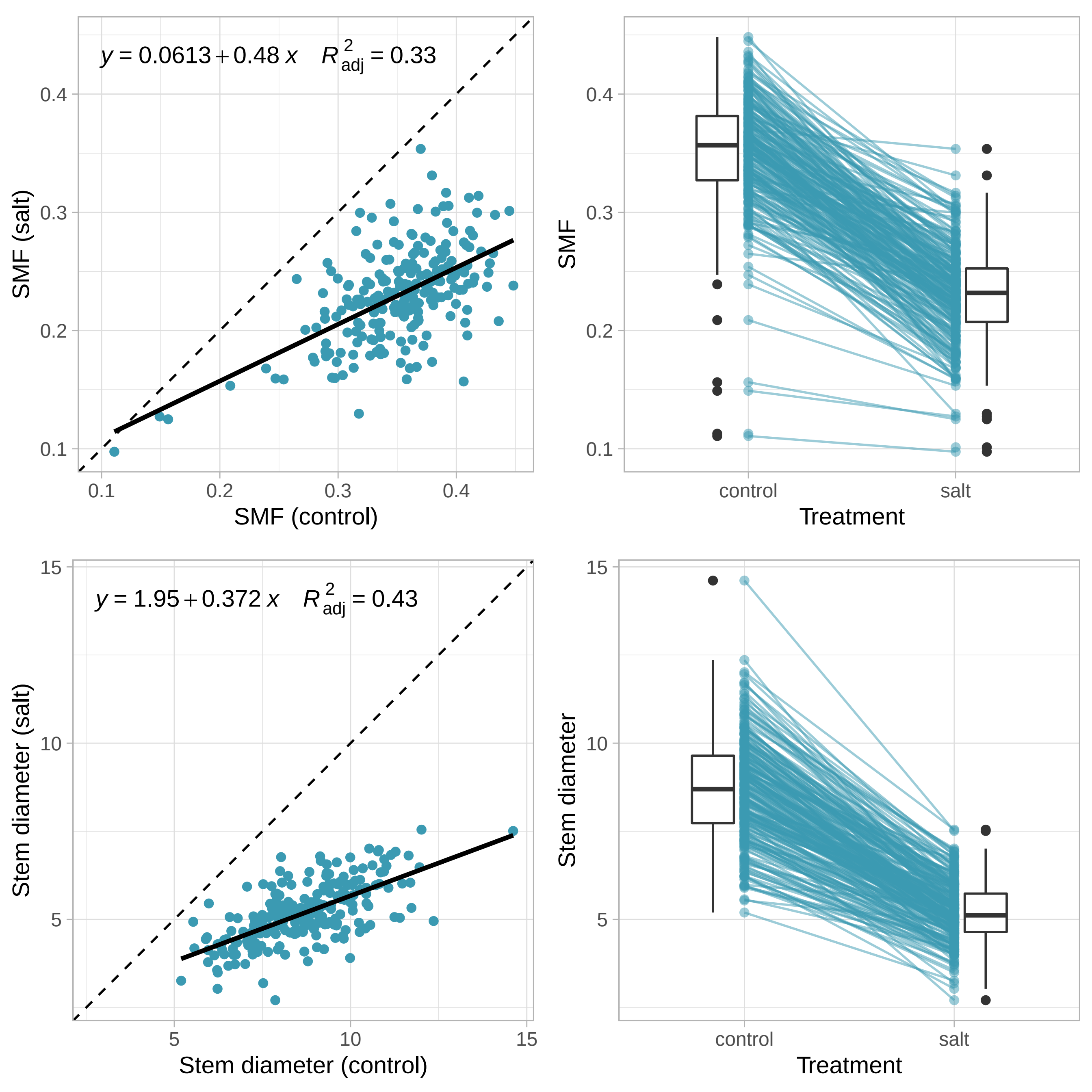

$y = 0.0613 + 0.48\,x \quad R^2_{adj} = 0.33$
SMF (salt)
SMF (control)
0.1
0.2
0.3
0.4
SMF
Treatment
control
salt
$y = 1.95 + 0.372\,x \quad R^2_{adj} = 0.43$
Stem diameter (salt)
Stem diameter (control)
5
10
15
Stem diameter

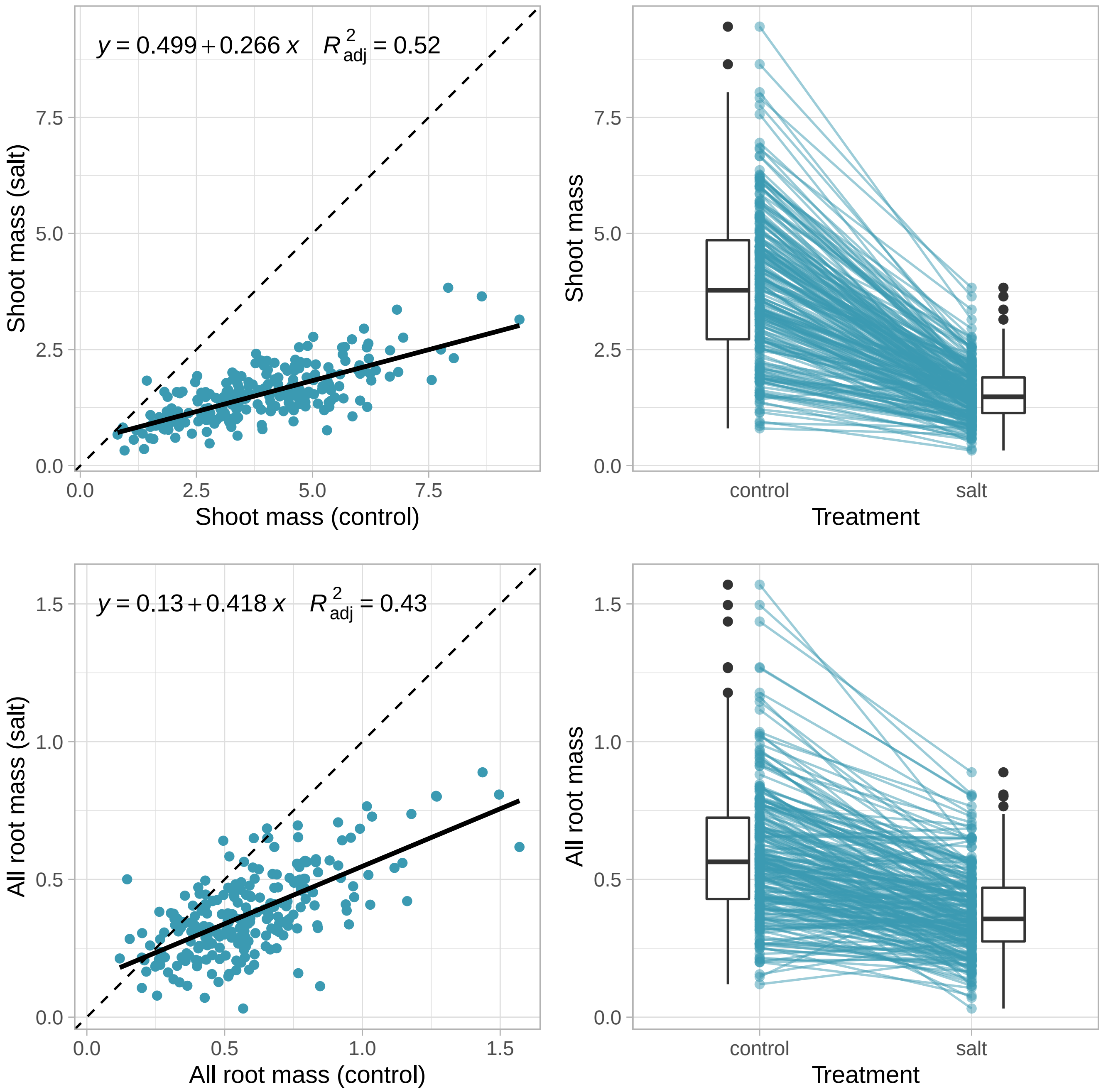

y = 0.499 + 0.266 x    R²adj = 0.52
Shoot mass (salt)
Shoot mass (control)
0.0
2.5
5.0
7.5
Shoot mass
Treatment
control
salt
y = 0.13 + 0.418 x    R²adj = 0.43
All root mass (salt)
All root mass (control)
0.0
0.5
1.0
1.5
All root mass
Treatment
control
salt

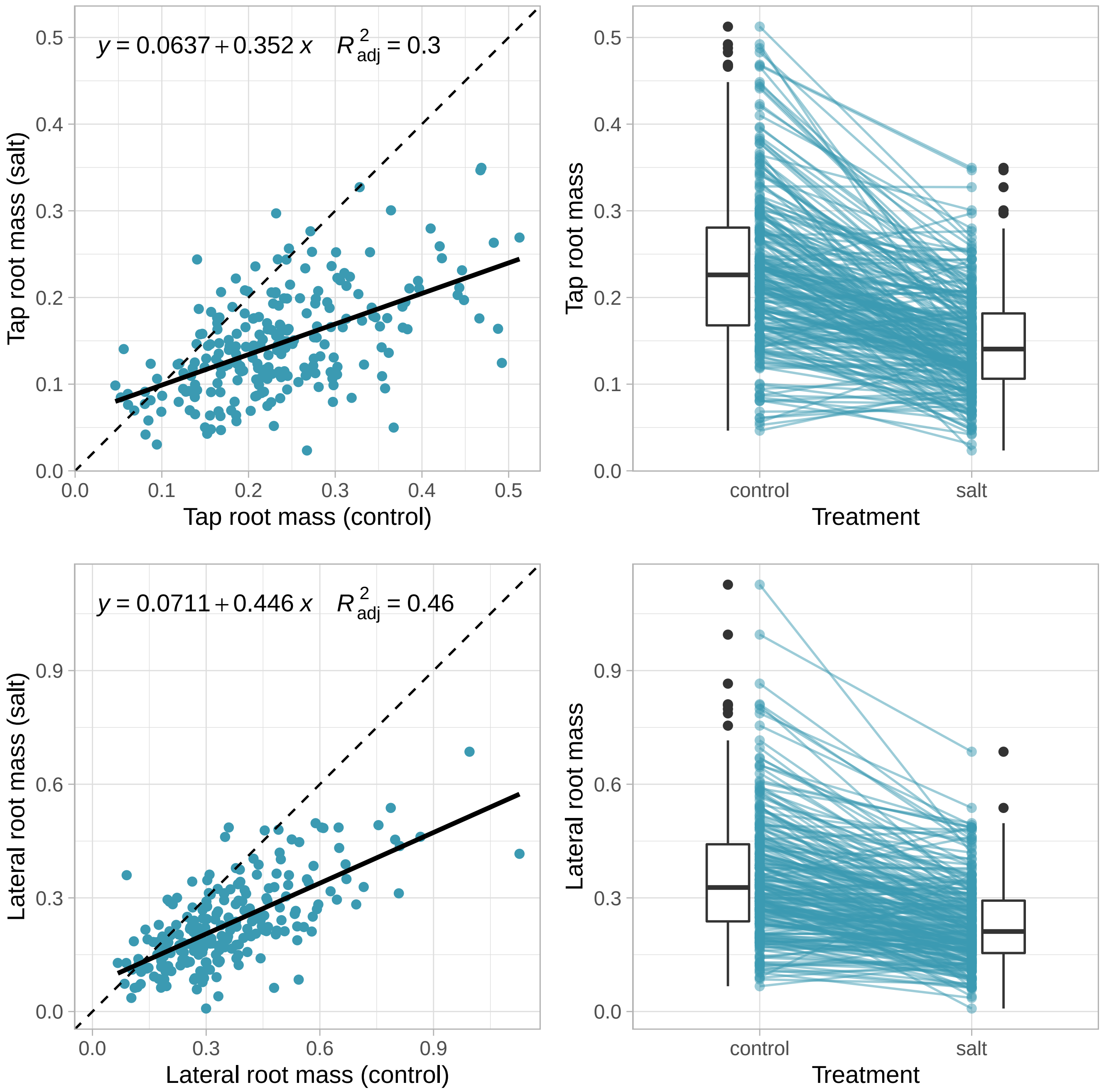

y = 0.0637 + 0.352 x    R²adj = 0.3
Tap root mass (salt)
Tap root mass (control)
Tap root mass
Treatment
control
salt
y = 0.0711 + 0.446 x    R²adj = 0.46
Lateral root mass (salt)
Lateral root mass (control)
Lateral root mass
Treatment
control
salt

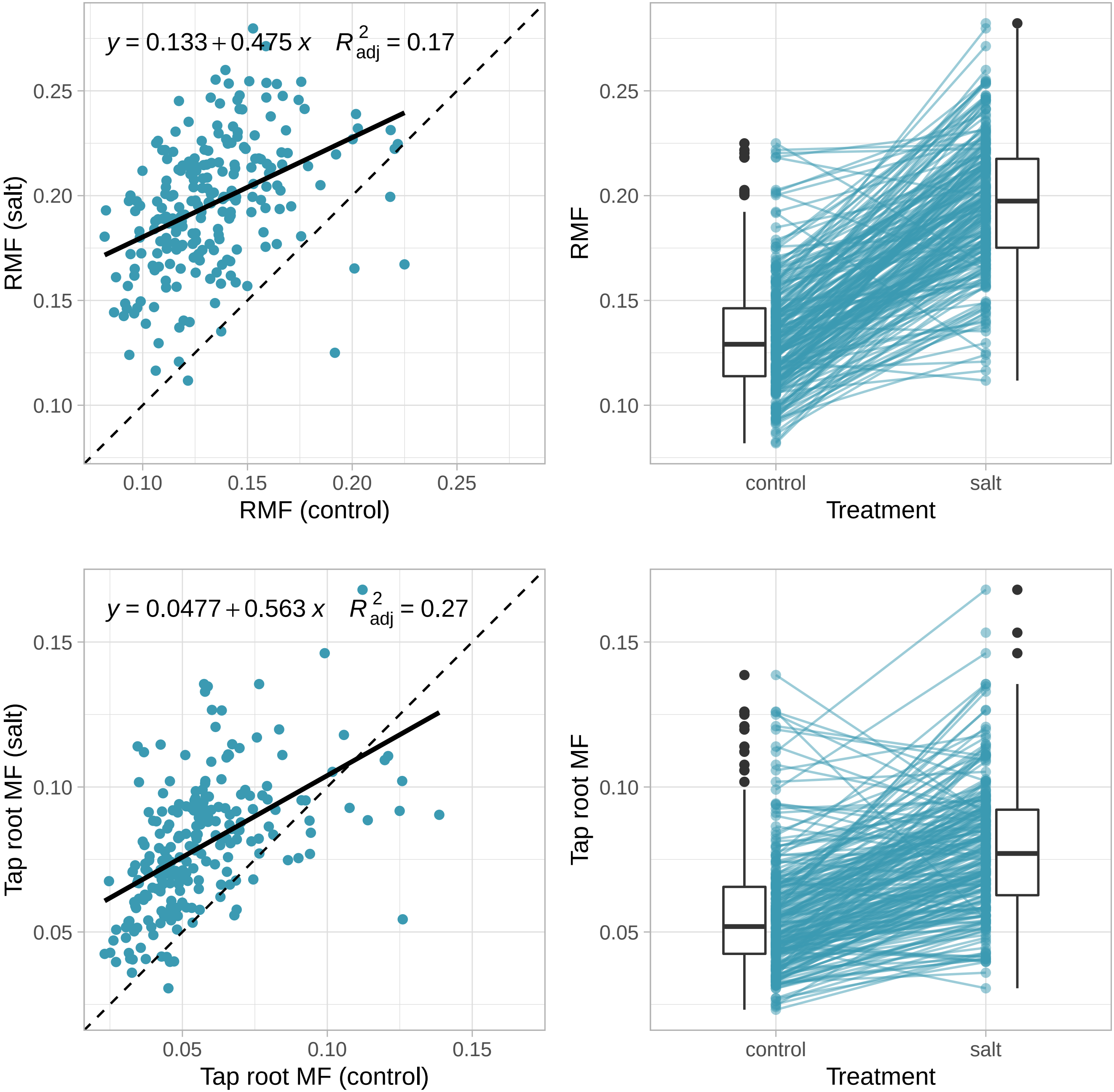

y = 0.133 + 0.475 x    R²adj = 0.17
RMF (salt)
RMF (control)
0.10
0.15
0.20
0.25
RMF
Treatment
control
salt
y = 0.0477 + 0.563 x    R²adj = 0.27
Tap root MF (salt)
Tap root MF (control)
0.05
0.10
0.15
Tap root MF
Treatment
control
salt

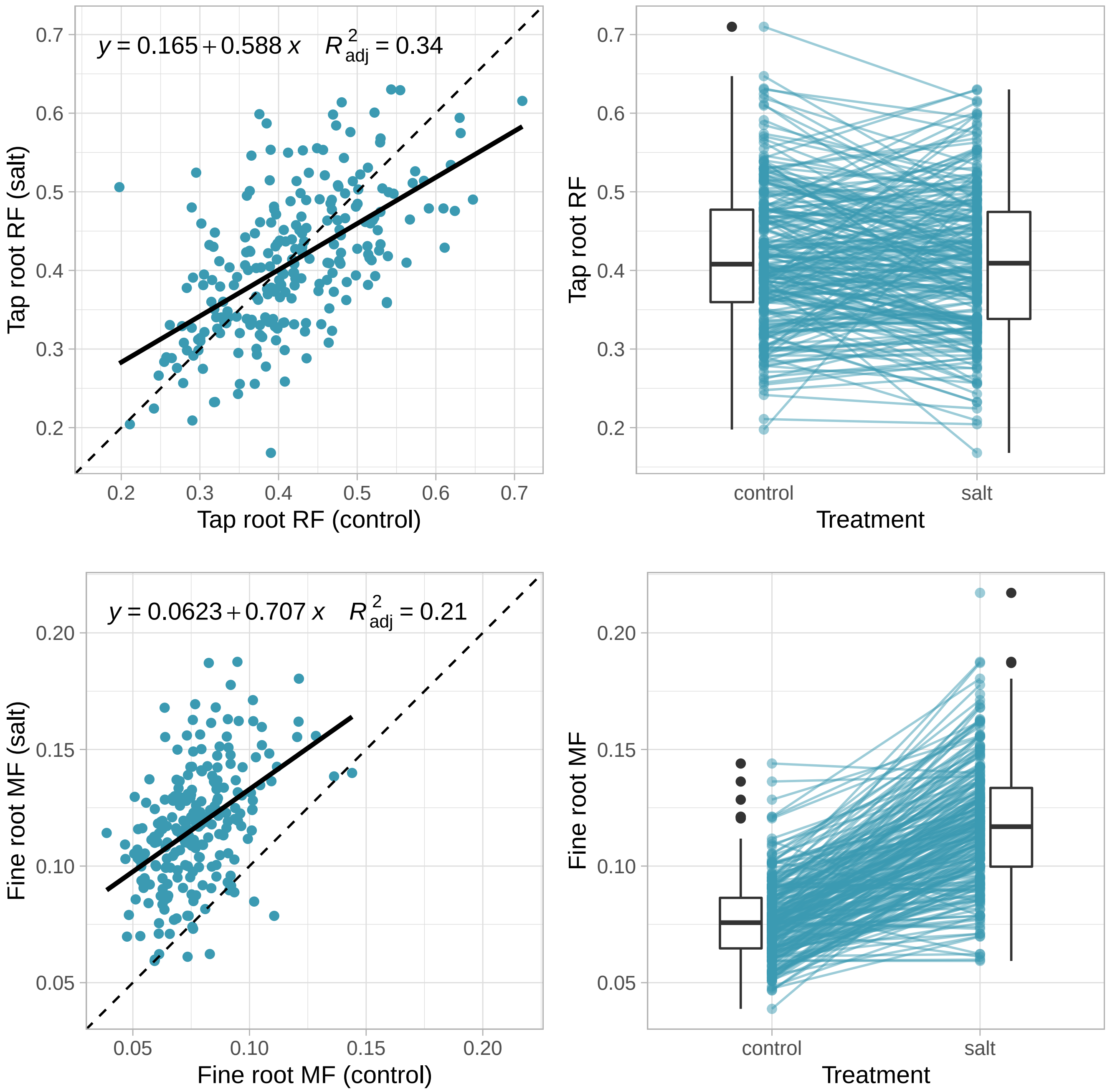

$y = 0.165 + 0.588\,x \quad R^2_{adj} = 0.34$
Tap root RF (salt)
Tap root RF (control)
0.2
0.3
0.4
0.5
0.6
0.7
Tap root RF
control
salt
Treatment
$y = 0.0623 + 0.707\,x \quad R^2_{adj} = 0.21$
Fine root MF (salt)
Fine root MF (control)
0.05
0.10
0.15
0.20
Fine root MF
control
salt
Treatment

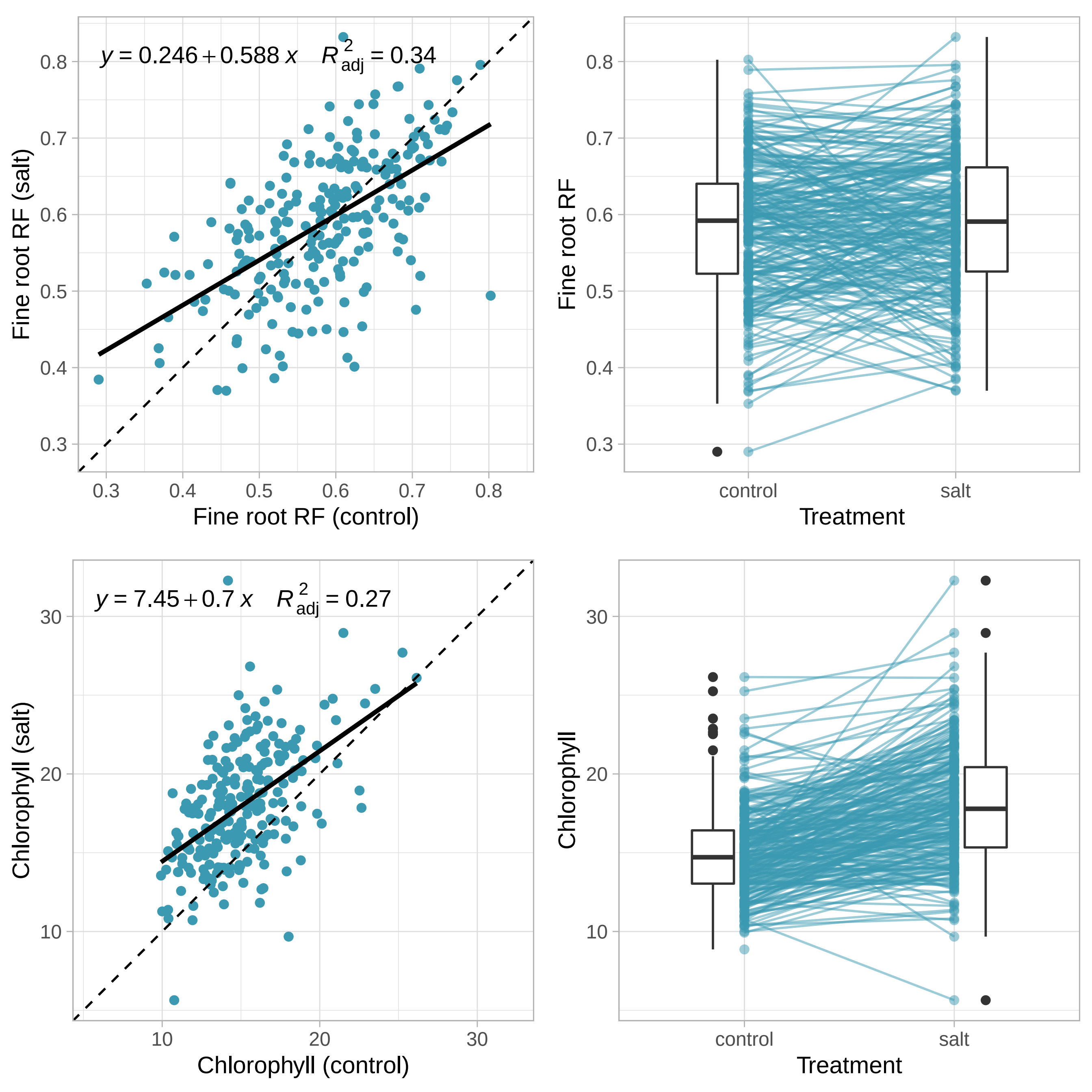

y = 0.246 + 0.588 x    R²adj = 0.34
Fine root RF (salt)
Fine root RF (control)
0.3
0.4
0.5
0.6
0.7
0.8
Fine root RF
Treatment
control
salt
y = 7.45 + 0.7 x    R²adj = 0.27
Chlorophyll (salt)
Chlorophyll (control)
10
20
30
Chlorophyll
Treatment
control
salt

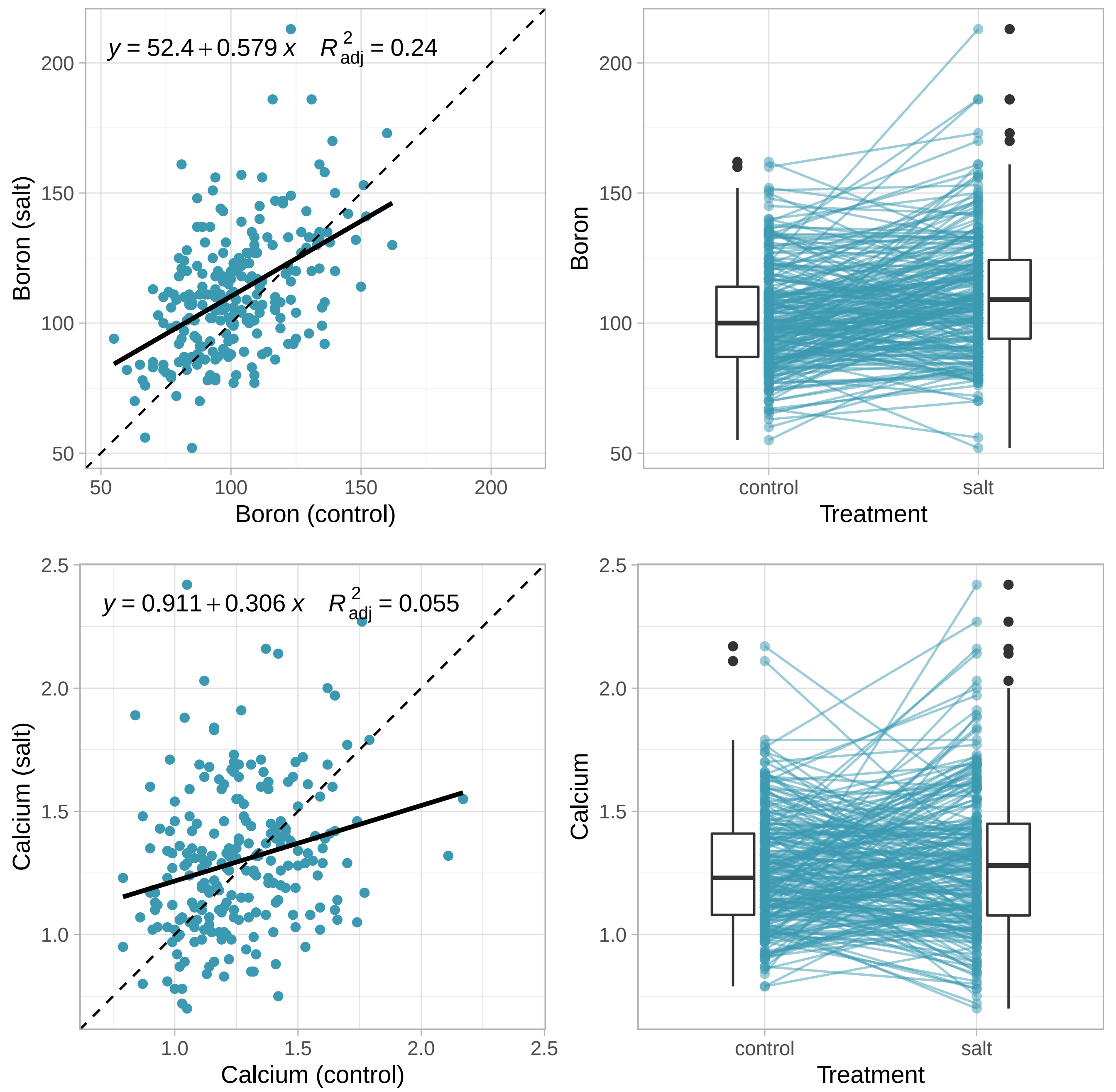

$y = 52.4 + 0.579\,x \quad R^2_{adj} = 0.24$
Boron (salt)
Boron (control)
Boron
Treatment
control
salt
$y = 0.911 + 0.306\,x \quad R^2_{adj} = 0.055$
Calcium (salt)
Calcium (control)
Calcium
Treatment
control
salt

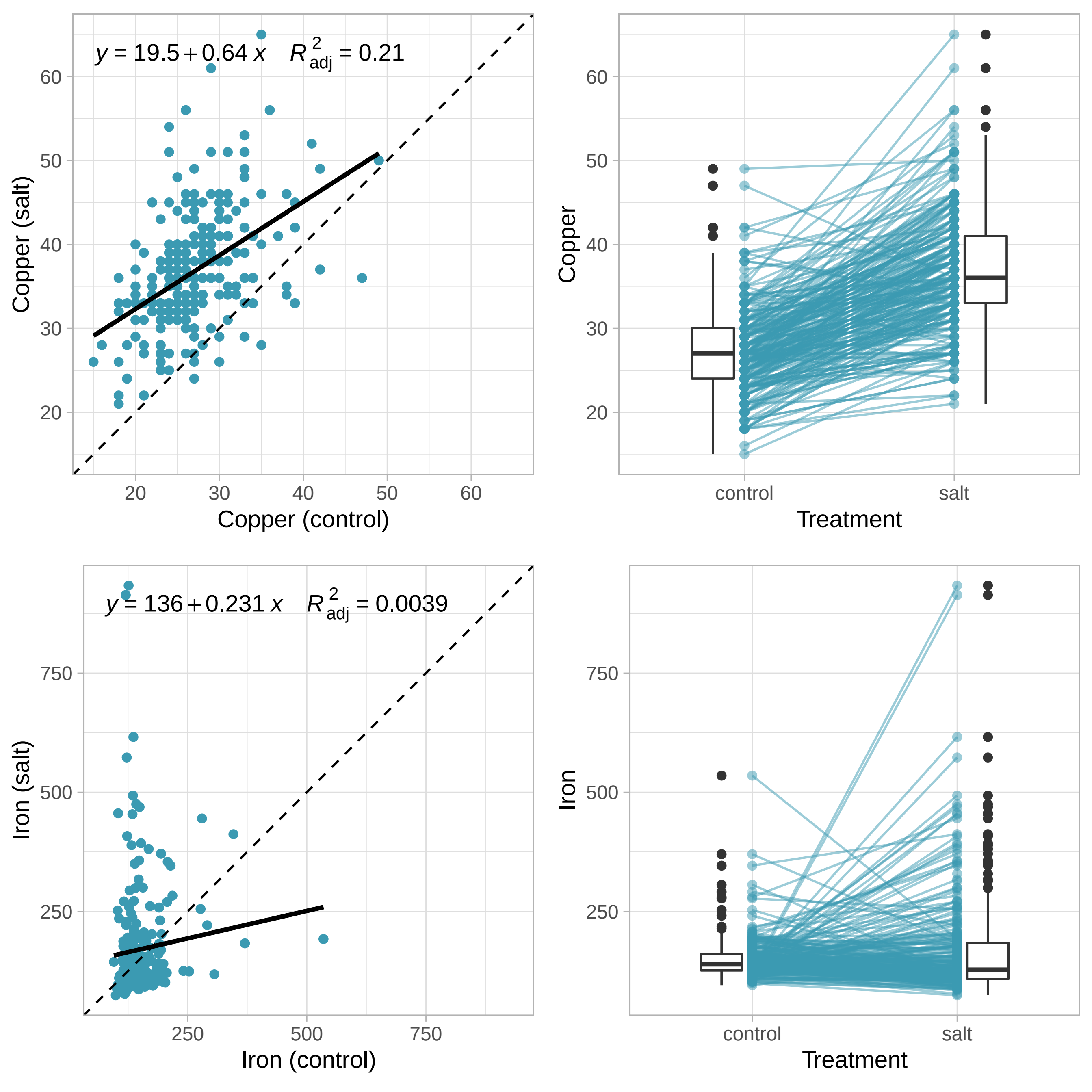

y = 19.5 + 0.64 x    R²adj = 0.21
Copper (salt)
Copper (control)
Copper
Treatment
control
salt
y = 136 + 0.231 x    R²adj = 0.0039
Iron (salt)
Iron (control)
Iron
Treatment
control
salt

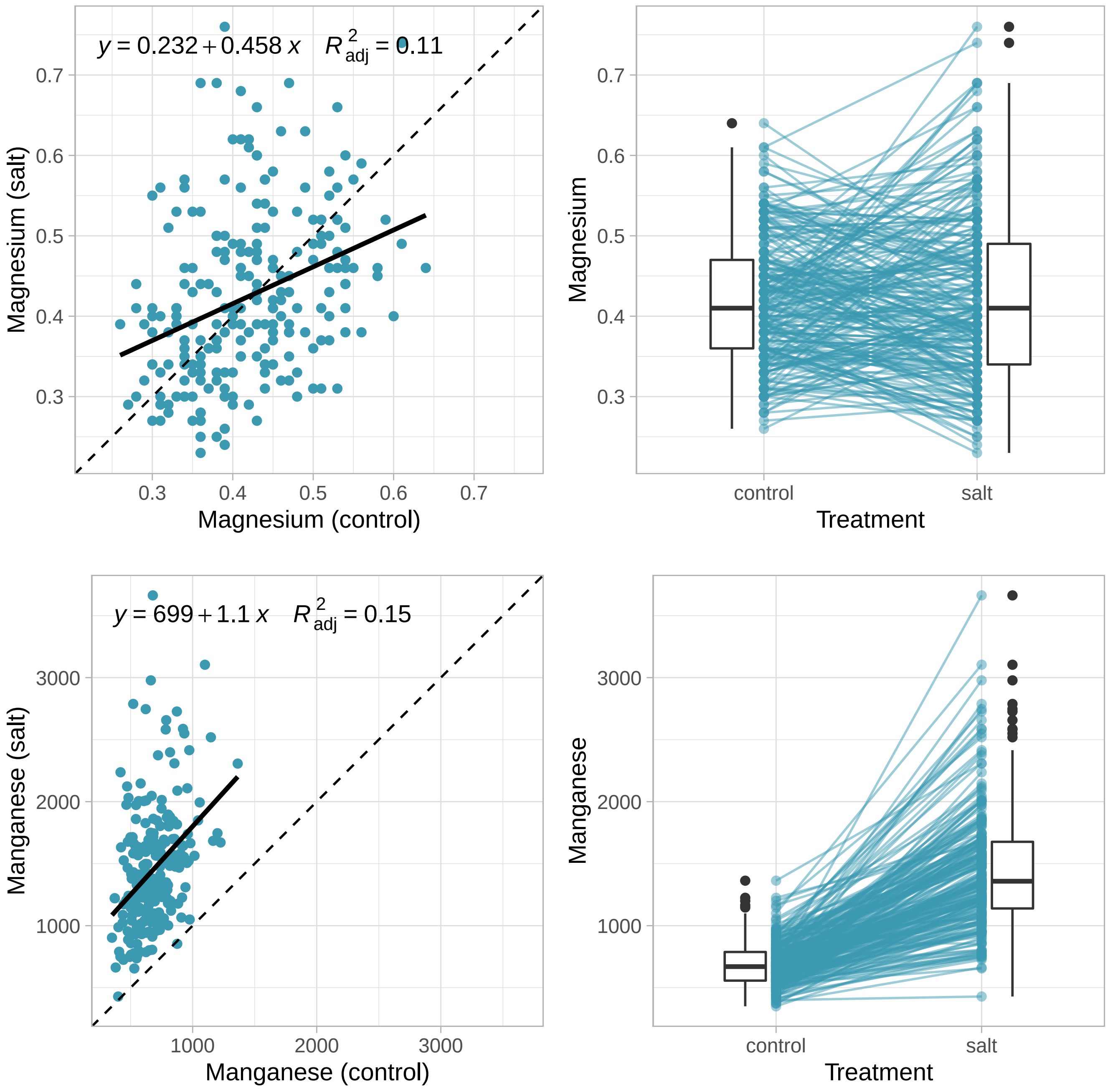

y = 0.232 + 0.458 x    R²adj = 0.11
Magnesium (salt)
Magnesium (control)
Magnesium
Treatment
control
salt
y = 699 + 1.1 x    R²adj = 0.15
Manganese (salt)
Manganese (control)
Manganese
Treatment
control
salt

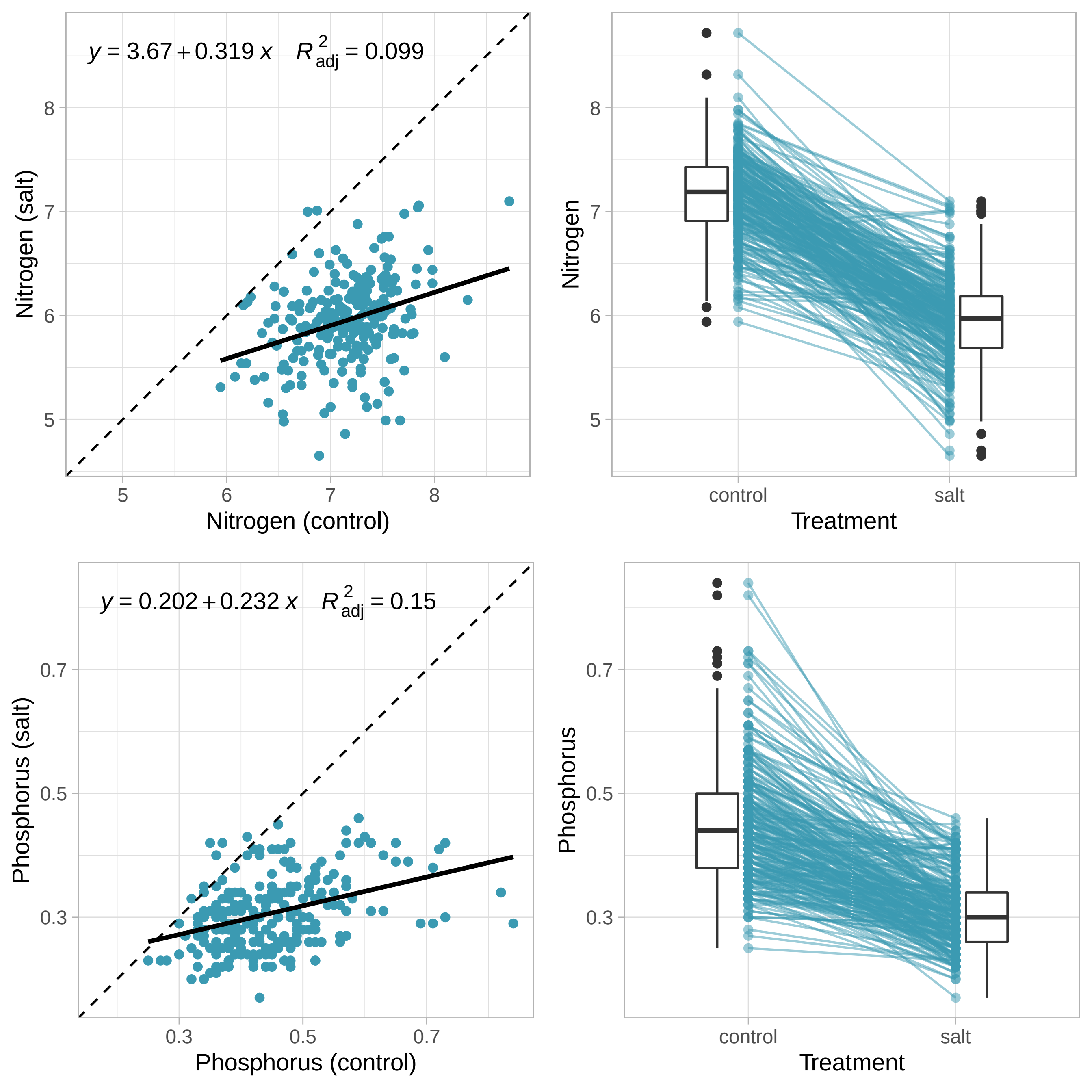

$y = 3.67 + 0.319\,x \quad R^2_{adj} = 0.099$
Nitrogen (salt)
Nitrogen (control)
Nitrogen
Treatment
control
salt
$y = 0.202 + 0.232\,x \quad R^2_{adj} = 0.15$
Phosphorus (salt)
Phosphorus (control)
Phosphorus

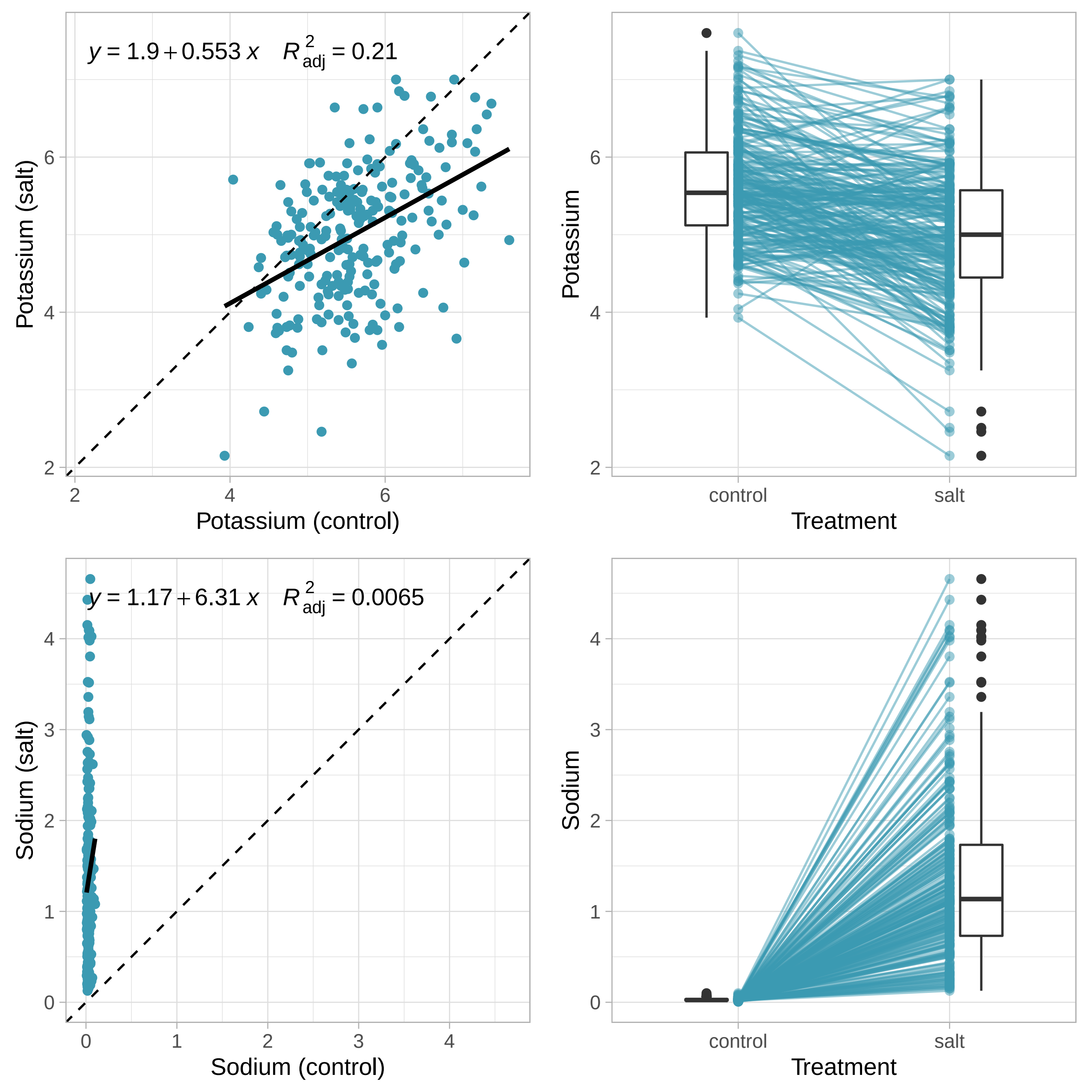

$y = 1.9 + 0.553\,x \quad R^2_{adj} = 0.21$
Potassium (salt)
Potassium (control)
Potassium
Treatment
control
salt
$y = 1.17 + 6.31\,x \quad R^2_{adj} = 0.0065$
Sodium (salt)
Sodium (control)
Sodium
Treatment
control
salt

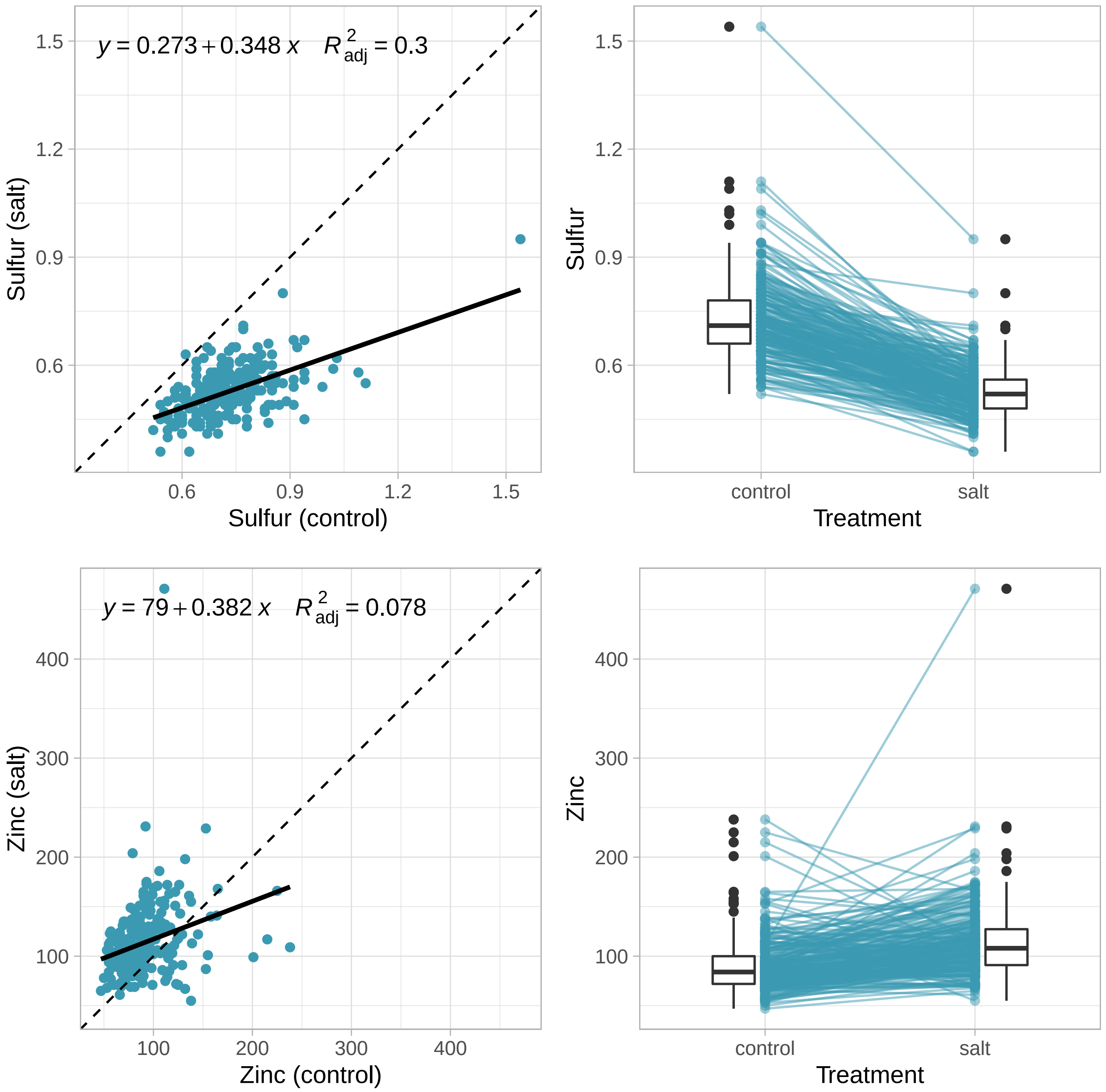

y = 0.273 + 0.348 x R2adj = 0.3
Sulfur (salt)
Sulfur (control)
1.5
1.2
0.9
0.6
Sulfur
Treatment
control
salt
y = 79 + 0.382 x R2adj = 0.078
Zinc (salt)
Zinc (control)
100
200
300
400
Zinc

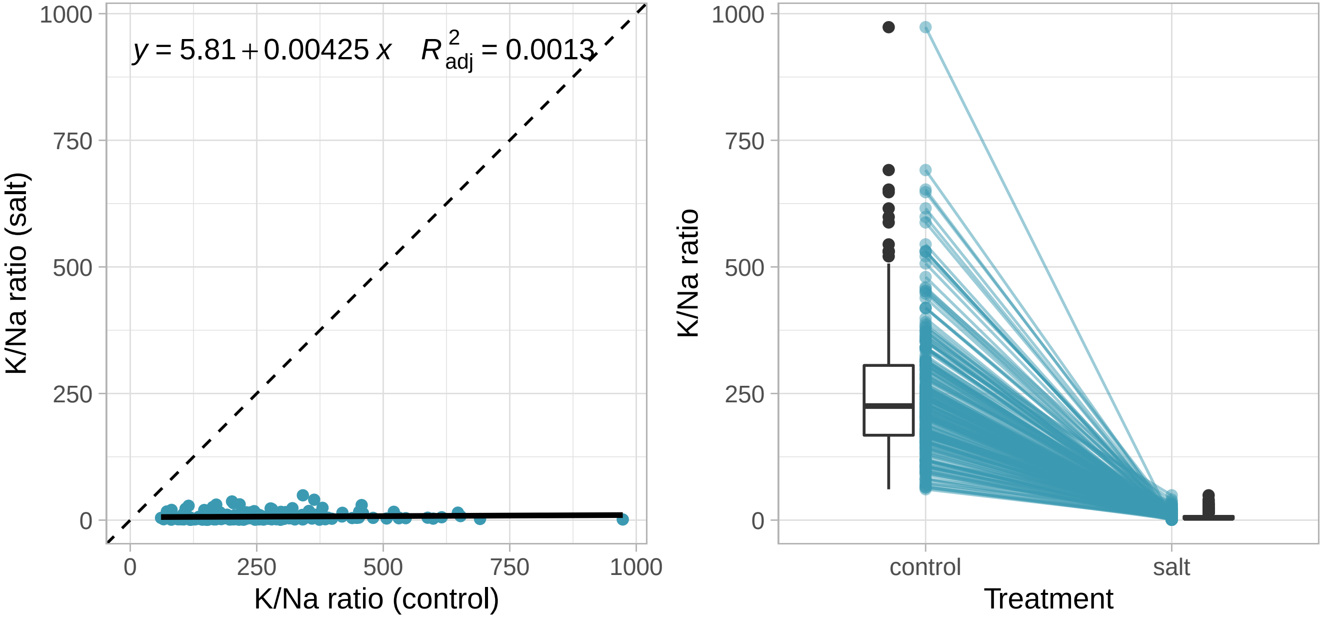

y = 5.81 + 0.00425 x    R²adj = 0.0013
1000
750
500
250
0
K/Na ratio (salt)
0
250
500
750
1000
K/Na ratio (control)
K/Na ratio
control
salt
Treatment

**Figure S3. Vigor and associated trait complexes.** Vigor (i.e., biomass in control conditions) as compared to the biomass independent traits in control conditions (a & b) and to the leaf ionome in control conditions (c & d). Points indicate genotypes, whereas arrows represent eigenvectors of traits.

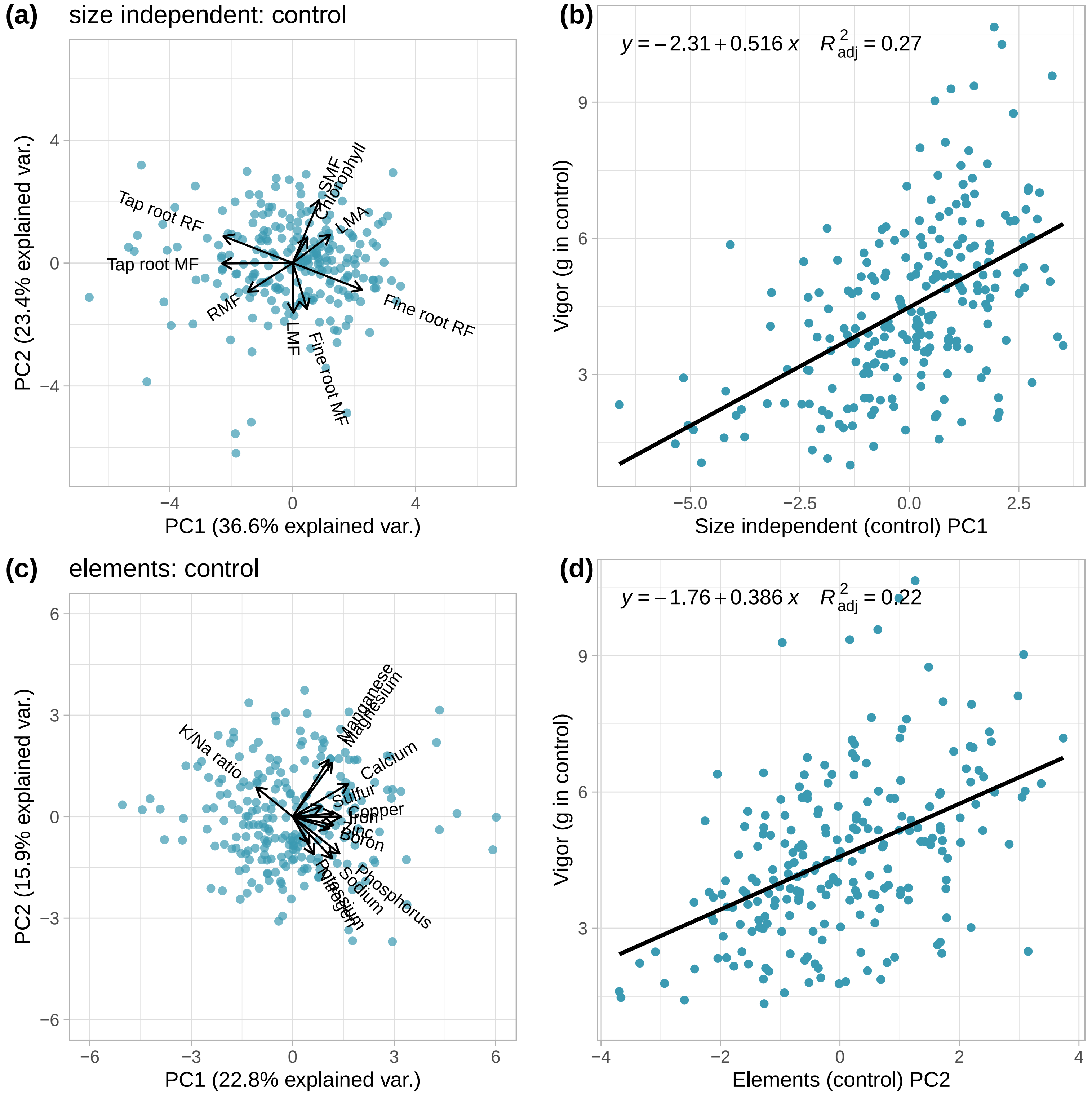

(a)
size independent: control
PC1 (36.6% explained var.)
PC2 (23.4% explained var.)
Tap root RF
Tap root MF
RMF
LMF
Fine root MF
Fine root RF
LMA
Chlorophyll
SMF
(b)
y = −2.31 + 0.516 x R²adj = 0.27
Size independent (control) PC1
Vigor (g in control)
(c)
elements: control
PC1 (22.8% explained var.)
PC2 (15.9% explained var.)
K/Na ratio
Manganese
Magnesium
Calcium
Sulfur
Copper
Iron
Zinc
Boron
Phosphorus
Sodium
Potassium
Nitrogen
(d)
y = −1.76 + 0.386 x R²adj = 0.22
Elements (control) PC2
Vigor (g in control)

**Figure S4. Linkage disequilibrium heatmaps for significant SNPs on each chromosome.** Linkage disequilibrium (estimated as $R^2$) between all SNPs that are significant for at least one trait within and between both treatments. Colored bar notes the haplotype blocks SNPs belong to based on the haplotype map presented in **Fig S3**. Block membership is indicated for the calculation based on the genome-wide collection of SNPs (“genome”) and for the re-calculation based on just the significant SNPs (“significant”).

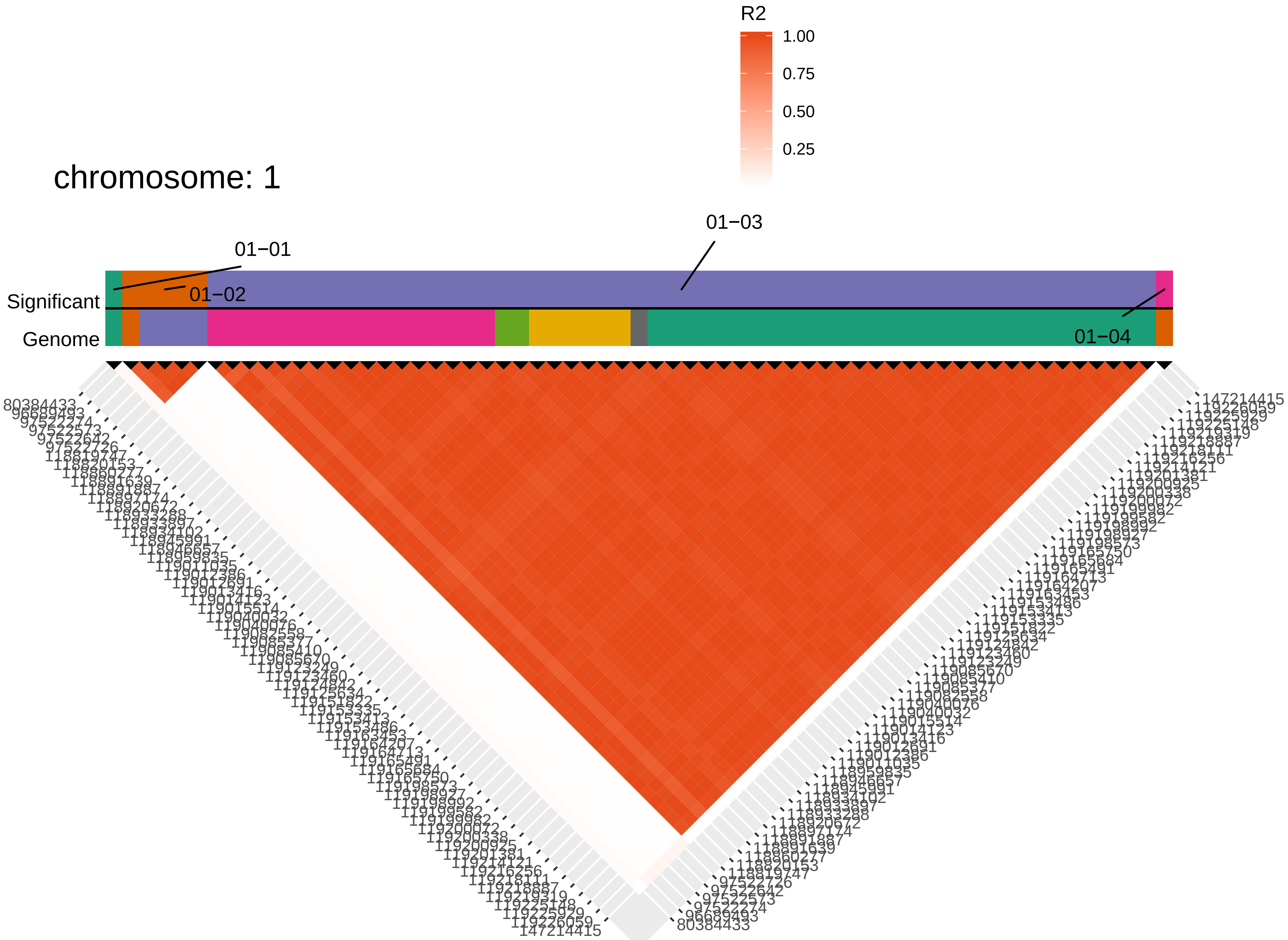
chromosome: 1
R2
1.00
0.75
0.50
0.25
01−01
01−02
01−03
01−04
Significant
Genome
80384433
147214415

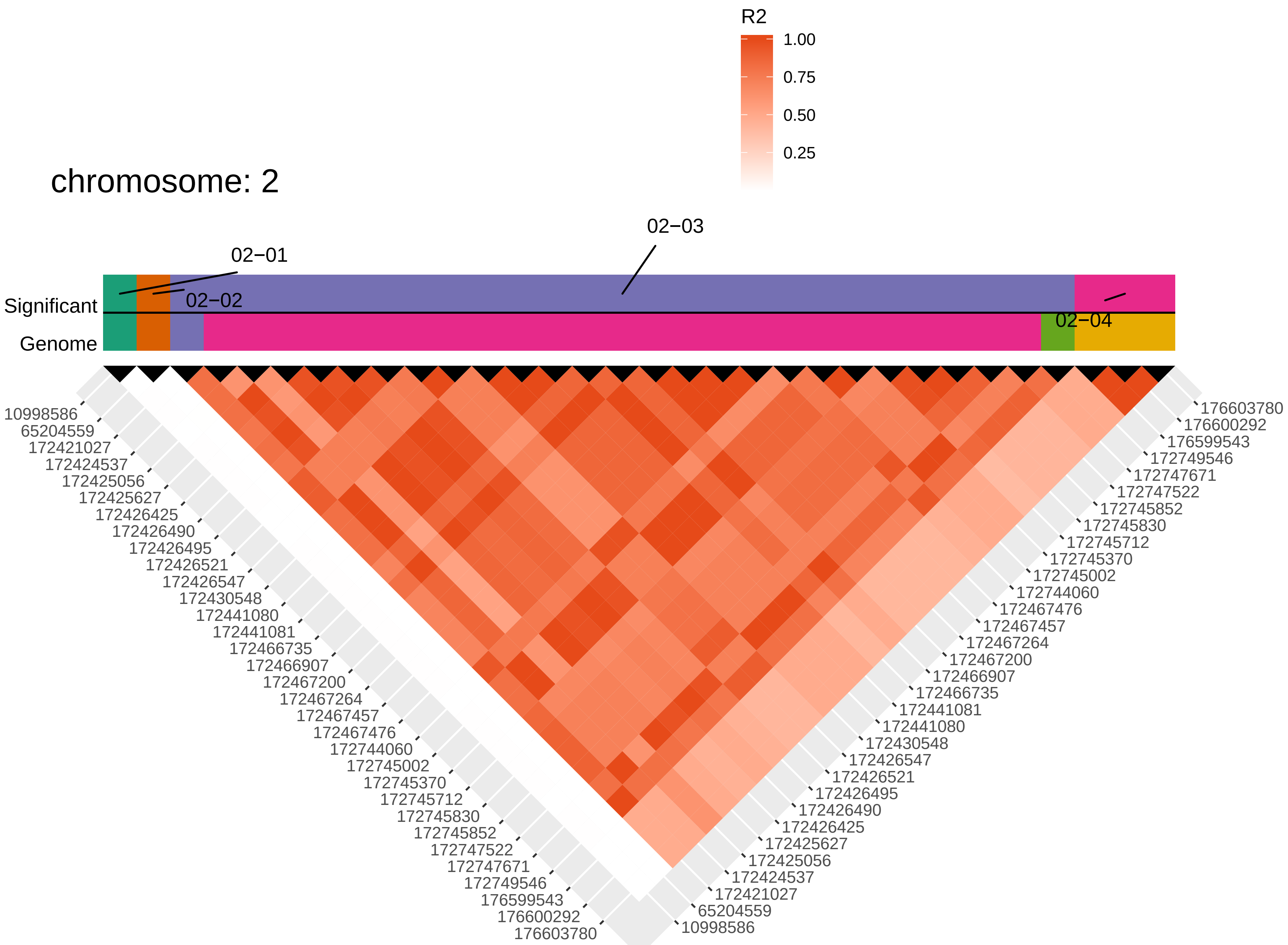
chromosome: 2
R2
1.00
0.75
0.50
0.25
02−01
02−02
02−03
02−04
Significant
Genome
10998586
65204559
172421027
172424537
172425056
172425627
172426425
172426490
172426495
172426521
172426547
172430548
172441080
172441081
172466735
172466907
172467200
172467264
172467457
172467476
172744060
172745002
172745370
172745712
172745830
172745852
172747522
172747671
172749546
176599543
176600292
176603780

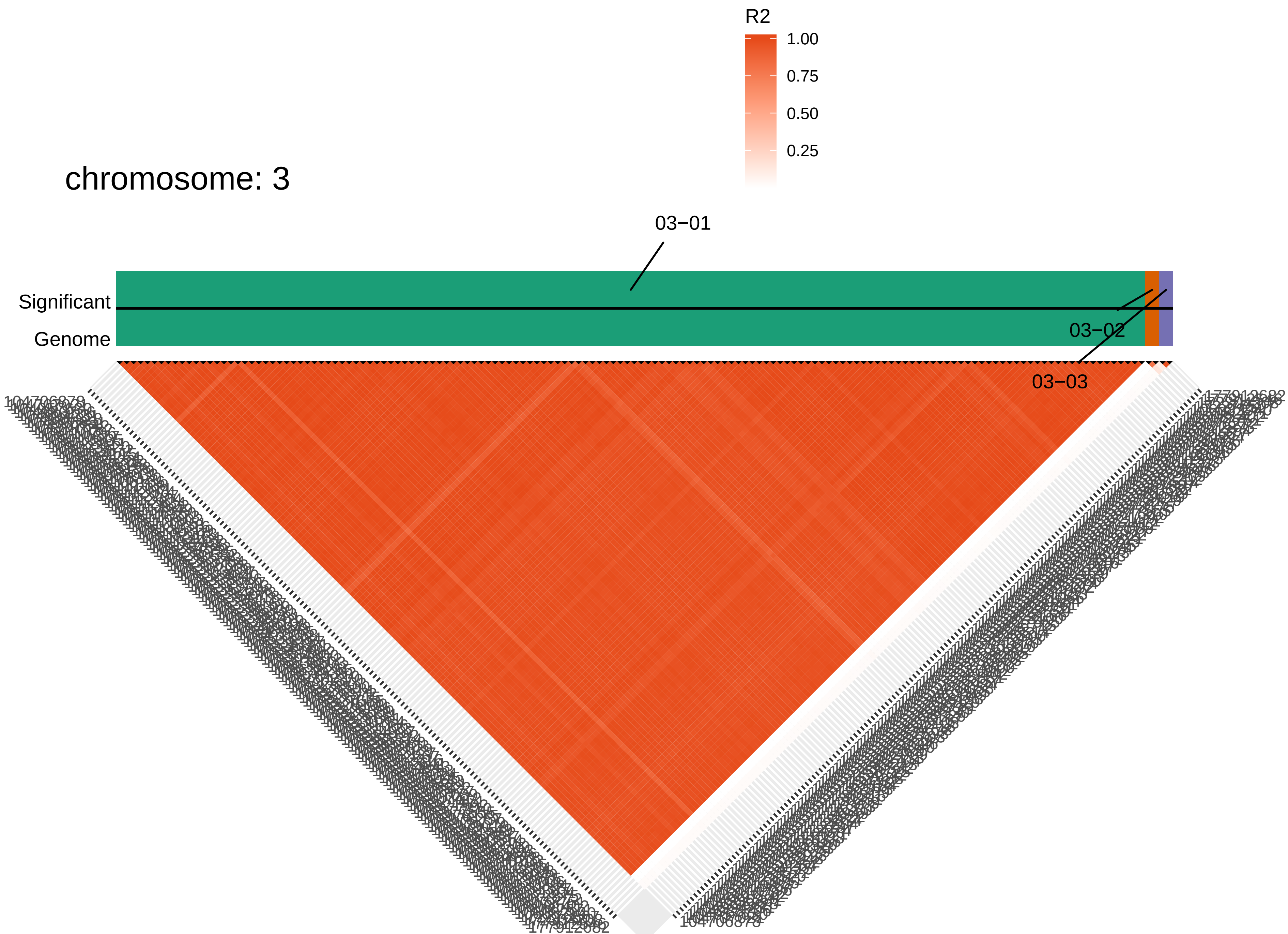
R2
1.00
0.75
0.50
0.25
chromosome: 3
03−01
Significant
Genome
03−02
03−03

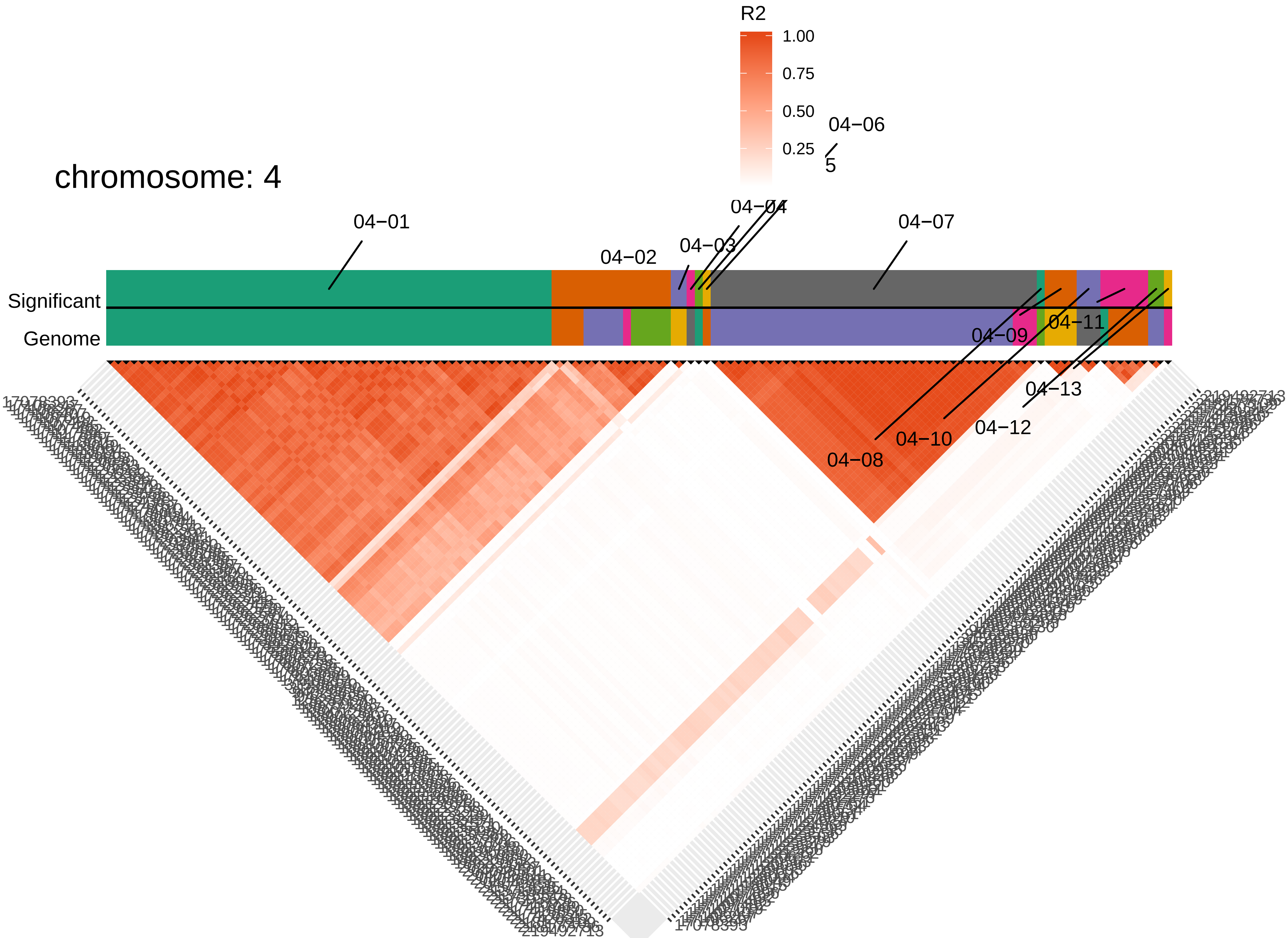
R2
1.00
0.75
0.50
0.25
chromosome: 4
04−01
04−02
04−03
04−06
04−07
04−08
04−09
04−10
04−11
04−12
04−13
Significant
Genome
219492713
17078393

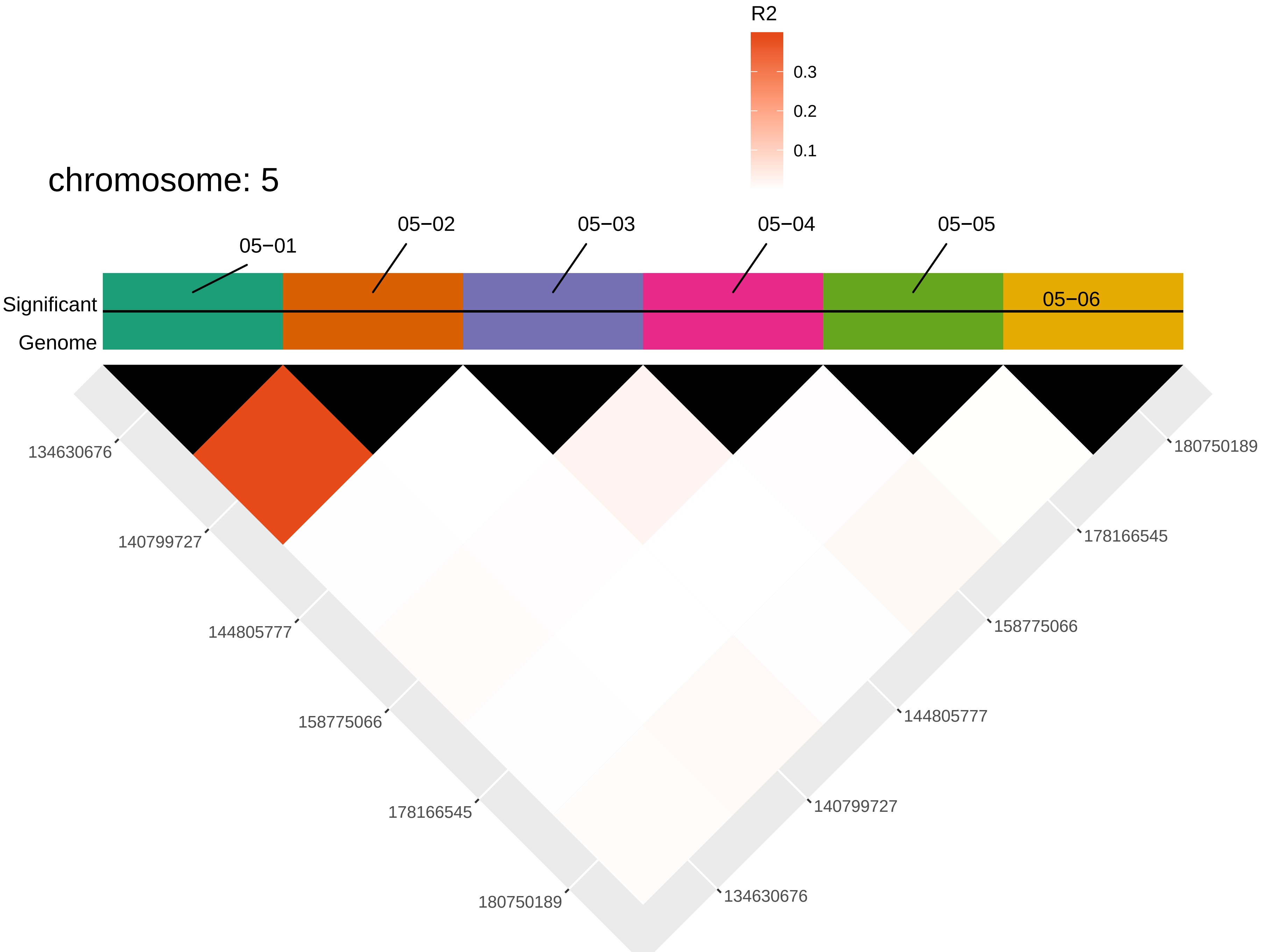
R2
0.3
0.2
0.1
chromosome: 5
05−01
05−02
05−03
05−04
05−05
05−06
Significant
Genome
134630676
140799727
144805777
158775066
178166545
180750189
180750189
178166545
158775066
144805777
140799727
134630676

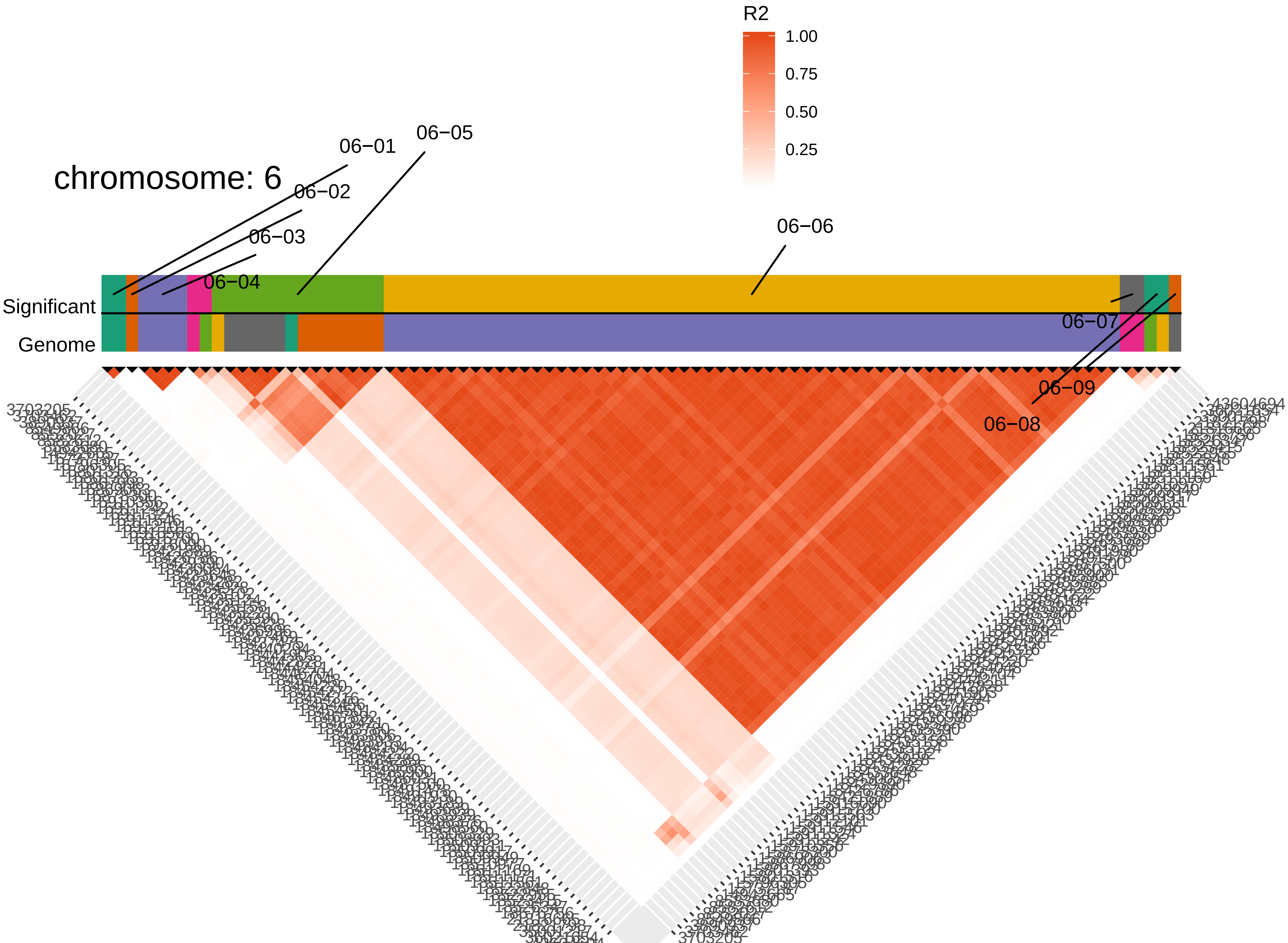
chromosome: 6
R2
1.00
0.75
0.50
0.25
06−01
06−02
06−03
06−04
06−05
06−06
06−07
06−08
06−09
Significant
Genome
3703205
43604694

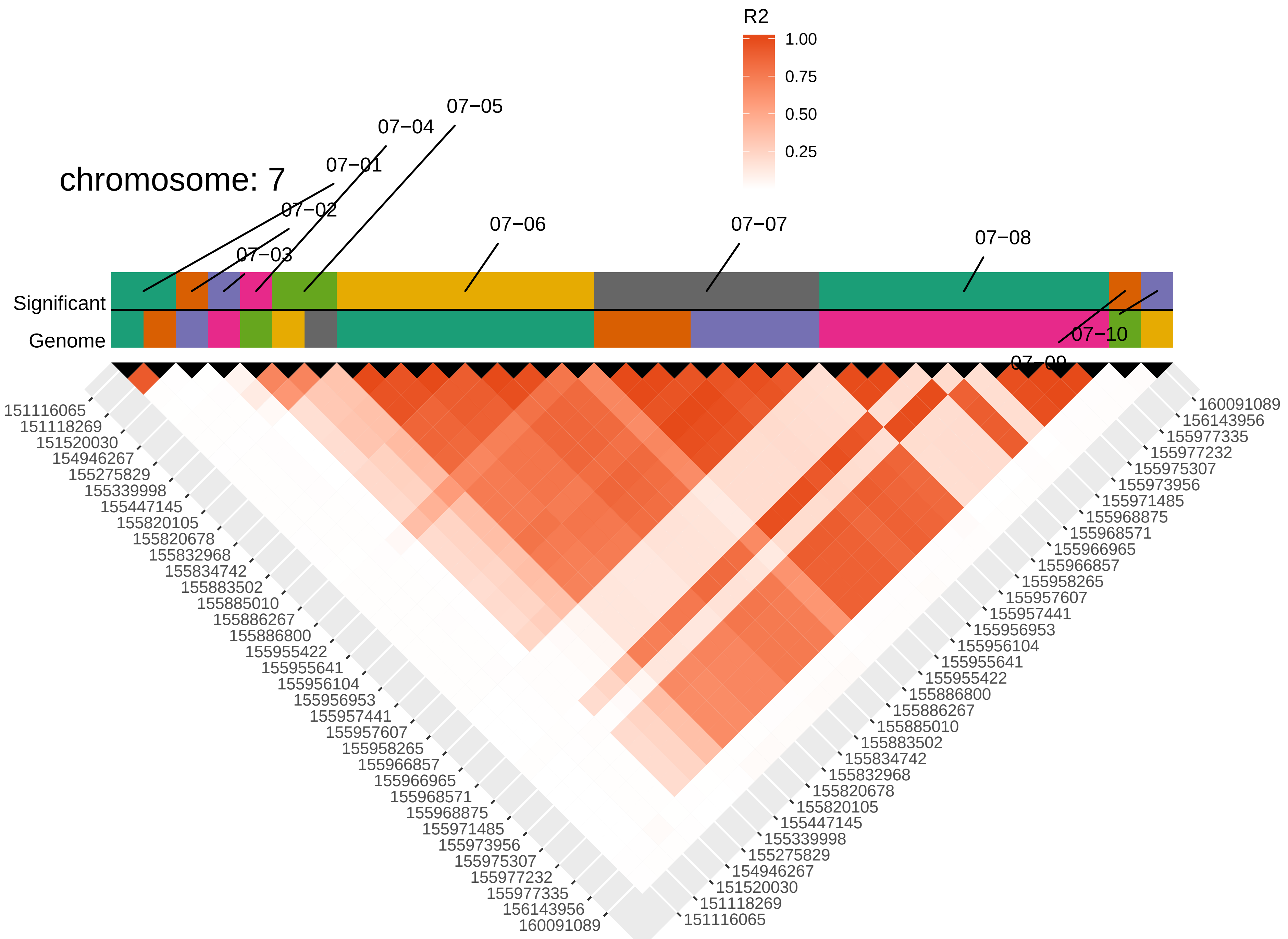
chromosome: 7
R2
1.00
0.75
0.50
0.25
07−01
07−02
07−03
07−04
07−05
07−06
07−07
07−08
07−09
07−10
Significant
Genome
151116065
151118269
151520030
154946267
155275829
155339998
155447145
155820105
155820678
155832968
155834742
155883502
155885010
155886267
155886800
155955422
155955641
155956104
155956953
155957441
155957607
155958265
155966857
155966965
155968571
155968875
155971485
155973956
155975307
155977232
155977335
156143956
160091089

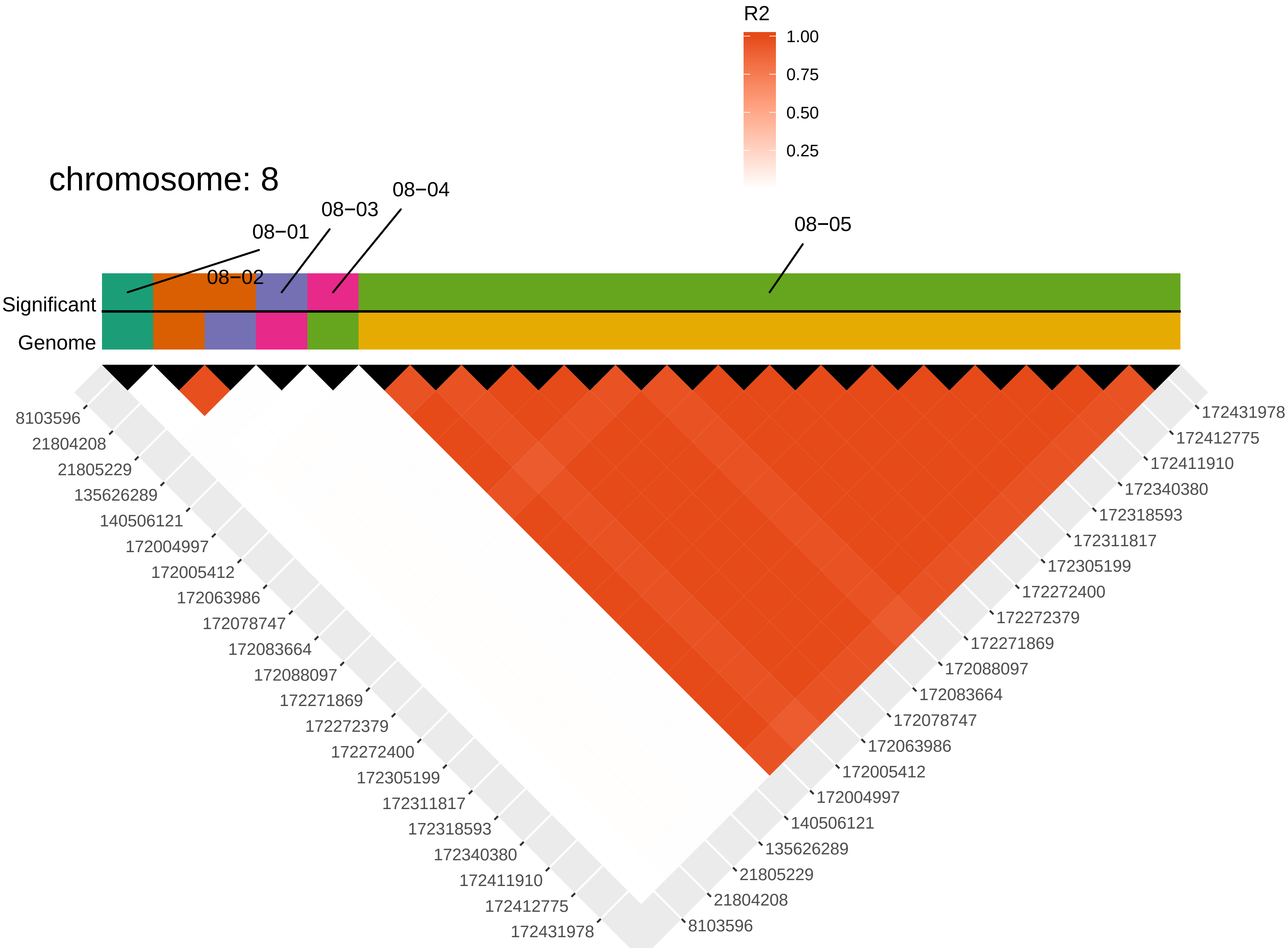
R2
1.00
0.75
0.50
0.25
chromosome: 8
08−04
08−03
08−01
08−05
08−02
Significant
Genome
8103596
21804208
21805229
135626289
140506121
172004997
172005412
172063986
172078747
172083664
172088097
172271869
172272379
172272400
172305199
172311817
172318593
172340380
172411910
172412775
172431978

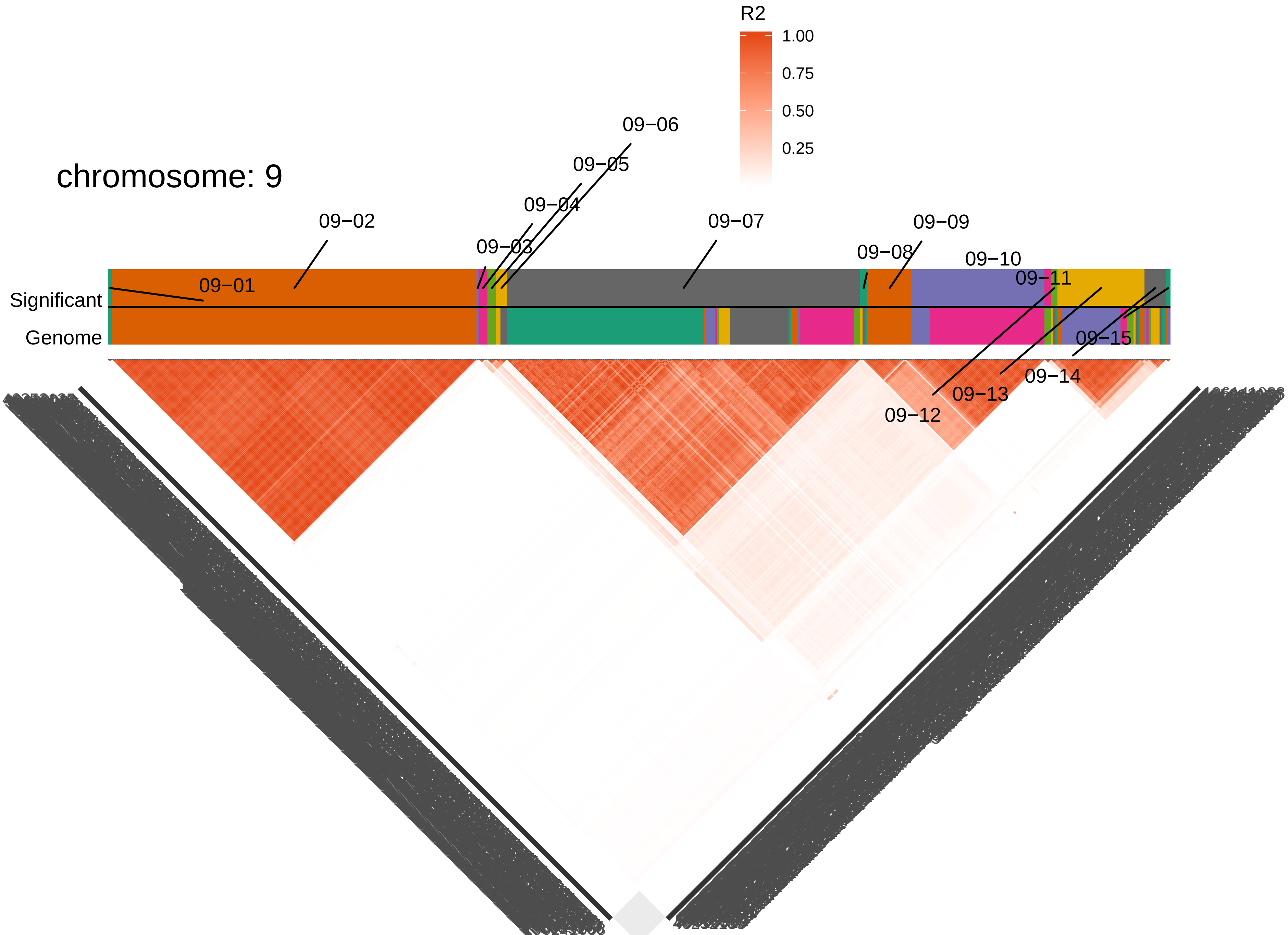
chromosome: 9
R2
1.00
0.75
0.50
0.25
Significant
Genome
09−01
09−02
09−03
09−04
09−05
09−06
09−07
09−08
09−09
09−10
09−11
09−12
09−13
09−14
09−15

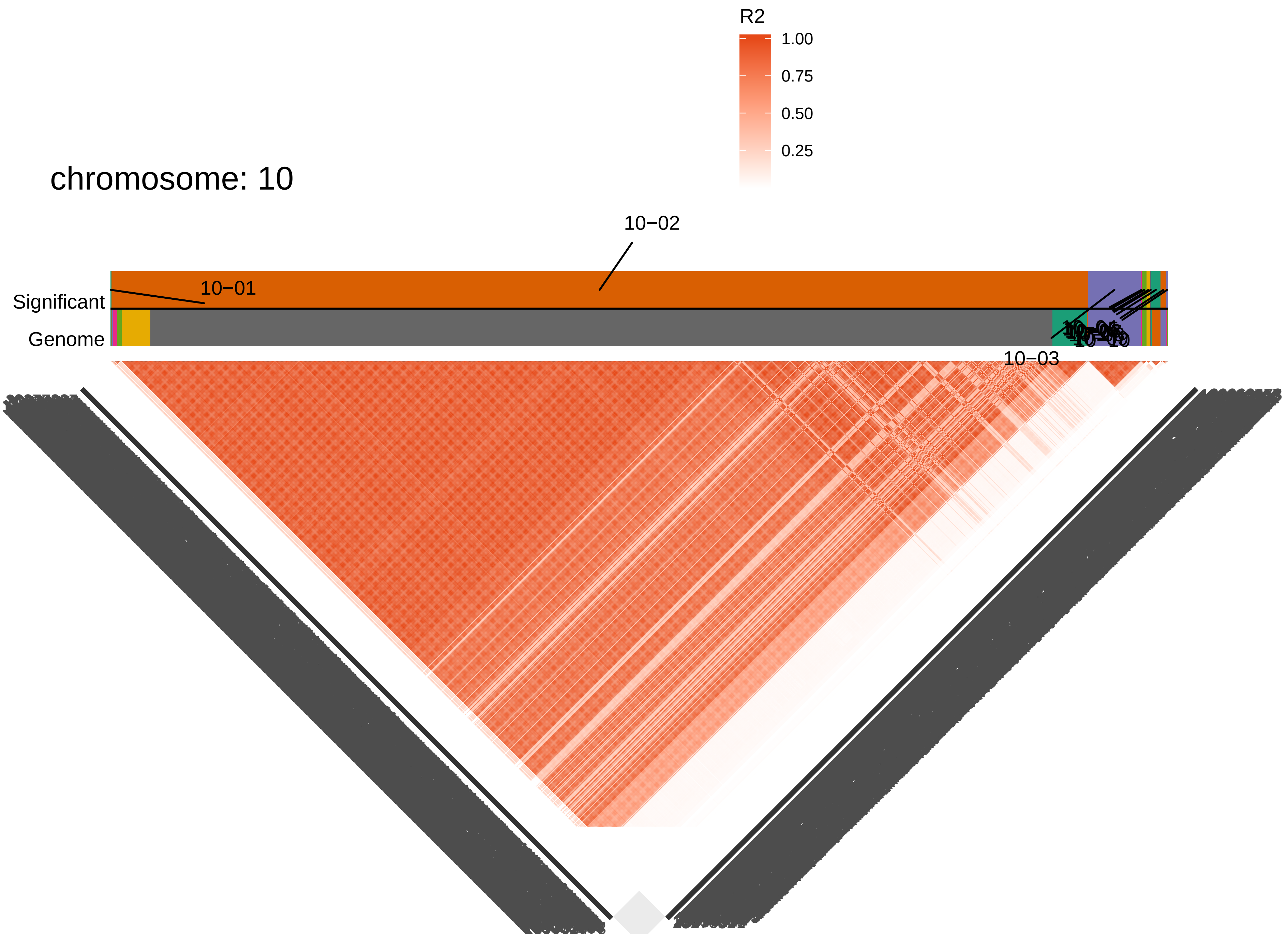

R2
1.00
0.75
0.50
0.25
chromosome: 10
10−02
10−01
Significant
Genome
10−03

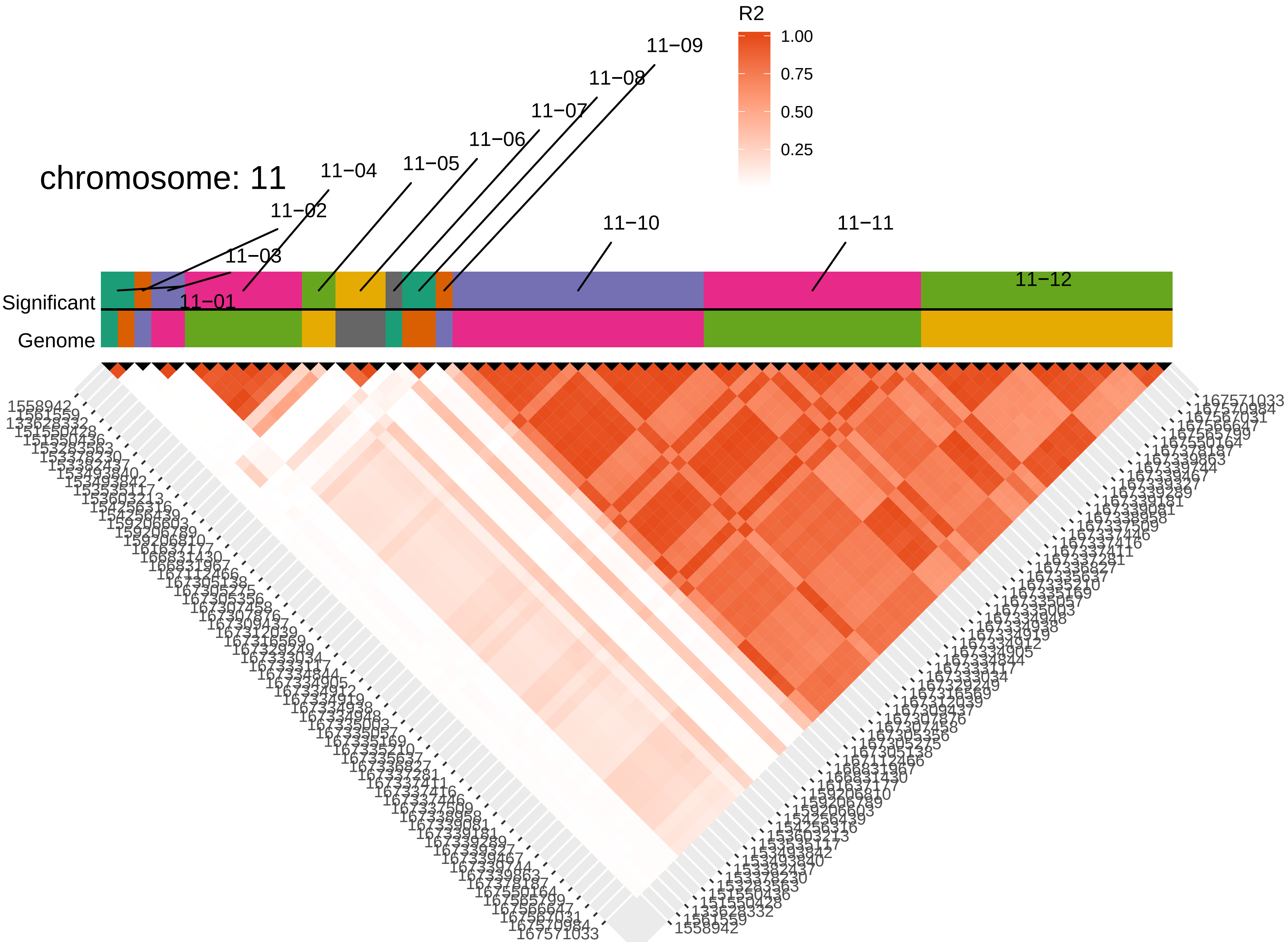
chromosome: 11
R2
1.00
0.75
0.50
0.25
11−01
11−02
11−03
11−04
11−05
11−06
11−07
11−08
11−09
11−10
11−11
11−12
Significant
Genome
1558942
167571033

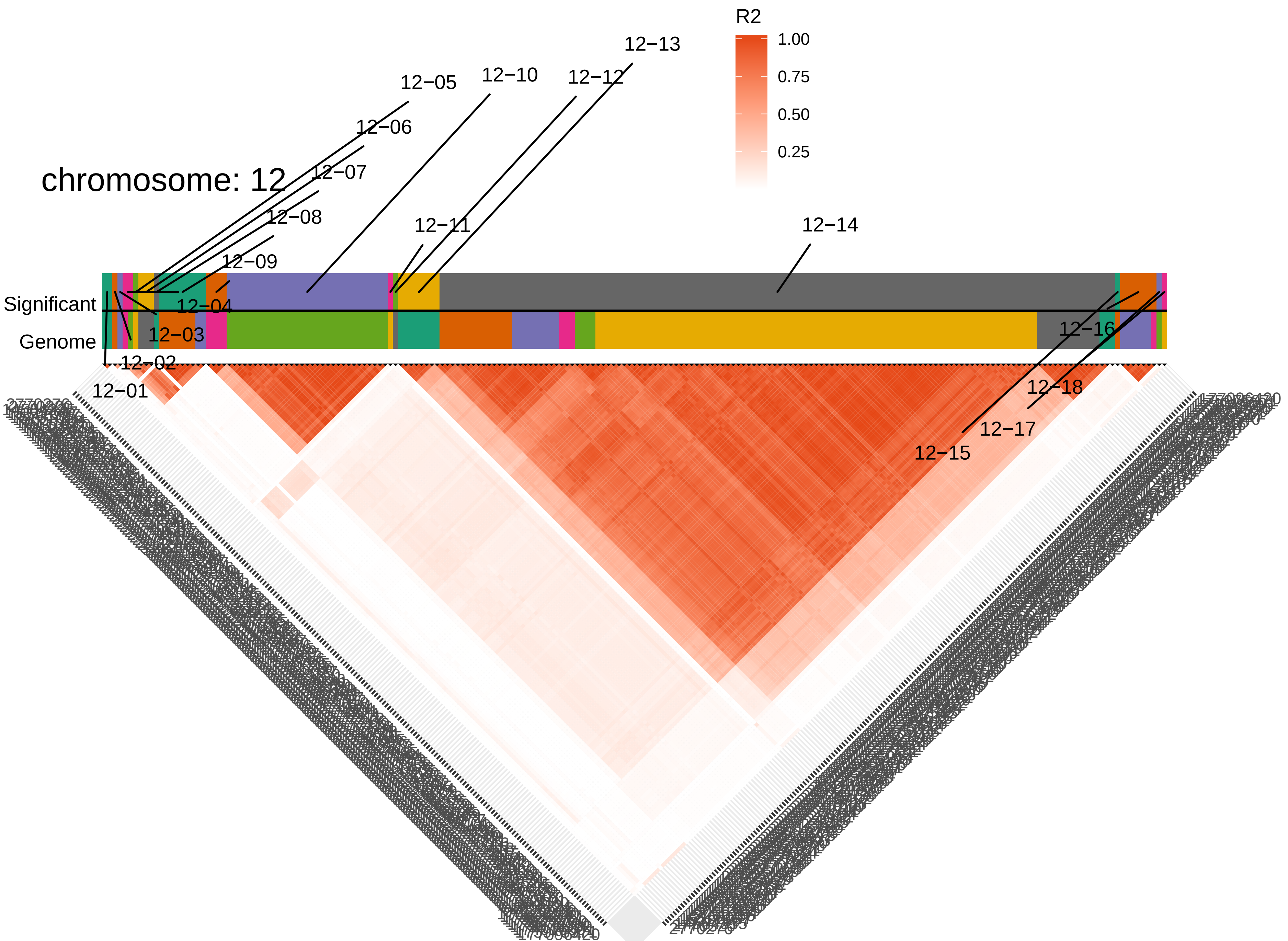
chromosome: 12
R2
1.00
0.75
0.50
0.25
Significant
Genome
12−01
12−02
12−03
12−04
12−05
12−06
12−07
12−08
12−09
12−10
12−11
12−12
12−13
12−14
12−15
12−16
12−17
12−18

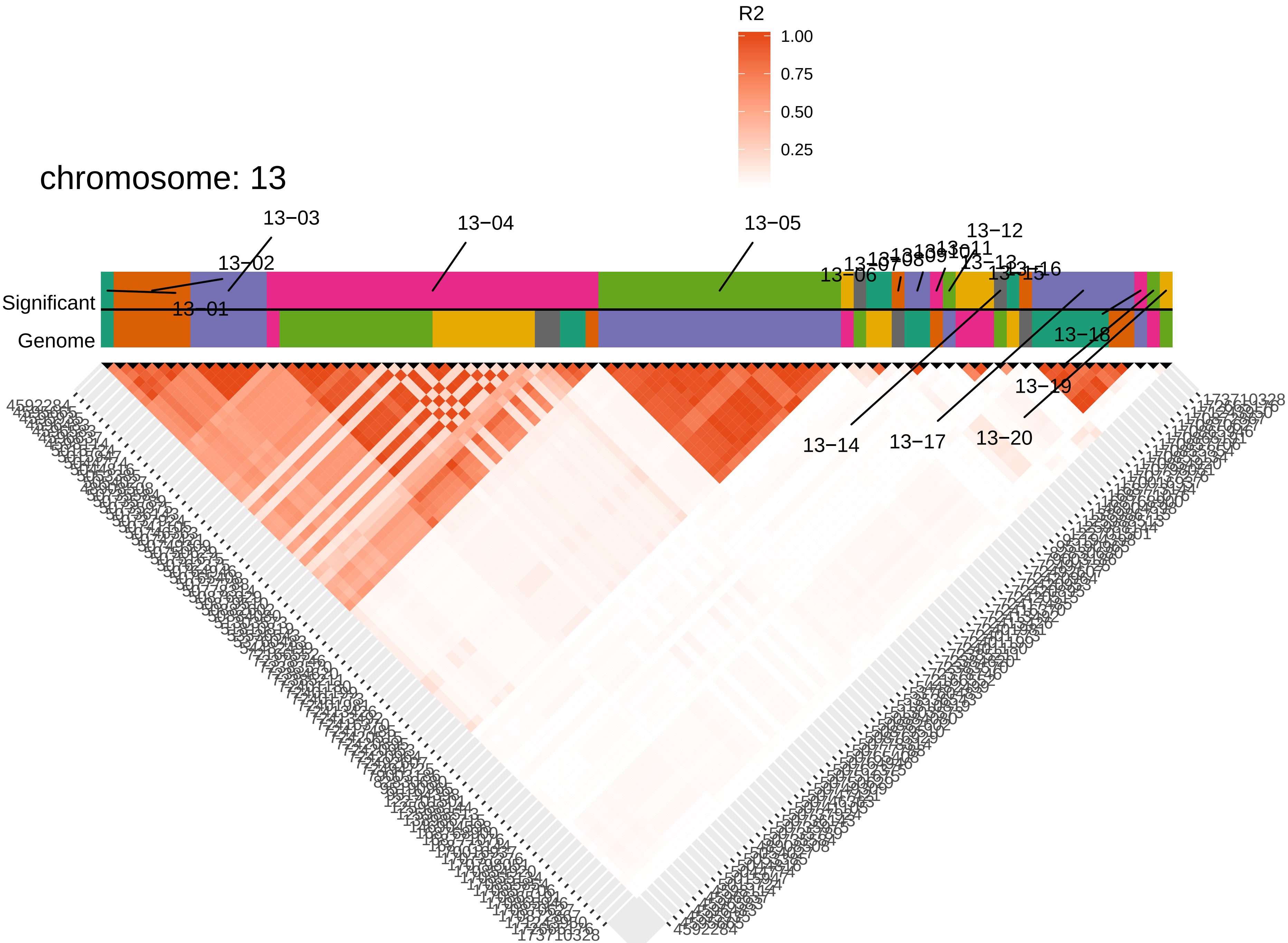

chromosome: 13
R2
1.00
0.75
0.50
0.25
Significant
Genome
13−01
13−02
13−03
13−04
13−05
13−06
13−12
13−14
13−17
13−18
13−19
13−20
4592284
173710328

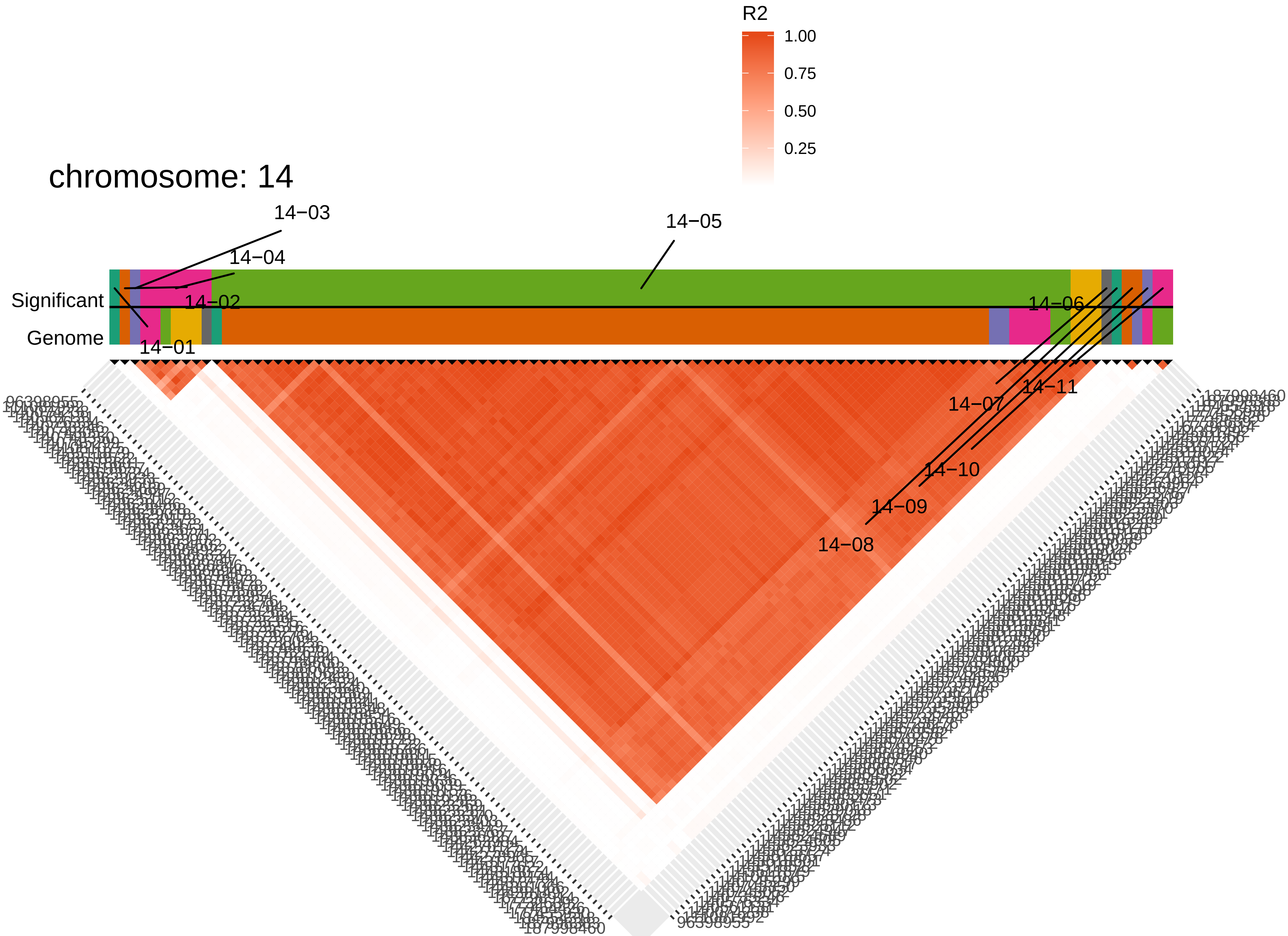
chromosome: 14
R2
1.00
0.75
0.50
0.25
14−01
14−02
14−03
14−04
14−05
14−06
14−07
14−08
14−09
14−10
14−11
Significant
Genome
187998460
96398955

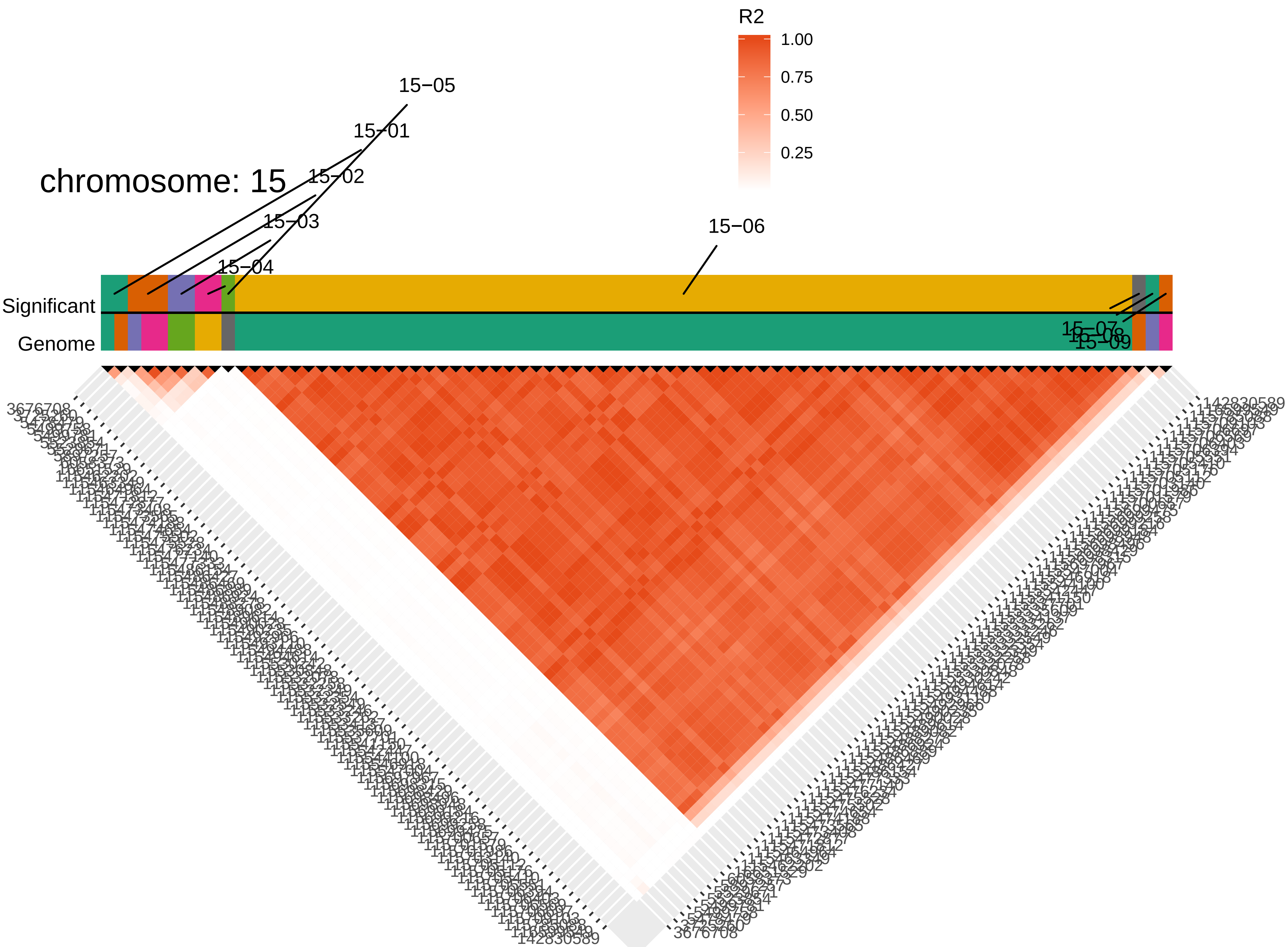

chromosome: 15
R2
1.00
0.75
0.50
0.25
15−05
15−01
15−02
15−03
15−04
15−06
Significant
Genome
15−07
15−09
3676708
142830589

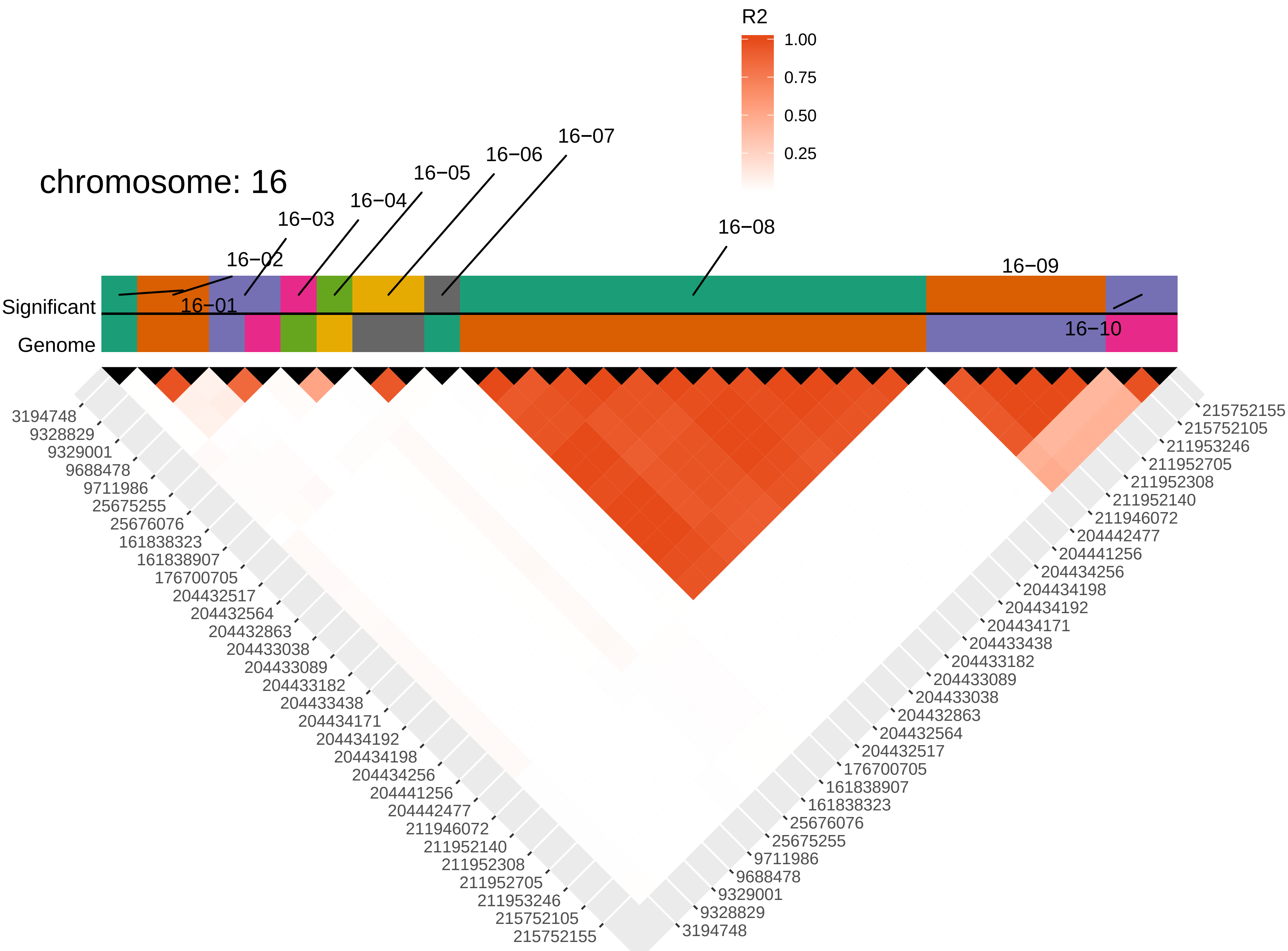
chromosome: 16
R2
1.00
0.75
0.50
0.25
16−01
16−02
16−03
16−04
16−05
16−06
16−07
16−08
16−09
16−10
Significant
Genome
3194748
9328829
9329001
9688478
9711986
25675255
25676076
161838323
161838907
176700705
204432517
204432564
204432863
204433038
204433089
204433182
204433438
204434171
204434192
204434198
204434256
204441256
204442477
211946072
211952140
211952308
211952705
211953246
215752105
215752155

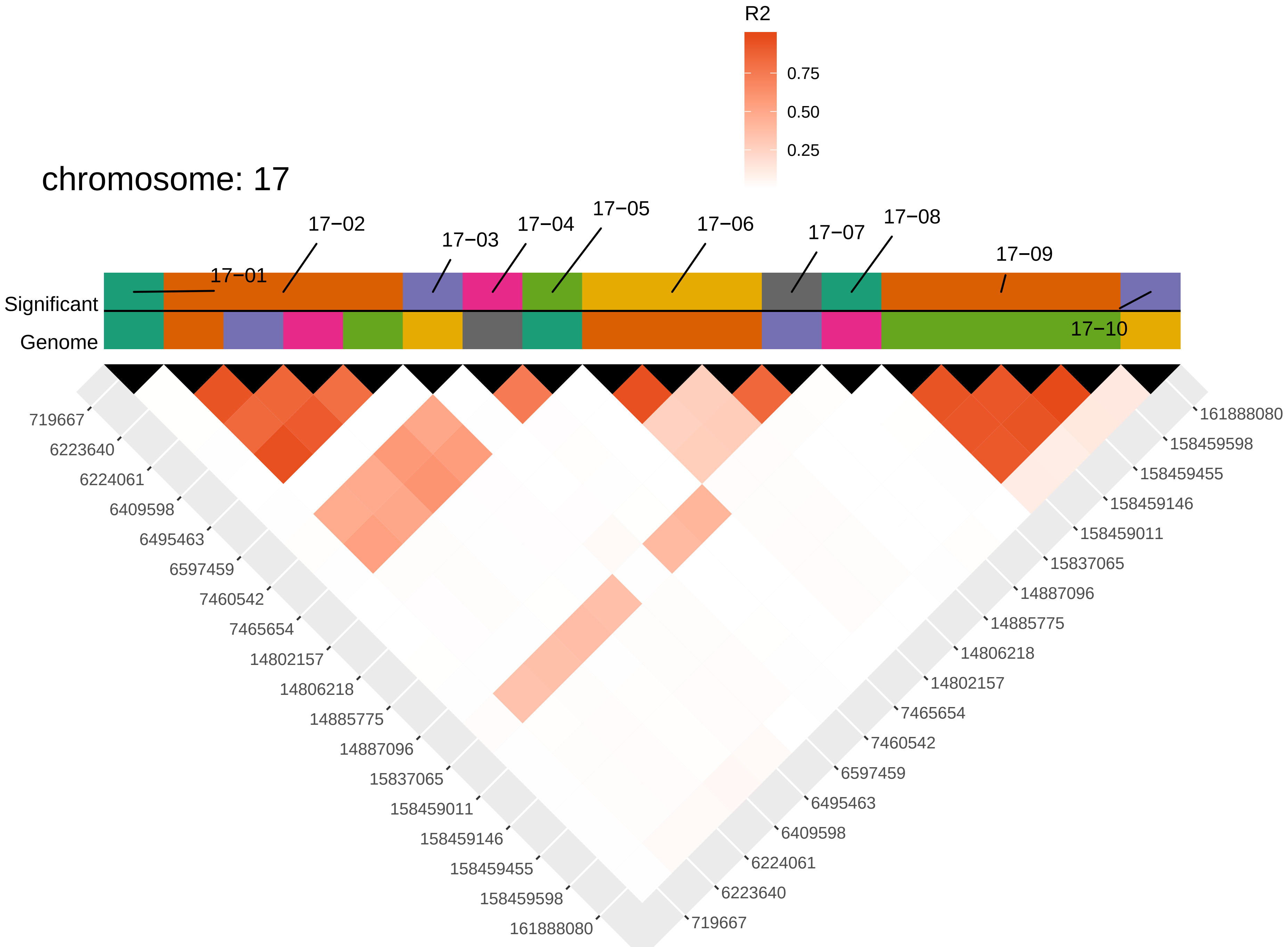

chromosome: 17
R2
0.75
0.50
0.25
17−01
17−02
17−03
17−04
17−05
17−06
17−07
17−08
17−09
17−10
Significant
Genome
719667
6223640
6224061
6409598
6495463
6597459
7460542
7465654
14802157
14806218
14885775
14887096
15837065
158459011
158459146
158459455
158459598
161888080

**Figure S5. Manhattan plots of all traits.** Per-trait Manhattan plots are shown for trait values under control and salt-stressed conditions, as well as the plasticity between treatments (log difference). SNPs above the red line are significant after multiple comparison correction, SNPs above the blue line are in the top 0.1% of *P*-values. SNPs are colored by the “significant” haplotype regions per chromosome as displayed in **Fig S3**.

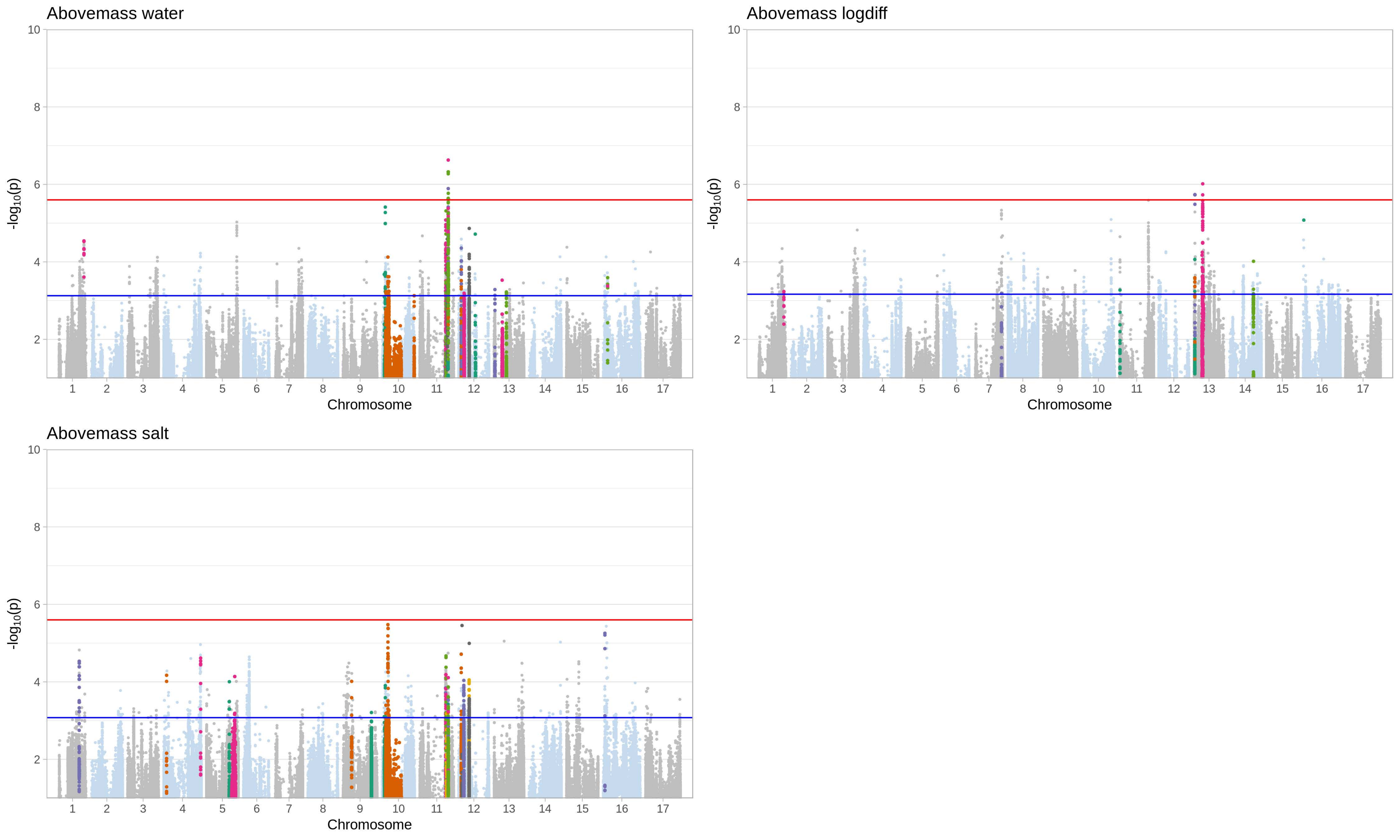
Abovemass water
Abovemass logdiff
Abovemass salt
-log10(p)
Chromosome
1 2 3 4 5 6 7 8 9 10 11 12 13 14 15 16 17
2 4 6 8 10

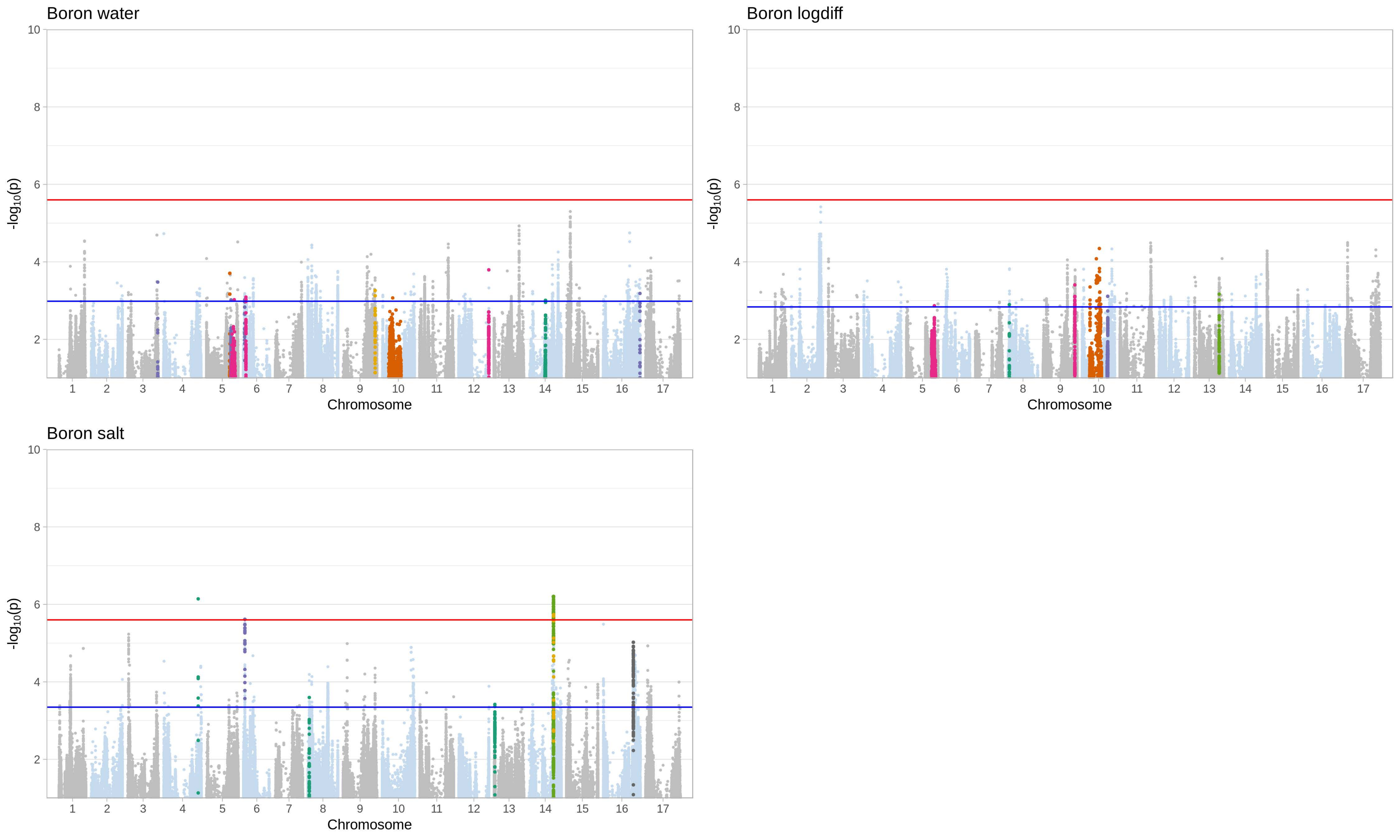
Boron water
Boron logdiff
Boron salt
-log10(p)
Chromosome
0
2
4
6
8
10

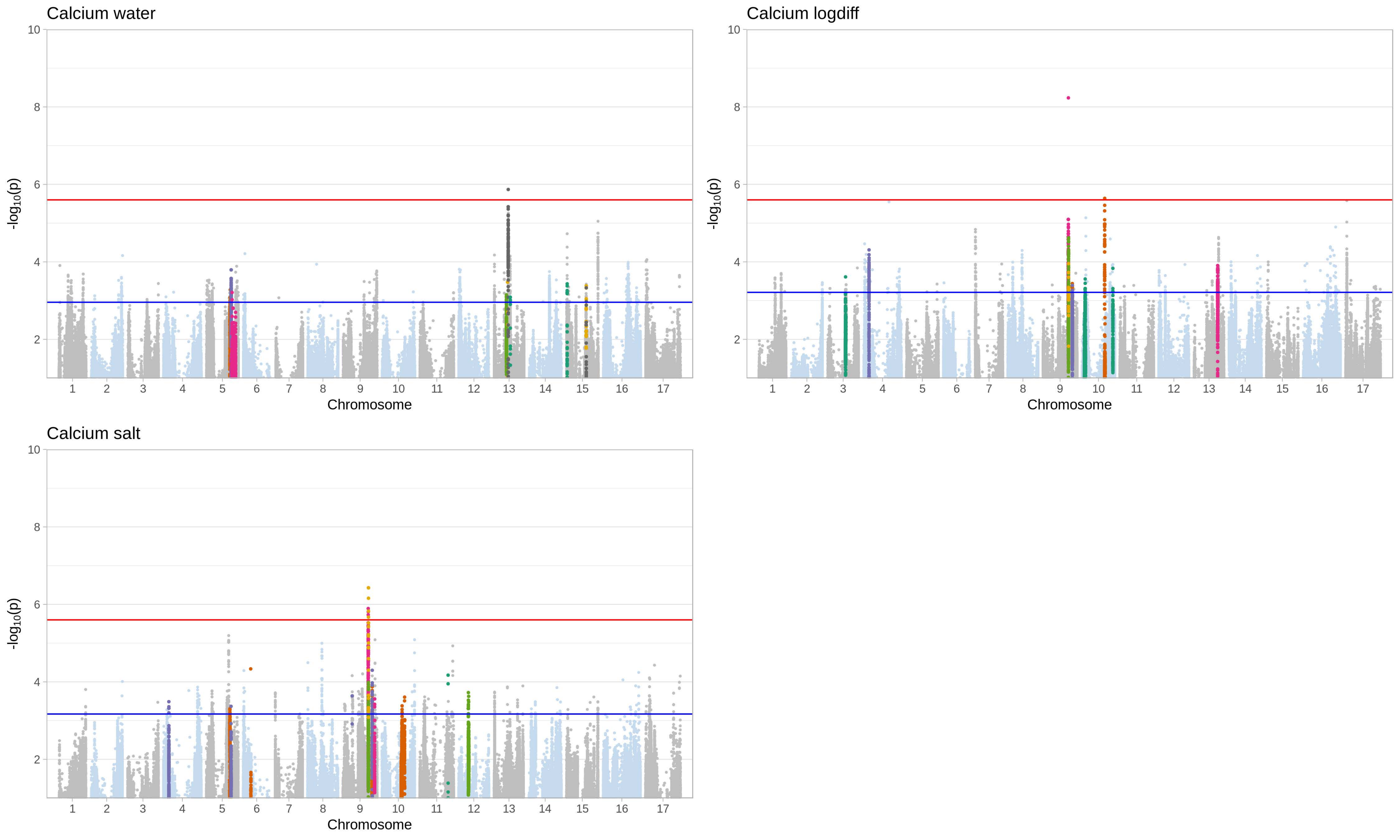
Calcium water
Calcium logdiff
Calcium salt
$-\log_{10}(p)$
Chromosome
10
8
6
4
2
1 2 3 4 5 6 7 8 9 10 11 12 13 14 15 16 17

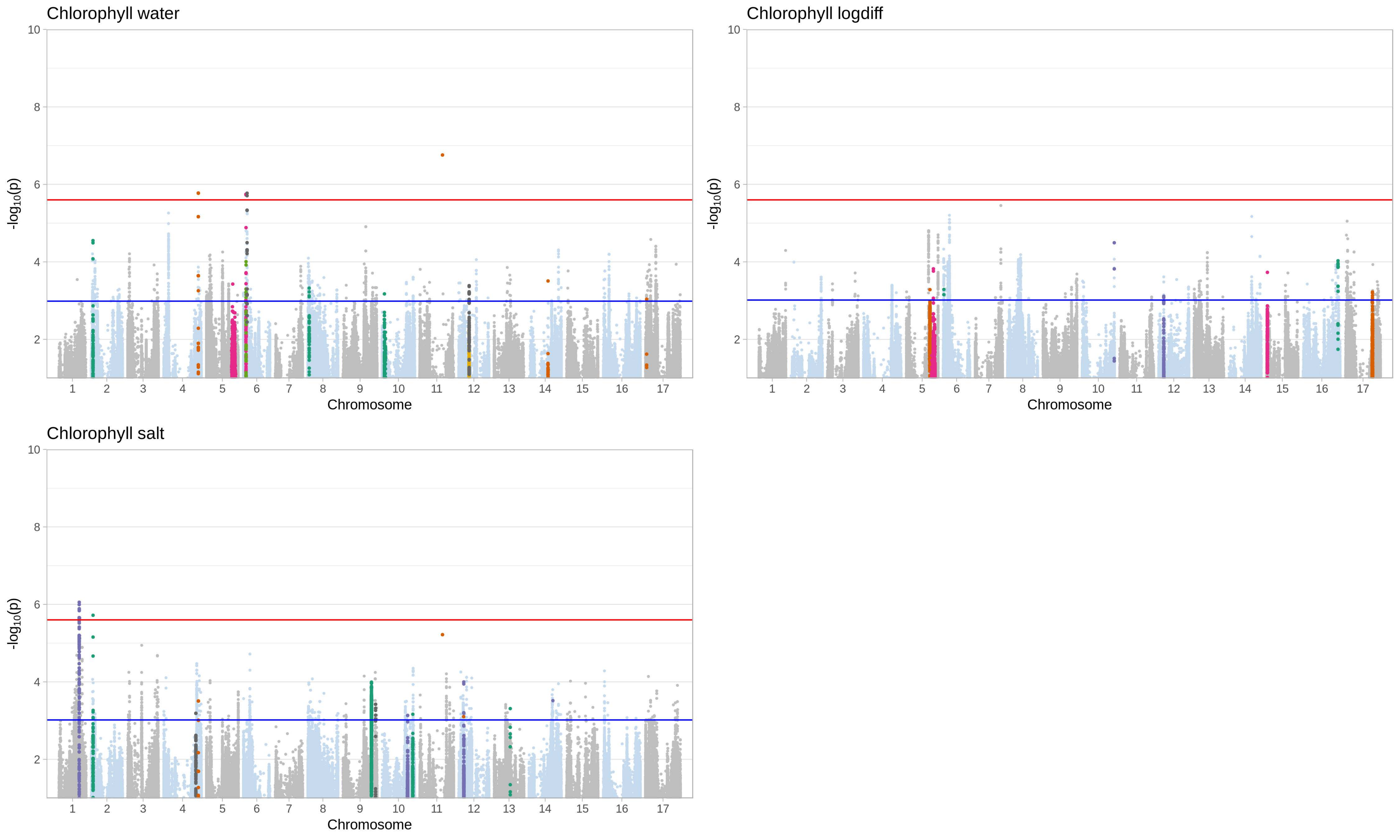
Chlorophyll water
Chlorophyll logdiff
Chlorophyll salt
-log10(p)
Chromosome
0
2
4
6
8
10

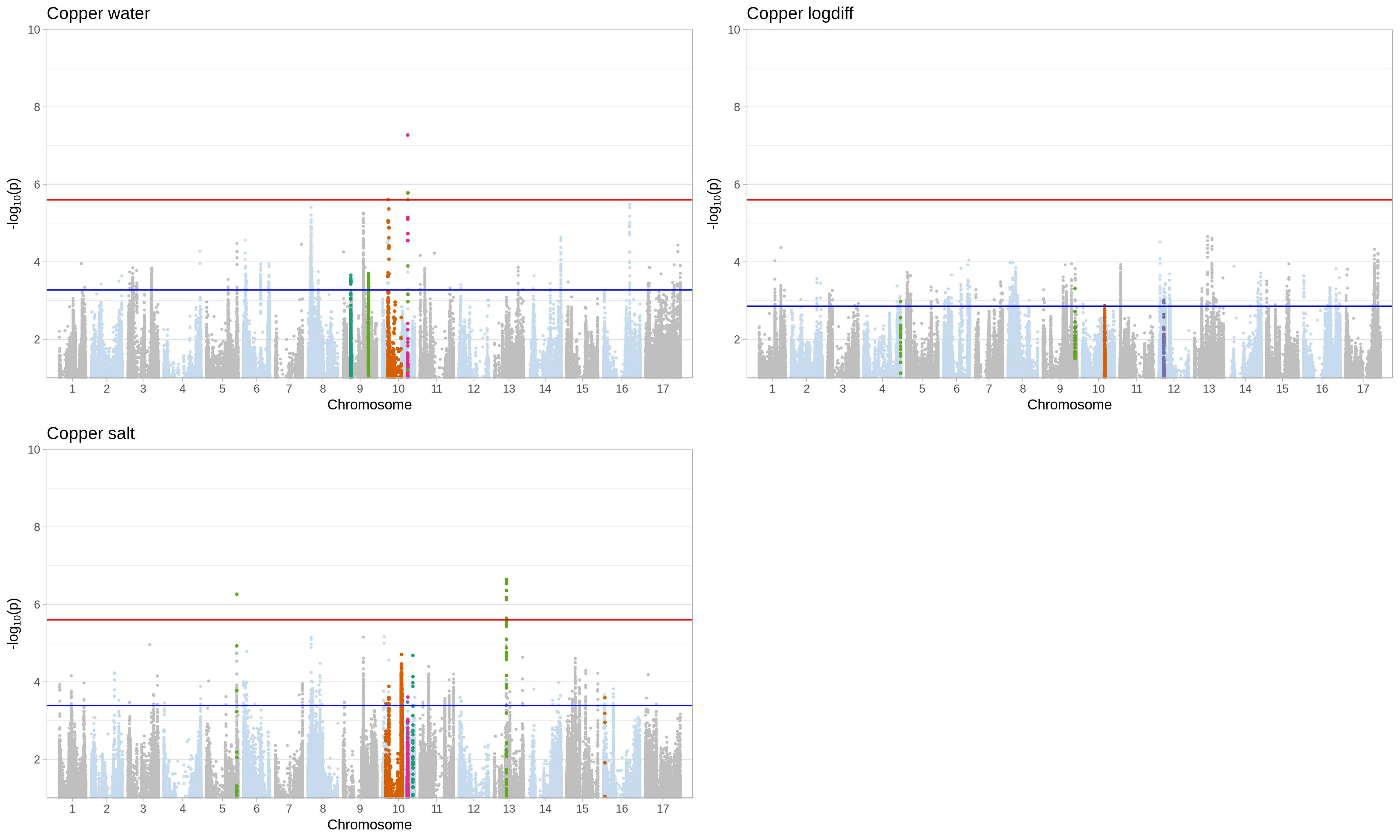

Copper water
Copper logdiff
Copper salt
-log10(p)
Chromosome
1 2 3 4 5 6 7 8 9 10 11 12 13 14 15 16 17
2 4 6 8 10

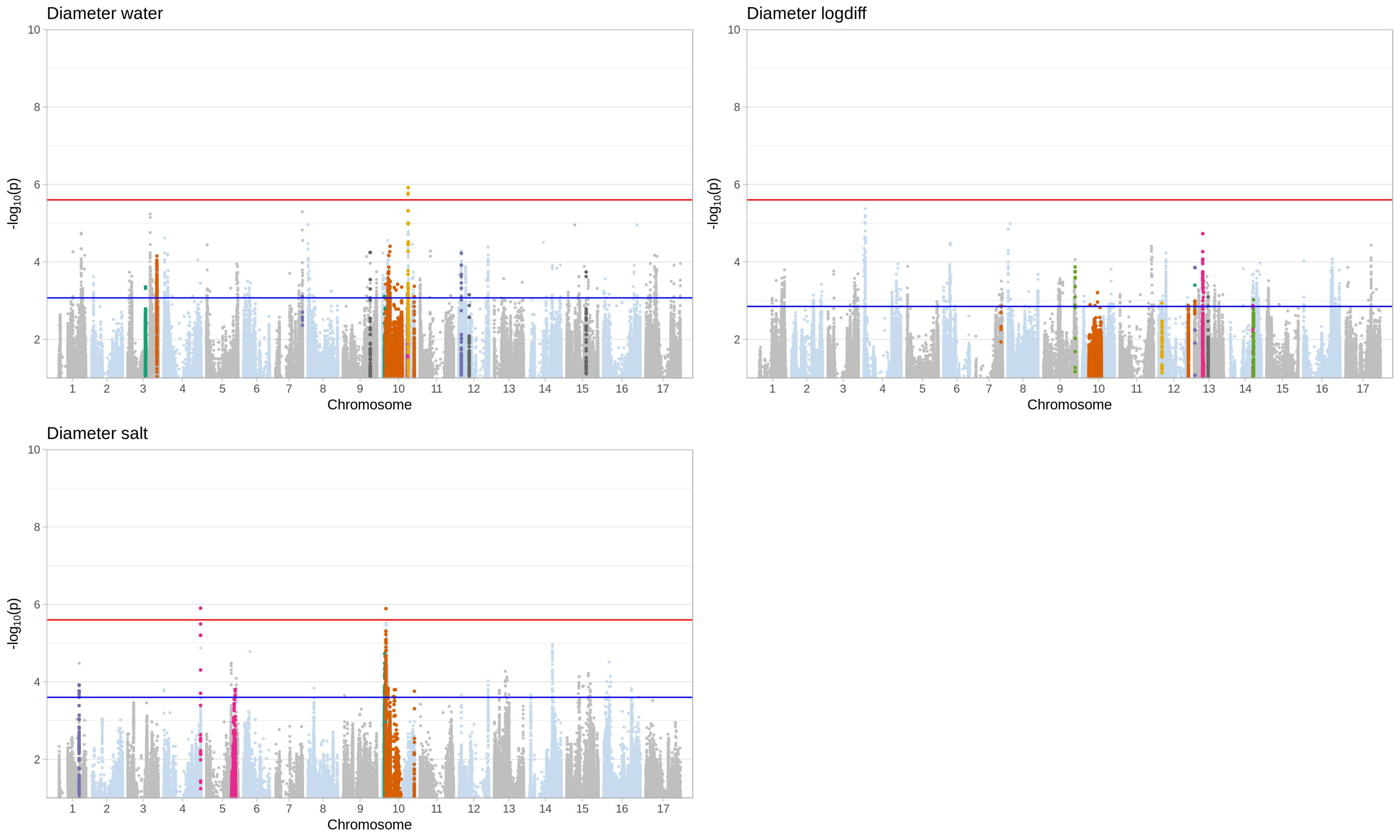

Diameter water
Diameter logdiff
Diameter salt
-log10(p)
Chromosome

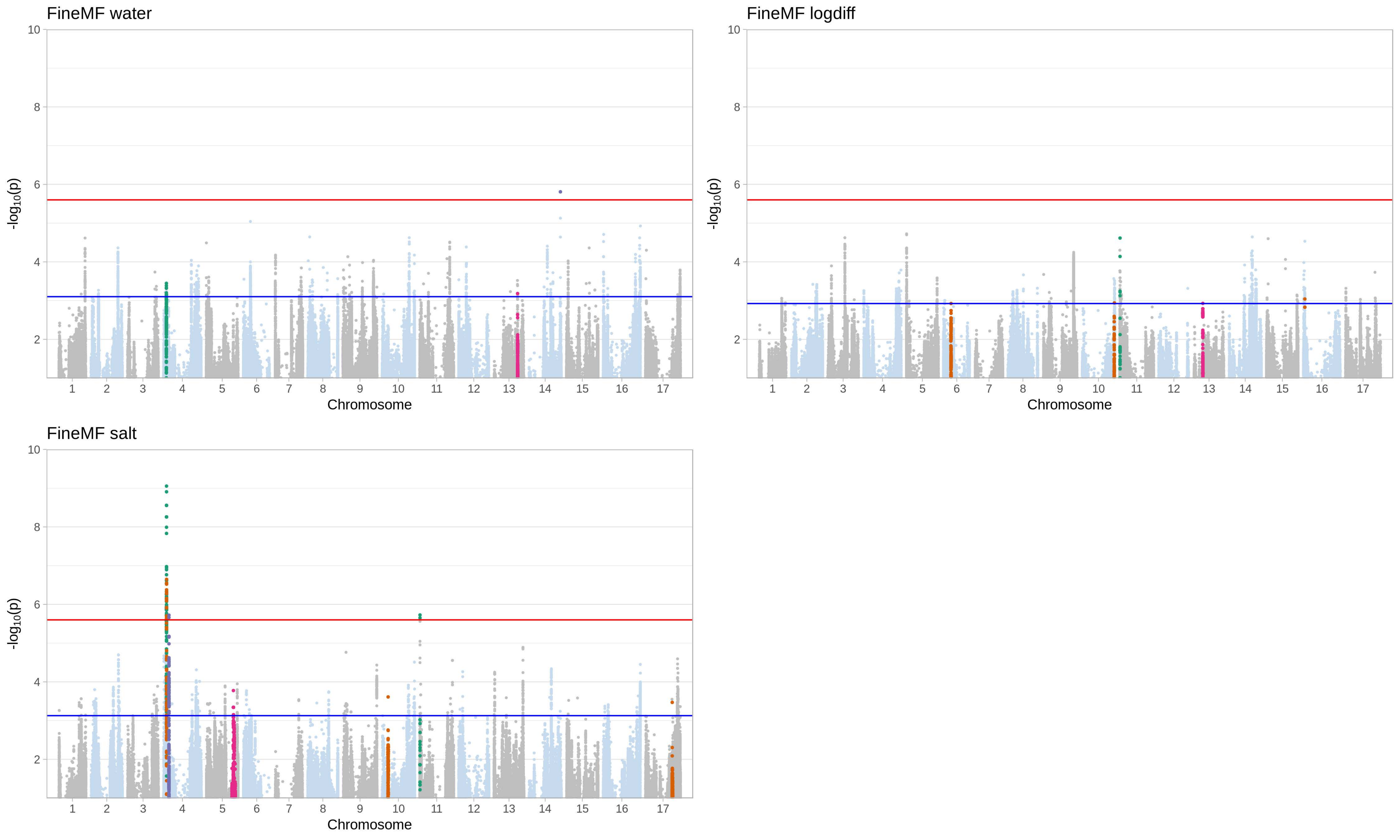
FineMF water
FineMF logdiff
FineMF salt
-log10(p)
Chromosome
1 2 3 4 5 6 7 8 9 10 11 12 13 14 15 16 17
2 4 6 8 10

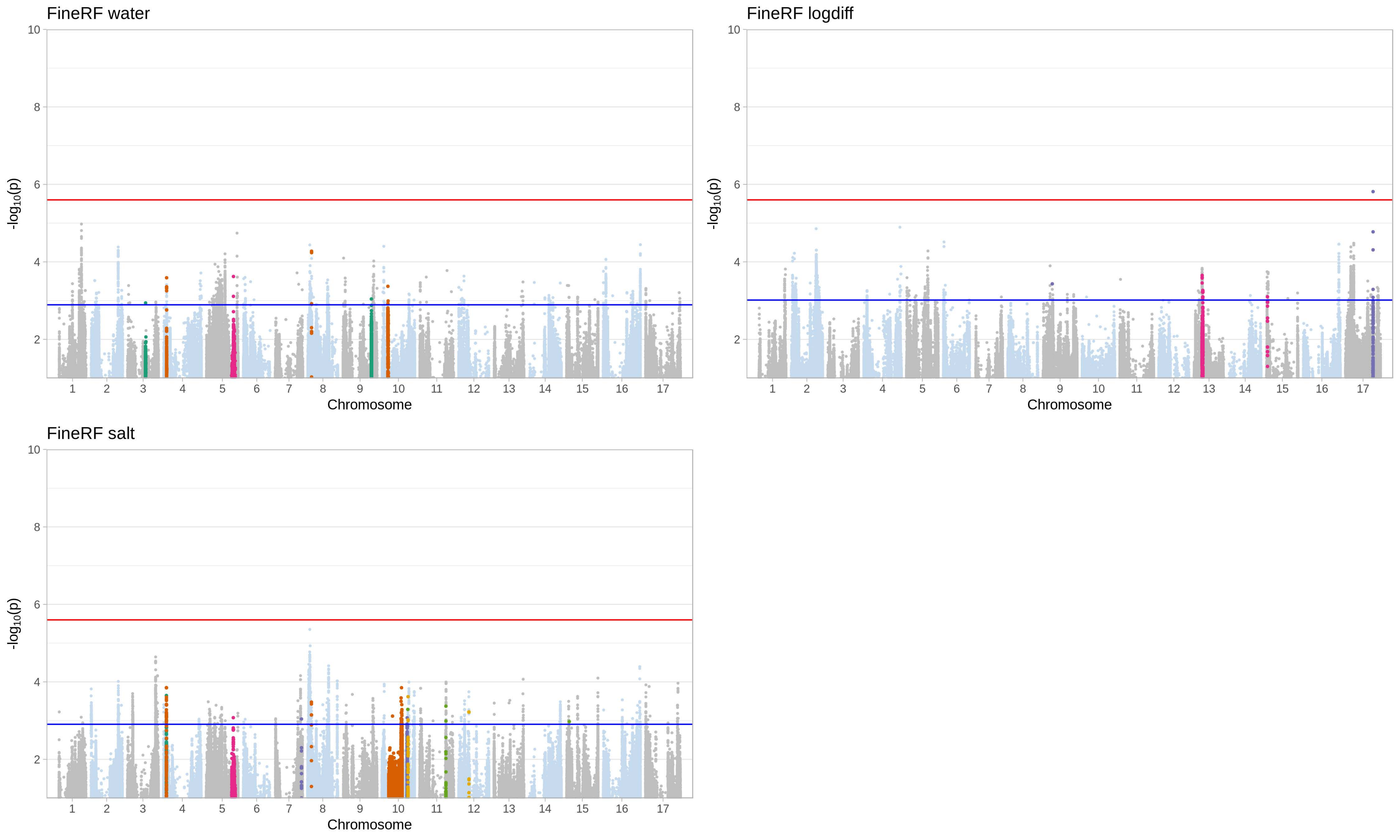
FineRF water
FineRF logdiff
FineRF salt
-log10(p)
10
8
6
4
2
Chromosome

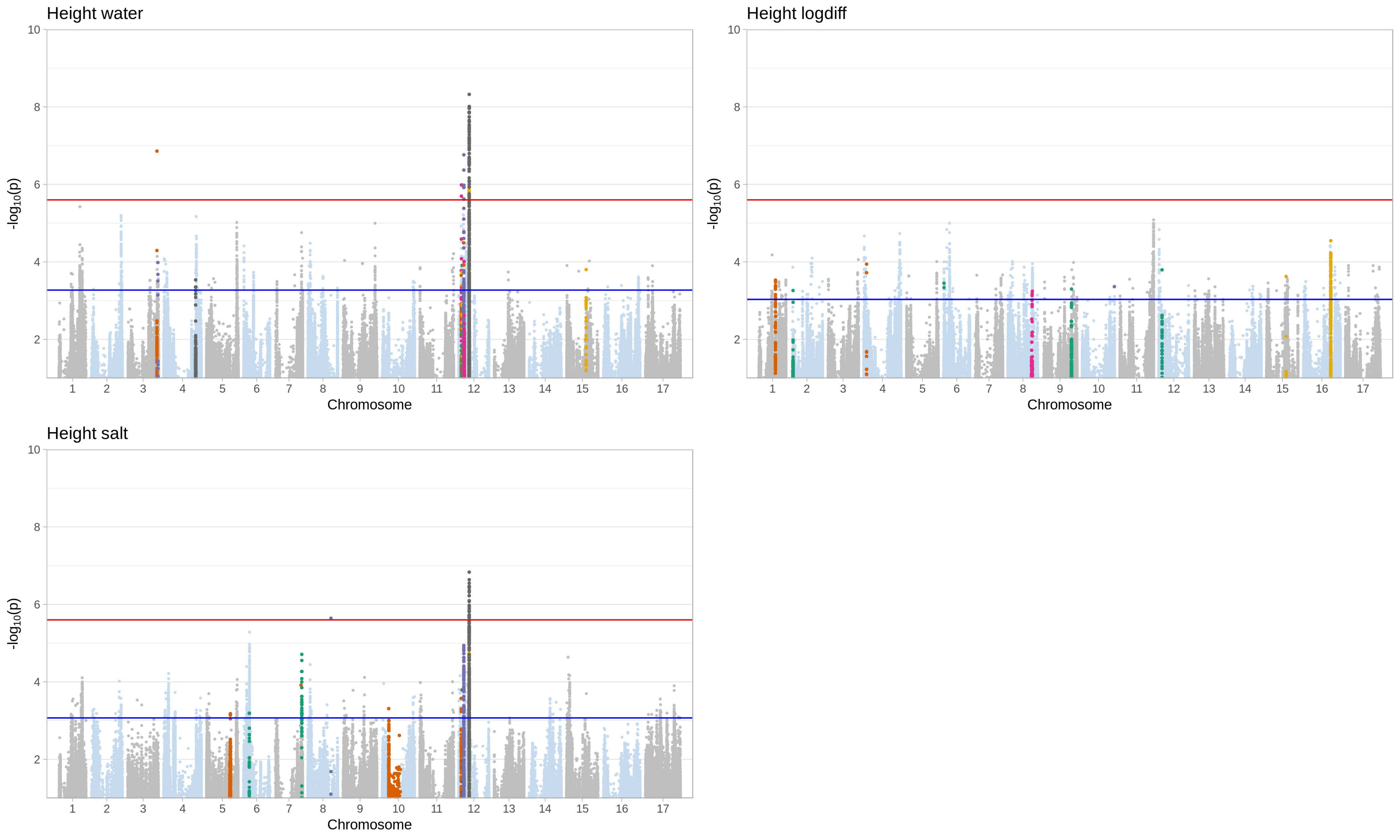

Height water
Height logdiff
Height salt
-log10(p)
Chromosome
10
8
6
4
2
1 2 3 4 5 6 7 8 9 10 11 12 13 14 15 16 17

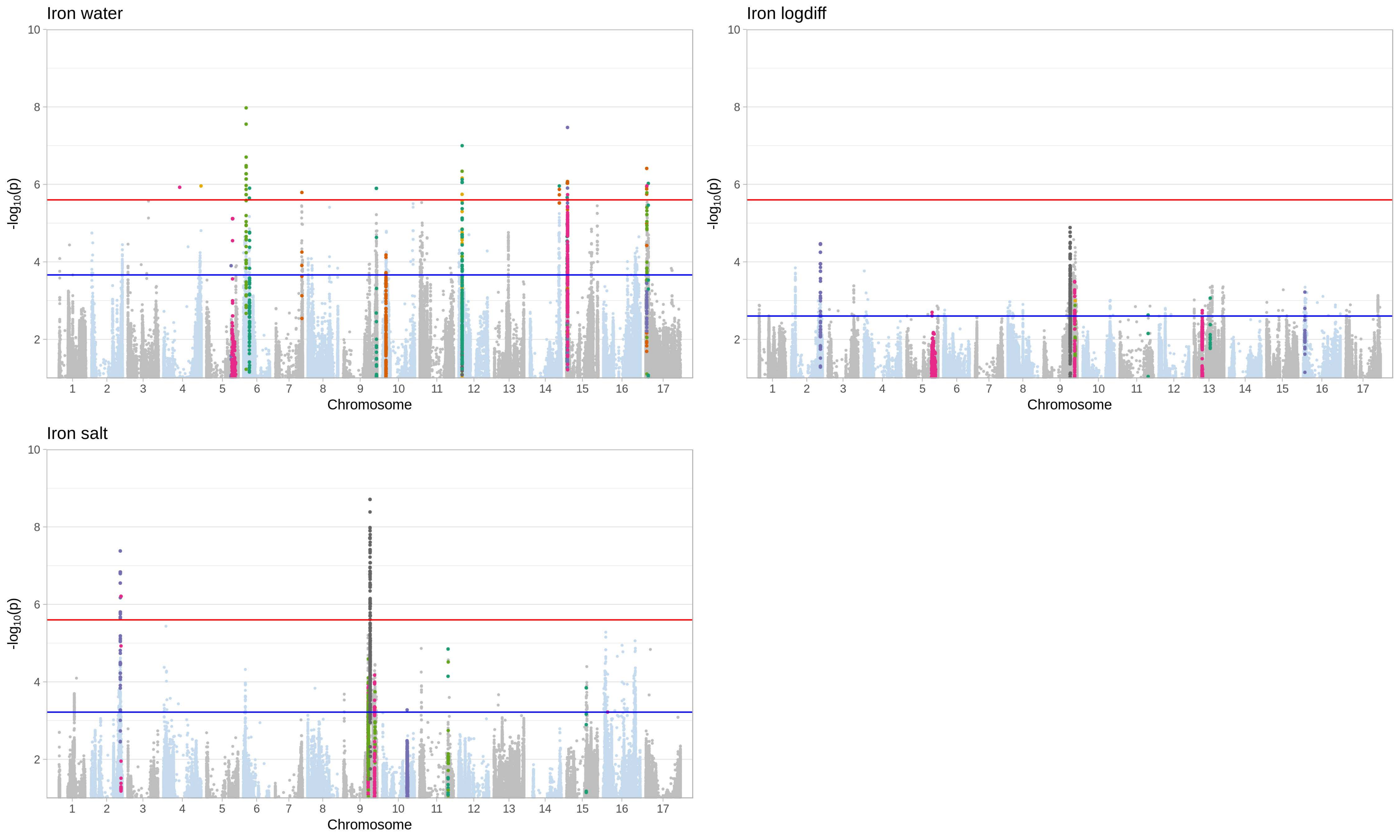

Iron water
Iron logdiff
Iron salt
-log10(p)
Chromosome
10
8
6
4
2

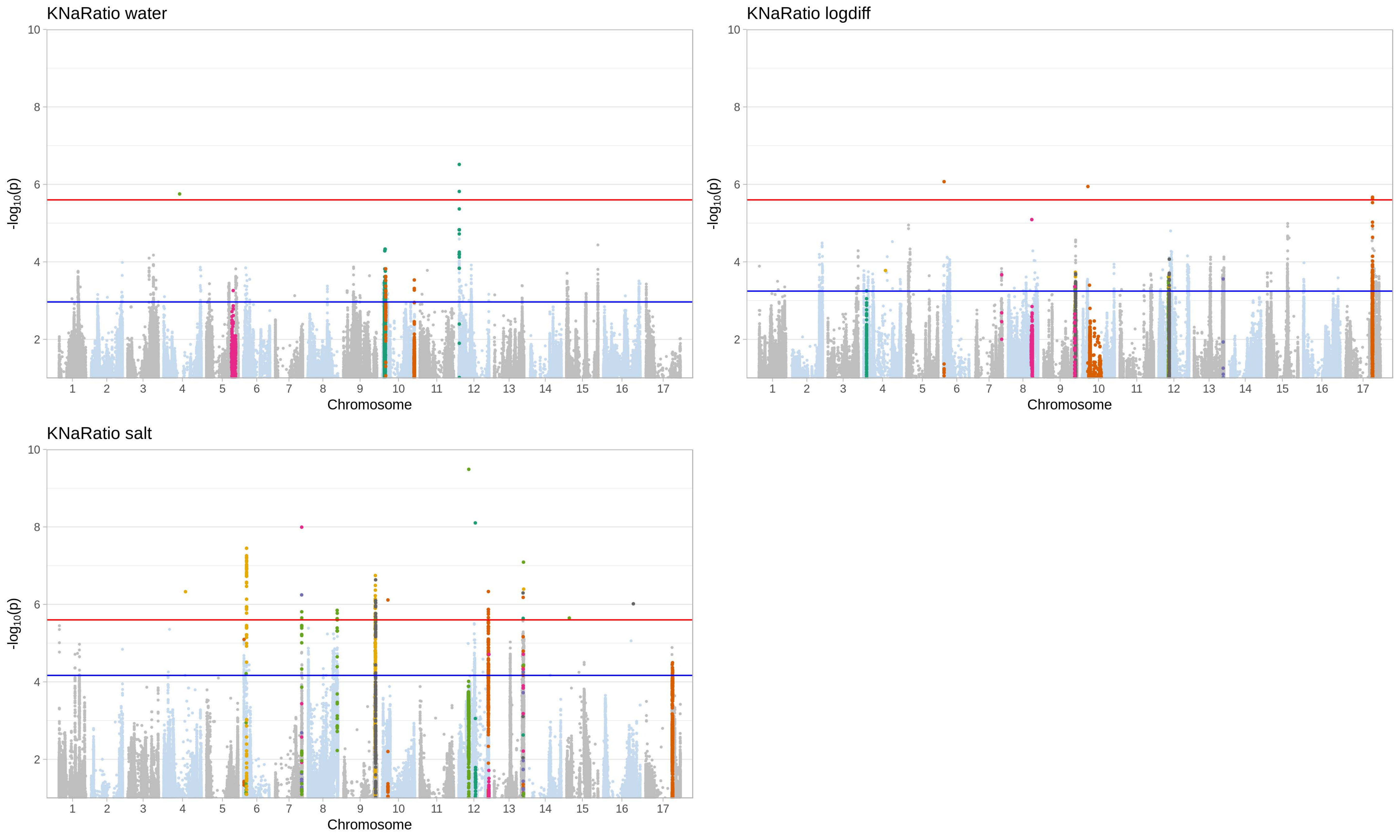
KNaRatio water
KNaRatio logdiff
KNaRatio salt
-log10(p)
Chromosome
1 2 3 4 5 6 7 8 9 10 11 12 13 14 15 16 17
2 4 6 8 10

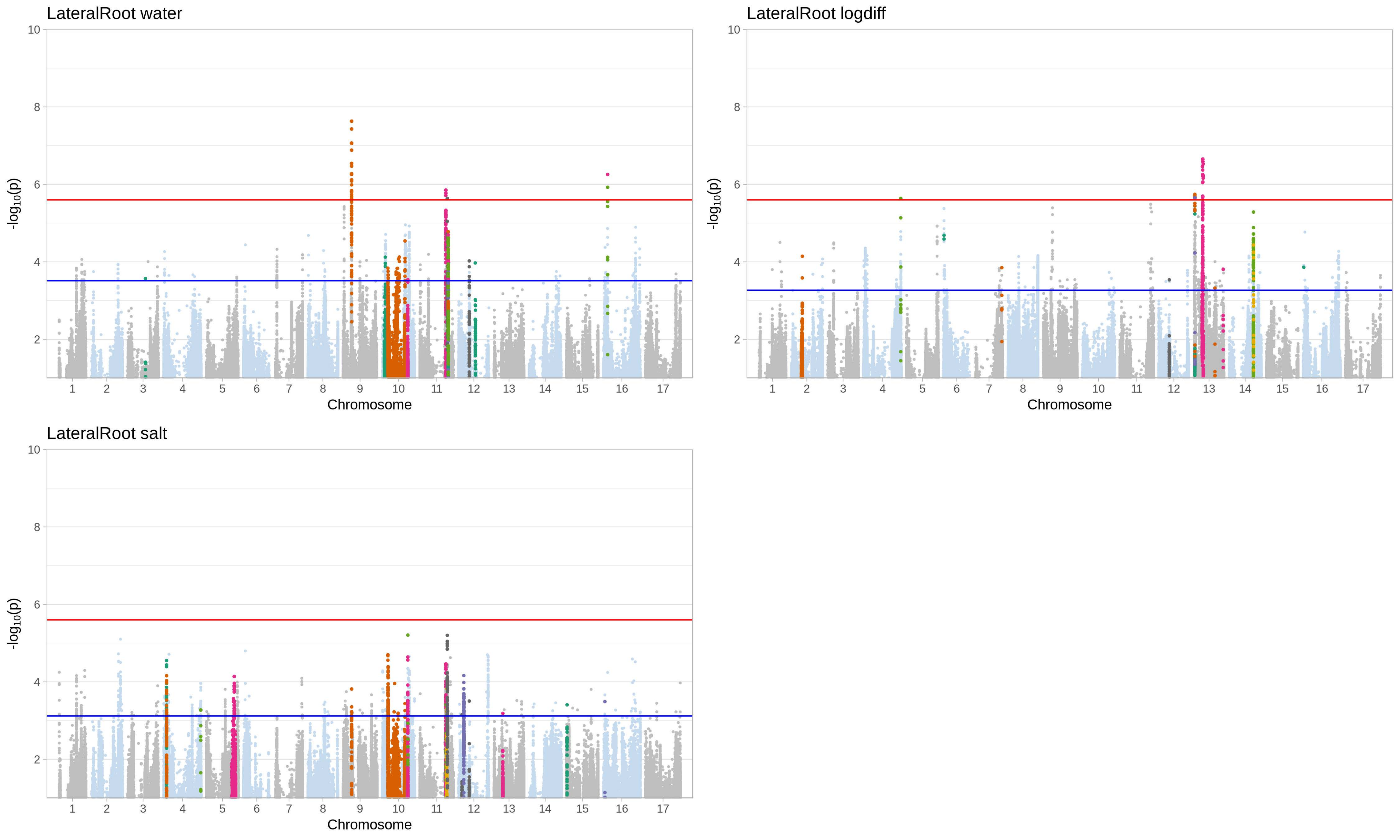

LateralRoot water
LateralRoot logdiff
LateralRoot salt
-log10(p)
Chromosome
1 2 3 4 5 6 7 8 9 10 11 12 13 14 15 16 17
2 4 6 8 10

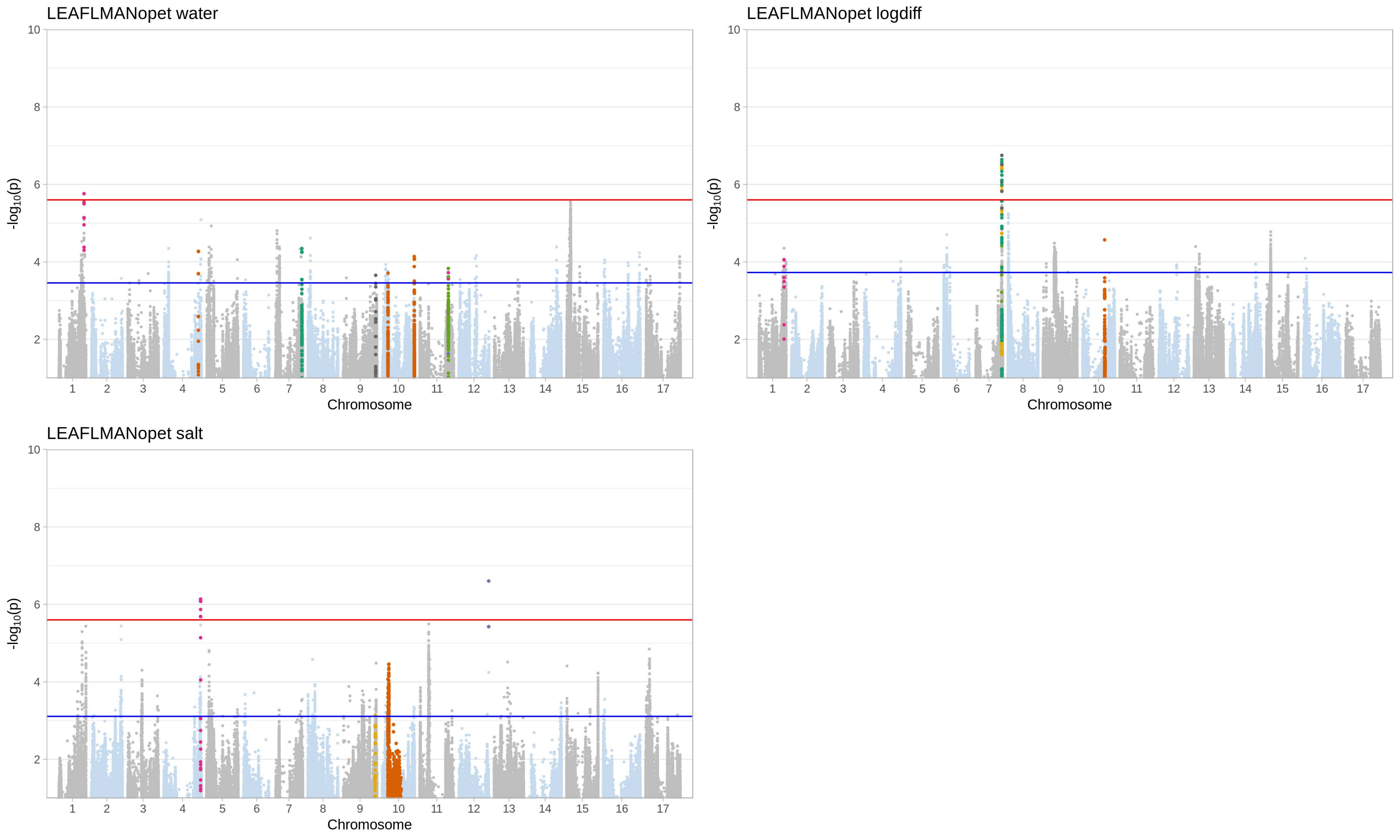
LEAFLMANopet water
LEAFLMANopet logdiff
LEAFLMANopet salt
-log10(p)
Chromosome
1 2 3 4 5 6 7 8 9 10 11 12 13 14 15 16 17
2 4 6 8 10

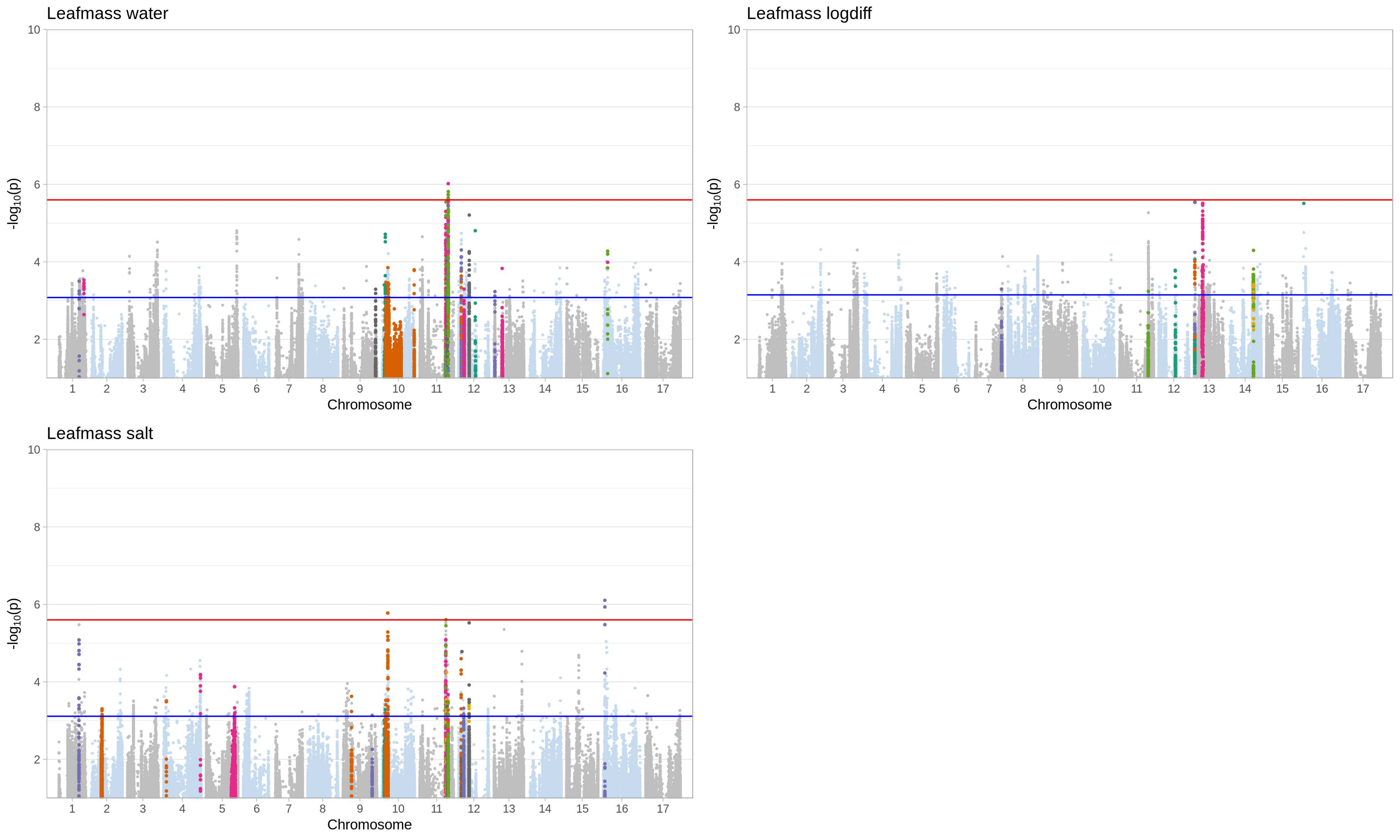

Leafmass water
Leafmass logdiff
Leafmass salt
-log10(p)
Chromosome
10
8
6
4
2

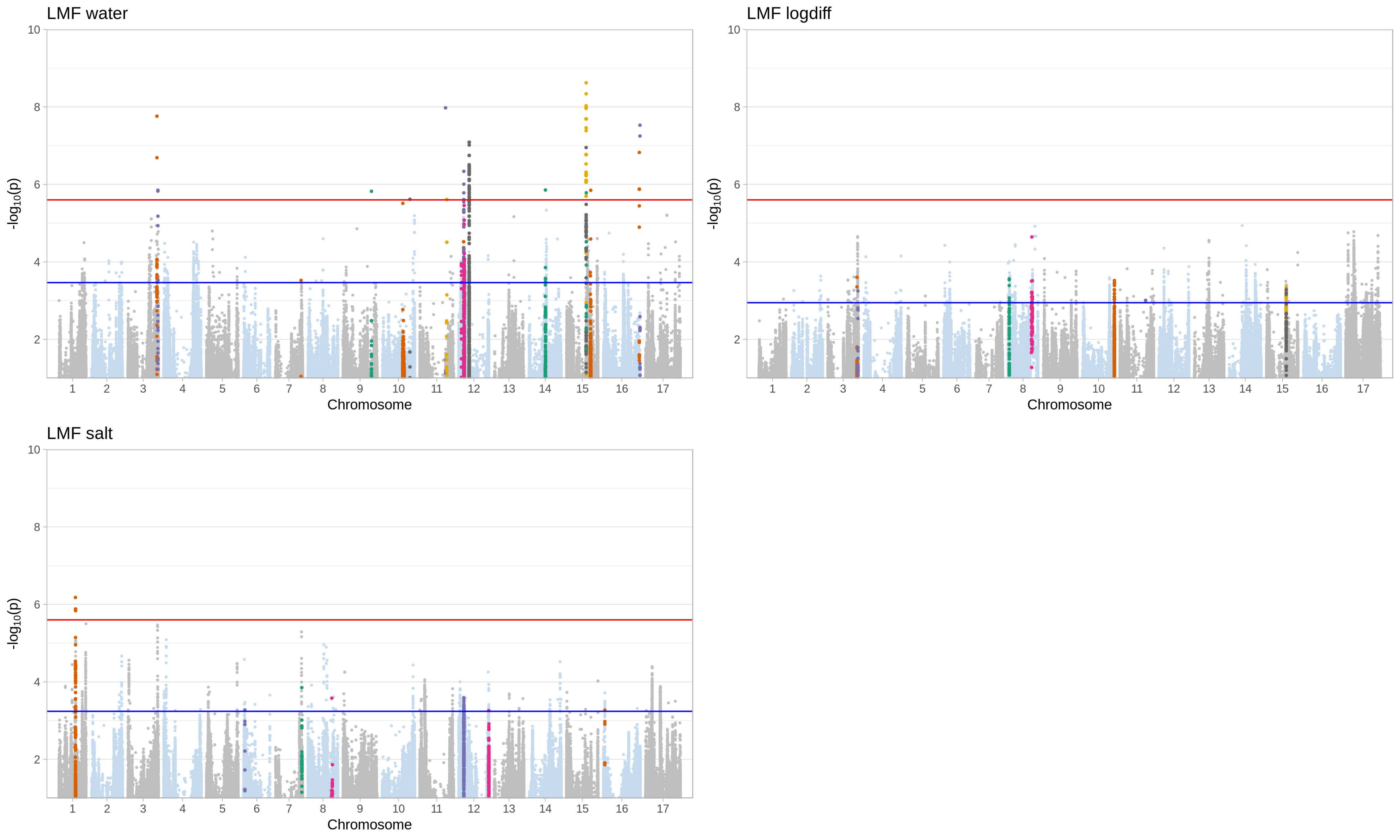

LMF water
LMF logdiff
LMF salt
-log10(p)
Chromosome
1 2 3 4 5 6 7 8 9 10 11 12 13 14 15 16 17
2 4 6 8 10

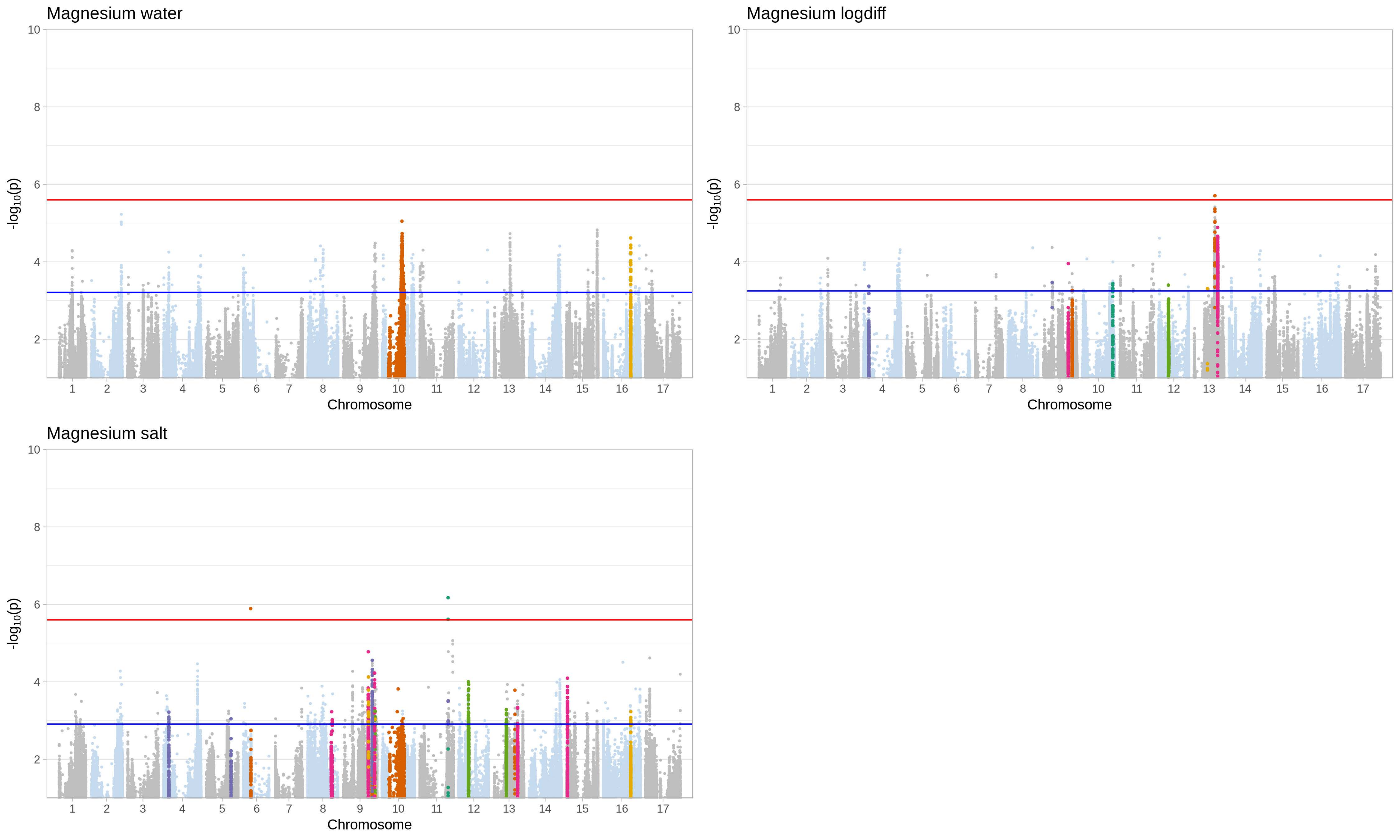
Magnesium water
Magnesium logdiff
Magnesium salt
-log10(p)
Chromosome
1 2 3 4 5 6 7 8 9 10 11 12 13 14 15 16 17
0 2 4 6 8 10

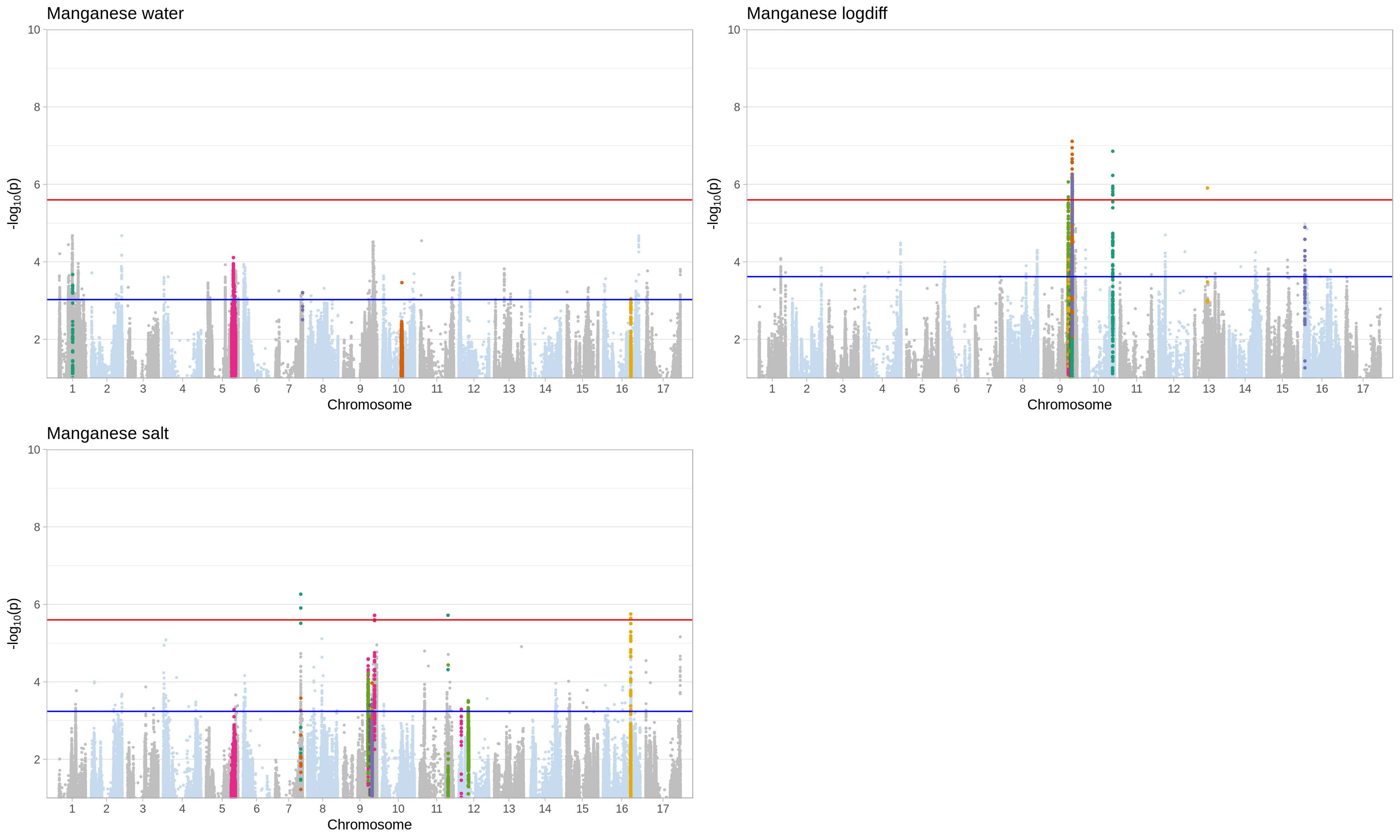

Manganese water
Manganese logdiff
Manganese salt
-log10(p)
Chromosome
10
8
6
4
2

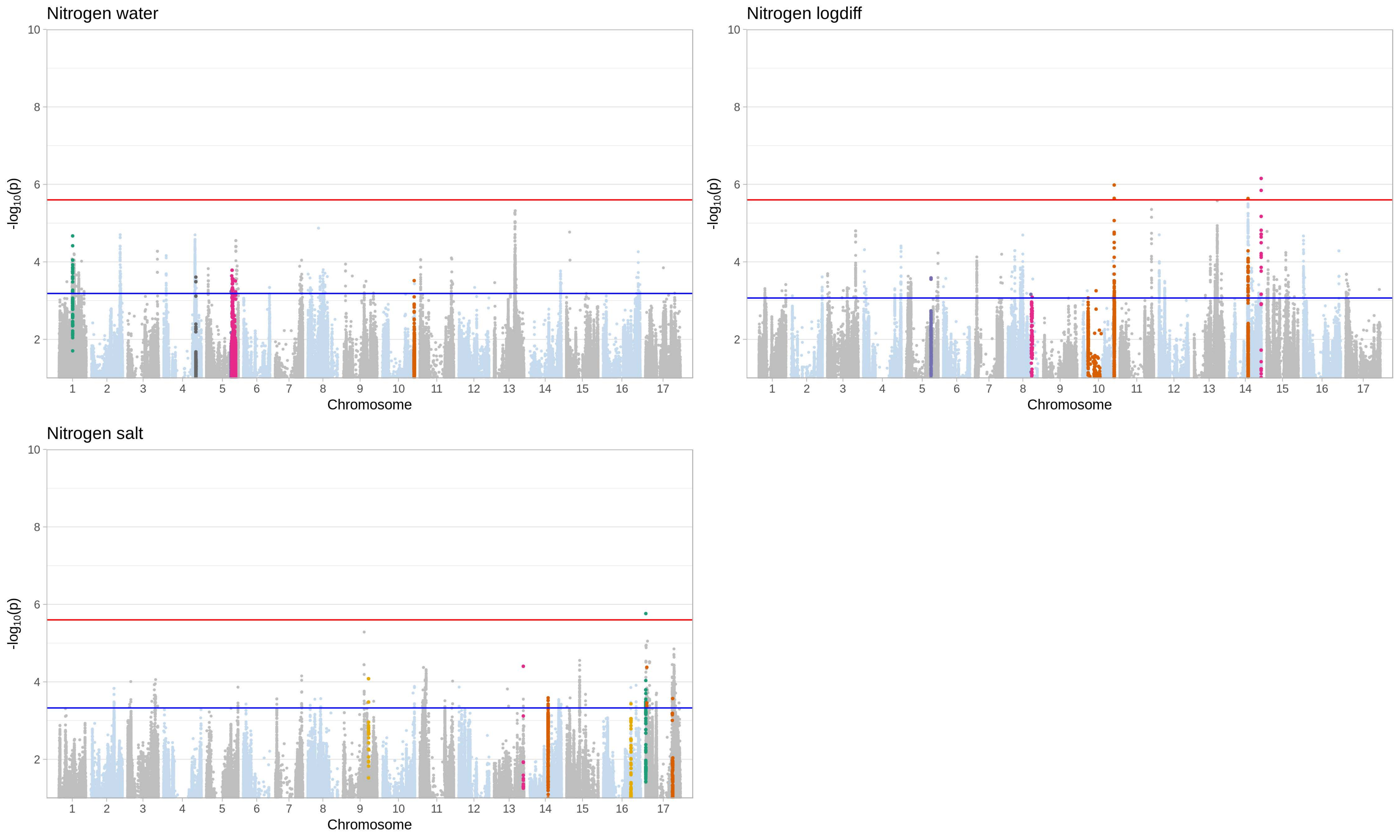
Nitrogen water
Nitrogen logdiff
Nitrogen salt
-log10(p)
Chromosome
1 2 3 4 5 6 7 8 9 10 11 12 13 14 15 16 17
2 4 6 8 10

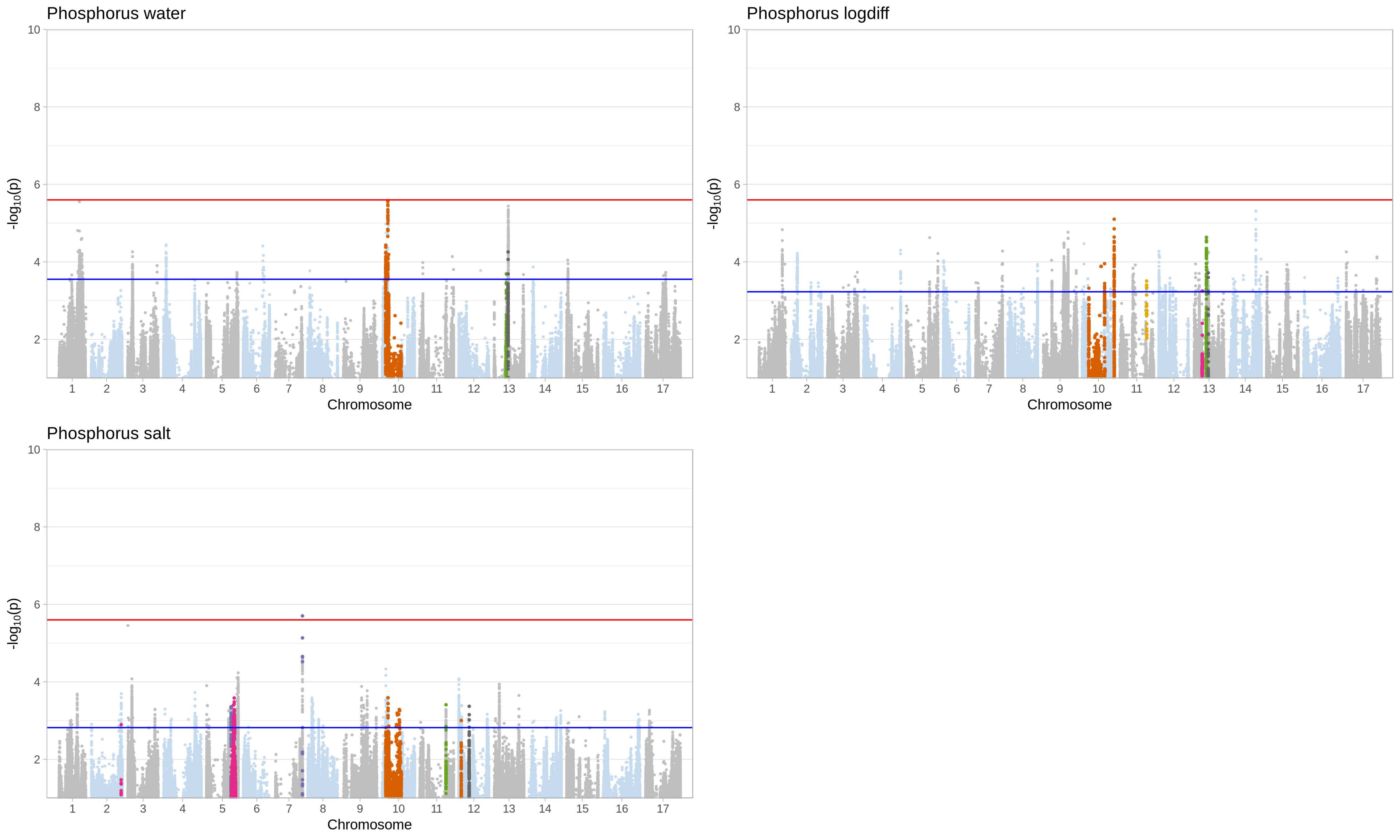
Phosphorus water
Phosphorus logdiff
Phosphorus salt
-log10(p)
Chromosome
0
2
4
6
8
10

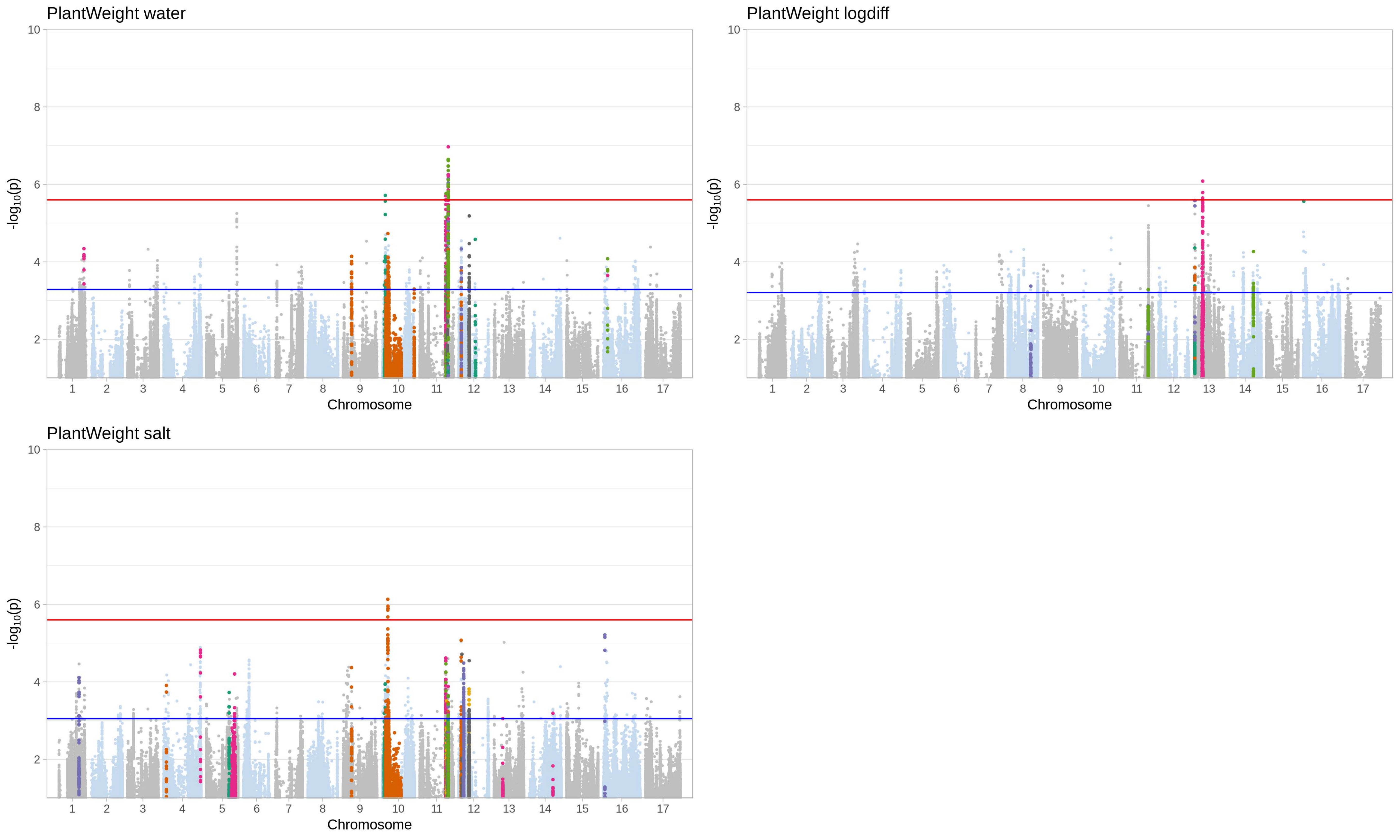
PlantWeight water
PlantWeight logdiff
PlantWeight salt
-log10(p)
Chromosome
10
8
6
4
2
1 2 3 4 5 6 7 8 9 10 11 12 13 14 15 16 17

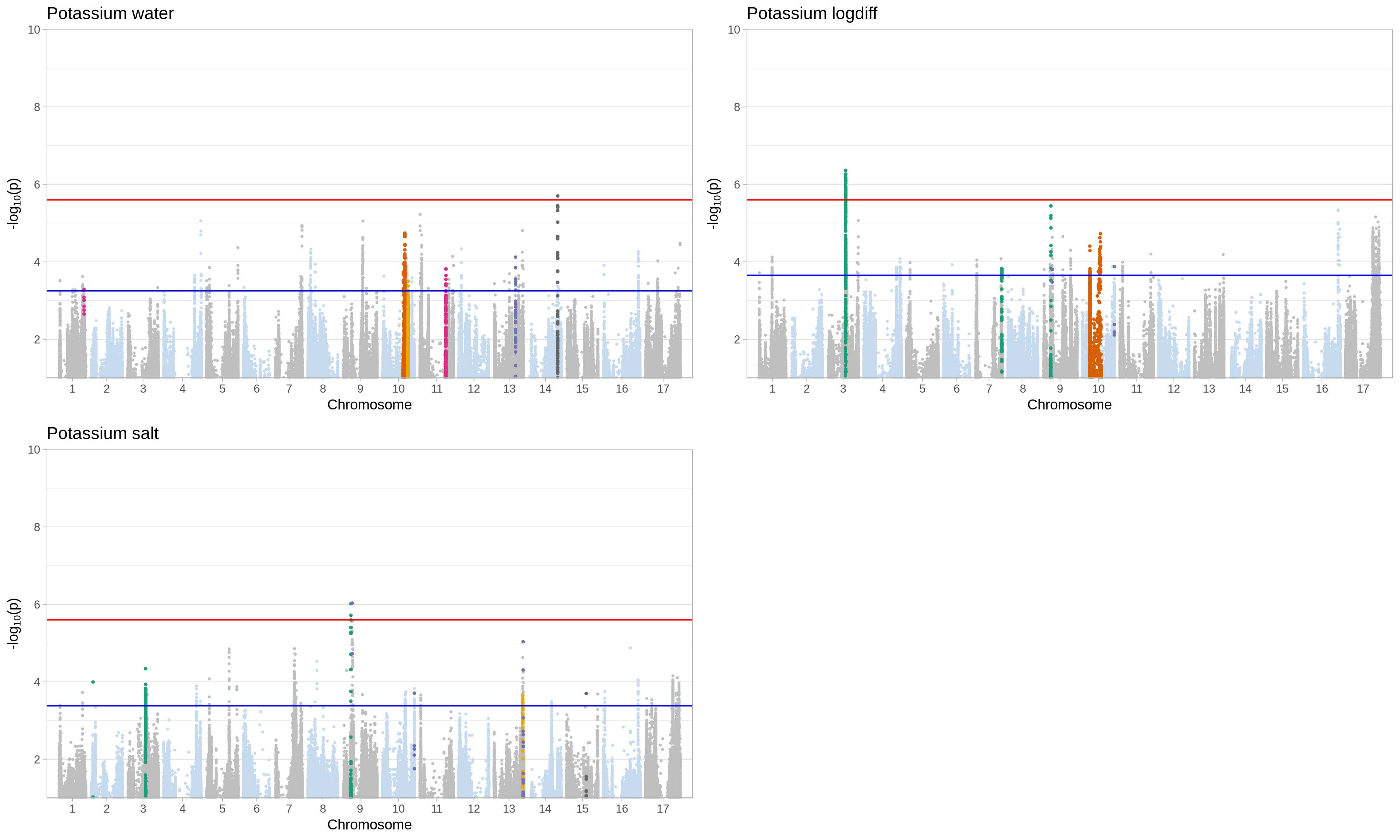
Potassium water
Potassium logdiff
Potassium salt
$-\log_{10}(p)$
Chromosome
10
8
6
4
2
1 2 3 4 5 6 7 8 9 10 11 12 13 14 15 16 17

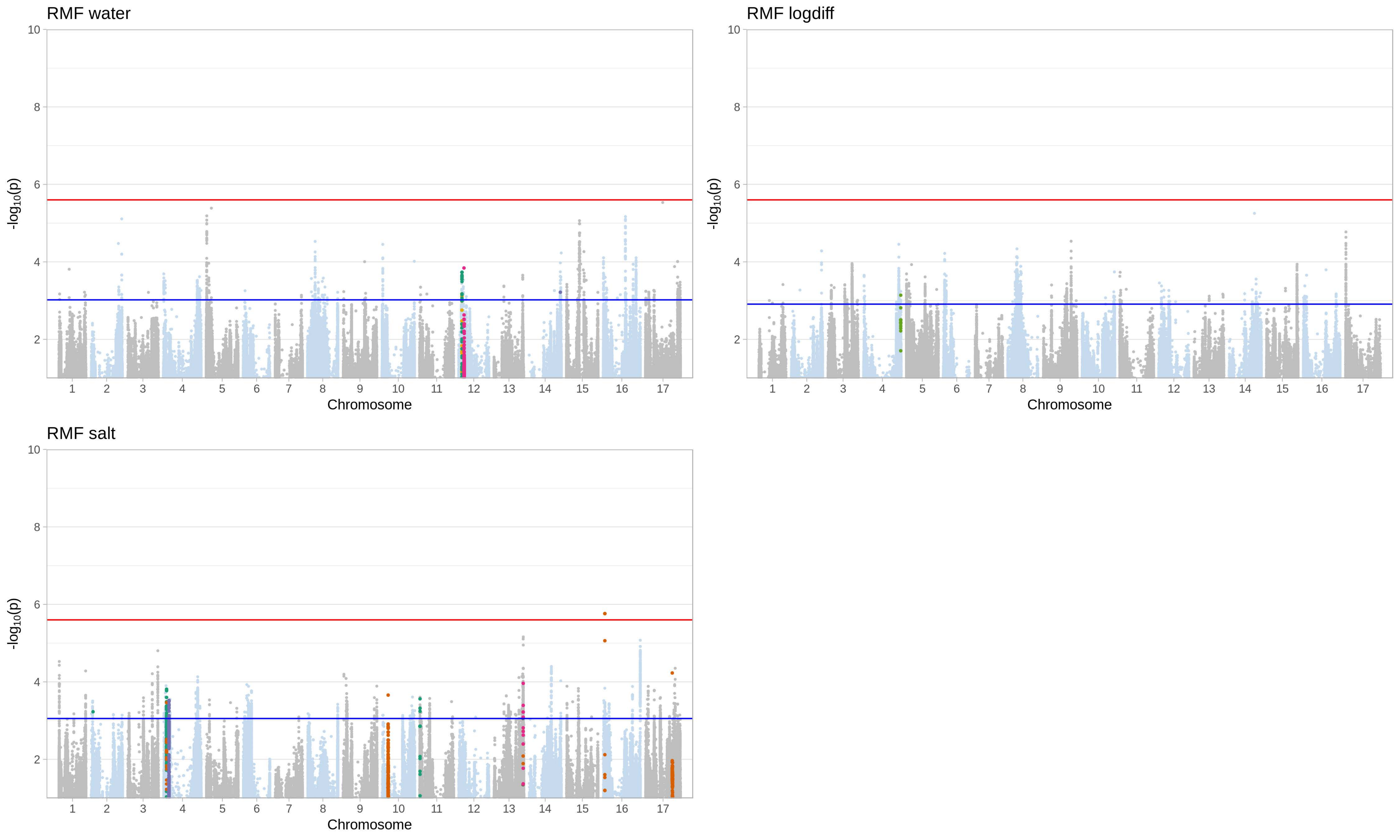

RMF water
RMF logdiff
RMF salt
-log10(p)
Chromosome
10
8
6
4
2
1 2 3 4 5 6 7 8 9 10 11 12 13 14 15 16 17

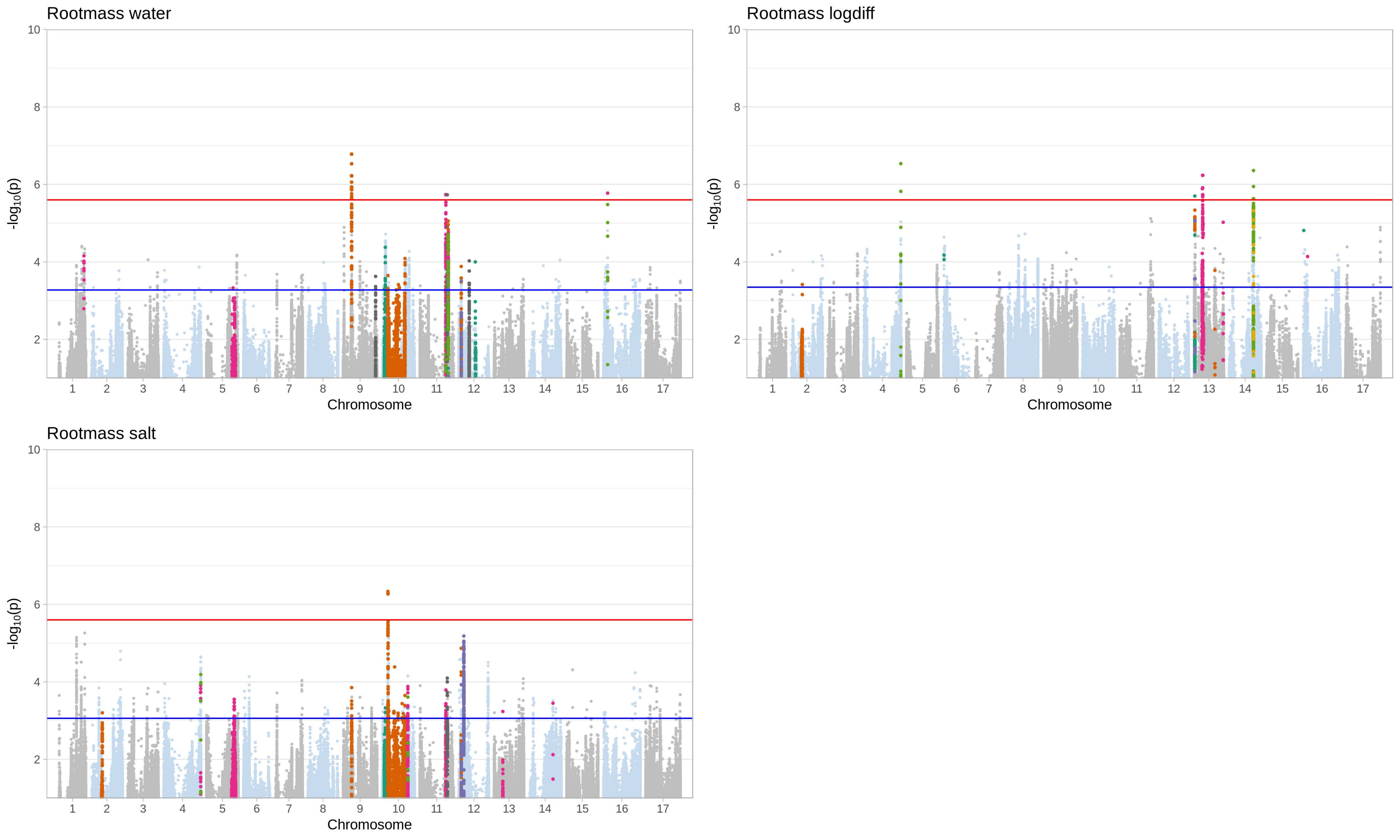
Rootmass water
Rootmass logdiff
Rootmass salt
-log10(p)
Chromosome
0
2
4
6
8
10

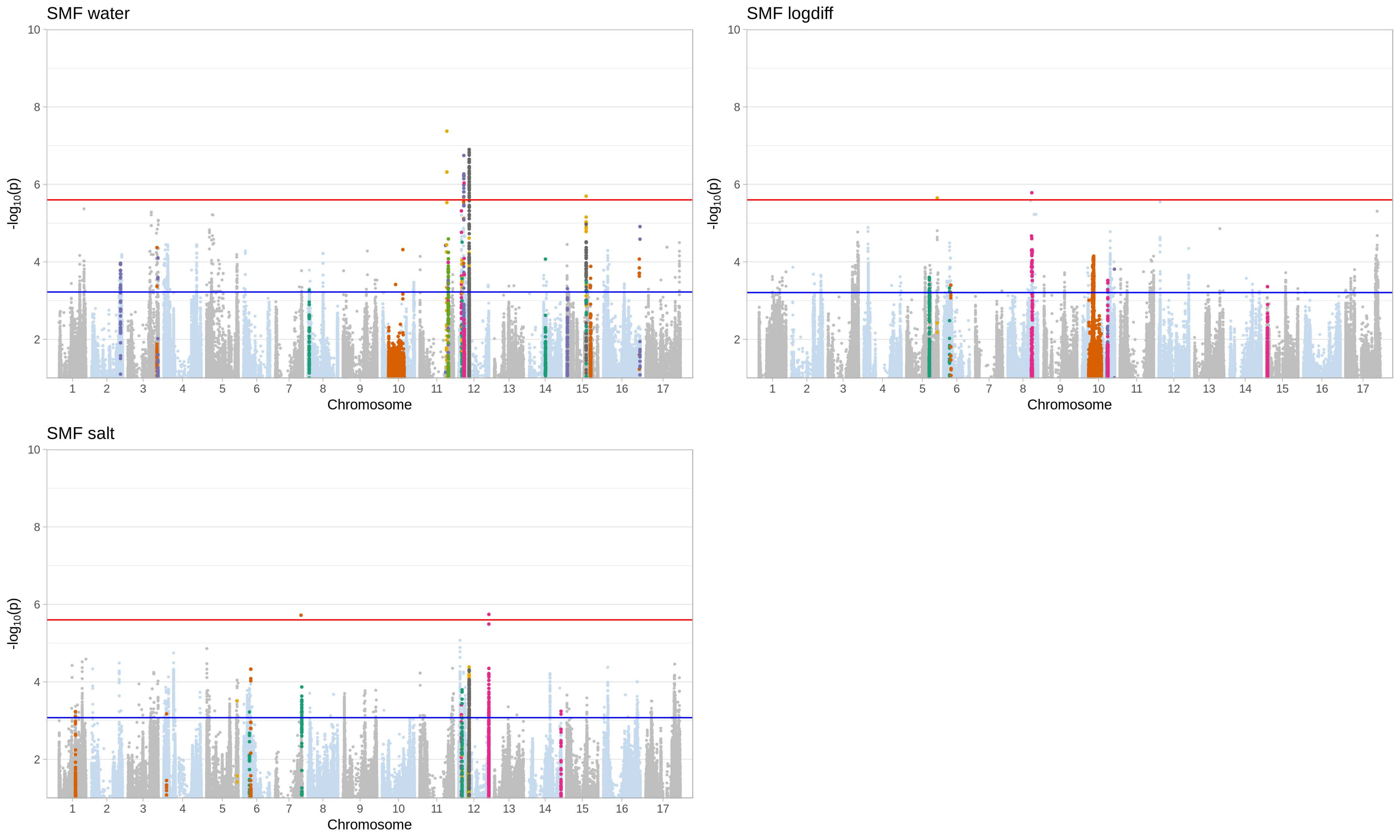
SMF water
SMF logdiff
SMF salt
-log10(p)
Chromosome
1 2 3 4 5 6 7 8 9 10 11 12 13 14 15 16 17
10
8
6
4
2

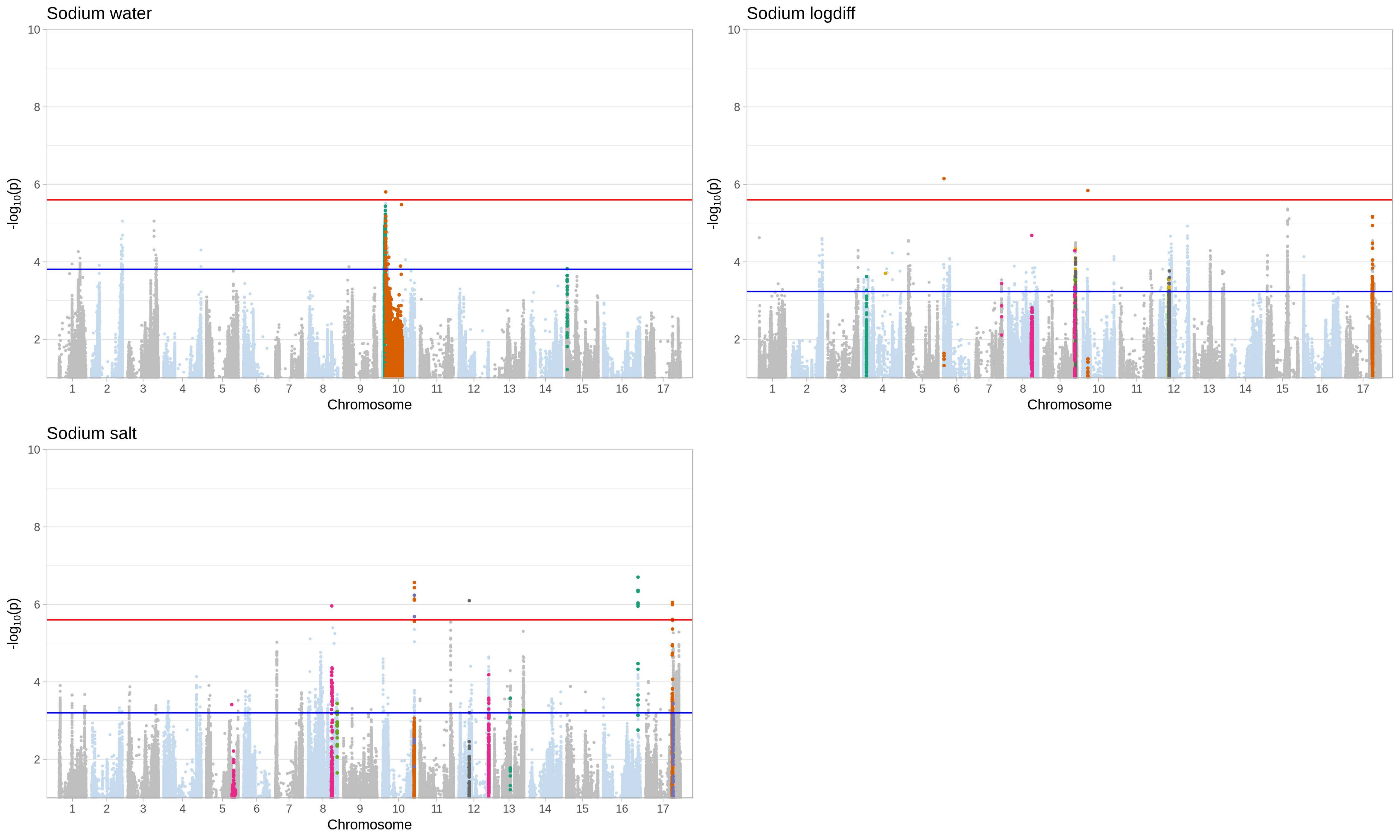

Sodium water
Sodium logdiff
Sodium salt
-log10(p)
Chromosome
10
8
6
4
2
1 2 3 4 5 6 7 8 9 10 11 12 13 14 15 16 17

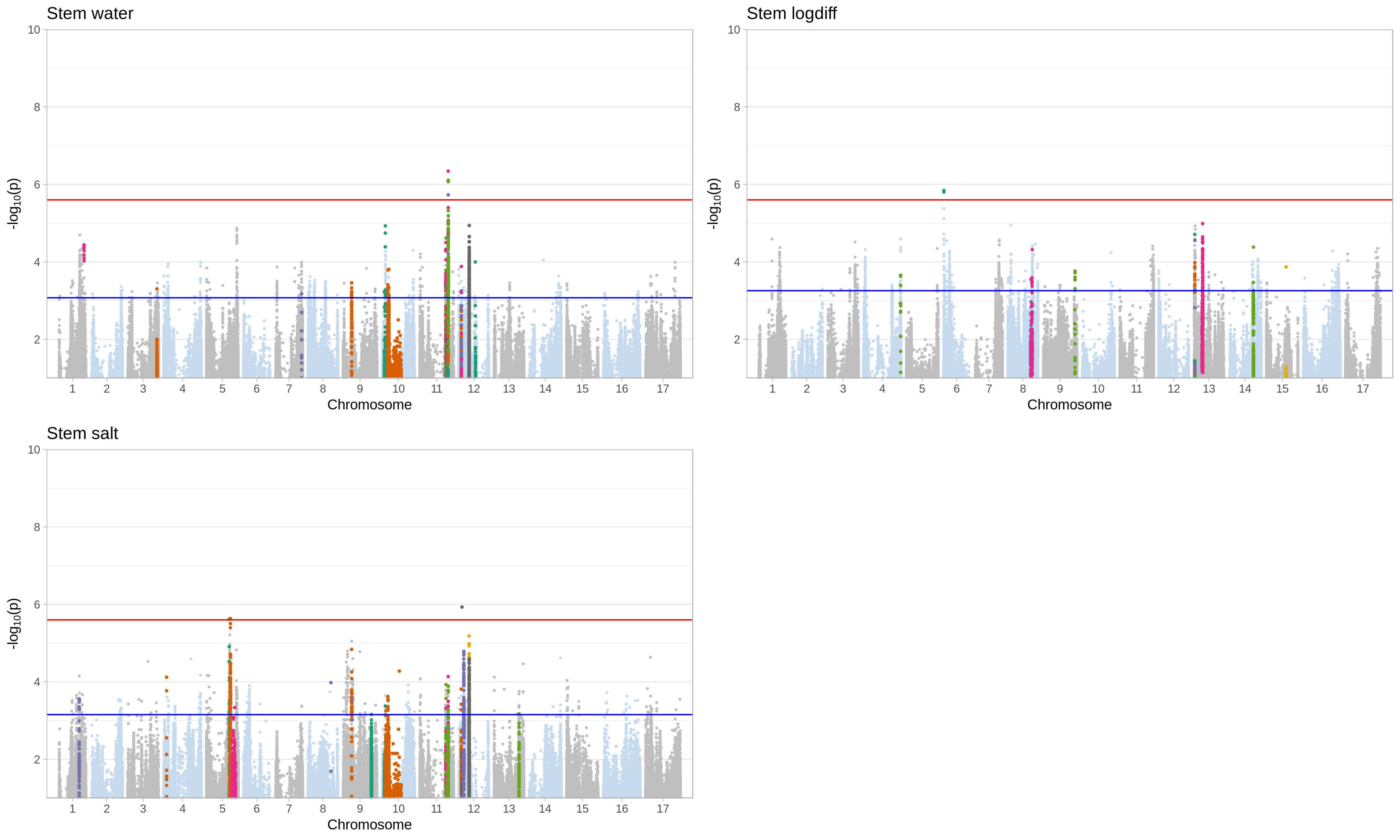

Stem water
Stem logdiff
Stem salt
-log10(p)
Chromosome
1 2 3 4 5 6 7 8 9 10 11 12 13 14 15 16 17
2 4 6 8 10

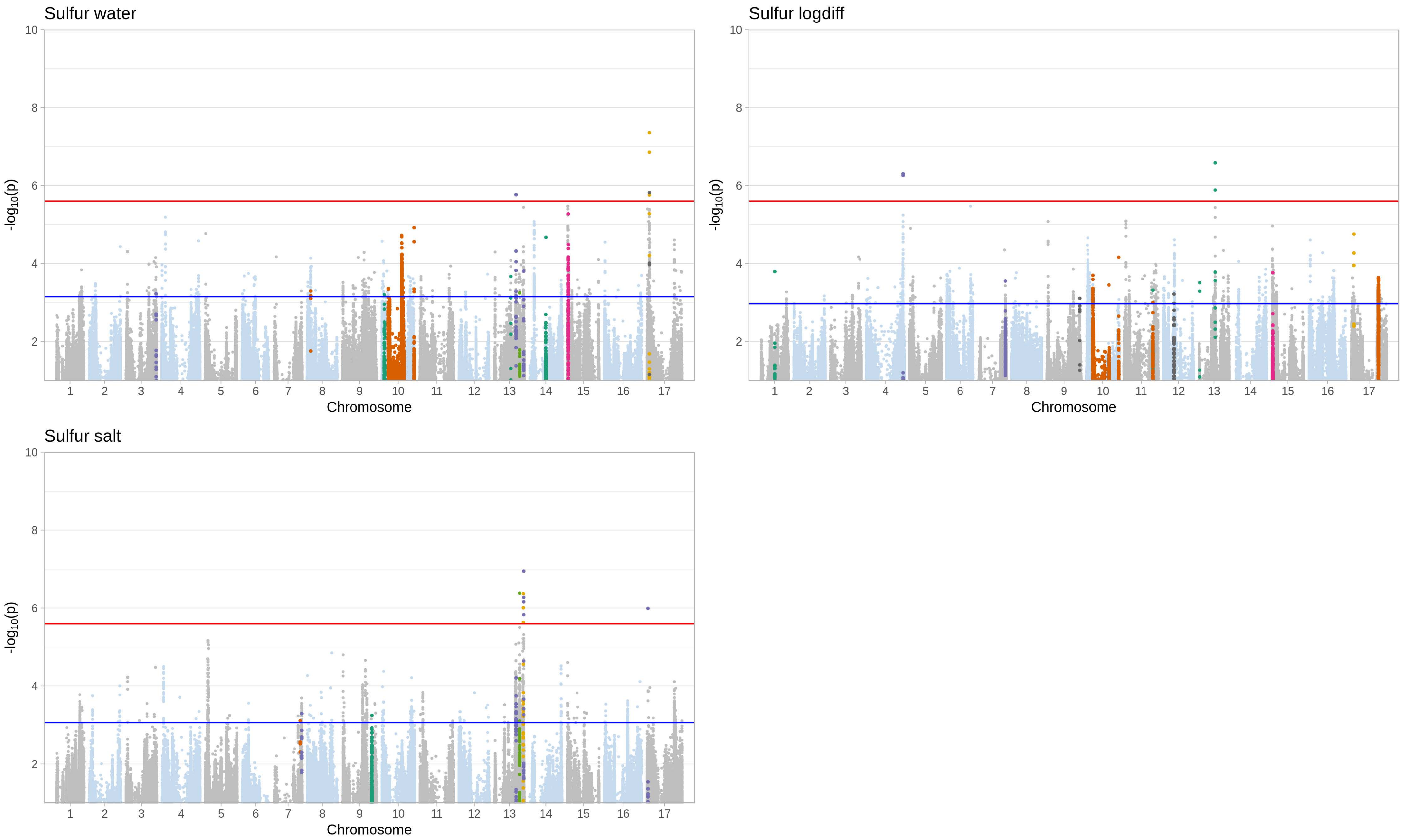

Sulfur water
Sulfur logdiff
Sulfur salt
-log10(p)
Chromosome

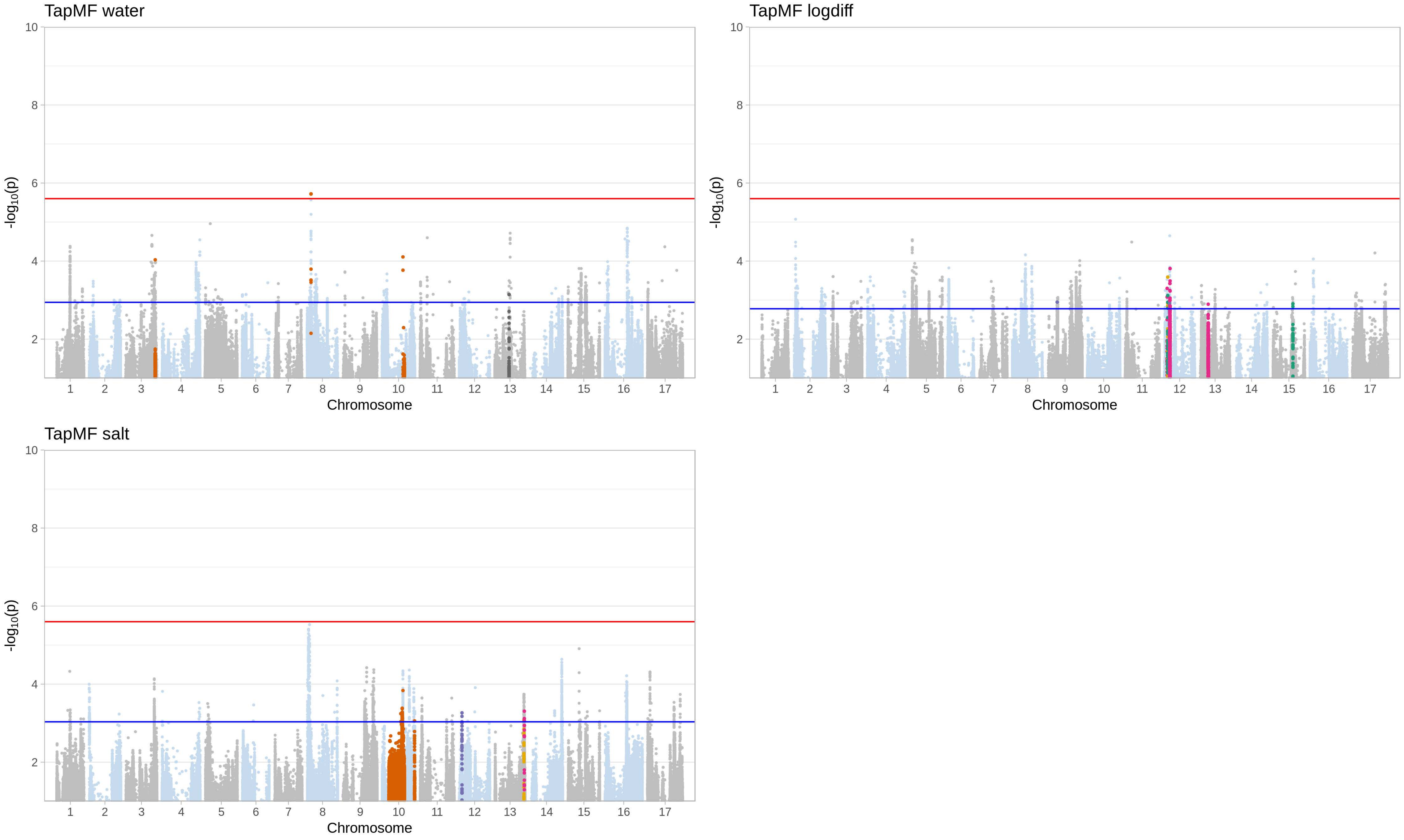

TapMF water
TapMF logdiff
TapMF salt
-log$_{10}$(p)
Chromosome
0
2
4
6
8
10
1 2 3 4 5 6 7 8 9 10 11 12 13 14 15 16 17

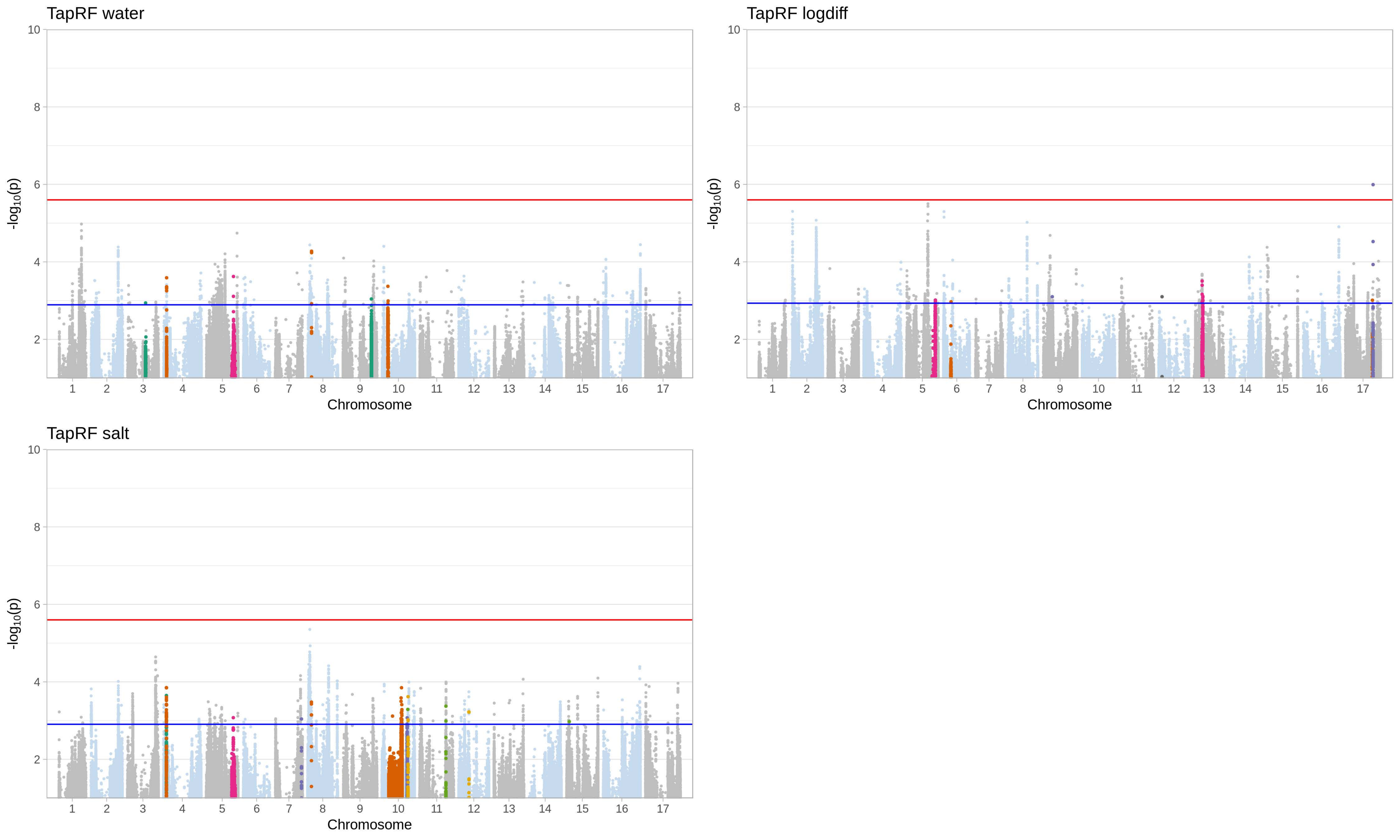
TapRF water
TapRF logdiff
TapRF salt
-log10(p)
Chromosome
1 2 3 4 5 6 7 8 9 10 11 12 13 14 15 16 17
2 4 6 8 10

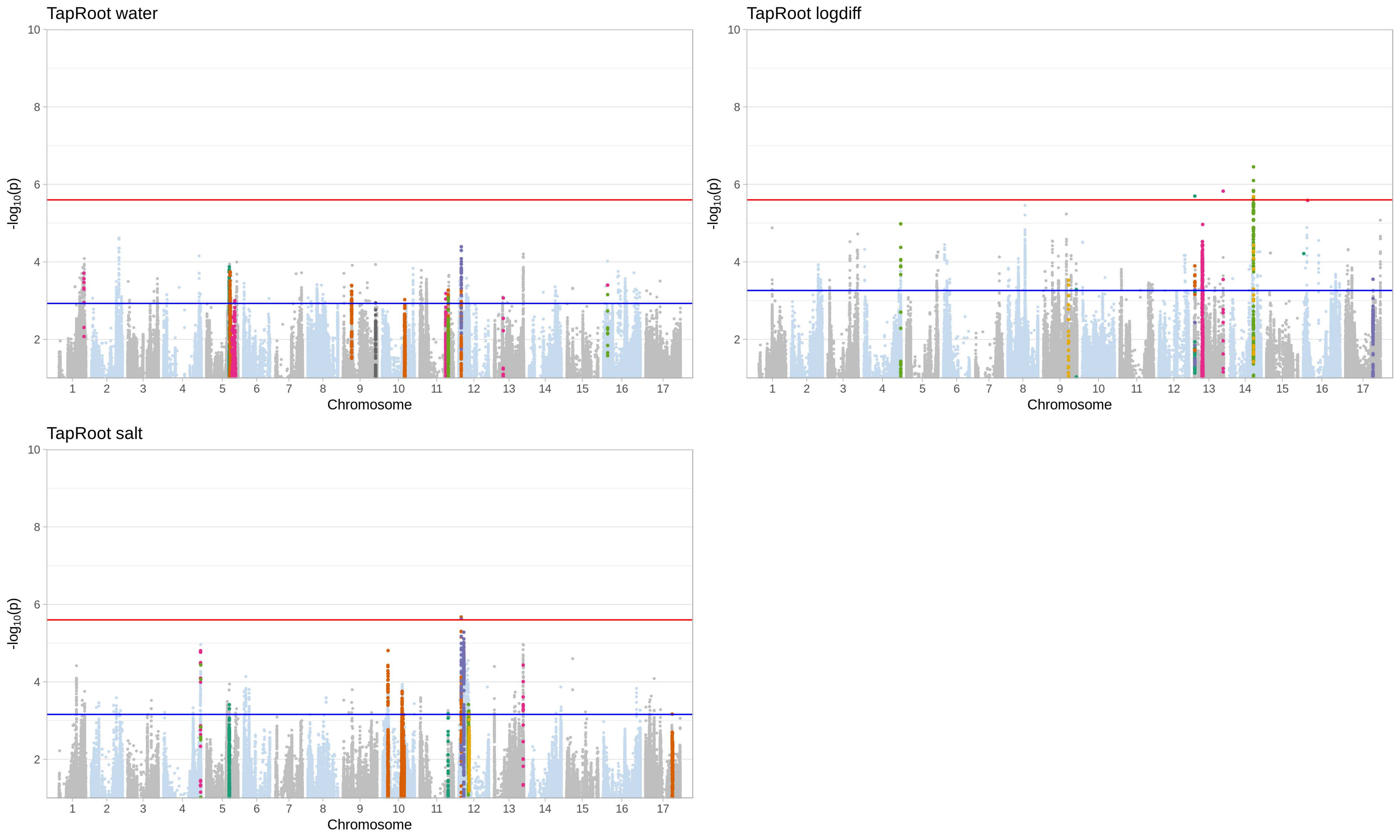

TapRoot water
TapRoot logdiff
TapRoot salt
-log10(p)
Chromosome
10
8
6
4
2

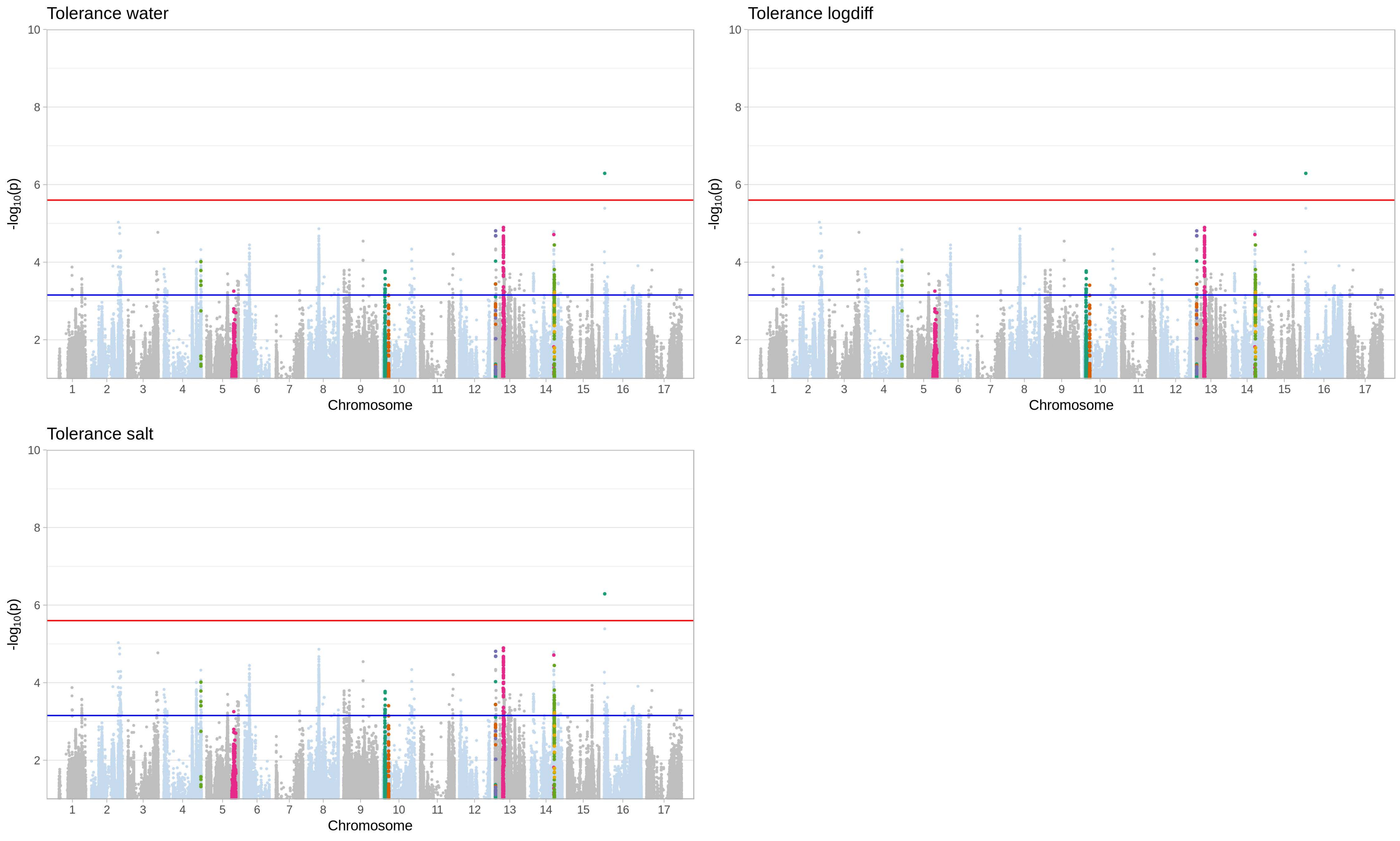

Tolerance water
10
8
6
4
2
-log10(p)
1 2 3 4 5 6 7 8 9 10 11 12 13 14 15 16 17
Chromosome
Tolerance logdiff
10
8
6
4
2
-log10(p)
1 2 3 4 5 6 7 8 9 10 11 12 13 14 15 16 17
Chromosome
Tolerance salt
10
8
6
4
2
-log10(p)
1 2 3 4 5 6 7 8 9 10 11 12 13 14 15 16 17
Chromosome

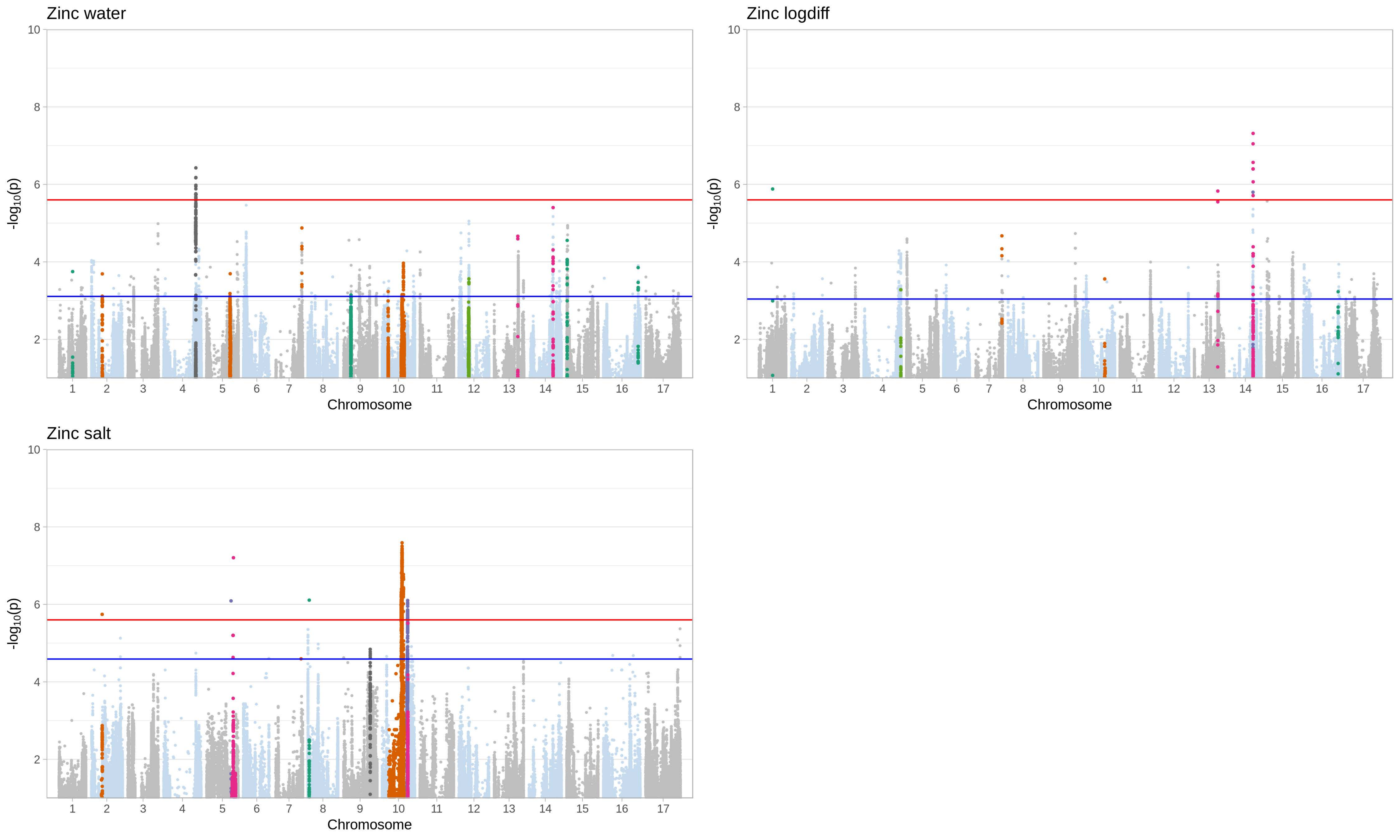
Zinc water
Zinc logdiff
Zinc salt
-log10(p)
Chromosome
10
8
6
4
2

**Figure S6. Trait co-localization under control and salt-stressed conditions as well as their plasticity.** Traits in (a) control, (b) salt stressed, and their (c) plasticity are clustered based on hierarchical clustering of their bivariate correlations. Traits grouped more closely together are more highly correlated. If a linkage block contained a significant SNP for a trait, all other traits were assessed to determine if any had a significant (green) or suggestive (i.e., top 0.1% of all SNPs; grey) in the same block . Plus/minus symbols in the tiles indicate the sign of $\beta$ (effect of minor allele on trait). Linkage blocks are ordered by chromosome with black lines separating them. Sizes of the linkage blocks vary both in physical size as well as gene number.

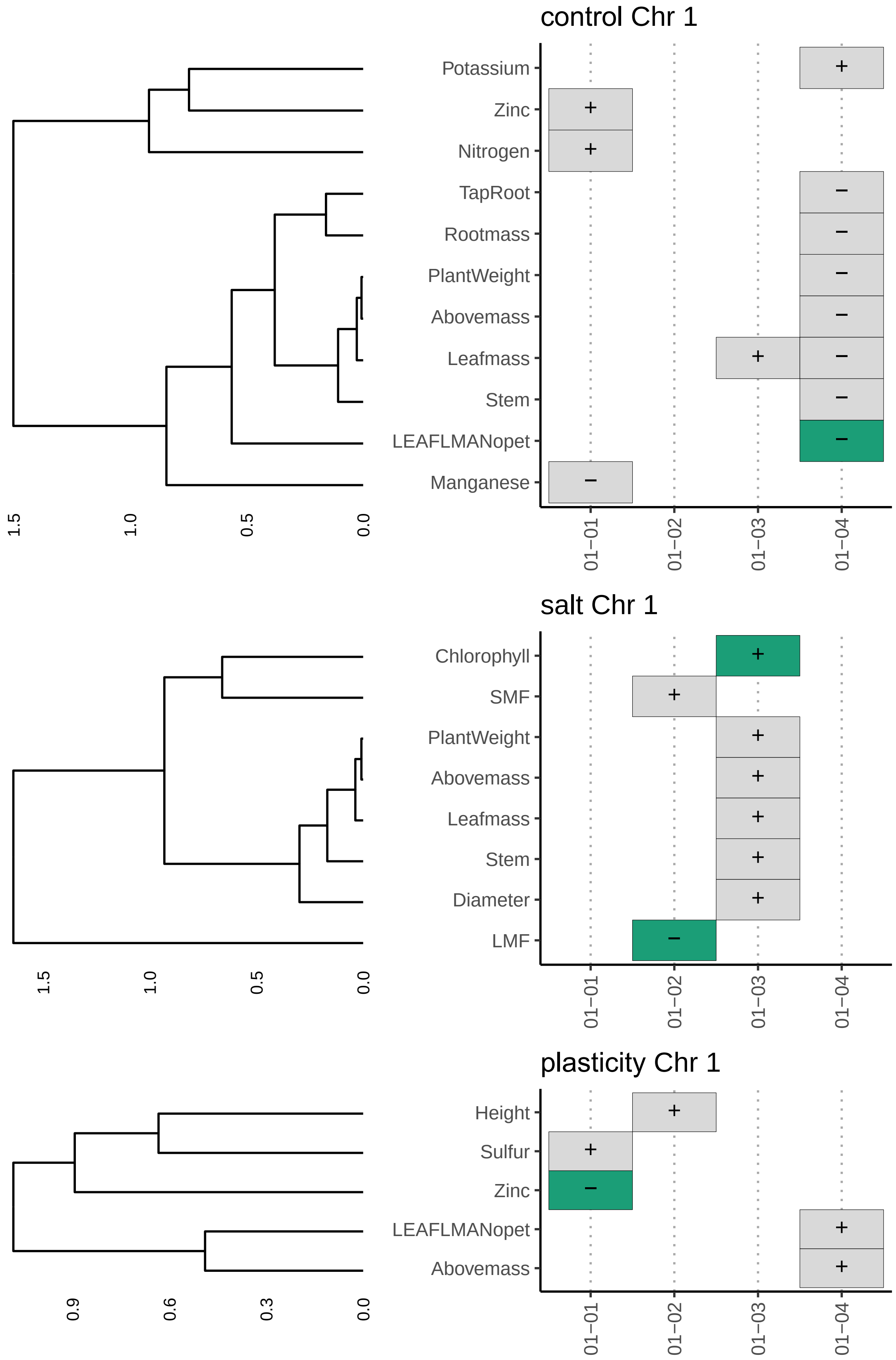

control Chr 1
Potassium
Zinc
Nitrogen
TapRoot
Rootmass
PlantWeight
Abovemass
Leafmass
Stem
LEAFLMANopet
Manganese
1.5
1.0
0.5
0.0
01−01
01−02
01−03
01−04
salt Chr 1
Chlorophyll
SMF
PlantWeight
Abovemass
Leafmass
Stem
Diameter
LMF
1.5
1.0
0.5
0.0
01−01
01−02
01−03
01−04
plasticity Chr 1
Height
Sulfur
Zinc
LEAFLMANopet
Abovemass
0.9
0.6
0.3
0.0
01−01
01−02
01−03
01−04

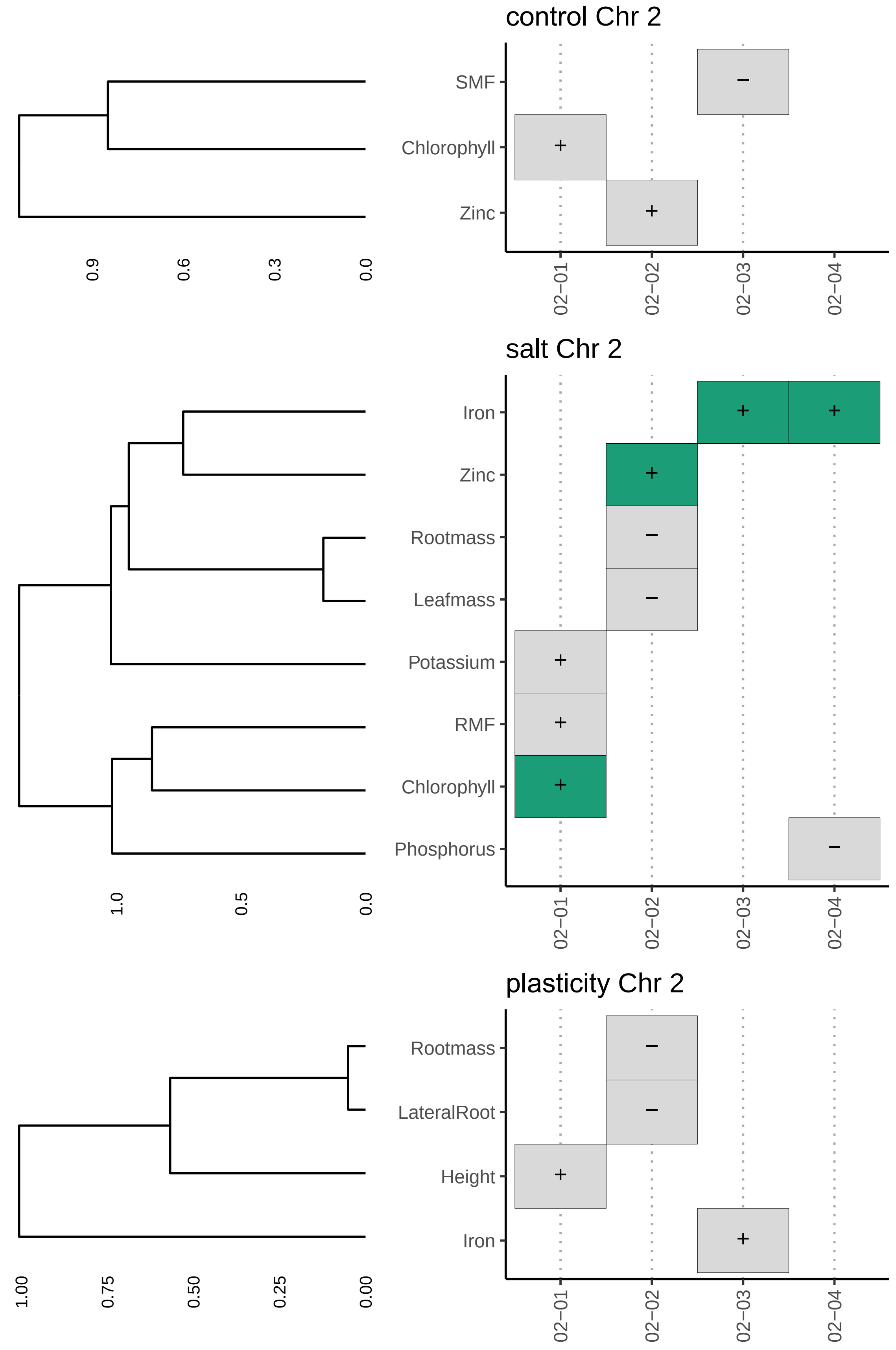

control Chr 2
SMF
Chlorophyll
Zinc
0.9
0.6
0.3
0.0
02−01
02−02
02−03
02−04
salt Chr 2
Iron
Zinc
Rootmass
Leafmass
Potassium
RMF
Chlorophyll
Phosphorus
1.0
0.5
0.0
02−01
02−02
02−03
02−04
plasticity Chr 2
Rootmass
LateralRoot
Height
Iron
1.00
0.75
0.50
0.25
0.00
02−01
02−02
02−03
02−04

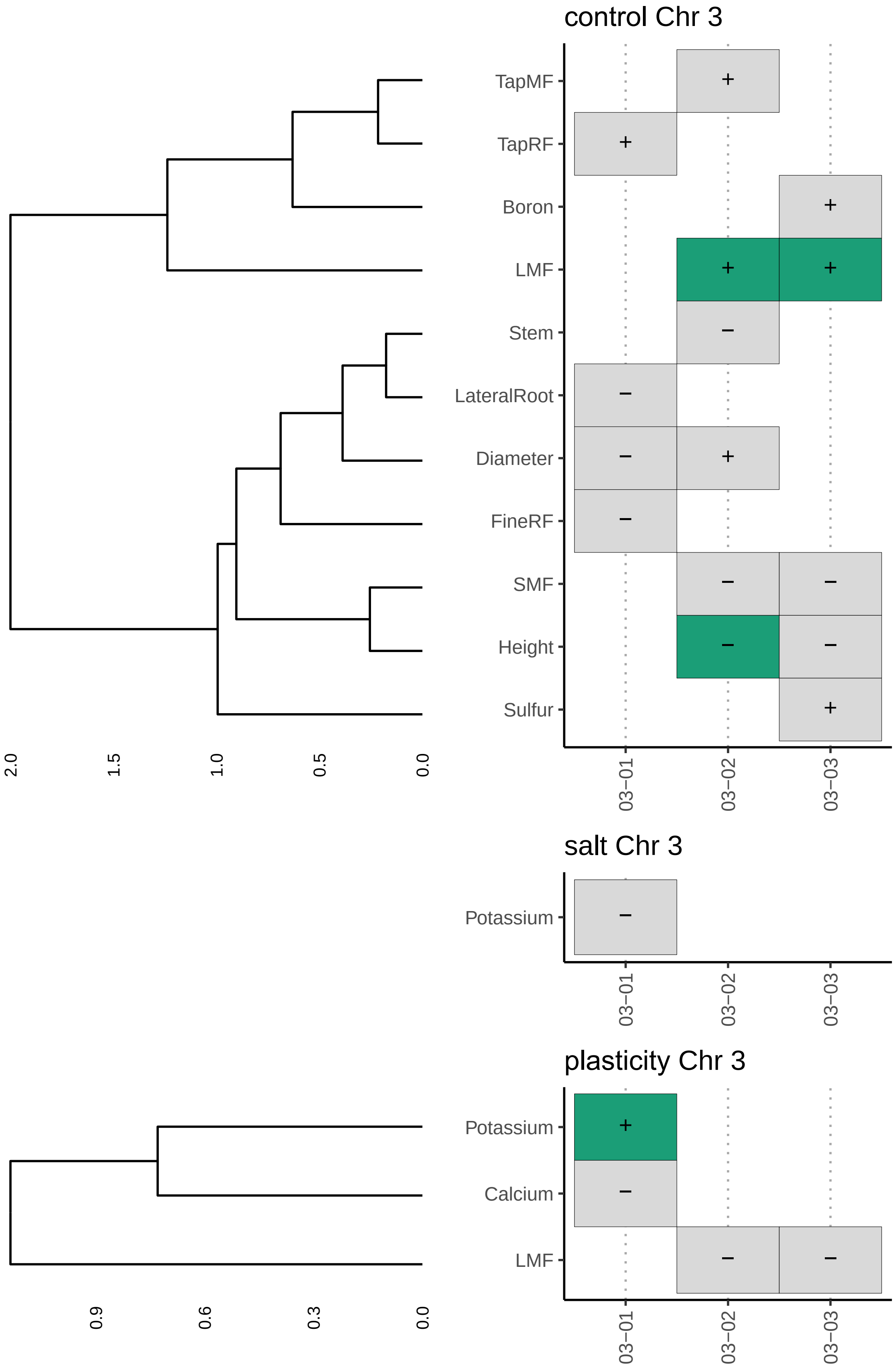
control Chr 3
TapMF
TapRF
Boron
LMF
Stem
LateralRoot
Diameter
FineRF
SMF
Height
Sulfur
2.0
1.5
1.0
0.5
0.0
03−01
03−02
03−03
salt Chr 3
Potassium
plasticity Chr 3
Potassium
Calcium
LMF
0.9
0.6
0.3
0.0

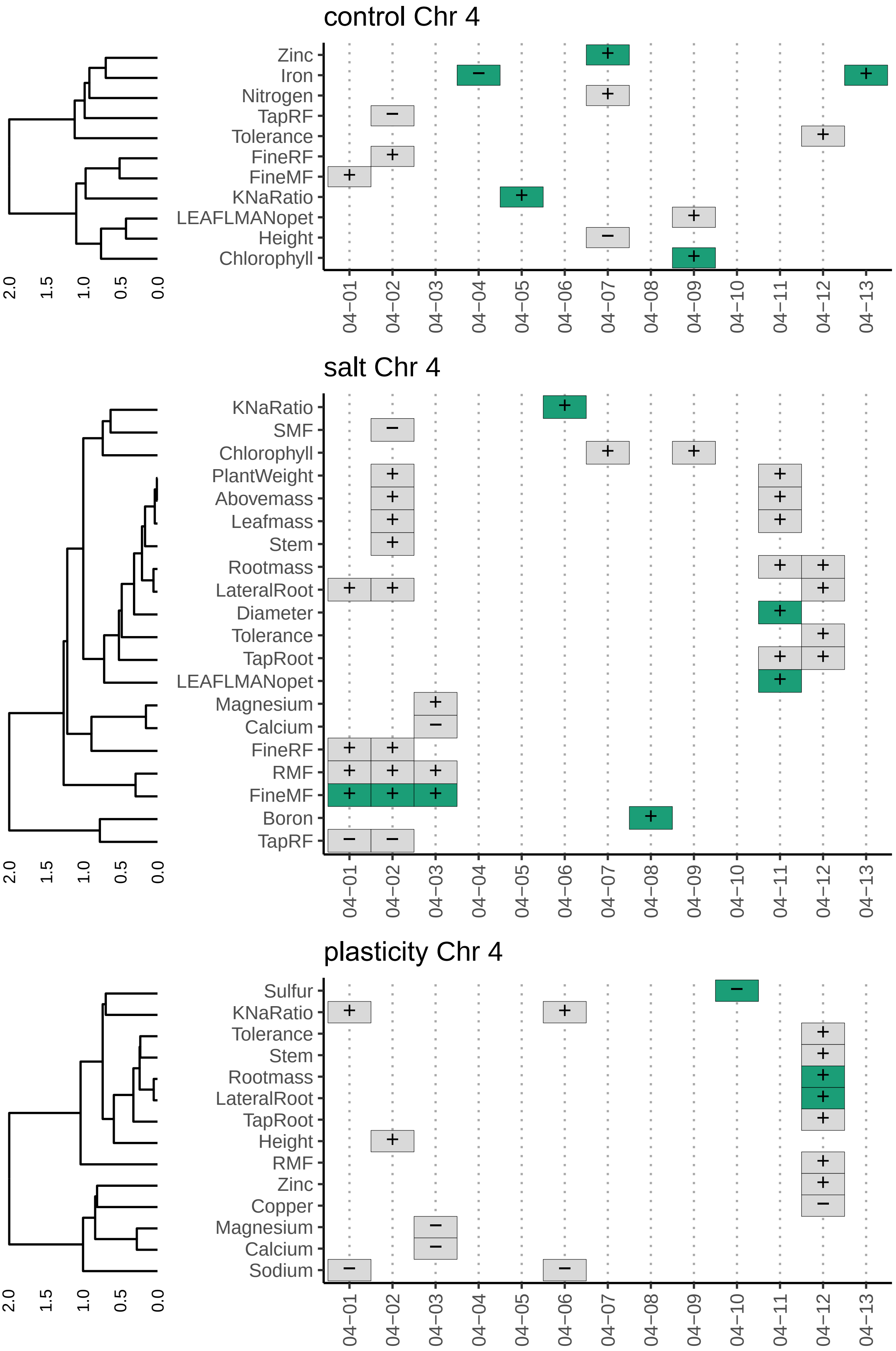
control Chr 4
Zinc
Iron
Nitrogen
TapRF
Tolerance
FineRF
FineMF
KNaRatio
LEAFLMANopet
Height
Chlorophyll
salt Chr 4
KNaRatio
SMF
Chlorophyll
PlantWeight
Abovemass
Leafmass
Stem
Rootmass
LateralRoot
Diameter
Tolerance
TapRoot
LEAFLMANopet
Magnesium
Calcium
FineRF
RMF
FineMF
Boron
TapRF
plasticity Chr 4
Sulfur
KNaRatio
Tolerance
Stem
Rootmass
LateralRoot
TapRoot
Height
RMF
Zinc
Copper
Magnesium
Calcium
Sodium
2.0 1.5 1.0 0.5 0.0
04−01 04−02 04−03 04−04 04−05 04−06 04−07 04−08 04−09 04−10 04−11 04−12 04−13

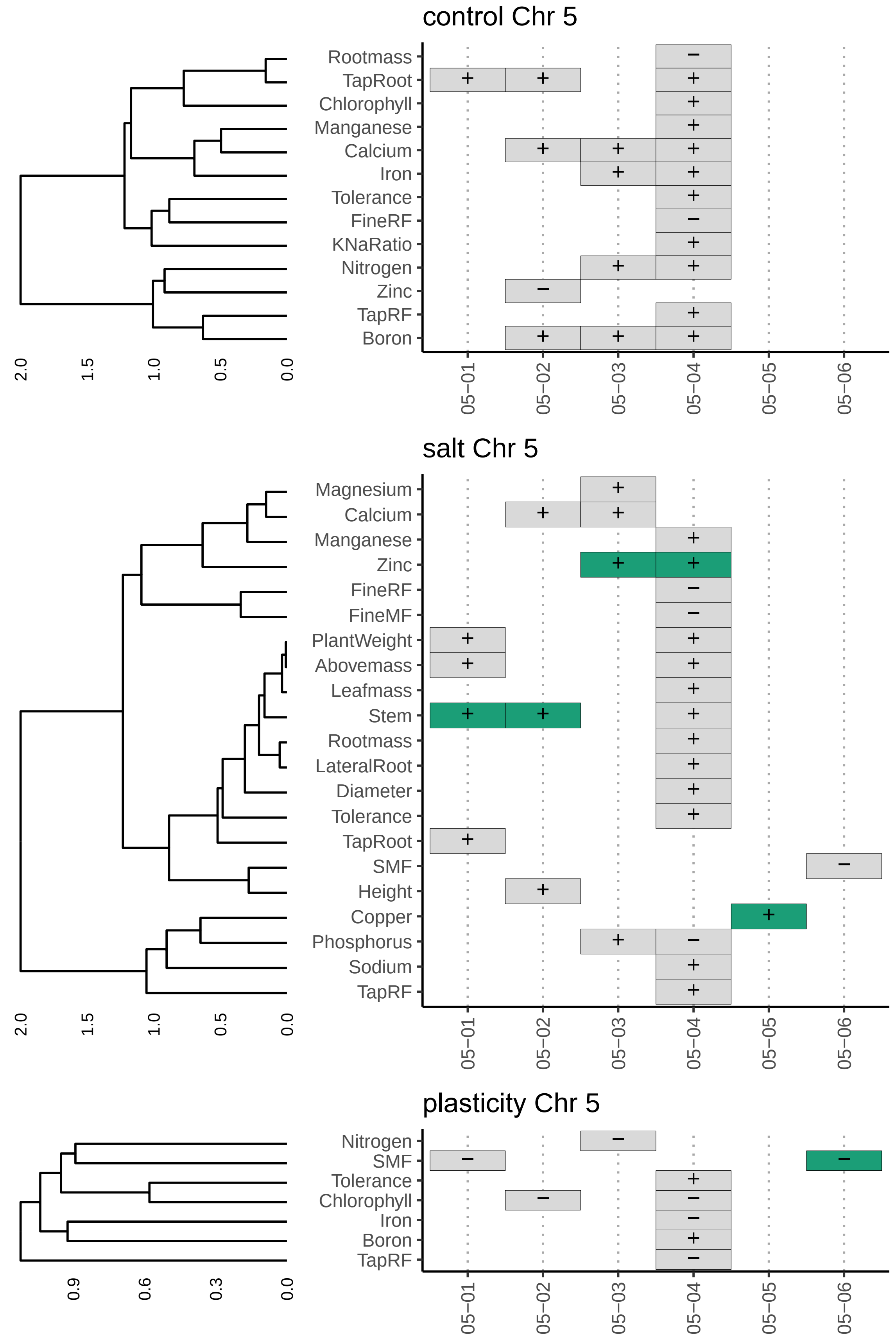

control Chr 5
Rootmass
TapRoot
Chlorophyll
Manganese
Calcium
Iron
Tolerance
FineRF
KNaRatio
Nitrogen
Zinc
TapRF
Boron
2.0
1.5
1.0
0.5
0.0
05−01
05−02
05−03
05−04
05−05
05−06
salt Chr 5
Magnesium
Calcium
Manganese
Zinc
FineRF
FineMF
PlantWeight
Abovemass
Leafmass
Stem
Rootmass
LateralRoot
Diameter
Tolerance
TapRoot
SMF
Height
Copper
Phosphorus
Sodium
TapRF
2.0
1.5
1.0
0.5
0.0
05−01
05−02
05−03
05−04
05−05
05−06
plasticity Chr 5
Nitrogen
SMF
Tolerance
Chlorophyll
Iron
Boron
TapRF
0.9
0.6
0.3
0.0
05−01
05−02
05−03
05−04
05−05
05−06

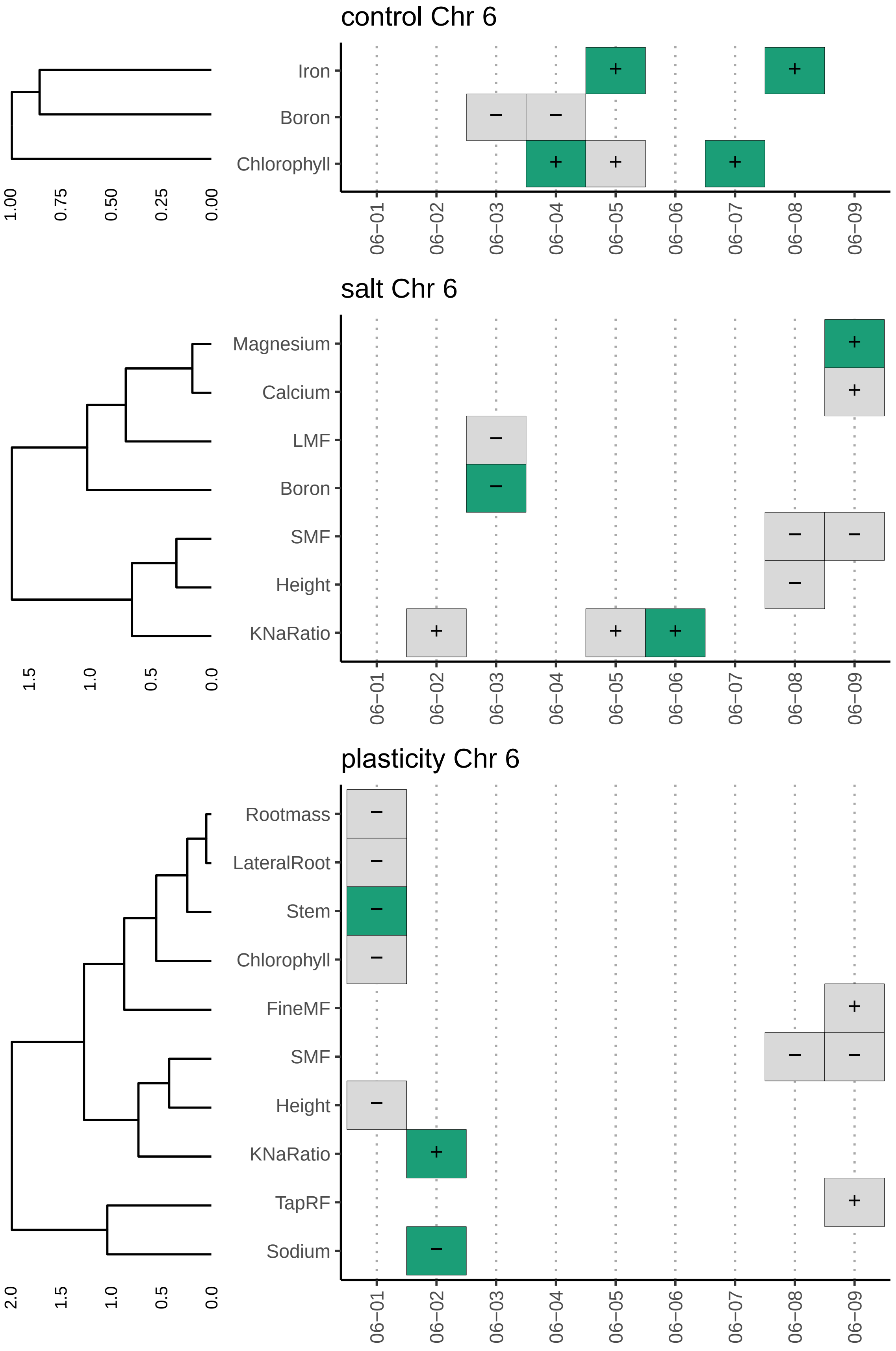

control Chr 6
Iron
Boron
Chlorophyll
1.00
0.75
0.50
0.25
0.00
06−01
06−02
06−03
06−04
06−05
06−06
06−07
06−08
06−09
salt Chr 6
Magnesium
Calcium
LMF
Boron
SMF
Height
KNaRatio
1.5
1.0
0.5
0.0
plasticity Chr 6
Rootmass
LateralRoot
Stem
Chlorophyll
FineMF
SMF
Height
KNaRatio
TapRF
Sodium
2.0
1.5
1.0
0.5
0.0

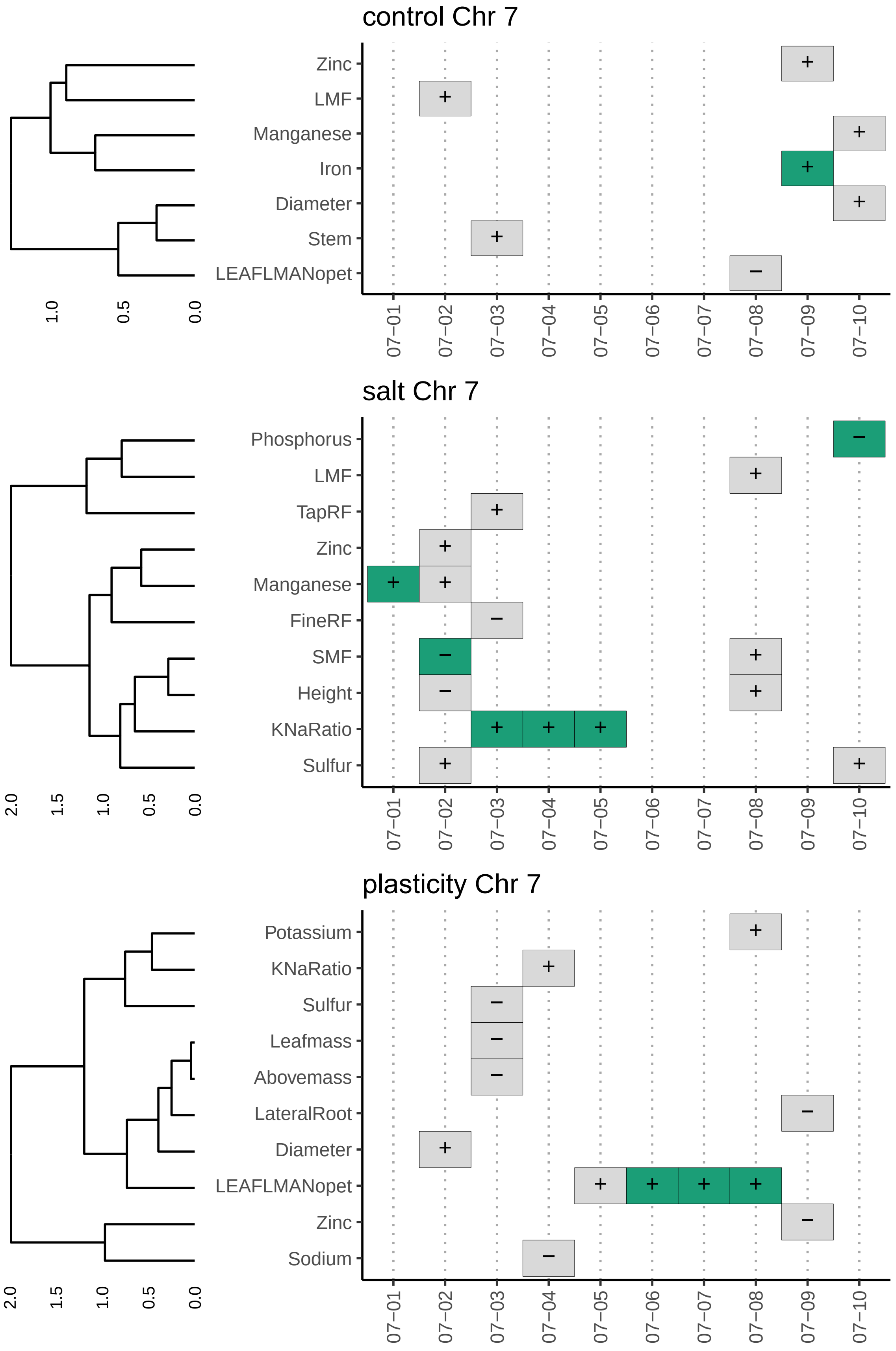

control Chr 7
Zinc
LMF
Manganese
Iron
Diameter
Stem
LEAFLMANopet
1.0
0.5
0.0
07−01
07−02
07−03
07−04
07−05
07−06
07−07
07−08
07−09
07−10
salt Chr 7
Phosphorus
LMF
TapRF
Zinc
Manganese
FineRF
SMF
Height
KNaRatio
Sulfur
2.0
1.5
1.0
0.5
0.0
plasticity Chr 7
Potassium
KNaRatio
Sulfur
Leafmass
Abovemass
LateralRoot
Diameter
LEAFLMANopet
Zinc
Sodium

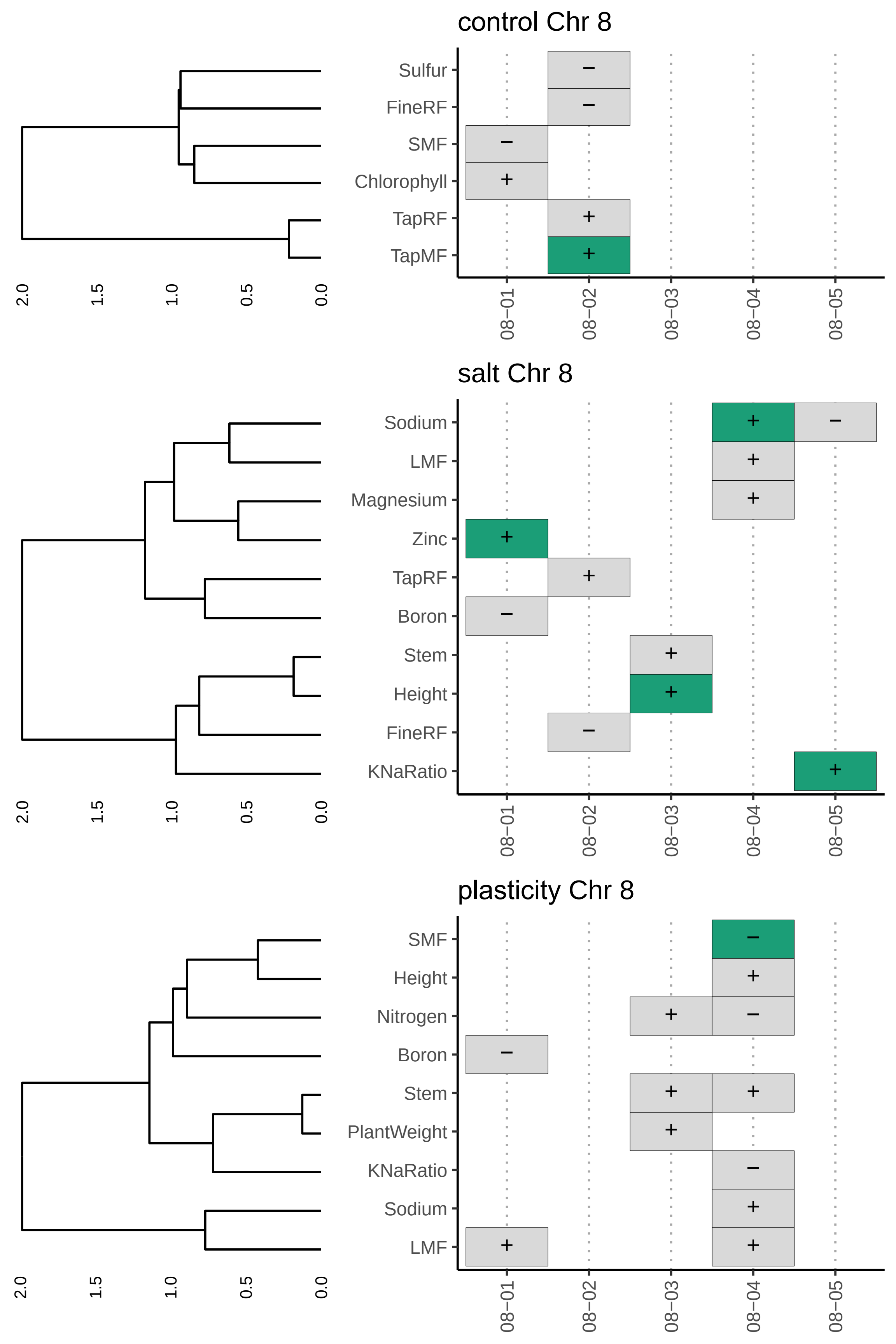

control Chr 8
Sulfur
FineRF
SMF
Chlorophyll
TapRF
TapMF
2.0
1.5
1.0
0.5
0.0
08−01
08−02
08−03
08−04
08−05
salt Chr 8
Sodium
LMF
Magnesium
Zinc
TapRF
Boron
Stem
Height
FineRF
KNaRatio
plasticity Chr 8
SMF
Height
Nitrogen
Boron
Stem
PlantWeight
KNaRatio
Sodium
LMF

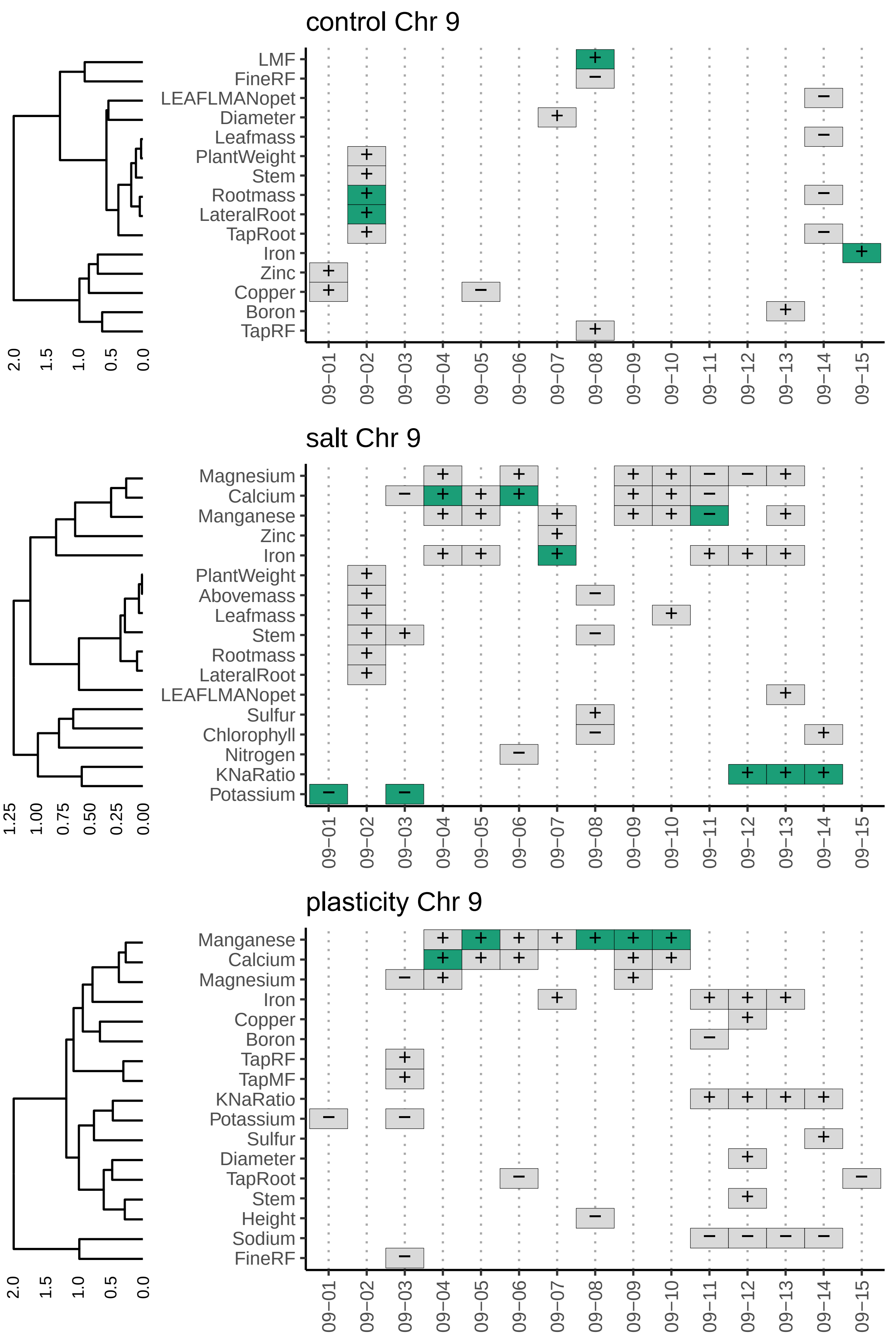

control Chr 9
LMF
FineRF
LEAFLMANopet
Diameter
Leafmass
PlantWeight
Stem
Rootmass
LateralRoot
TapRoot
Iron
Zinc
Copper
Boron
TapRF
2.0 1.5 1.0 0.5 0.0
09−01 09−02 09−03 09−04 09−05 09−06 09−07 09−08 09−09 09−10 09−11 09−12 09−13 09−14 09−15
salt Chr 9
Magnesium
Calcium
Manganese
Zinc
Iron
PlantWeight
Abovemass
Leafmass
Stem
Rootmass
LateralRoot
LEAFLMANopet
Sulfur
Chlorophyll
Nitrogen
KNaRatio
Potassium
1.25 1.00 0.75 0.50 0.25 0.00
09−01 09−02 09−03 09−04 09−05 09−06 09−07 09−08 09−09 09−10 09−11 09−12 09−13 09−14 09−15
plasticity Chr 9
Manganese
Calcium
Magnesium
Iron
Copper
Boron
TapRF
TapMF
KNaRatio
Potassium
Sulfur
Diameter
TapRoot
Stem
Height
Sodium
FineRF
2.0 1.5 1.0 0.5 0.0
09−01 09−02 09−03 09−04 09−05 09−06 09−07 09−08 09−09 09−10 09−11 09−12 09−13 09−14 09−15

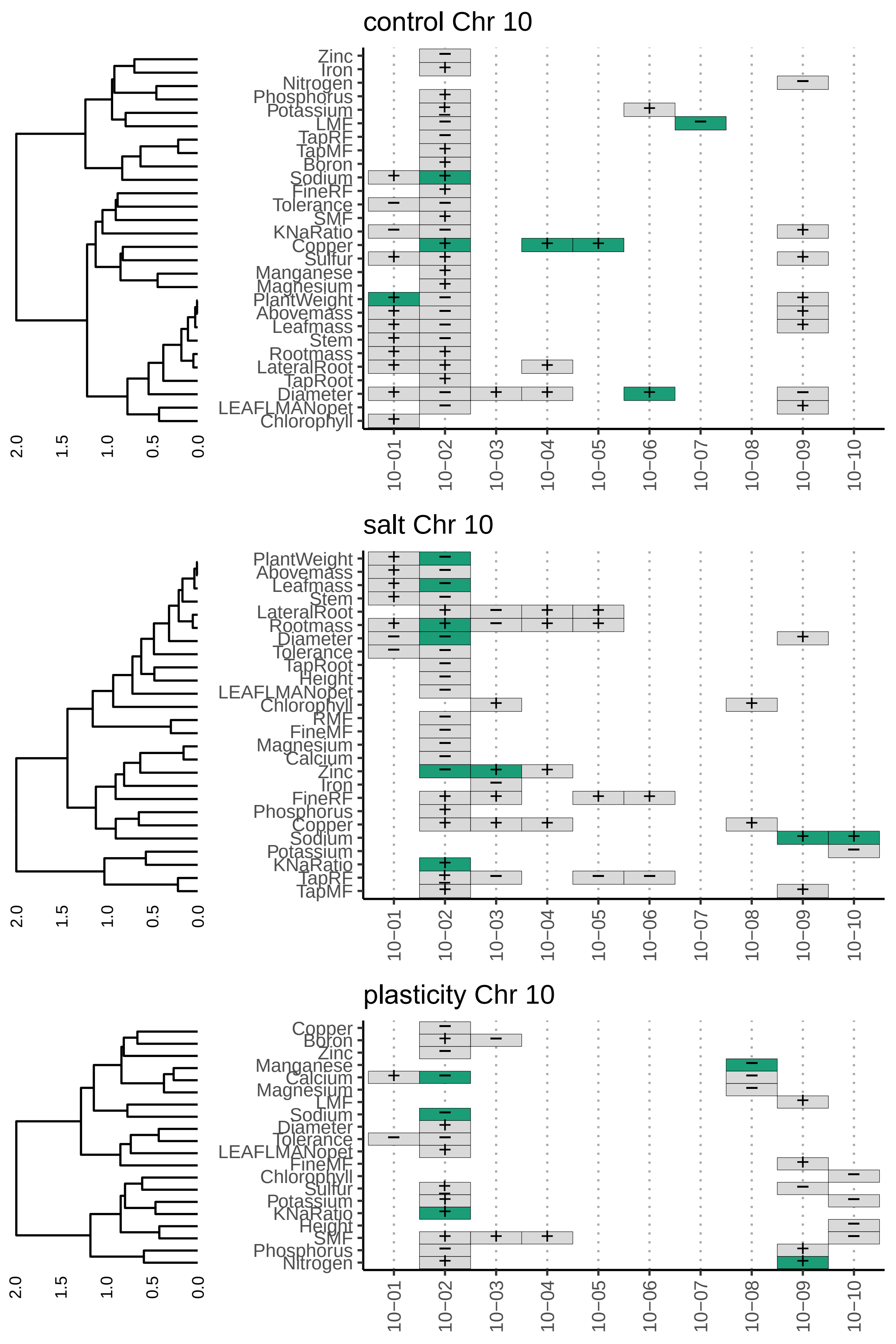

control Chr 10
Zinc
Iron
Nitrogen
Phosphorus
Potassium
LMF
TapRF
TapMF
Boron
Sodium
FineRF
Tolerance
SMF
KNaRatio
Copper
Sulfur
Manganese
Magnesium
PlantWeight
Abovemass
Leafmass
Stem
Rootmass
LateralRoot
TapRoot
Diameter
LEAFLMANopet
Chlorophyll
2.0 1.5 1.0 0.5 0.0
10−01 10−02 10−03 10−04 10−05 10−06 10−07 10−08 10−09 10−10
salt Chr 10
PlantWeight
Abovemass
Leafmass
Stem
LateralRoot
Rootmass
Diameter
Tolerance
TapRoot
Height
LEAFLMANopet
Chlorophyll
RMF
FineMF
Magnesium
Calcium
Zinc
Iron
FineRF
Phosphorus
Copper
Sodium
Potassium
KNaRatio
TapRF
TapMF
2.0 1.5 1.0 0.5 0.0
10−01 10−02 10−03 10−04 10−05 10−06 10−07 10−08 10−09 10−10
plasticity Chr 10
Copper
Boron
Zinc
Manganese
Calcium
Magnesium
LMF
Sodium
Diameter
Tolerance
LEAFLMANopet
FineMF
Chlorophyll
Sulfur
Potassium
KNaRatio
Height
SMF
Phosphorus
Nitrogen
2.0 1.5 1.0 0.5 0.0
10−01 10−02 10−03 10−04 10−05 10−06 10−07 10−08 10−09 10−10

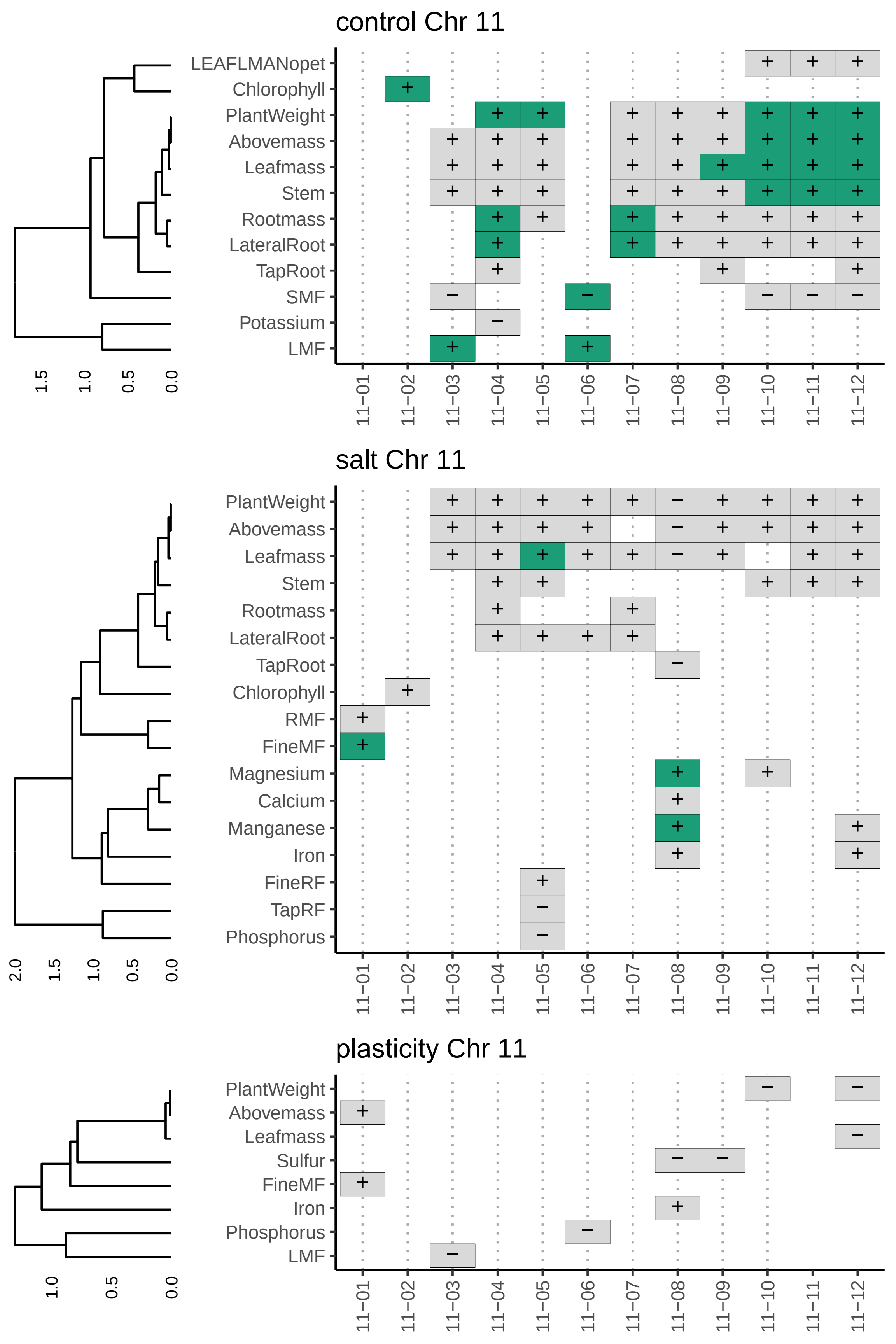

control Chr 11
LEAFLMANopet
Chlorophyll
PlantWeight
Abovemass
Leafmass
Stem
Rootmass
LateralRoot
TapRoot
SMF
Potassium
LMF
1.5
1.0
0.5
0.0
11−01
11−02
11−03
11−04
11−05
11−06
11−07
11−08
11−09
11−10
11−11
11−12
salt Chr 11
PlantWeight
Abovemass
Leafmass
Stem
Rootmass
LateralRoot
TapRoot
Chlorophyll
RMF
FineMF
Magnesium
Calcium
Manganese
Iron
FineRF
TapRF
Phosphorus
2.0
1.5
1.0
0.5
0.0
plasticity Chr 11
PlantWeight
Abovemass
Leafmass
Sulfur
FineMF
Iron
Phosphorus
LMF
1.0
0.5
0.0

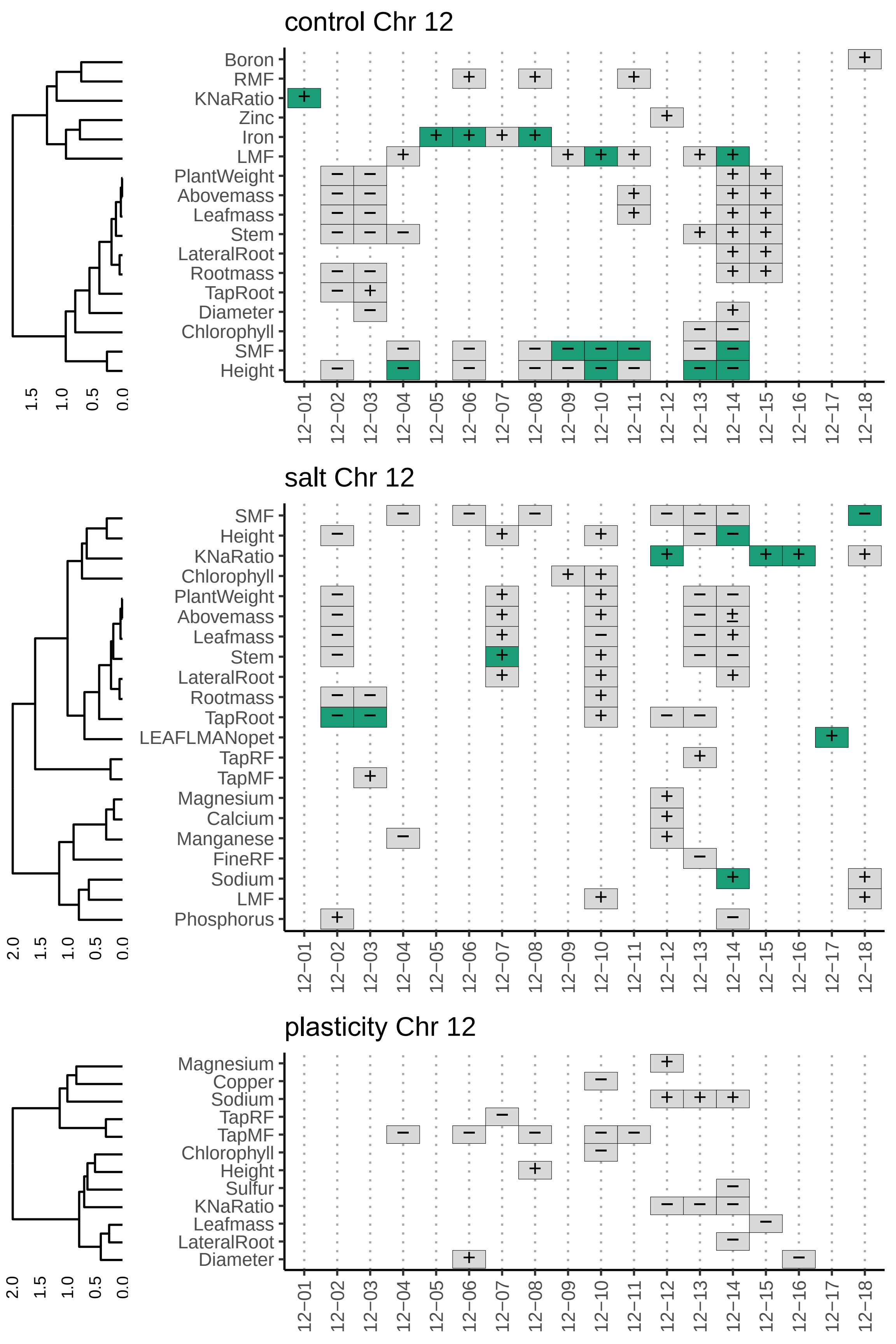

control Chr 12
Boron
RMF
KNaRatio
Zinc
Iron
LMF
PlantWeight
Abovemass
Leafmass
Stem
LateralRoot
Rootmass
TapRoot
Diameter
Chlorophyll
SMF
Height
1.5 1.0 0.5 0.0
12−01 12−02 12−03 12−04 12−05 12−06 12−07 12−08 12−09 12−10 12−11 12−12 12−13 12−14 12−15 12−16 12−17 12−18
salt Chr 12
SMF
Height
KNaRatio
Chlorophyll
PlantWeight
Abovemass
Leafmass
Stem
LateralRoot
Rootmass
TapRoot
LEAFLMANopet
TapRF
TapMF
Magnesium
Calcium
Manganese
FineRF
Sodium
LMF
Phosphorus
2.0 1.5 1.0 0.5 0.0
12−01 12−02 12−03 12−04 12−05 12−06 12−07 12−08 12−09 12−10 12−11 12−12 12−13 12−14 12−15 12−16 12−17 12−18
plasticity Chr 12
Magnesium
Copper
Sodium
TapRF
TapMF
Chlorophyll
Height
Sulfur
KNaRatio
Leafmass
LateralRoot
Diameter
2.0 1.5 1.0 0.5 0.0
12−01 12−02 12−03 12−04 12−05 12−06 12−07 12−08 12−09 12−10 12−11 12−12 12−13 12−14 12−15 12−16 12−17 12−18

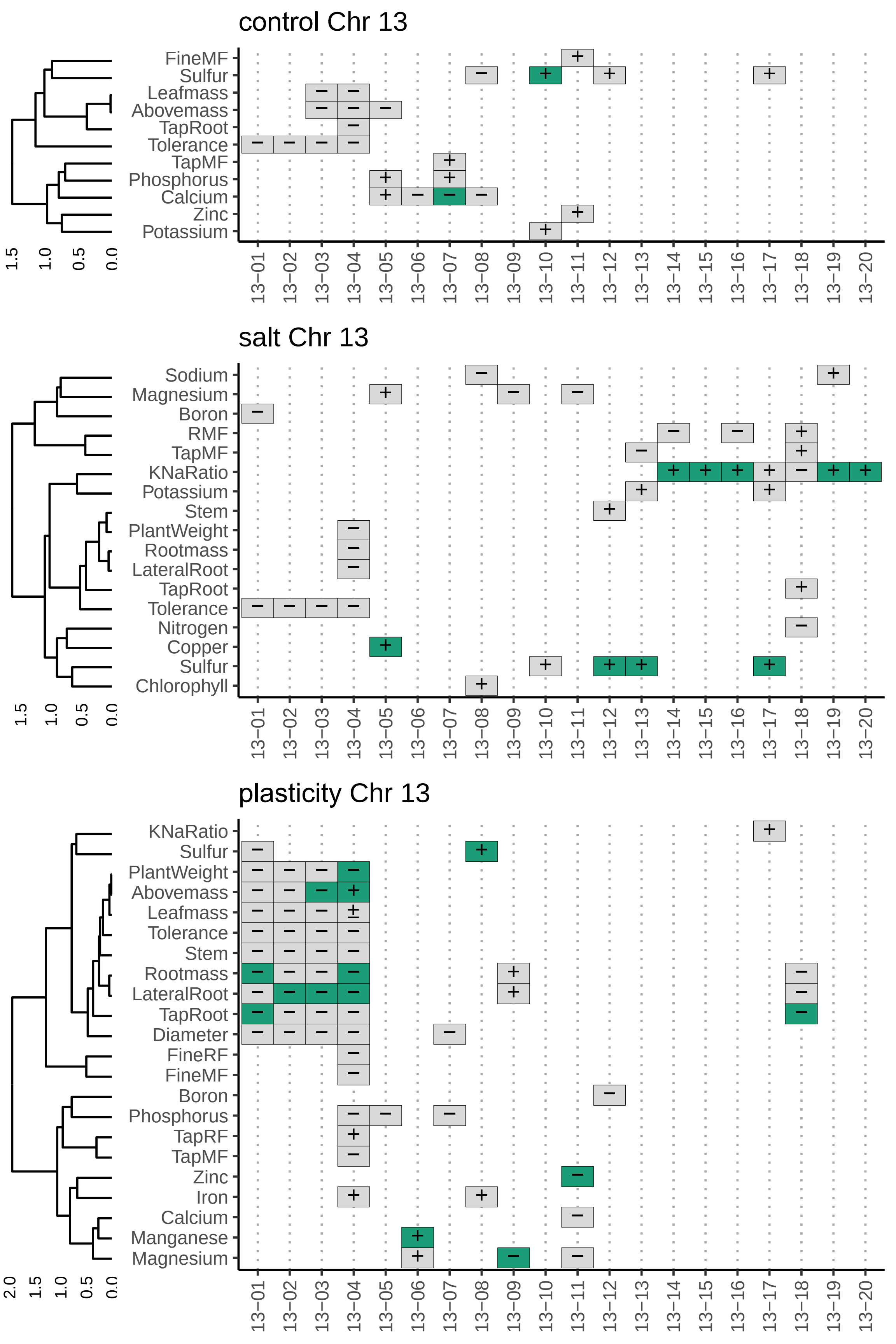
control Chr 13
FineMF
Sulfur
Leafmass
Abovemass
TapRoot
Tolerance
TapMF
Phosphorus
Calcium
Zinc
Potassium
salt Chr 13
Sodium
Magnesium
Boron
RMF
TapMF
KNaRatio
Potassium
Stem
PlantWeight
Rootmass
LateralRoot
TapRoot
Tolerance
Nitrogen
Copper
Sulfur
Chlorophyll
plasticity Chr 13
KNaRatio
Sulfur
PlantWeight
Abovemass
Leafmass
Tolerance
Stem
Rootmass
LateralRoot
TapRoot
Diameter
FineRF
FineMF
Boron
Phosphorus
TapRF
TapMF
Zinc
Iron
Calcium
Manganese
Magnesium
2.0 1.5 1.0 0.5 0.0
13−01 13−02 13−03 13−04 13−05 13−06 13−07 13−08 13−09 13−10 13−11 13−12 13−13 13−14 13−15 13−16 13−17 13−18 13−19 13−20

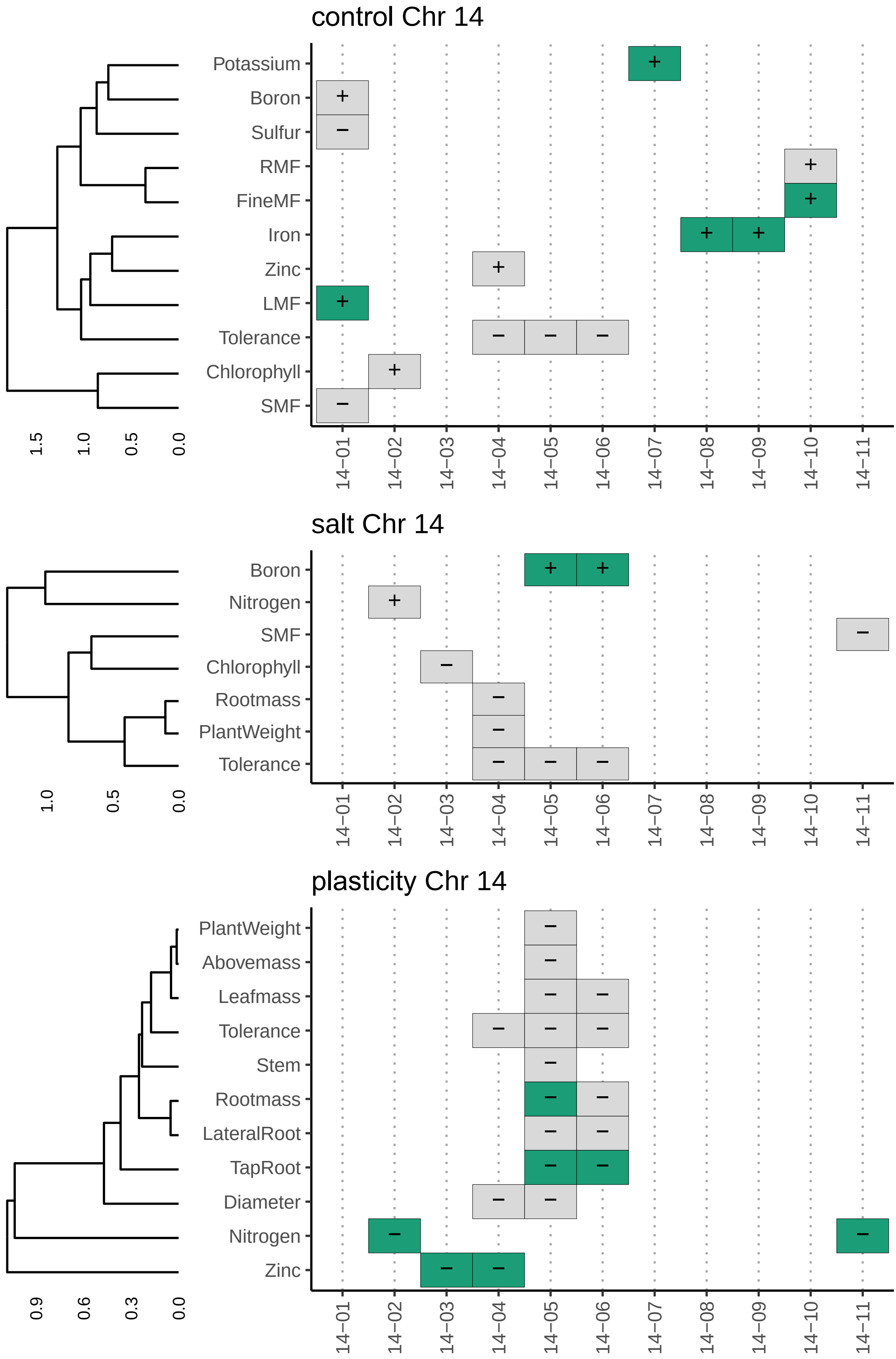
control Chr 14
Potassium
Boron
Sulfur
RMF
FineMF
Iron
Zinc
LMF
Tolerance
Chlorophyll
SMF
1.5
1.0
0.5
0.0
14−01
14−02
14−03
14−04
14−05
14−06
14−07
14−08
14−09
14−10
14−11
salt Chr 14
Boron
Nitrogen
SMF
Chlorophyll
Rootmass
PlantWeight
Tolerance
1.0
0.5
0.0
plasticity Chr 14
PlantWeight
Abovemass
Leafmass
Tolerance
Stem
Rootmass
LateralRoot
TapRoot
Diameter
Nitrogen
Zinc
0.9
0.6
0.3
0.0

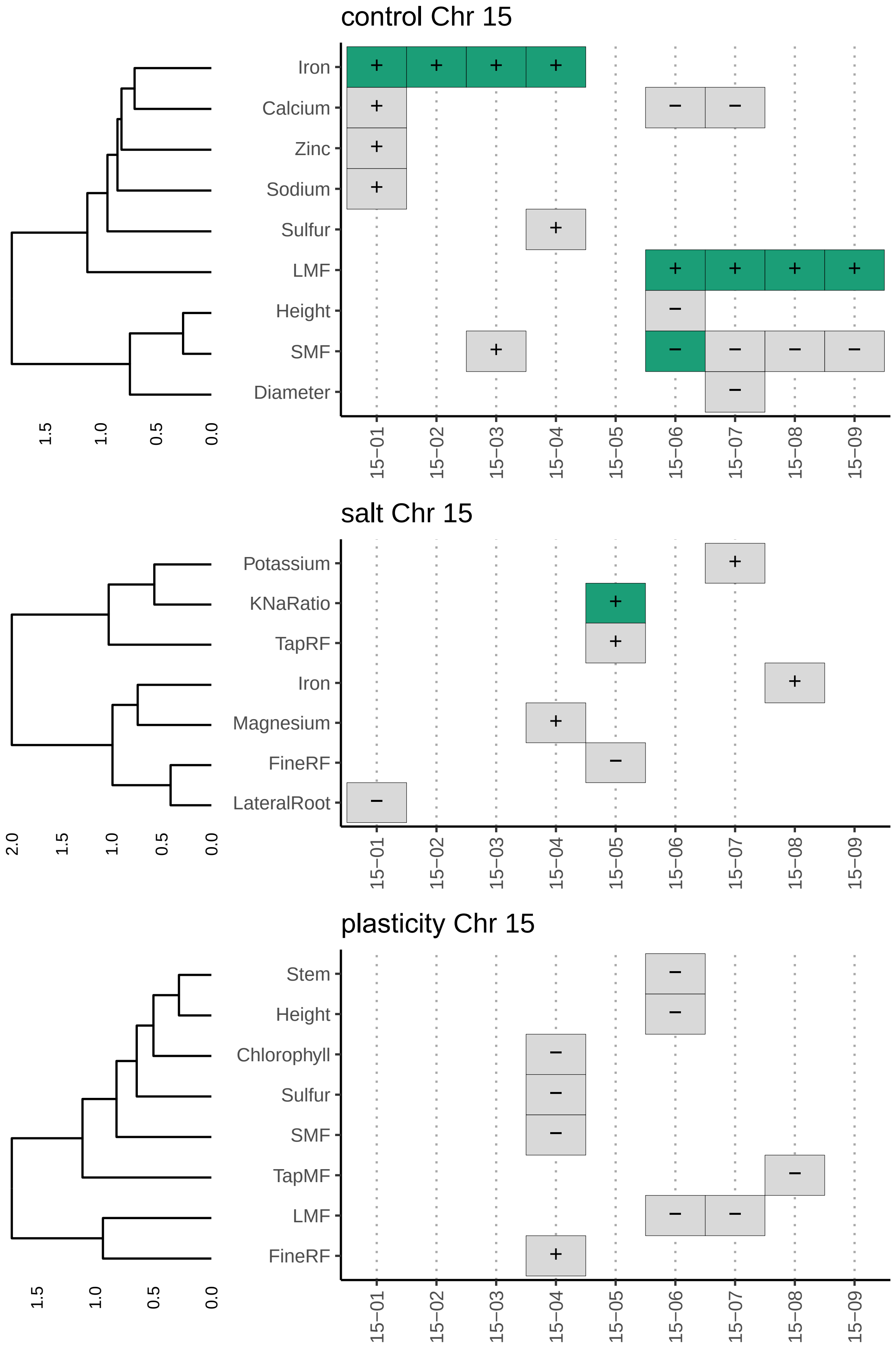

control Chr 15
Iron
Calcium
Zinc
Sodium
Sulfur
LMF
Height
SMF
Diameter
1.5
1.0
0.5
0.0
15−01
15−02
15−03
15−04
15−05
15−06
15−07
15−08
15−09
salt Chr 15
Potassium
KNaRatio
TapRF
Iron
Magnesium
FineRF
LateralRoot
2.0
1.5
1.0
0.5
0.0
15−01
15−02
15−03
15−04
15−05
15−06
15−07
15−08
15−09
plasticity Chr 15
Stem
Height
Chlorophyll
Sulfur
SMF
TapMF
LMF
FineRF
1.5
1.0
0.5
0.0
15−01
15−02
15−03
15−04
15−05
15−06
15−07
15−08
15−09

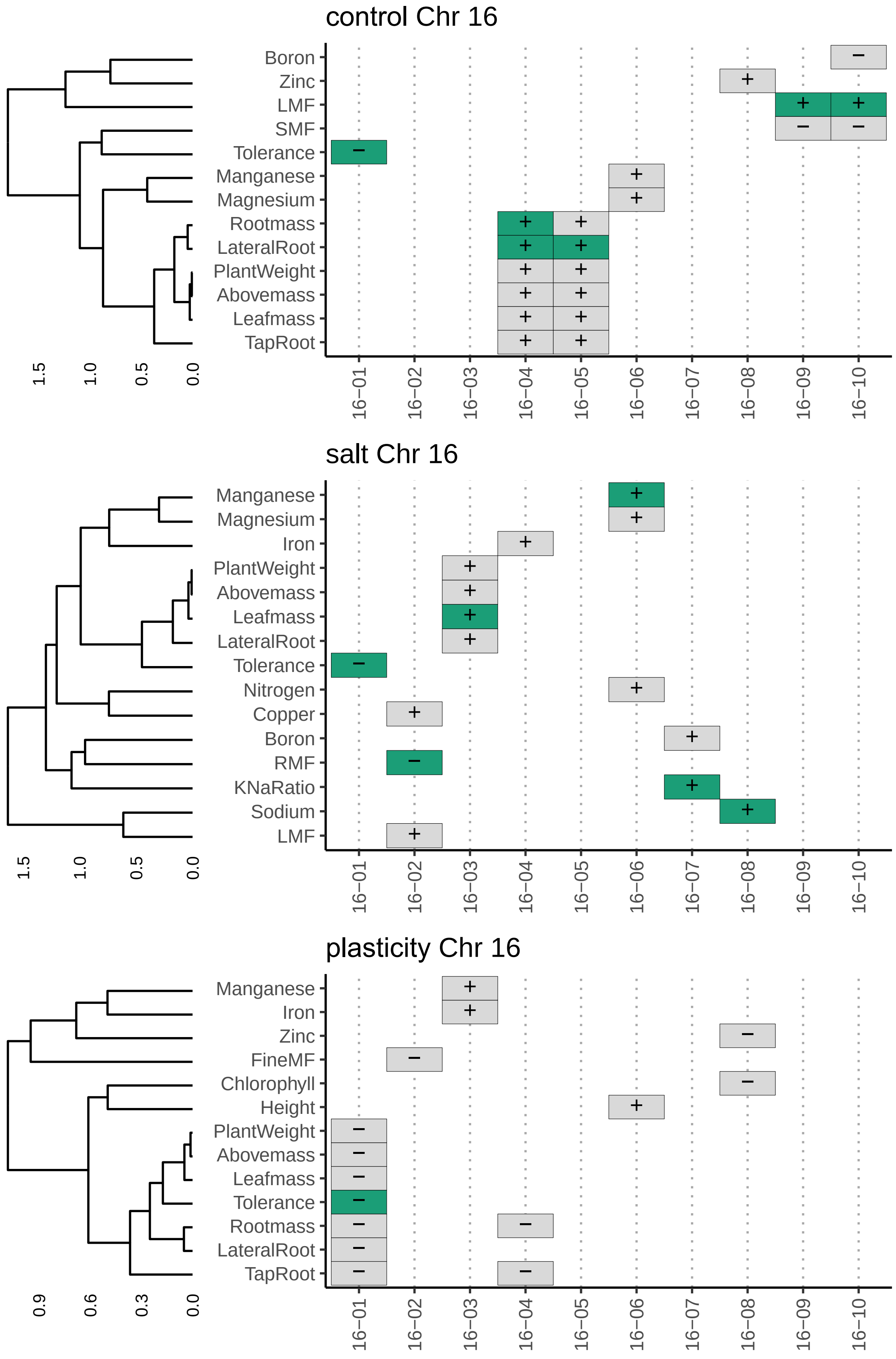
control Chr 16
Boron
Zinc
LMF
SMF
Tolerance
Manganese
Magnesium
Rootmass
LateralRoot
PlantWeight
Abovemass
Leafmass
TapRoot
1.5
1.0
0.5
0.0
16−01
16−02
16−03
16−04
16−05
16−06
16−07
16−08
16−09
16−10
salt Chr 16
Manganese
Magnesium
Iron
PlantWeight
Abovemass
Leafmass
LateralRoot
Tolerance
Nitrogen
Copper
Boron
RMF
KNaRatio
Sodium
LMF
1.5
1.0
0.5
0.0
16−01
16−02
16−03
16−04
16−05
16−06
16−07
16−08
16−09
16−10
plasticity Chr 16
Manganese
Iron
Zinc
FineMF
Chlorophyll
Height
PlantWeight
Abovemass
Leafmass
Tolerance
Rootmass
LateralRoot
TapRoot
0.9
0.6
0.3
0.0
16−01
16−02
16−03
16−04
16−05
16−06
16−07
16−08
16−09
16−10

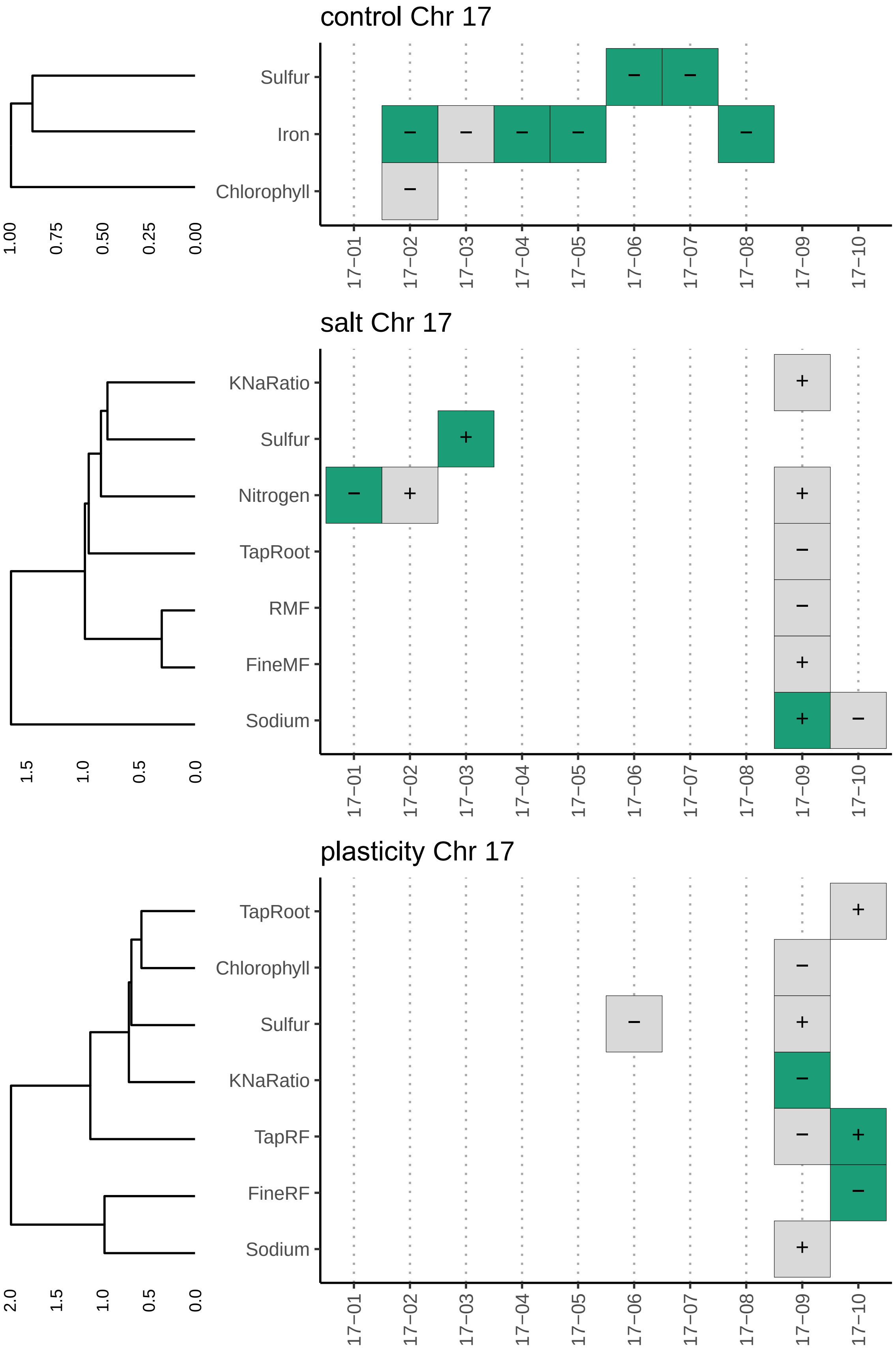

control Chr 17
Sulfur
Iron
Chlorophyll
1.00
0.75
0.50
0.25
0.00
17−01
17−02
17−03
17−04
17−05
17−06
17−07
17−08
17−09
17−10
salt Chr 17
KNaRatio
Sulfur
Nitrogen
TapRoot
RMF
FineMF
Sodium
1.5
1.0
0.5
0.0
plasticity Chr 17
TapRoot
Chlorophyll
Sulfur
KNaRatio
TapRF
FineRF
Sodium
2.0
1.5
1.0
0.5
0.0

**Figure S7. Co-localization of traits across environments.** If a linkage block contained a significant SNP for a trait in one environment, or trait plasticity across environments, the alternate values were assessed to determine if there were significant (green) or suggestive (i.e., top 0.1% of all SNPs; grey) associations across environments. Plus/minus symbols in the tiles indicate the sign of $\beta$ (effect of minor allele on trait). Linkage blocks are ordered by chromosome with black lines separating them. Sizes of the linkage blocks vary both in physical size as well as gene number.

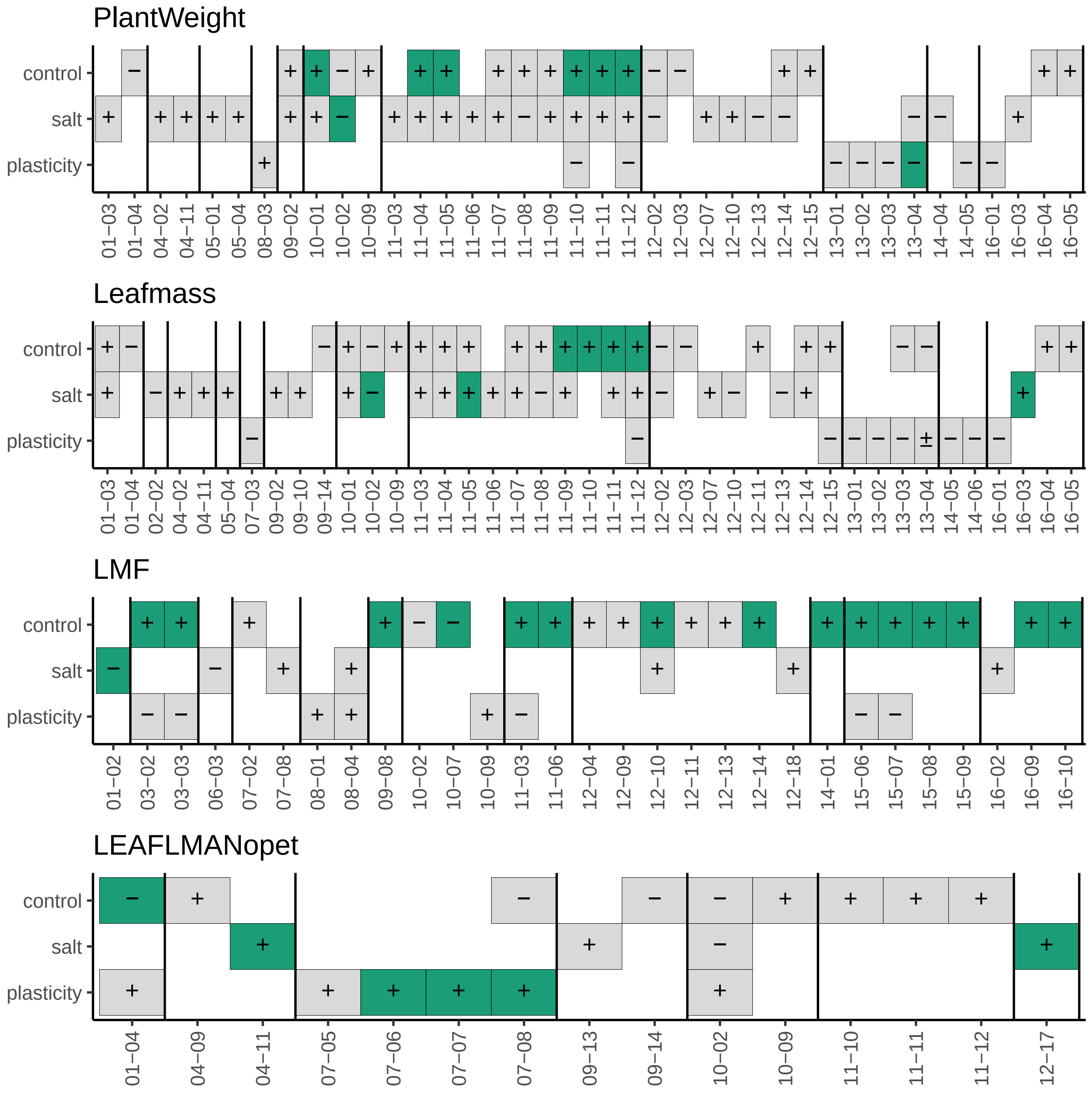
PlantWeight
control
salt
plasticity
01−03 01−04 04−02 04−11 05−01 05−04 08−03 09−02 10−01 10−02 10−09 11−03 11−04 11−05 11−06 11−07 11−08 11−09 11−10 11−11 11−12 12−02 12−03 12−07 12−10 12−13 12−14 12−15 13−01 13−02 13−03 13−04 14−04 14−05 16−01 16−03 16−04 16−05
Leafmass
control
salt
plasticity
01−03 01−04 02−02 04−02 04−11 05−04 07−03 09−02 09−10 09−14 10−01 10−02 10−09 11−03 11−04 11−05 11−06 11−07 11−08 11−09 11−10 11−11 11−12 12−02 12−03 12−07 12−10 12−11 12−13 12−14 12−15 13−01 13−02 13−03 13−04 14−05 14−06 16−01 16−03 16−04 16−05
LMF
control
salt
plasticity
01−02 03−02 03−03 06−03 07−02 07−08 08−01 08−04 09−08 10−02 10−07 10−09 11−03 11−06 12−04 12−09 12−10 12−11 12−13 12−14 12−18 14−01 15−06 15−07 15−08 15−09 16−02 16−09 16−10
LEAFLMANopet
control
salt
plasticity
01−04 04−09 04−11 07−05 07−06 07−07 07−08 09−13 09−14 10−02 10−09 11−10 11−11 11−12 12−17

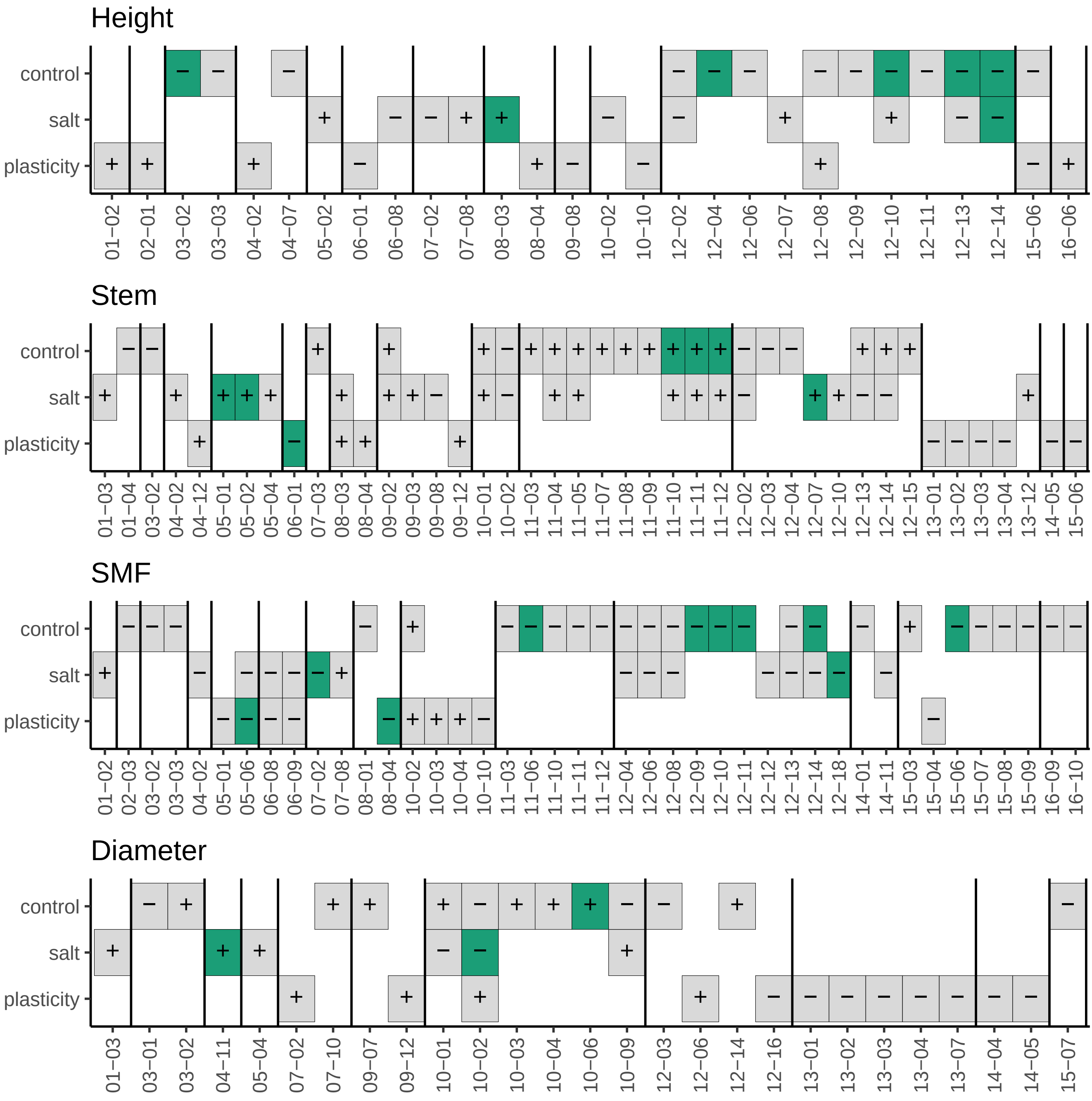
Height
Stem
SMF
Diameter
control
salt
plasticity

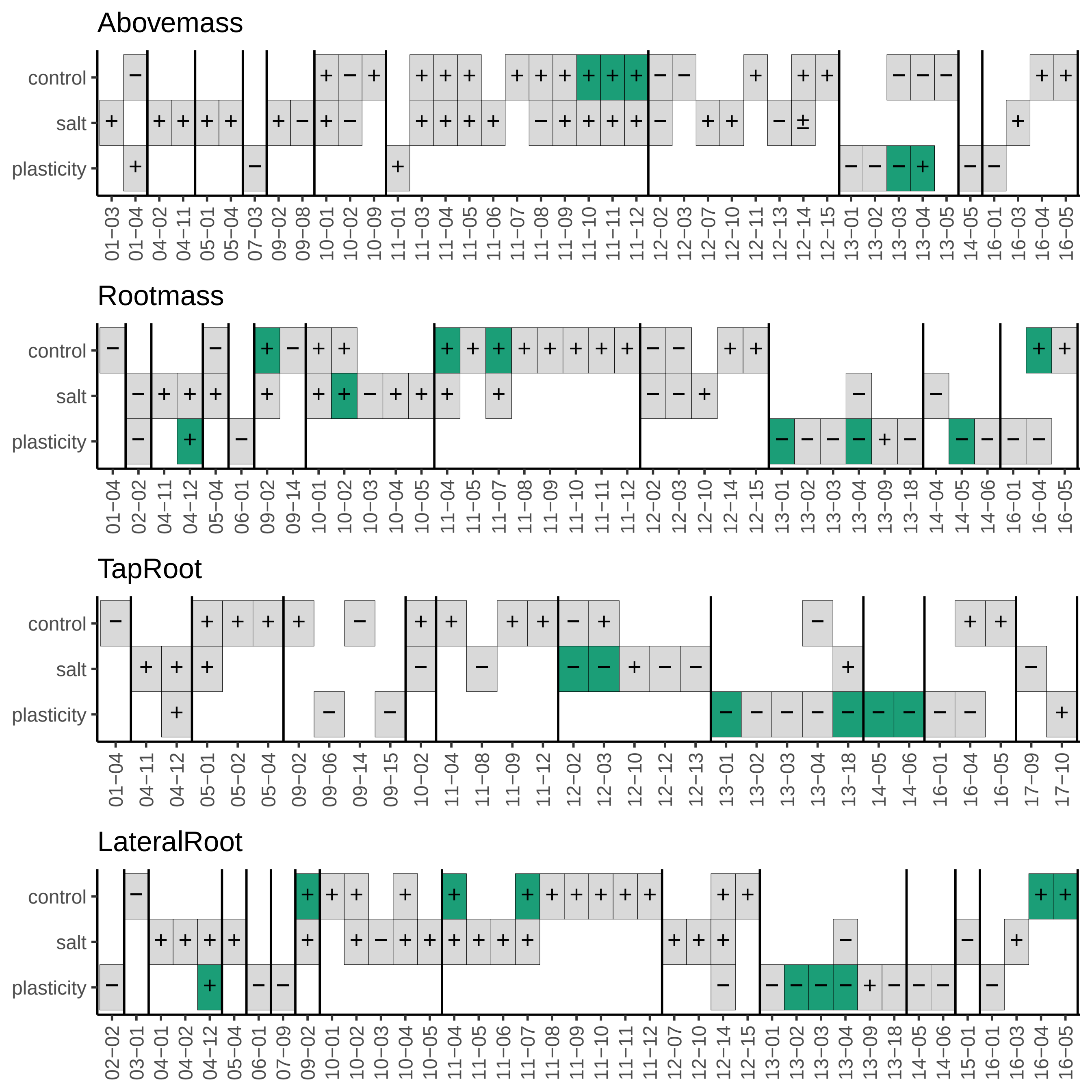
Abovemass
control
salt
plasticity
01−03 01−04 04−02 04−11 05−01 05−04 07−03 09−02 09−08 10−01 10−02 10−09 11−01 11−03 11−04 11−05 11−06 11−07 11−08 11−09 11−10 11−11 11−12 12−02 12−03 12−07 12−10 12−11 12−13 12−14 12−15 13−01 13−02 13−03 13−04 13−05 14−05 16−01 16−03 16−04 16−05
Rootmass
control
salt
plasticity
01−04 02−02 04−11 04−12 05−04 06−01 09−02 09−14 10−01 10−02 10−03 10−04 10−05 11−04 11−05 11−07 11−08 11−09 11−10 11−11 11−12 12−02 12−03 12−10 12−14 12−15 13−01 13−02 13−03 13−04 13−09 13−18 14−04 14−05 14−06 16−01 16−04 16−05
TapRoot
control
salt
plasticity
01−04 04−11 04−12 05−01 05−02 05−04 09−02 09−06 09−14 09−15 10−02 11−04 11−08 11−09 11−12 12−02 12−03 12−10 12−12 12−13 13−01 13−02 13−03 13−04 13−18 14−05 14−06 16−01 16−04 16−05 17−09 17−10
LateralRoot
control
salt
plasticity
02−02 03−01 04−01 04−02 04−12 05−04 06−01 07−09 09−02 10−01 10−02 10−03 10−04 10−05 11−04 11−05 11−06 11−07 11−08 11−09 11−10 11−11 11−12 12−07 12−10 12−14 12−15 13−01 13−02 13−03 13−04 13−09 13−18 14−05 14−06 15−01 16−01 16−03 16−04 16−05

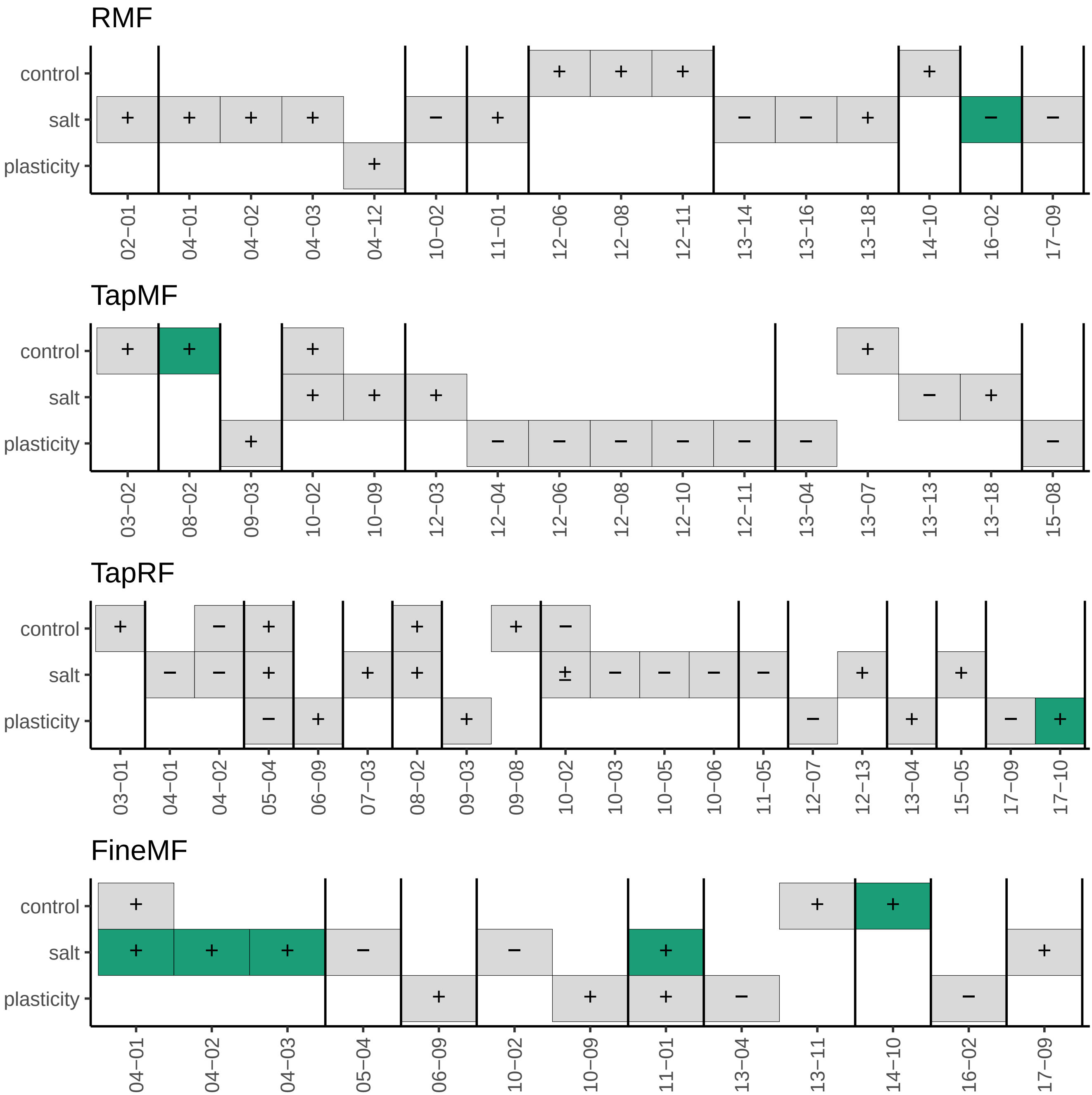
RMF
control
salt
plasticity
02−01 04−01 04−02 04−03 04−12 10−02 11−01 12−06 12−08 12−11 13−14 13−16 13−18 14−10 16−02 17−09
TapMF
control
salt
plasticity
03−02 08−02 09−03 10−02 10−09 12−03 12−04 12−06 12−08 12−10 12−11 13−04 13−07 13−13 13−18 15−08
TapRF
control
salt
plasticity
03−01 04−01 04−02 05−04 06−09 07−03 08−02 09−03 09−08 10−02 10−03 10−05 10−06 11−05 12−07 12−13 13−04 15−05 17−09 17−10
FineMF
control
salt
plasticity
04−01 04−02 04−03 05−04 06−09 10−02 10−09 11−01 13−04 13−11 14−10 16−02 17−09

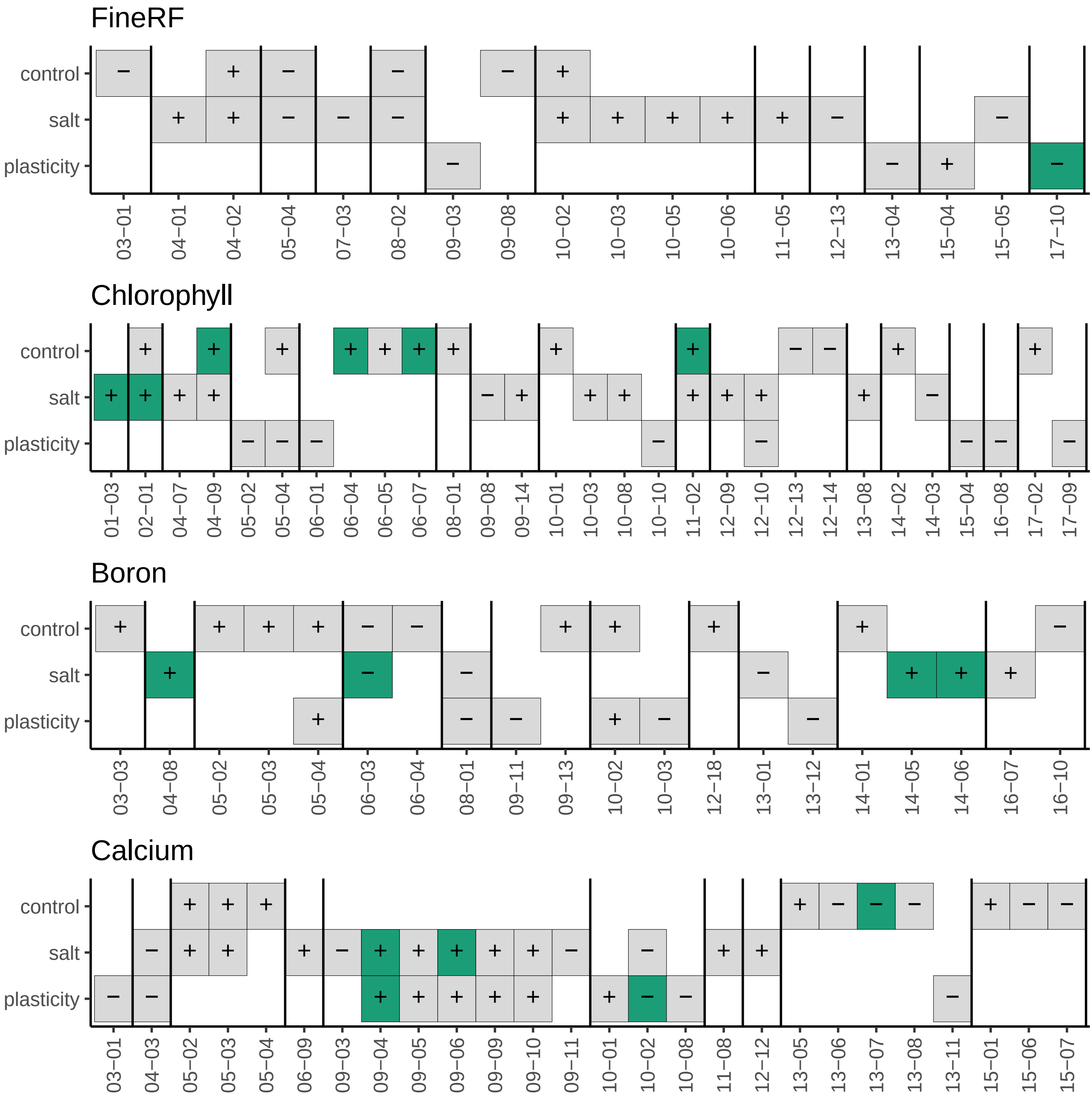

FineRF
control
salt
plasticity
03−01
04−01
04−02
05−04
07−03
08−02
09−03
09−08
10−02
10−03
10−05
10−06
11−05
12−13
13−04
15−04
15−05
17−10
Chlorophyll
control
salt
plasticity
01−03
02−01
04−07
04−09
05−02
05−04
06−01
06−04
06−05
06−07
08−01
09−08
09−14
10−01
10−03
10−08
10−10
11−02
12−09
12−10
12−13
12−14
13−08
14−02
14−03
15−04
16−08
17−02
17−09
Boron
control
salt
plasticity
03−03
04−08
05−02
05−03
05−04
06−03
06−04
08−01
09−11
09−13
10−02
10−03
12−18
13−01
13−12
14−01
14−05
14−06
16−07
16−10
Calcium
control
salt
plasticity
03−01
04−03
05−02
05−03
05−04
06−09
09−03
09−04
09−05
09−06
09−09
09−10
09−11
10−01
10−02
10−08
11−08
12−12
13−05
13−06
13−07
13−08
13−11
15−01
15−06
15−07

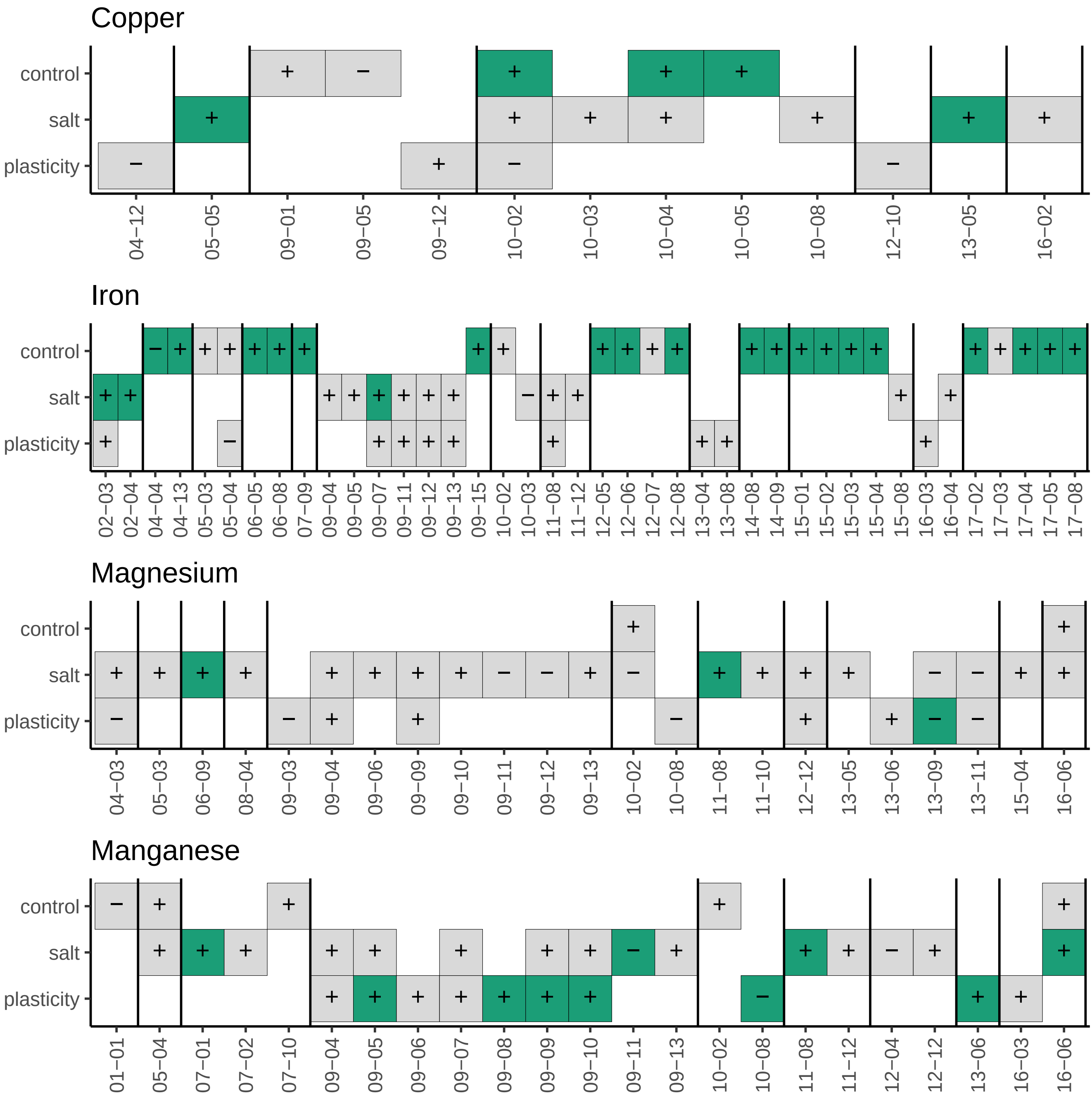

Copper
Iron
Magnesium
Manganese
control
salt
plasticity

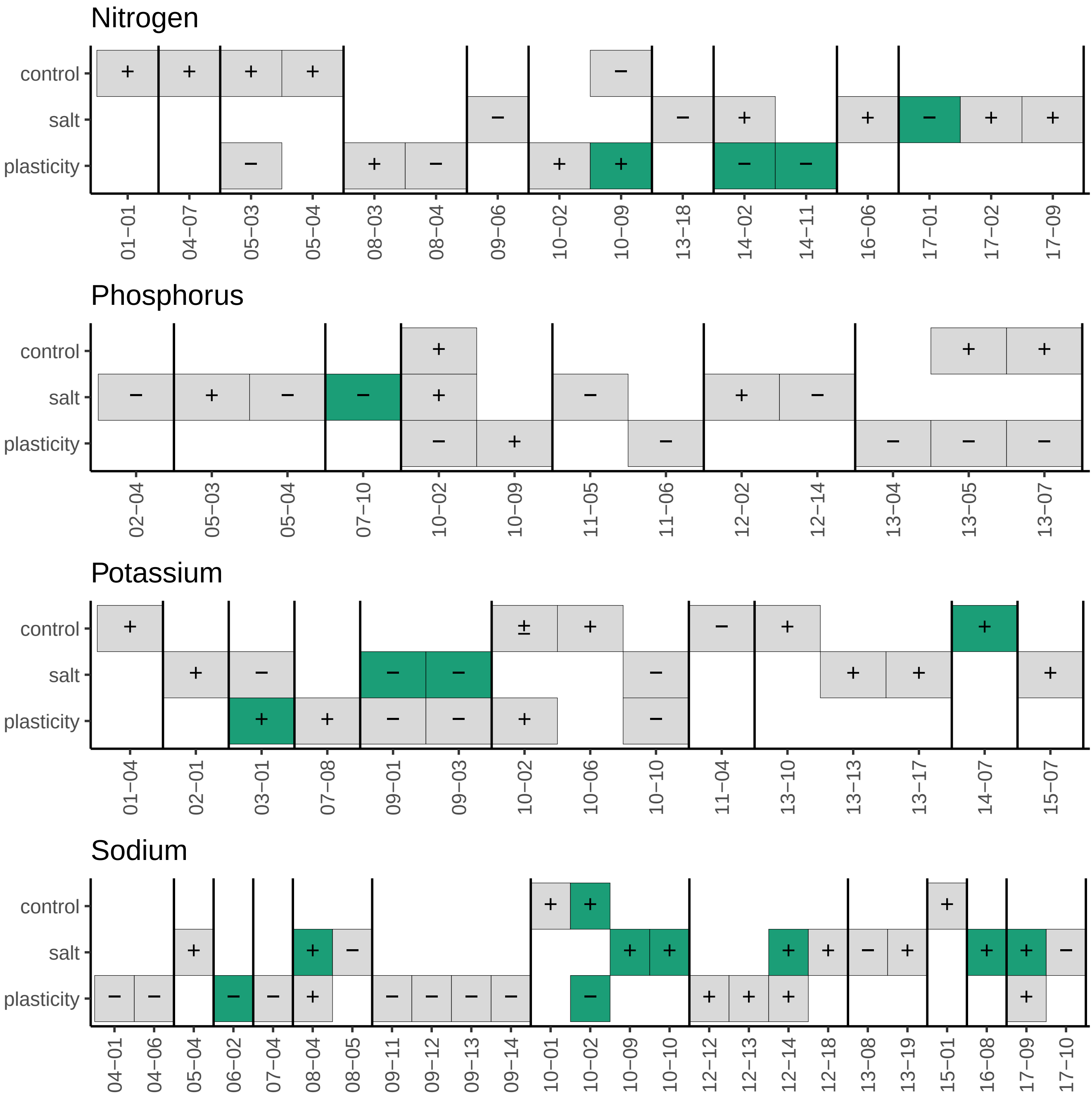
Nitrogen
control
salt
plasticity
01−01
04−07
05−03
05−04
08−03
08−04
09−06
10−02
10−09
13−18
14−02
14−11
16−06
17−01
17−02
17−09
Phosphorus
02−04
05−03
05−04
07−10
10−02
10−09
11−05
11−06
12−02
12−14
13−04
13−05
13−07
Potassium
01−04
02−01
03−01
07−08
09−01
09−03
10−02
10−06
10−10
11−04
13−10
13−13
13−17
14−07
15−07
Sodium
04−01
04−06
05−04
06−02
07−04
08−04
08−05
09−11
09−12
09−13
09−14
10−01
10−02
10−09
10−10
12−12
12−13
12−14
12−18
13−08
13−19
15−01
16−08
17−09
17−10

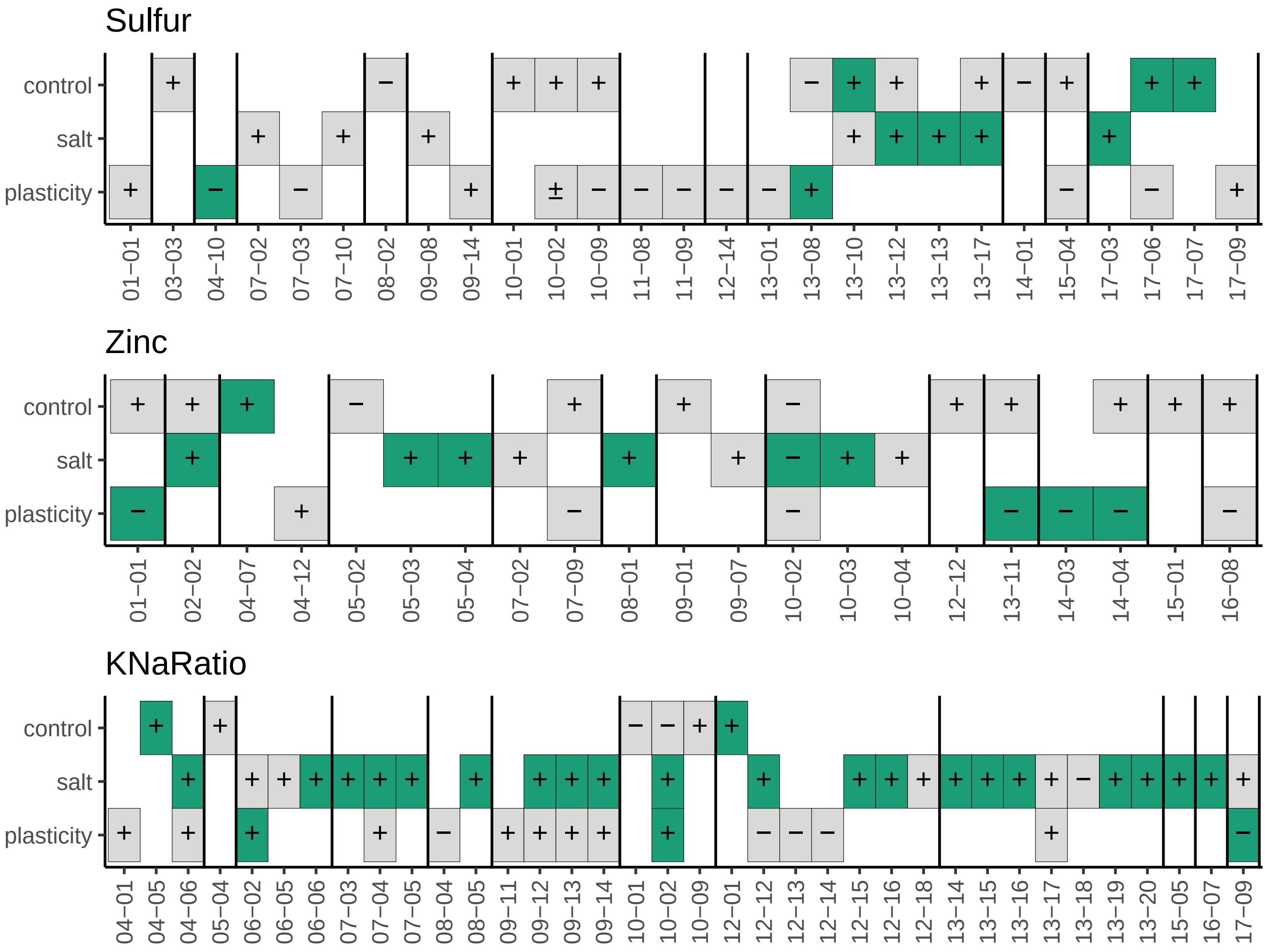

Sulfur
control
salt
plasticity
01−01
03−03
04−10
07−02
07−03
07−10
08−02
09−08
09−14
10−01
10−02
10−09
11−08
11−09
12−14
13−01
13−08
13−10
13−12
13−13
13−17
14−01
15−04
17−03
17−06
17−07
17−09
Zinc
control
salt
plasticity
01−01
02−02
04−07
04−12
05−02
05−03
05−04
07−02
07−09
08−01
09−01
09−07
10−02
10−03
10−04
12−12
13−11
14−03
14−04
15−01
16−08
KNaRatio
control
salt
plasticity
04−01
04−05
04−06
05−04
06−02
06−05
06−06
07−03
07−04
07−05
08−04
08−05
09−11
09−12
09−13
09−14
10−01
10−02
10−09
12−01
12−12
12−13
12−14
12−15
12−16
12−18
13−14
13−15
13−16
13−17
13−18
13−19
13−20
15−05
16−07
17−09

**Figure S8. Number of genes per significant region**. Points indicate the number of genes present in the 189 linkage blocks identified from GWAS of all traits/trait plasticities. Note the log scale on the y-axis to aid in visualization.

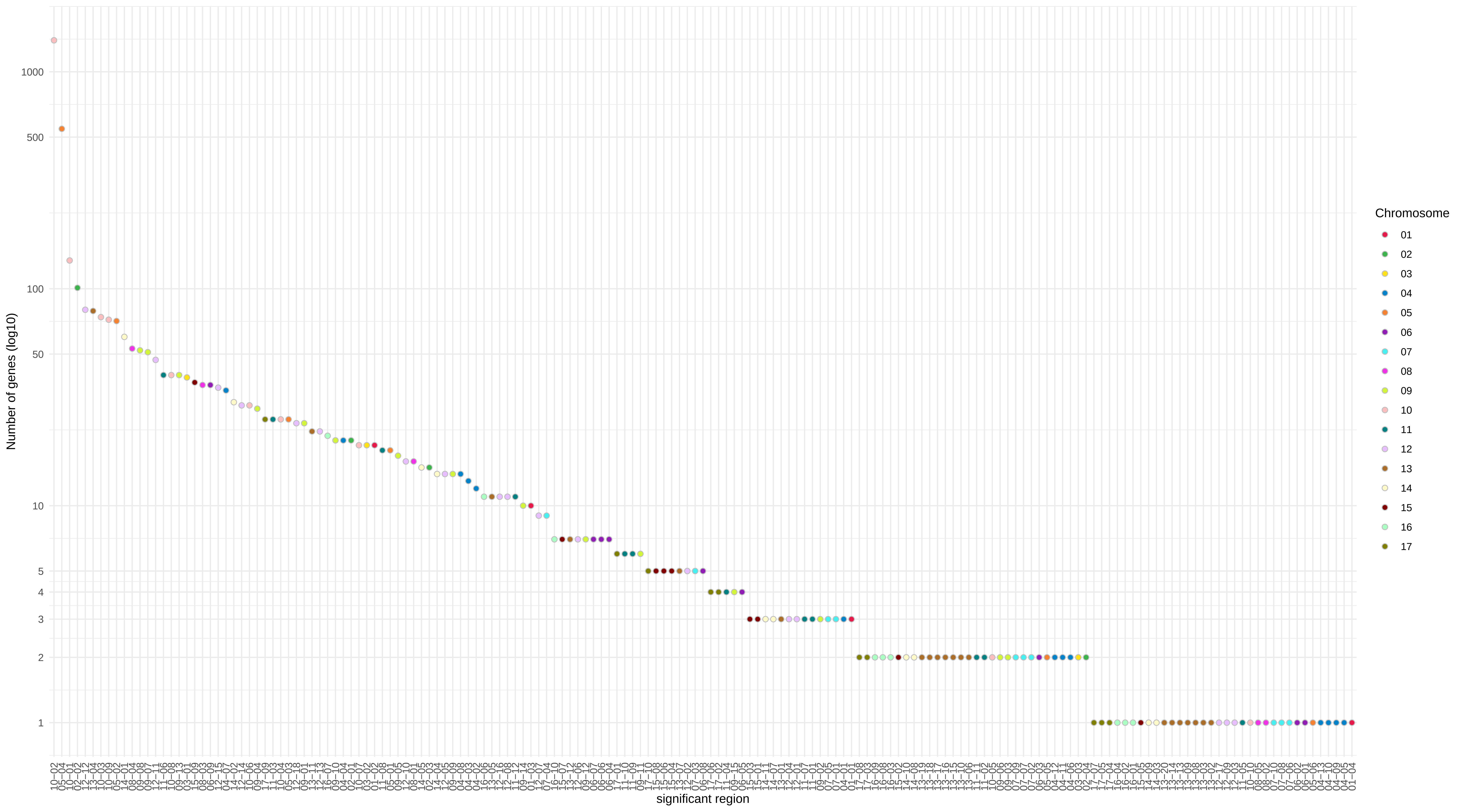

Number of genes (log10)
1000
500
100
50
10
5
4
3
2
1
significant region
Chromosome
